# Conservation of copy number profiles during engraftment and passaging of patient-derived cancer xenografts

**DOI:** 10.1101/861393

**Authors:** Xing Yi Woo, Jessica Giordano, Anuj Srivastava, Zi-Ming Zhao, Michael W. Lloyd, Roebi de Bruijn, Yun-Suhk Suh, Rajesh Patidar, Li Chen, Sandra Scherer, Matthew Bailey, Chieh-Hsiang Yang, Emilio Cortes-Sanchez, Yuanxin Xi, Jing Wang, Jayamanna Wickramasinghe, Andrew V. Kossenkov, Vito Rebecca, Hua Sun, R. Jay Mashl, Sherri Davies, Ryan Jeon, Christian Frech, Jelena Randjelovic, Jacqueline Rosains, Francesco Galimi, Andrea Bertotti, Adam Lafferty, Alice C. O’Farrell, Elodie Modave, Diether Lambrechts, Petra ter Brugge, Violeta Serra, Elisabetta Marangoni, Rania El Botty, Hyunsoo Kim, Jong-Il Kim, Han-Kwang Yang, Charles Lee, Dennis A. Dean, Brandi Davis-Dusenbery, Yvonne A. Evrard, James H. Doroshow, Alana L. Welm, Bryan E. Welm, Michael T. Lewis, Bingliang Fang, Jack A. Roth, Funda Meric-Bernstam, Meenhard Herlyn, Michael Davies, Li Ding, Shunqiang Li, Ramaswamy Govindan, Claudio Isella, Jeffrey A. Moscow, Livio Trusolino, Annette T. Byrne, Jos Jonkers, Carol J. Bult, Enzo Medico, Jeffrey H. Chuang, PDXNET consortium, EurOPDX consortium

**Affiliations:** The Jackson Laboratory for Genomic Medicine, Farmington, CT; Department of Oncology, University of Torino, Candiolo, Italy; Candiolo Cancer Institute, FPO-IRCCS, Candiolo, Italy; The Jackson Laboratory, Bar Harbor, ME; Netherlands Cancer Institute, Amsterdam, Netherlands; College of Medicine, Seoul National University, South Korea; Frederick National Laboratory for Cancer Research, Frederick, MD; University of Utah Huntsman Cancer Institute, Salt Lake City, UT; Department of Human Genetics, University of Utah, Salt Lake City, UT; The University of Texas M.D. Anderson Cancer Center, Houston, TX; The Wistar Institute, Philadelphia, PA; Washington University School of Medicine in St. Louis, St. Louis, MO; Seven Bridges Genomics, Inc., Cambridge, Charlestown, MA; Department of Physiology and Medical Physics, Centre for Systems Medicine, Royal College of Surgeons in Ireland, Dublin, Ireland; Center for Cancer Biology, VIB, Leuven, Belgium; Laboratory of Translational Genetics, Department of Human Genetics, KU Leuven, Leuven, Belgium; Vall d’Hebron Institute of Oncology, Barcelona, Spain; Institut Curie, Paris, France; Division of Cancer Treatment and Diagnosis, National Cancer Institute, Bethesda, MD; Lester and Sue Smith Breast Center, Baylor College of Medicine, Houston, TX; Investigational Drug Branch, National Cancer Institute, Bethesda, MD; Center to Reduce Cancer Health Disparities, National Cancer Institute, Bethesda, MD; Hamon Center For Therapeutic Oncology, UT Southwestern Medical Center, Dallas; Abramson Cancer Center, University of Pennsylvania, Philadelphia, PA; Department of Pathology and Laboratory Medicine, Hospital of the University of Pennsylvania, Philadelphia, PA; University of California Davis, Sacramento, CA; Baylor College of Medicine, Houston, TX; Manchester Cancer Research Centre, University of Manchester, UK; University of Basel, University Hospital Basel, Switzerland; University of Glasgow, Institute of Cancer Sciences, UK; Cancer Research UK Cambridge Institute, Cambridge Cancer Centre, UK; Catalan Institute of Oncology, Bellvitge Biomedical Research Institute, L’Hopistalet de Llobregat, Barcelona, Spain; European Bioinformatics Institute, European Molecular Biology Laboratory, Wellcome Trust Genome Campus, Hinxton, Cambridge, UK; Institute of Computer Science, MazarykUniversity, Brno, Czech Republic; Cornell University, Weill Cornell Medical College, New York, NY, United States; NorLux Neuro-Oncology Laboratory, Department of Oncology, Luxembourg Institute of Health, Luxembourg, Luxembourg; University Medical Centre Groningen, Groningen, The Netherlands; Czech Center for Phenogenomics, Institute of Molecular Genetics, Vestec, Czech Republic; TRACE PDX platform, Katholieke Universiteit Leuven, Leuven, Belgium; European Institute of Oncology, Milan, Italy; Oslo University Hospital, Oslo, Norway; seeding science SPRL, Limelette, Belgium

## Abstract

Patient-derived xenografts (PDXs) are resected human tumors engrafted into mice for preclinical studies and therapeutic testing. It has been proposed that the mouse host affects tumor evolution during PDX engraftment and propagation, impacting the accuracy of PDX modeling of human cancer. Here we exhaustively analyze copy number alterations (CNAs) in 1451 PDX and matched patient tumor (PT) samples from 509 PDX models. CNA inferences based on DNA sequencing and microarray data displayed substantially higher resolution and dynamic range than gene expression-based inferences, and they also showed strong CNA conservation from PTs through late-passage PDXs. CNA recurrence analysis of 130 colorectal and breast PT/PDX-early/PDX-late trios confirmed high-resolution CNA retention. We observed no significant enrichment of cancer-related genes in PDX-specific CNAs across models. Moreover, CNA differences between patient and PDX tumors were comparable to variations in multi-region samples within patients. Our study demonstrates the lack of systematic copy number evolution driven by the PDX mouse host.

## MAIN

A variety of models of human cancer have been used to study basic biological processes and predict responses to treatment. For example, mouse models with genetically engineered mutations in oncogenes and tumor suppressor genes have clarified the genetic and molecular basis of tumor initiation and progression^1,2^, though responses sometimes differ between human and mouse^3^. Cell lines have also been widely used to study cancer cells, but they lack the heterogeneity and microenvironment of *in vivo* tumors and have shown limitations for predicting clinical response^4^. Human tumors engrafted into transplant-compliant recipient mice (patient-derived xenografts, PDX) have advantages over prior systems for preclinical drug efficacy studies because they allow researchers to directly study human cells and tissues *in vivo*^5–8^. Comparisons of genome characteristics and histopathology of primary tumors and xenografts of human breast cancer^9–13^, ovarian cancer^14^, colorectal cancer^15^ and lung cancer^16–18^, have demonstrated that the biological properties of patient-derived tumors are largely preserved in xenografts. A growing body of literature supports their use in cancer drug discovery and development^19–21^.

A caveat to PDX models is that intratumoral evolution can occur during engraftment and passaging^11,22–25^. Such evolution could potentially modify treatment response of PDXs with respect to the patient tumors^23,26,27^, particularly if the evolution were to systematically alter cancer-related genes. This issue is related to multi-region comparisons of patient tumors^28–31^, for which local mutational and immune infiltration variations have suggested differential phenotypes among multi-region samples^32^. However, it remains unclear how therapies should be designed with respect to this variation. Comparing patient tumor-PDX evolution to the multi-region variations within the patient tumor would clarify the importance of primary-PDX divergence for treatment.

Recently, Ben-David et al.^26^ reported extensive PDX copy number divergence from the patient tumor of origin and across passages, based mainly on large-scale assessment of CNA profiles inferred from gene expression microarray data, which allowed analysis of aberrations at the scale of chromosomal arms. They raised concerns about genetic evolution in PDXs as a consequence of mouse-specific selective pressures, which could impact the capacity of PDXs for faithful modeling of patient treatment response. Such results contrast with reports that have observed genomic fidelity of PDX models with respect to the originating patient tumors and from early to late passages by direct DNA measurements (DNA sequencing or SNP arrays) in several dozen PDX models^9,10,33^.

Here we resolve these contradicting observations by systematically evaluating CNA changes and the genes they affect during engraftment and passaging in a large, internationally collected set of PDX models, comparing both RNA and DNA-based approaches. The data collected, as part of the U.S. National Cancer Institute (NCI) PDXNet (PDX Development and Trial Centers Research Network) Consortium and EurOPDX consortium, comprises 1548 PT and PDX datasets (1451 unique samples) from 509 models derived from American, European and Asian cancer patients. Our study demonstrates that prior reports of systematic copy number divergence between patient tumors and PDXs are incorrect, and that there is high retention of copy number during PDX engraftment and passaging. This work also finely enumerates the copy number profiles in hundreds of publicly available models, which will enable researchers to assess the suitability of each for individualized treatment studies.

## RESULTS

### Catalog of copy number alterations in PDXs

We have assembled copy number alteration (CNA) profiles of 1451 unique samples (324 patient tumor, PT, and 1127 PDX samples) corresponding to 509 PDX models contributed by participating centers of the PDXNET, the EurOPDX consortium, and other published datasets^9,34^ (see METHODS, Supplementary Table 1 and Supplementary Fig. 1). We estimated copy number (CN) from five data types: single nucleotide polymorphism (SNP) array, whole-exome sequencing (WES), low-pass whole-genome sequencing (WGS), RNA sequencing (RNA-Seq) and gene expression array data, yielding 1548 tumor datasets including samples assayed on multiple platforms. Paired-normal DNA and in some cases, paired normal RNA, were also obtained to calibrate WES and RNA-Seq tumor samples. To estimate the CNA profiles for the different data types, we used tools including ASCAT for SNP arrays^35^, Sequenza for tumor-normal WES^36^, qDNAseq^37^ and ASCAT for WGS and e-karyotyping^38^ for gene expression (RNA-Seq and gene expression array) data (see METHODS). Copy number segments for each sample were filtered for measurement noise, median-centered, and intersected with gene coordinates (see METHODS, Supplementary Data 1).

The combined PDX data represent 16 broad tumor types (see METHODS), with 64% (n=324) of the models having their corresponding patient tumors assayed and another 64% (n=328) having multiple PDX samples of either varying passages (ranging from P0 – P21) or varying lineages from propagation into distinct mice (Fig. 1a, Supplementary Table 2). The distributions of PT and PDX samples across different tumor types, passages, and assay platforms (Fig. 1b, Supplementary Fig. 2-12) show the wide spectrum of this combined dataset, which is the most comprehensive copy number profiling of PDXs compiled to date. Additionally, our data include 7 patients with multiple tumors collected either from different relapse time points or different metastatic sites, resulting in multiple PDX models derived from a single patient.

**Figure 1.**
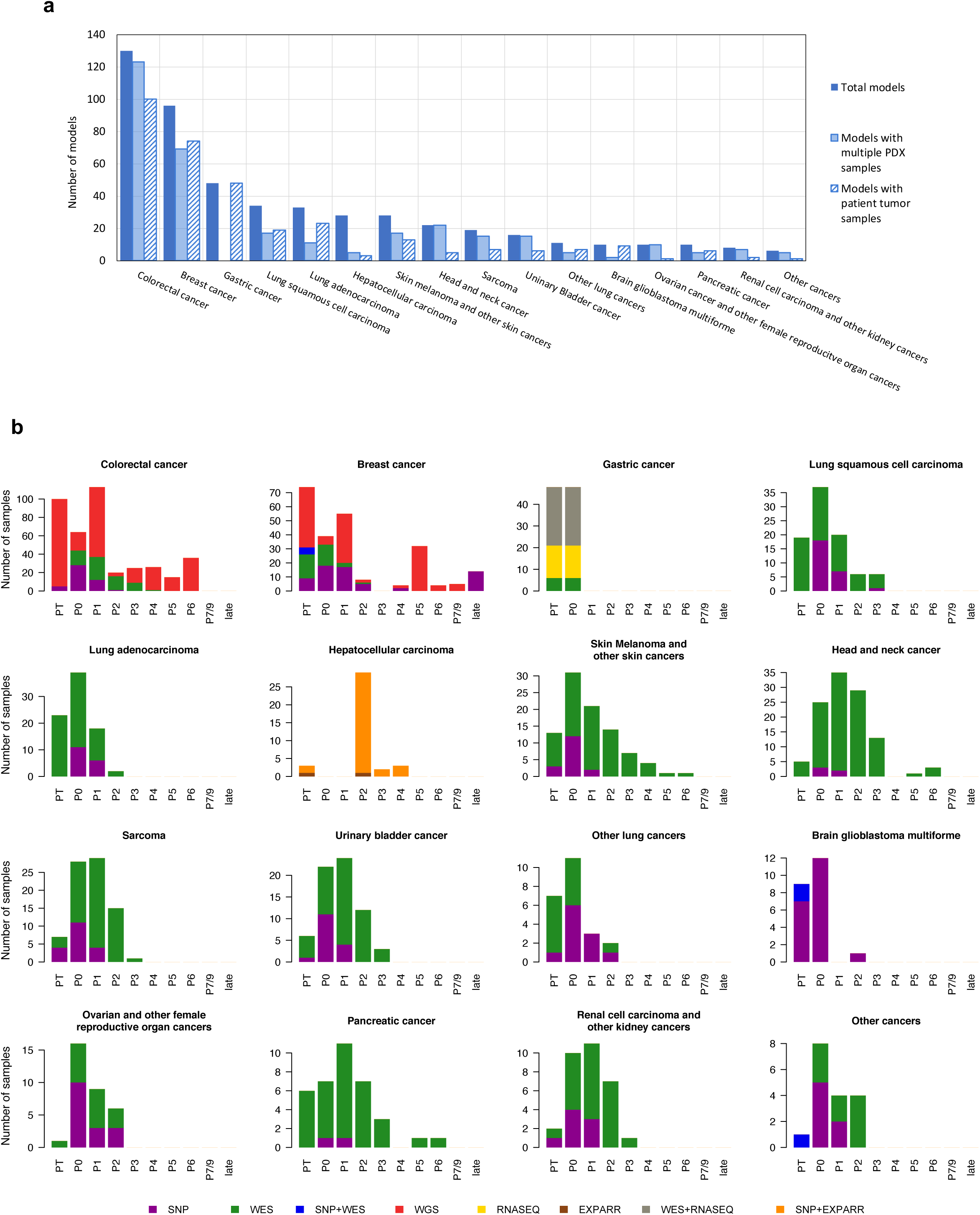
PDXNet and EurOPDX patient derived xenograft datasets used for copy number profiling across 16 tumor types. (**a**) Numbers of PDX models for each tumor type, with models also having multiple PDX samples or having matched patient tumor samples specified. (**b**) Distributions of datasets by passage number and assay platform for patient tumors and PDX samples, separated by tumor type. “Late” passages include P18, P19 and P21 samples.

### Comparison of CNA profiles from SNP array, WES and gene expression data

To compare the CNA profiles from different platforms in a controlled fashion, we assembled a benchmarking dataset with matched measurements across multiple platforms (Supplementary Table 3, Supplementary Fig. 13 – 17). Copy number calling has been reported to be noisy for several data types^39,40^, and we observed that quantitative comparisons between CNA profiles are sensitive to: (1) the thresholds and baselines used to define gains and losses, (2) the dynamic range of copy number values from each platform, and (3) the differential impacts of normal cell contamination for different measurements. To control for such systematic biases, we assessed the similarity between two CNA profiles using the Pearson correlation of their log_2_(CN ratio) values across the genome in 100kb windows. Regions with discrepant copy number were identified as those with outlier values from the linear regression model (see METHODS).

#### CNAs from WES are consistent with CNAs from SNP array data

While SNP arrays are widely accepted for estimating tumor CNA profiles^41,42^, CNA estimates from WES data have more uncertainty^36,43^. We implemented a WES-based CNA pipeline and benchmarked it against SNP array-based estimates for matched samples, which we used as a gold standard. Copy number gain or loss segments (see METHODS) from SNP arrays were of a higher resolution (Fig. 2a; median/mean segment size: 1.49/4.05 Mb for SNP, 4.70/14.6 Mb for WES, *p* < 2.2e^−16^) and wider dynamic range (Fig. 2b; range of log_2_(CN ratio): –8.62 – 2.84 for SNP, –3.04 – 1.85 for WES, *p* < 2.2e^−16^). The difference in range is apparent in the linear regressions between platforms (Supplementary Fig. 19a). These observations take into account the broad factors affecting CNA estimates across platforms, such as the positional distribution of sequencing loci; the sequencing depth of WES (10 – 280X); and the superior removal of normal cell contamination by SNP array CNA analysis workflows using SNP allele frequencies^35^.

**Figure 2.**
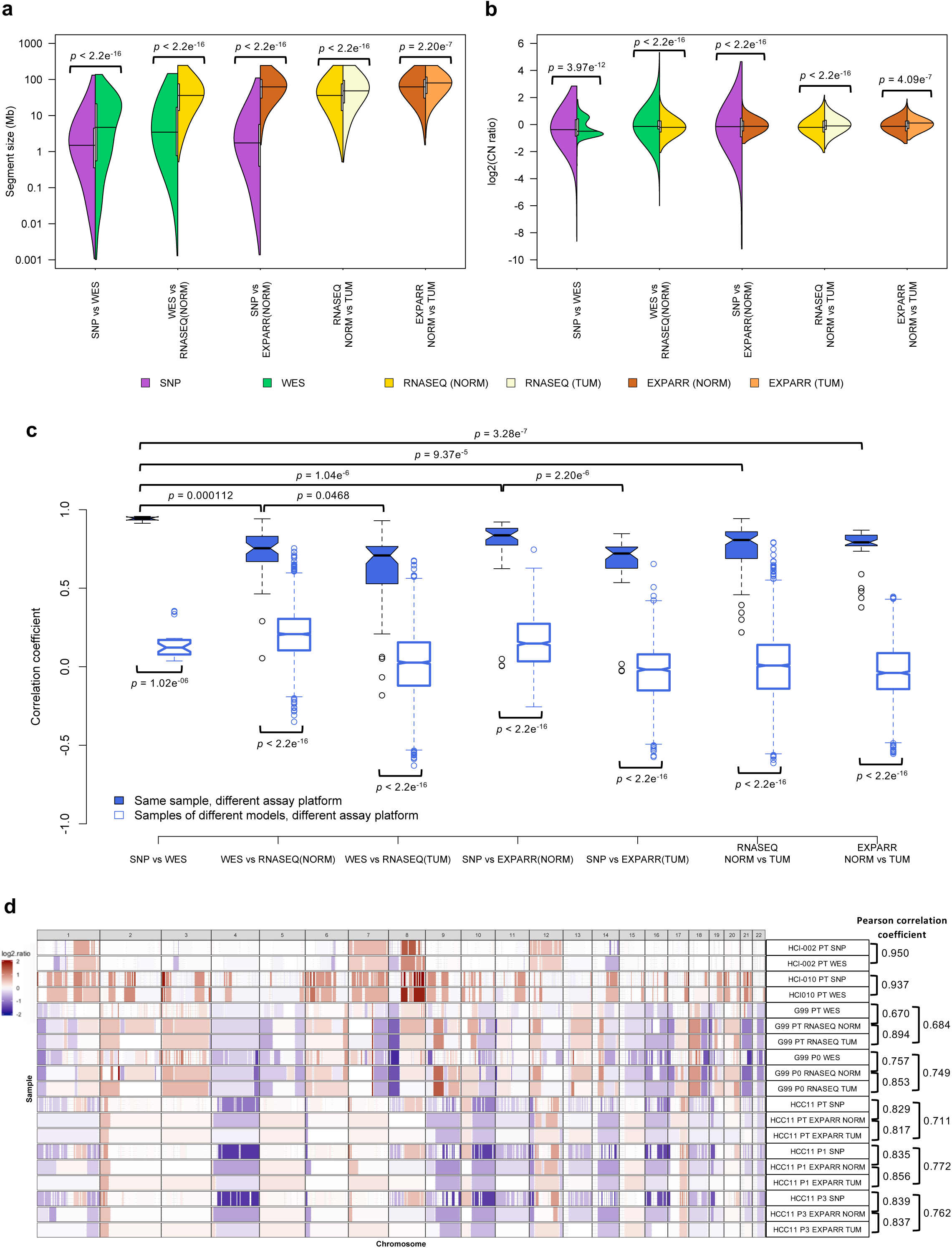
Comparisons of resolution and accuracy for copy number alterations estimated by DNA-based and expression-based methods. (**a**) Pairwise comparisons of distributions of segment size (Mb) of CNAs estimated by different measurement platforms in the benchmarking dataset (see Supplementary Table 3). CNAs are regions with (|log_2_(CN ratio)| ≥ 0.1). P-values indicate significance of difference between distributions by Wilcoxon rank sum test. (**b**) Pairwise comparisons of distributions of log_2_(CN ratio) of CNA segments. P-values were computed by Kolmogorov-Smirnov test. (**c**) Distributions of Pearson correlation coefficient of median-centered log_2_(CN ratio) in 100-kb windows from CNA segments between pairs of samples estimated by different platforms (see Supplementary Table 3). Samples with non-aberrant profiles in SNP array and WES data are omitted (Range (5-95 percentile) of log_2_(CN ratio) < 0.3). P-values indicate Wilcoxon rank sum test. (**d**) Examples of CNA profiles in comparisons of different platforms. Pearson correlation coefficients of CNA segments between pairs of samples are shown on the right. (SNP: SNP array, WES: whole-exome sequencing, RNASEQ: RNA sequencing, EXPARR: gene expression array, NORM: normalization by median expression of normal samples, TUM: normalization by median expression of tumor samples)

Despite the superiority of SNP arrays, we observed strong agreement between SNP arrays and WES, with significantly higher Pearson correlation coefficients on matched samples than samples of different models (range: 0.913 – 0.957 for matched samples, 0.0366 – 0.354 for unmatched samples, *p* = 1.02e^−06^), with the exception of 2 samples that lacked CNA aberrations (Fig. 2c, Supplementary Fig. 13, 18 and 19a). Regions with discordant copy number between platforms could also be identified (Supplementary Fig. 19a, see METHODS). The discordant copy number regions largely correspond to small focal events (average size 1.53Mb) detectable by SNP arrays but missed by WES (Supplementary Fig. 19b). Still, CNA profiling by WES is reliable in most cases, with 99% of the genome locations across the samples consistent with the values from SNP arrays.

#### Low accuracy for gene expression-derived CNA profiles

To compare the suitability of gene expression for quantifying evolutionary changes in CNA, we adapted the e-karyotyping method used in Ben-David et al.^26,38,44^ for RNA-Seq and gene expression array data. For each tumor type, the expression values were calibrated relative to either median expression of non-tumor tissue RNA samples, or relative to median expression of tumor samples when normal samples were not available (Supplementary Fig. 15 and 17). Copy number segments calibrated by non-tumor expression were of higher resolution (Fig. 2a; median/mean segment size: 36.0/51.9 Mb for RNASEQ NORM, 48.2/65.3 Mb for RNASEQ TUM, *p* < 2.2e^−16^; 62.0/72.4 Mb for EXPARR NORM, 80.1/85.2 Mb for EXPARR TUM, *p* = 2.20e^−07^) and wider dynamic range (Fig. 2b; range of log_2_(CN ratio): –2.07 – 2.17 for RNASEQ NORM, –1.79 – 1.81 for RNASEQ TUM, *p* < 2.2e^−16^; –1.40 – 1.89 for EXPARR NORM, –1.13 – 1.59 for EXPARR TUM, *p* = 4.09e^−07^) compared to segments calculated by calibration with tumor samples. This was true for both RNA-Seq and gene expression array platforms.

A notable problem with the expression-based calls is that the alternative expression calibrations can have a major impact on called gains and losses. This is especially apparent for regions frequently called as gains or losses in specific tumor types (Supplementary Fig. 20), e.g. as identified in other studies^45–47^. Chromosomes 8q and 13 were almost exclusively identified as gains and chromosomes 21 and 22 were almost exclusively as losses in the gastric cancer RNA-Seq dataset when normal samples were used for calibration. Similarly, we called exclusive gains in chromosomes 7q and 20 and losses in chromosomes 4q31-35, 8p,16q and 21 using normal samples for calibration for the hepatocellular carcinoma expression array dataset. However, changing the calibration to use tumor samples resulted in these regions being erroneously called with approximately equal frequencies of gains and losses. These alternate methodologies yielded strong variability in the CNA calls, and this was the case for each of the RNAseq and expression array datasets (Pearson correlation range: 0.218 – 0.943 for RNASEQ NORM vs TUM, 0.377 – 0.869 for EXPARR NORM vs TUM, Fig. 2c and Supplementary Fig. 21). For each, this range of correlations was far greater than was observed in comparisons between the DNA-based methods (*p* = 9.37e^−5^ and *p* = 3.28e^−07^ relative to SNP vs WES). This indicates the problematic nature of RNA-based CNA calling with calibration by tumor samples, which has been used when normal samples are not available.

We observed other measures showing the limitations of RNA-based CNA calling. Expression-based calling had segmental resolution an order of magnitude worse than the DNA-based methods (Fig. 2a and Supplementary Fig. 14 – 17; median/mean segment size: 3.45/14.0 Mb for WES, 36.0/51.9 Mb for RNASEQ NORM, *p* < 2.2e^−16^; 1.73/5.18 Mb for SNP, 62.0/72.4 Mb for EXPARR NORM, *p* < 2.2e^−16^). The range of detectable copy number values was also superior for DNA-based methods (Fig. 2b; range of log_2_(CN ratio): –6.00 – 5.33 for WES, –2.07 – 2.17 for RNASEQ NORM, *p* < 2.2e^−16^; –9.19 – 4.65 for SNP, –1.40 – 1.89 for EXPARR NORM, *p* < 2.2e^−16^). In addition, there was a lack of correlation between the expression-based and DNA-based methods (range: 0.0541 – 0.942 for WES vs RNASEQ (NORM); 0.00517 – 0.921 for SNP vs EXPARR (NORM)) (Fig. 2c and Supplementary Fig. 22 and 23). CNA estimates after tumor-based expression normalization resulted in further discordance with DNA-based copy number results (range: –0.182 – 0.929, *p* = 0.0468 for WES vs RNASEQ (TUM); –0.0274 – 0.847, *p* = 2.20e^−06^ for SNP vs EXPARR (TUM)). Many focal copy number events detected by DNA-based methods, as well as some larger segments, were missed by the expression-based methods (Supplementary Fig. 24). Representative examples illustrating the superior resolution and accuracy from DNA-based estimates are given in Fig. 2d (see also Supplementary Fig. 19a and 25).

### Concordance of PDXs with patient tumors and during passaging

We tracked the similarity of CNA profiles during tumor engraftment and passaging by calculating the Pearson correlation of gene-level copy-number for samples measured on the same platform (see METHODS, Supplementary Fig. 26-64). All pairs of samples derived from the same PDX model were compared – yielding 501 PT-PDX and 1257 PDX-PDX pairs.

For all DNA-based platforms we observed strong concordance between matched PT-PDX and PDX-PDX pairs, significantly higher than between different models from the same tumor type and the same center (*p* < 2.2e^−16^) (Fig. 3a – c, correlation heatmaps in Supplementary Fig. 27 – 63). We observed no significant difference in the correlation values between PT-PDX and PDX-PDX pairs for SNP array data (median correlation PT-PDX = 0.950, PDX-PDX = 0.964; *p* > 0.05), though there were small but statistically significant shifts for WES (PT-PDX = 0.874, PDX-PDX = 0.936; *p* = 2.31e^−16^) and WGS data (PT-PDX = 0.914, PDX-PDX = 0.931; *p* = 0.000299). PT samples have a smaller CNA range than their derived PDXs (median ratio PT/PDX / PDX/PDX: 0.832/0.982, *p* = 0.000120 for SNP; 0.626/0.996, *p* < 2.2e^−16^ for WES; 0.667/1.00, *p* < 2.2e^−16^ for WGS; Supplementary Fig. 64b and 65), which can be attributed to stromal DNA in PT samples “diluting” the CNA signal. In PDXs, the human stromal DNA is reduced^9,15^. The minimal effect for SNP array data confirm this interpretation – human stromal DNA contributions to CNA estimates can be removed from SNP arrays based on allele frequencies of germline heterozygous sites, while such contributions to WES and WGS have higher uncertainties.

**Figure 3.**
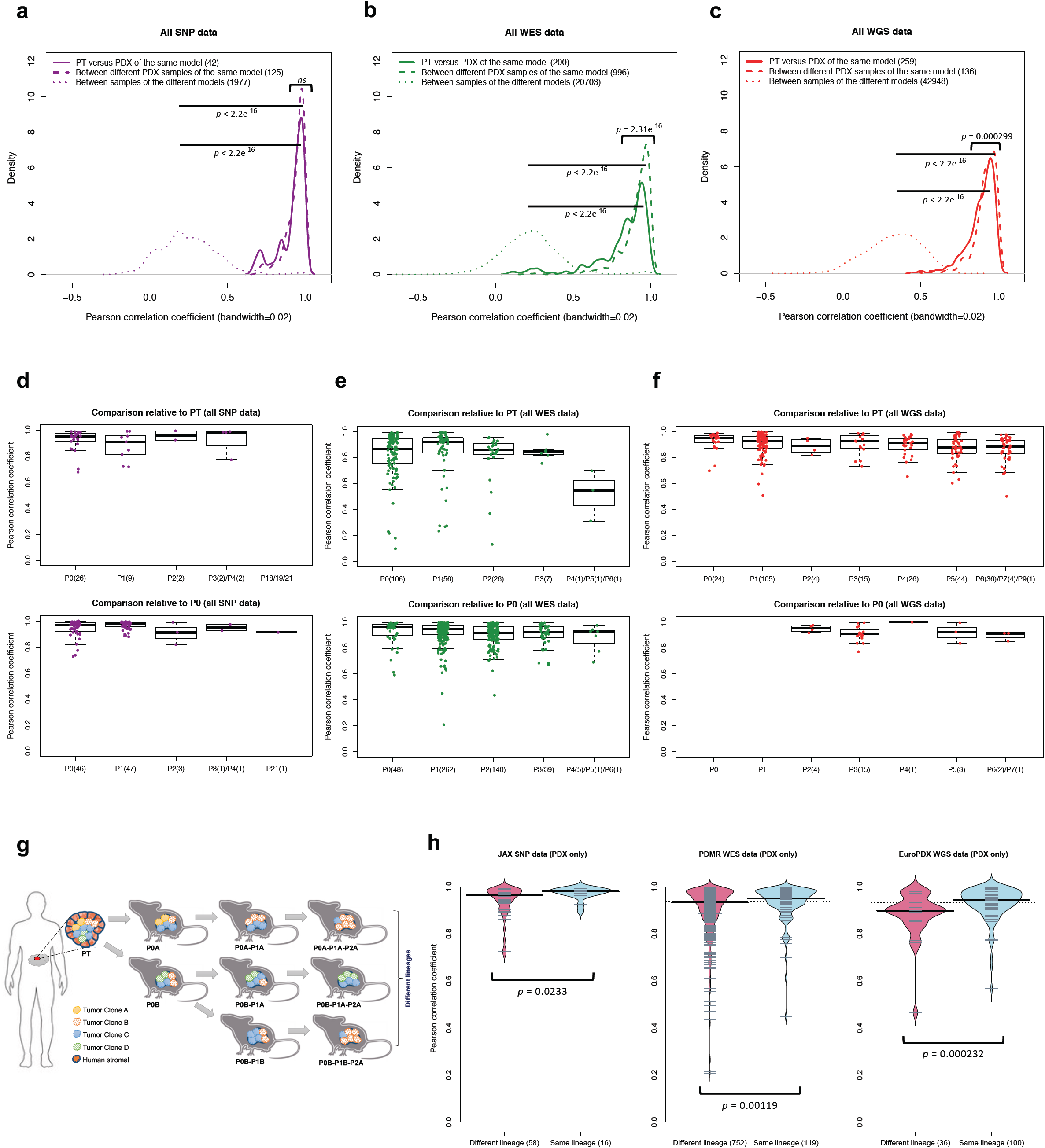
Comparisons of copy number alterations from patient tumor to early and late PDX passages. (**a-c**) Distributions of Pearson correlation coefficient of gene-based copy number, estimated by (**a**) SNP array, (**b**) WES, and (**c**) WGS, between: PT-PDX samples from the same model; PDX-PDX samples of the same model; and samples of different models from a common tumor type and contributing center. P-values were computed by Wilcoxon rank sum test (ns: not significant p-value > 0.05). (**d-f**) Distributions of Pearson correlation coefficients of gene-based copy number, estimated by (**d**) SNP array, (**e**) WES, and (**f**) WGS, among patient tumor and PDX passages of the same model. Comparisons relative to PT and P0 are shown (higher passages are shown in Supplementary Fig. 66). (**g**) Schematic of lineage splitting during passaging and expansion of tumors into multiple mice. This is a simplified illustration for passaging procedures in which different fragments of a tumor are implanted into different mice. (**h**) Pearson correlation distributions for PDX sample pairs of different lineages and sample pairs within the same lineage: for JAX SNP array, PDMR WES, and EuroPDX WGS datasets. P-values were computed by Wilcoxon rank sum test. The numbers in the parentheses represent the number of pairwise correlations.

We also performed intra-model comparisons using RNA-based approaches, but the Pearson correlations between pairs of samples did not clearly reproduce the Pearson correlations from DNA-based platforms for those same sample pairs (Supplementary Fig. 66a). To clarify this, we considered just the highly-correlated cases (>0.8 for SNU-JAX Gastric cancer WES, >0.9 for SIBS HCC SNP). We observed that the correlation values for the corresponding RNA-based methods were lower and had higher variance (*p* < 0.05, Supplementary Fig. 66b). In particular, the tumor-median normalization for expression data resulted in significant differences from DNA-based methods.

#### Late PDX passages maintain CNA profiles similar to early passages

Next, we asked if there is any systematic evolution of copy number during engraftment and passaging. Mouse environment-driven evolution, if present, should reduce CN correlations relative to early samples, such as the primary tumor or first engraftment (P0). However, we observed no apparent effect during passaging on the SNP, WES, or WGS platforms (Fig. 3d – f, Supplementary Fig. 67). For example, the SNP data showed no significant difference between passages (Fig. 3d and Supplementary Fig. 67a). For those models having very late passages (14 breast cancer models, P18 to P21), there was a small but statistically significant correlation decrease compared to models with earlier passages (*p* < 8.98e^−05^, Supplementary Fig. 68), indicating some copy number changes can occur over long-term passaging (Supplementary Fig. 38). However even at these late passages, the correlations to early passages remained high (median = 0.896). In any given comparison, only a small proportion of the genes were affected by copy number changes (median: 2.72%, range: 1.03% – 11.9%). Genes that are deleted and subsequently gained in the later passages (top left quadrant of regression plots, Supplementary Fig. 69) suggest selection of preexisting minor clones as the key mechanism in these regions. For WES and WGS data, more variability in the correlations can be observed (Fig. 3e and f, Supplementary Fig. 66b and c), likely due to a few samples having more stromal contamination or low aberration levels (Supplementary Fig. 64b and 65). However, the lack of downward trend over passaging was also apparent in these sets.

#### PDX copy number profiles trace lineages

We next compared the similarity of engrafted PDXs of the same model with the same passage number (i.e. all P0s, all P1s, all P2s, etc.). Surprisingly, we discovered that these fragments were not more similar than PDXs from different passage numbers (Fig. 3d – e and Supplementary Fig. 66b, IQR of correlation coefficient for same-passages/different-passages: 0.0700/0.0619 for SNP and 0.103/0.0979 for WES). To further this analysis, we defined, for JAX SNP array and PDMR WES datasets, samples within a lineage as those differing only by consecutive serial passages, while we defined lineages as split when a tumor was divided and propagated into multiple mice (Fig. 3g). For the EurOPDX CRC and BRCA WGS datasets, such lineage splitting was due only to cases with initial engraftment of different fragments of the PT, i.e., PDX samples of different passages were considered as different lineages if they originate from different PT fragments. We observed lower correlation between PDX samples from different lineages compared to within a lineage (Fig. 3h, *p* = 0.0233 for SNP, *p* = 0.00119 for WES, *p* = 0.000232 for WGS), despite a majority of these pairwise comparisons exhibiting high correlation (>0.9). A few examples of models exhibiting large drift between lineages include TM01500 (Supplementary Fig. 29); 416634, 558786 and 665939 (Supplementary Fig. 50); 135848 and 762968 (Supplementary Fig. 51); 245127 and 959717 (Supplementary Fig. 52); 287954 and 594176 (Supplementary Fig. 56); 174316 and 695221 (Supplementary Fig. 57).

We next asked if the phylogenetic distance between samples could explain the observed shifts in the correlations. These distance relationships are clearest for the CRC and BRCA WGS sets because these models have only one lineage split occurring at the engraftment stage. We compared correlation as a function of phylogenetic distance within a lineage, which in this phylogeny is simply equal to the difference in passage number between the two samples. Increase in passage difference did not consistently reduce the correlation between samples (Supplementary Fig. 70). This suggests that lineage-splitting is often responsible for deviations in CNAs between samples, and that copy number evolution during passaging mainly arises from evolved spatial heterogeneity^27^.

### Genes with copy number alterations acquired during engraftment and passaging show no preference for cancer or treatment-related functions

Next, we investigated which genes tend to undergo copy number changes. Genes with changes during engraftment or during passaging were identified based on a residual threshold with respect to the improved linear regression^48^ (see METHODS, Supplementary Fig. 26). A low copy number change threshold (|log_2_(CN ratio) change| > 0.5) was selected to include genes with subclonal alterations. To test for functional biases, we compared CNA-altered genes to gene sets with known cancer- and treatment-related functions, notably genes in TCGA oncogenic signaling pathways^49^; genes with copy number and expression changes associated with therapeutic sensitivity, resistance or changes in drug response from the JAX Clinical Knowledgebase^50,51^; and genes with frequent amplifications or deletions in the Cancer Gene Census^52^ (Cosmic version 89). We calculated the proportion of altered genes for sample pairs from each model across all platforms and tumor types. In agreement with the high maintenance of CNA profiles described above, we found the proportion of altered protein-coding genes to be low (median/IQR: 1.90%/4.11% PT-PDX, 1.25%/3.60% PDX-PDX pairs, Fig. 4a). Only 8.78% of PT-PDX pairs and 4.53% PDX-PDX pairs showed >10% of their protein-coding genes altered. We observed no significant increase (*p* < 0.1) in alterations among any of the cancer gene sets compared to the background of all protein-coding genes, for either the PT-PDX or PDX-PDX comparisons. This provides evidence that there is no systematic selection for CNAs in oncogenic or treatment-related pathways during engraftment or passaging. We next considered tumor-type-specific effects, focusing on types with larger numbers of models to ensure statistical power (breast cancer, colorectal cancer, lung adenocarcinoma and lung squamous cell carcinoma). Genomic Identification of Significant Targets in Cancer (GISTIC)^53,54^ analysis of TCGA tumors has previously identified significantly altered genomic driver regions which can be used to differentiate tumor types and subtypes^55–58^. We observed no significant increase in alterations in tumor-type-specific GISTIC gene sets compared to the background (*p* < 0.1) for either PT-PDX or PDX-PDX comparisons (Fig. 4b).

**Figure 4.**
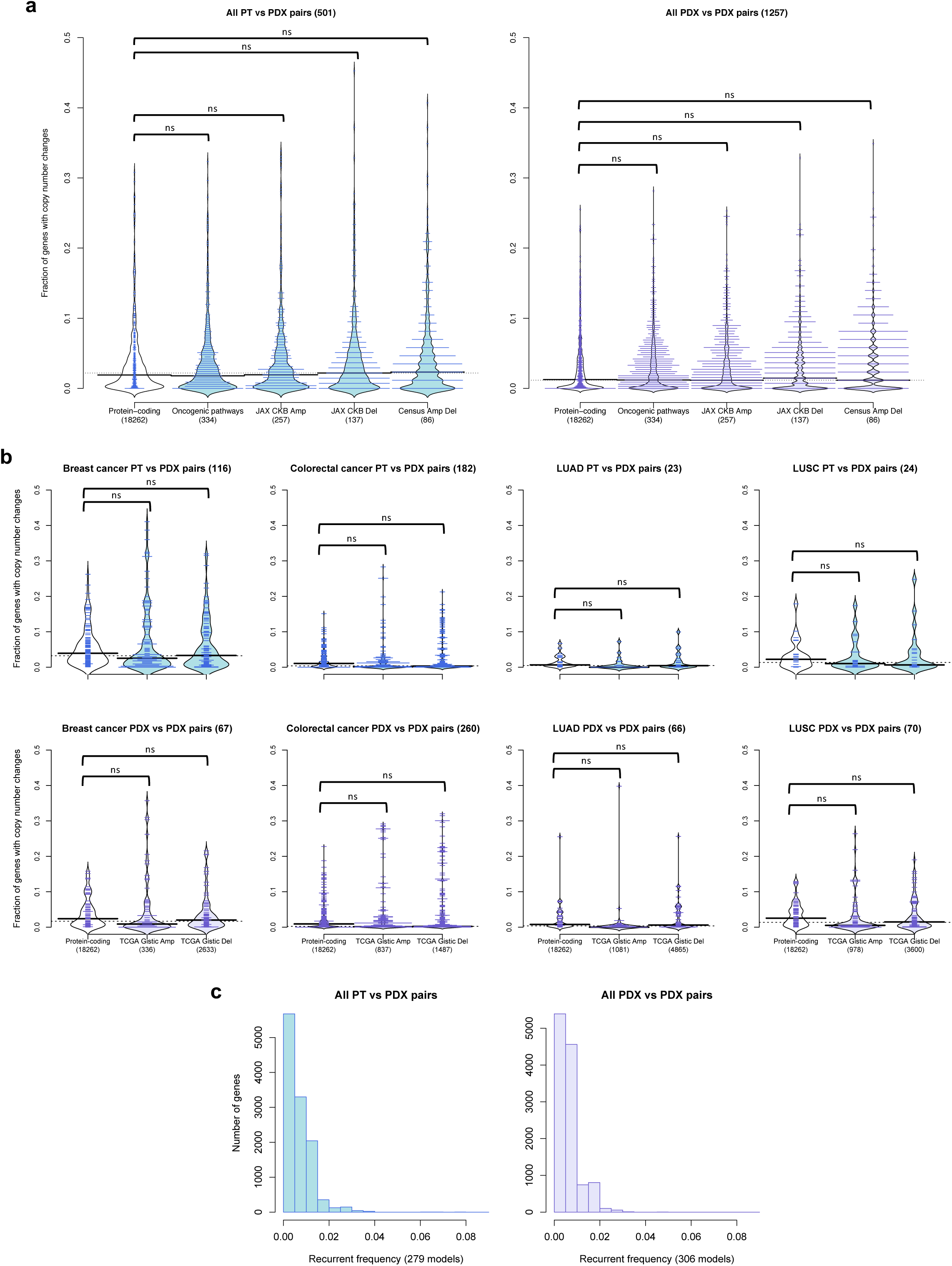
Cancer pathway analysis for copy number altered genes during engraftment and passaging. (**a**) Distribution of proportion of altered genes for pairwise comparisons of PDX samples for various gene sets: Protein-coding genes annotated by Ensembl; Oncogenic signaling pathways identified by TCGA^49^; JAX CKB^50,51^ Amp indicates genes with copy number gain or over-expression associated with therapeutic sensitivity or resistance; JAX CKB Del indicates genes with copy number loss or under-expression associated with therapeutic sensitivity or resistance; Census Amp Del indicates genes with frequent amplifications or deletions in the Cancer Gene Census^52^. CNA genes were identified by |residual| > 0.5 from linear regression model. P-values were computed by Wilcoxon rank sum test (ns: not significant, p > 0.1). (**b**) Distribution of proportion of altered genes for pairwise comparisons within breast cancer, colorectal cancer, lung adenocarcinoma (LUAD) and lung squamous cell carcinoma (LUSC) models. Prevalence of alterations in significantly amplified (TCGA Gistic Amp) or deleted (TCGA Gistic Del) genes for the corresponding tumor type are shown. P-values were computed by Wilcoxon rank sum test (ns: not significant, p > 0.1). The numbers in the parentheses in the horizontal axis represent the number of genes, and those in the plot title represent the number of pairwise correlations. (**c**) Recurrence frequency of protein coding genes with copy number alterations, |residual| > 1, across all models in PT-PDX and PDX-PDX comparisons.

#### Low recurrence of altered genes across models

We tested if any particular genes often recurred in CNAs across models. Using a stringent CNA threshold (|log2(CN ratio) change| > 1.0 with respect to linear regression model) to distinguish genes with possible functional impact (see METHODS), we observed a very low recurrent frequency (Fig. 4c), with only 12 and 2 genes recurring at > 5% frequency for PT-PDX and PDX-PDX comparisons, respectively (Supplementary Table 4). No gene had a recurrence frequency higher than 8.96%. We observed that all these recurrent genes overlapped models in which one sample displayed an unusually large gain or loss (|log2 (CN ratio)| > 1.5). This suggests that these regions may be subject to more noise in the CNA estimation procedure at these loci (Supplementary Fig. 71). None of these recurrent genes overlapped cancer- or treatment-related gene sets, nor did they intersect genes (n=3) reported by Ben-David et al.^26^ to have mouse-induced copy number changes associated with drug response in the CCLE^59,60^ database. We further queried from CCLE data whether any of these recurrent genes showed evidence for copy number-related drug response (see METHODS, Supplementary Table 5). For the 6 genes with sufficient data available, we found no association between copy number and drug response mediated by gene expression (*q-value* < 1).

#### Absence of CNA shifts in 130 WGS patient tumour, early passage PDX and late passage PDX trios

We next investigated whether recurrent CNA changes occur in PDXs in a tumor-type specific fashion. To this aim, we analysed further the WGS-based CNA profiles of large metastatic colorectal (CRC) and breast cancer (BRCA) series (see METHODS), respectively composed of 87 and 43 matched trios of patient tumour (PT), PDX at early passage (PDX-early) and PDX at later passage (PDX-late). We carried out GISTIC analysis to identify recurrent CNAs by evaluating the frequency and amplitude of observed events^53,54^. GISTIC was applied separately for each PT, PDX-early (P0-P1 for CRC, P0-P2 for BRCA) and PDX-late (P2-P7 for CRC, P3-P9 for BRCA) cohorts of CRC and BRCA (Supplementary Table 6). As expected, CRCs and BRCAs generated different patterns of significant CNAs, with each similar to the GISTIC patterns in their respective TCGA series (Supplementary Fig. 72). However, within each tumour type GISTIC profiles of the PT, PDX-early, and PDX-late cohorts were virtually indistinguishable (Fig 5a and Supplementary Fig. 72), demonstrating no gross genomic alteration systematically acquired or lost in PDXs.

**Figure 5.**
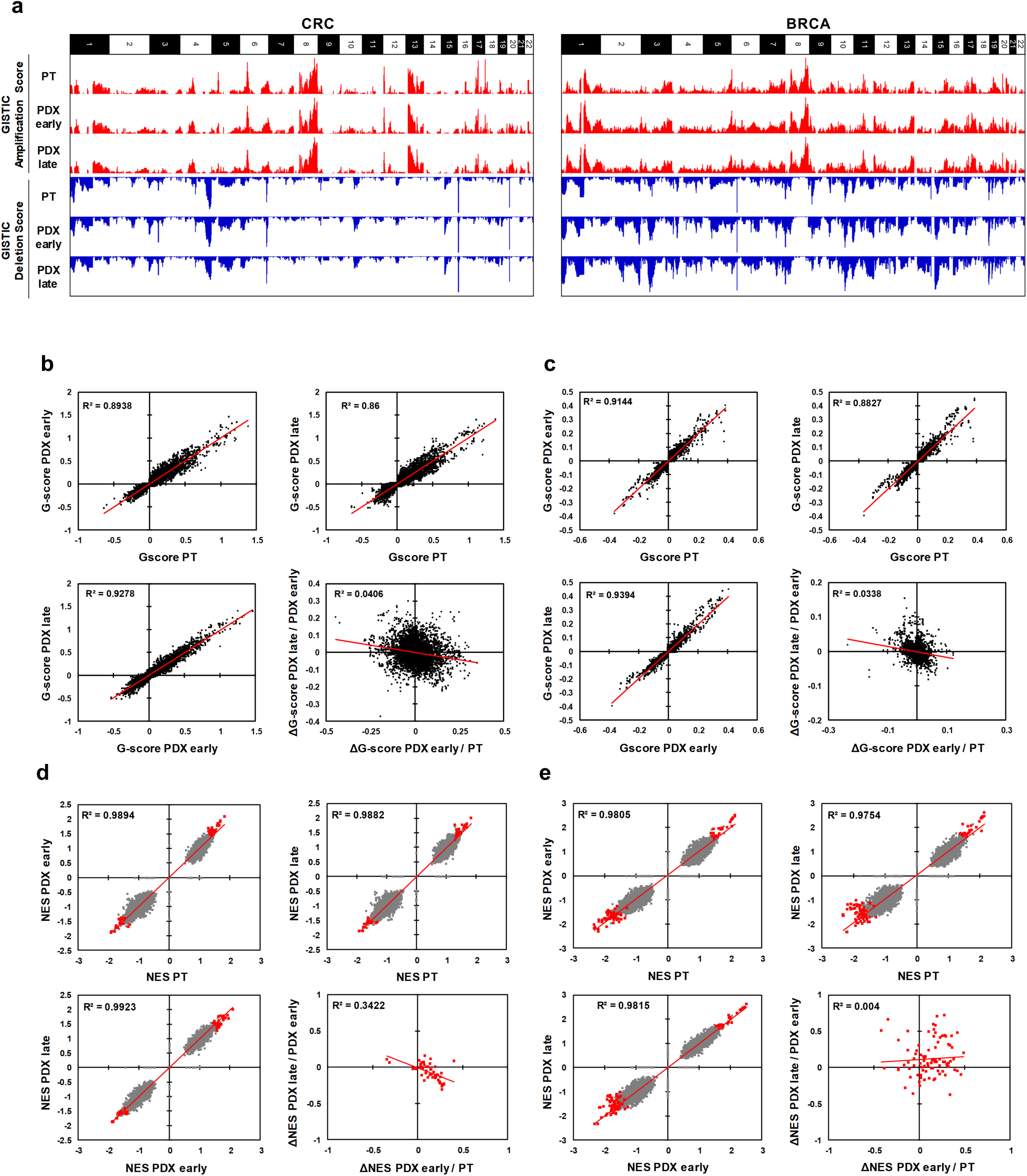
Absence of mouse-driven recurrent CNAs during engraftment and propagation of colorectal and breast cancer PDXs. (**a**) Bar charts representing genome-wide GISTIC G-score for amplifications (red) and deletions (blue) in each of the three cohorts (PT, PDX-early, PDX-late) for CRC and BRCA. (**b-c**) Scatter plots comparing gene-level GISTIC G-score between each of the three cohorts, for (**b**) CRC and (**c**) BRCA. Bottom-right panels of (**b**) and (**c**): scatter plots comparing ΔG-scores from PT to PDX-early and from PDX early to PDX-late. (**d-e**) Scatter plots comparing GSEA Normalized Enrichment Score (NES) for gene sets between each of the three cohorts, for (**d**) CRC (**e**) and BRCA. Grey dots: all gene sets; red dots: gene sets significantly enriched in at least one among the PT, PDX-early, PDX-late cohorts. Bottom-right panels of (**d**) and (**e**): scatter plots comparing ΔNES from PT to PDX-early and from PDX-early to PDX-late.

To clarify these behaviors, we carried out gene-level analysis, where each gene was attributed the GISTIC score (G-score) of the respective segment (Supplementary Table_7). In both the CRC and BRCA cohorts, gene-level G-scores of the PTs were highly correlated with the respective PDX-early and PDX-late cohorts (Fig. 5b and c). Moreover, PT versus PDX correlations were comparable to PDX-early versus PDX-late correlations. To search for progressive shifts, we compared the change in G-score (ΔG): (i) from tumor to PDX-early and (ii) from PDX-early to PDX-late. Correlations in these two ΔG values, as shown in the bottom-right panels of Fig. 5b and c, was absent or even slightly negative. These results confirmed the absence of systematic CNA shifts in PDXs even under high resolution, gene-level analysis.

#### Lack of CNA-based functional shifts in trios confirms the absence of mouse-specific evolution

We then considered the possibility of systematic copy number evolution at the pathway level in these triplets. To evaluate this, we performed Gene Set Enrichment Analysis (GSEA)^61,62^ using G-scores to rank genes in each cohort (See METHODS). Consistent with the known recurrence of cancer CNAs at driver genes, multiple gene sets displayed significant enrichment in individual cohorts. To avoid spurious apparent enrichment for sets of genes with adjacent chromosomal location, we implemented an additional filter based on G-score significance (see METHODS and Supplementary Table 8). After applying the Normalized Enrichment Score (NES), FDR q-value and G-score filters, 49 gene sets were found to be significant in at least one of the three CRC cohorts, and 89 gene sets in at least one of the three BRCA cohorts (Supplementary Table 9). Importantly, control gene sets composed of GISTIC hits identified in TCGA CRC and BRCA datasets were all significant, confirming that the WGS cohorts used here correctly recapitulate the major CNA features of these two cancer types.

However, differences associated with PDX engraftment and passage were negligible. For both CRC and BRCA, the NES profiles for the ~8000 gene sets of PTs were highly correlated with the respective PDX-early and PDX-late cohorts (Fig. 5d and e). Moreover, PT versus PDX correlations were comparable to PDX-early versus PDX-late correlations. To search for progressive shifts, we calculated for each significant gene set ΔNES values between PT and PDX-early, as well as between early and late PDX. Similarly to what was observed for the ΔG-scores, as shown in the bottom-right panels of Fig. 5d and e, correlations were absent or at most slightly negative, confirming the absence of systematic CNA-based functional shifts in PDXs.

### CNA evolution across PDXs is no greater than variation in patient multi-region samples

As a reference for the treatment relevance of PDX-specific evolution, we compared to levels of copy number variation in multi-region samples of patient tumors. For this we used copy number data from multi-region sampling of non-small-cell lung cancer (92 patient tumors, 295 multi-region samples) from the TRACERx Consortium^31^, performing analogous CNA correlation and gene analyses (|residual| > 0.5) between multi-region pairs (Supplementary Fig. 73). We observed no significant differences in correlation (*p* > 0.05) between patient multi-region and lung cancer PT-PDX pairs, while PDX-PDX pairs in fact showed significantly better correlation than the multi-region pairs (*p* < 0.05, Fig. 6a). These findings were consistent when tumors were grouped as adenocarcinomas, squamous cell carcinomas, or others. Cancer gene set analyses confirmed these results, with multi-region samples showing greater differences than either PT-PDX or PDX-PDX comparisons, across all the cancer gene sets considered (*p* < 0.05, Fig. 6b and Supplementary Fig. 74). These results show that PDX-associated CNA evolution is no greater than what patients experience naturally within their tumors. Our PDX collection also contains a few cases in which the patient tumor was assayed at multiple time points (relapse/metastasis) or multiple metastatic sites, allowing for controlled comparison of intra-patient variation versus PDX evolution (Supplementary Fig. 3, 4 and 7). We observed no significant difference between the CNA evolution during engraftment and passaging compared to the intra-patient samples (Fig. 6c). CNA profiles for these samples are shown visually in Fig. 6d.

**Figure 6.**
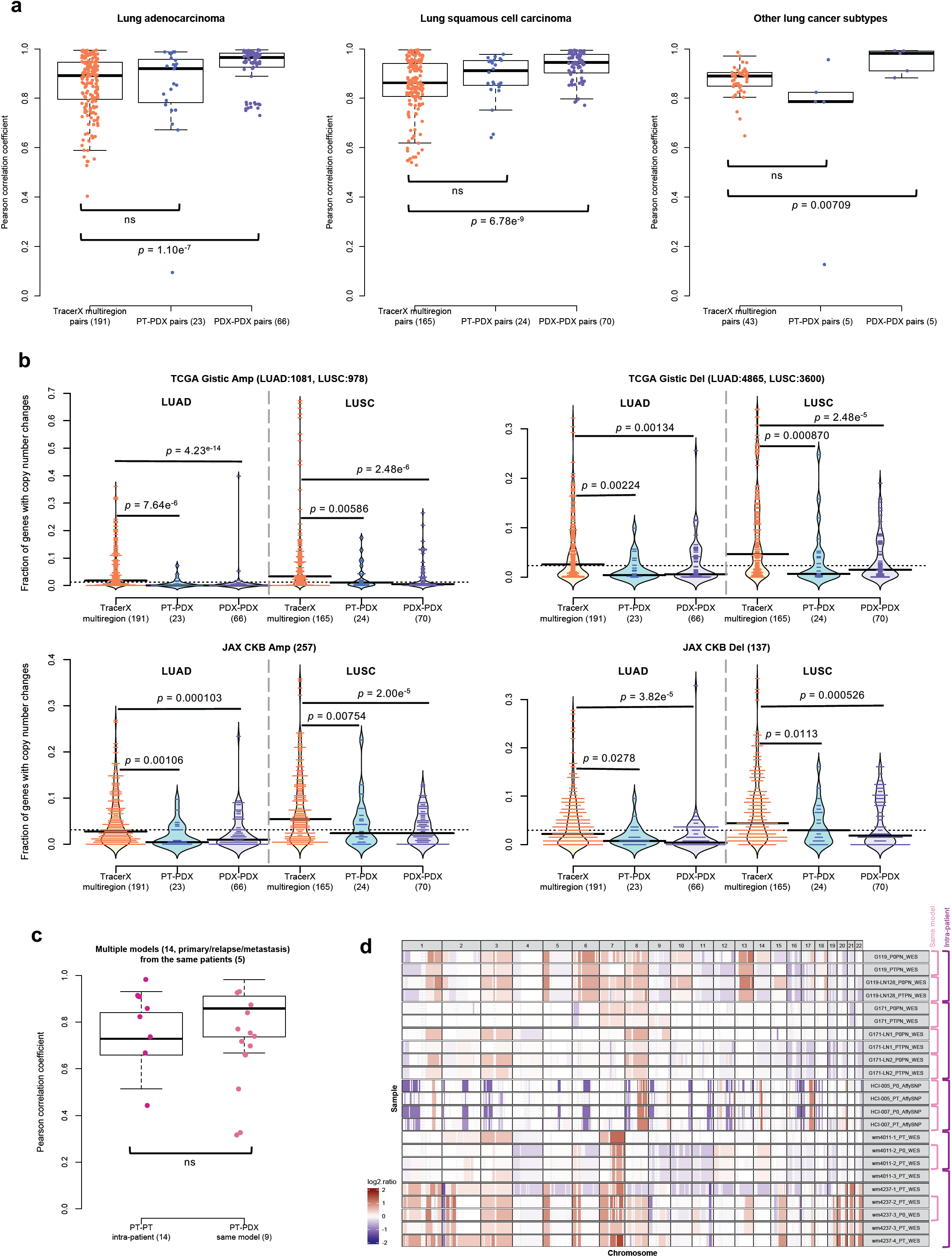
Comparison of CNA variation during PDX engraftment and passaging to CNA variation among patient multi-region, tumor relapse, and metastasis samples. (**a**) Distributions of Pearson correlation coefficients of gene-based copy number for lung adenocarcinoma (LUAD), lung squamous cell carcinoma (LUSC), and other lung cancer subtypes. Columns compare: multi-region tumor samples from TRACERx^31^; PT-PDX samples from the same model; and PDX-PDX samples from the same model. P-values indicate Wilcoxon rank sum test (ns: p-value > 0.05). (**b**) Distributions of proportion of altered genes between multi-region tumor pairs from TRACERx, and PT-PDX and PDX-PDX pairs for various gene sets for LUAD and LUSC. Gene sets are the same as in Fig. 4. TCGA Gistic and JAX CKB gene sets are shown (other gene sets are shown in Supplementary Fig. 76). (**c**) Distributions of Pearson correlation coefficients of gene-based copy number between intra-patient PT (primary/relapse/metastasis) pairs from the same patient and corresponding PT-PDX/PDX-PDX (derived from the same model; a different PT sample from the same patient generates a different model) pairs from the same set of patients. P-values were computed by Wilcoxon rank sum test (ns: p-value > 0.05). (**d**) CNA profiles of PT and PDX samples from patients with multiple PDX models from multiple PT collection (primary/relapse/metastasis).

## DISCUSSION

Here we have investigated the evolutionary stability of patient-derived xenografts, an important model system for which there have been prior reports of mouse-induced copy number evolution. To better address this, we assembled the largest collection of CNA profiles of PDX models reported to date, comprising over 1500 datasets from PDX samples of multiple passages and their originating patient tumors from more than 500 PDX models across a variety of tumor types. Our analysis demonstrated the reliability of copy number estimation by DNA-based measurements over RNA-based inferences, which are substantially inferior in terms of resolution and accuracy. The importance of DNA measurements is supported by the inconsistent conclusions by two independent studies on the same PDX expression array datasets by Gao et al.^19^ Ben-David et al.^26,63^ concluded that drastic copy number changes, driven by mouse-specific selection, often occur within a few passages. On the other hand, Mer et al.^64^ reported high similarity between passages of the same PDX model based on direct correlations of gene expression, consistent with our findings in large, independent DNA-based datasets.

To understand this, we note that the CN shifts inferred by Ben-David et al. are inherently impacted by major technical issues. First, the microarray signal for PT samples is diluted by introgressed human stromal cells, while in PDXs mouse stromal transcripts hybridize only to a fraction of the human probes^65^. As a consequence, PT samples with substantial stromal content would display a reduced signal compared to the corresponding PDX, which can lead to an erroneous inference of systematic increase in aberrations during PDX engraftment. Second, the mouse host microenvironment can affect the transcriptional profile of the PDX tumor^66^ and the quantity of mouse stroma can vary across passages. This can result in variability in the expression signal which can be wrongly inferred as CN changes, both from the tumor itself and through cross hybridization of mouse RNA to the human microarray. Although improved concordance in expression between PT and PDX can be achieved with RNA sequencing with the removal of mouse reads^67,68^, we observed that expression-based copy number inferences still have low resolution and robustness. Hence, many cancer-driving genes, which are found mainly in focal events with a size of 3Mb or lower^69–72^, cannot be evaluated for PDX-specific alterations. These issues are further worsened by the lack of tissue-matched normal gene expression profiles for calibration^38^, which have been only intermittently available but can substantially impact copy number inferences. Because of these considerations, the question of how much PDXs evolve as a consequence of mouse-specific selective pressures cannot be adequately addressed by expression data.

The studies we have presented here take into account the above issues by use of DNA data, as well as by assessing copy number changes by pairwise correlation/residual analysis to control for systematic biases, and they overall confirm the high retention of CNA profiles from PDX engraftment to passaging. We do observe larger deviations between PT-PDX than in PDX-PDX comparisons, though this is likely due to dilution of PT signal by human stromal cells. Interestingly, we found that a major contributor to the differences between PDX samples is lineage-specific drift associated with splitting of tumors into fragments during PDX propagation. This spatial evolution within tumors appears to affect sample comparisons more than time or the number of passages.

A challenge for evaluating any model system is that there is no clear threshold for genomic change that determines whether the model will still reflect patient response. Genetic variation among multi-region samples within a patient can shed light on this point, since the goal of a successful treatment would be to eradicate all of the multiple regions of the tumor. We found that the copy number differences between PT and PDX are no greater than the variations among multi-region tumor samples or intra-patient samples. Thus concerns about the genetic stability of the PDX system are likely to be less important than the spatial heterogeneity of solid tumors themselves. This result is consistent with our results on lineage effects during passaging, which indicate that intratumoral spatial evolution is the major reason for genetic drift.

We observed no evidence for systematic mouse environment-induced selection for cancer or treatment-related genes via copy number changes, though individual cases vary (see example in Supplementary Fig. 75). Moreover, only a small fraction of sample pairs (2.44%, 43 out of 1758) shows large CNA discordance (see METHODS), suggesting that clonal selection out of a complex population is rare. These results indicate that the variations observed in PDXs are mainly due to spontaneous intratumoral evolution rather than murine pressures. The extreme cases (see Supplementary Fig. 76 for examples with same lineage) may be informative for future studies of the evolutionary process, especially through consideration of repeated spatial sampling. It may be informative to compare such examples to those reported by Eirew et al.^22^, who described a variety of clonal selection dynamics during engraftment and passaging for breast cancer PDXs, as well as by Ding et al.^11^, who demonstrated the possibility of cellular selection during xenograft formation similar to that during metastasis. While such cases are uncommon in our study, further subclonal analysis may be useful for clarifying potential selection pressures.

In summary, our in-depth tracking of CNAs throughout PDX engraftment and passaging confirms that tumors engrafted and passaged in PDX models maintain a high degree of molecular fidelity to the original patient tumors and their suitability for pre-clinical drug testing. Overall, we find that PDX are highly concordant with the originating patient tumor and stable through multiple passaging, and that differences are no greater than those observed spatially within patient solid tumors. At the same time, our study does not rule out that PDXs will evolve in individual trajectories over time, and for therapeutic dosing studies, the best practice is to confirm the existence of expected molecular targets and obtain sequence characterizations in the cohorts used for testing as close to the time of the treatment study as is practicable.

## ACKNOWLEDGEMENTS

Support for the PDXNET consortium included funding provided by the NIH to the PDXNet Data Commons and Coordination Center (NCI U24-CA224067), to the PDX Development and Trial Centers (NCI U54-CA224083, NCI U54-CA224070, NCI U54-CA224065, NCI U54-CA224076, NCI U54-CA233223, and NCI U54-CA233306) and to the National Cancer Institute Cancer Genomics Cloud (HHSN261201400008C and HHSN261201500003I). The Jackson Laboratory (JAX) PDX resource data were supported by the National Cancer Institute of the National Institutes of Health under the JAX Cancer Center NCI Grant (Award Number P30CA034196). The content is solely the responsibility of the authors and does not necessarily represent the official views of the National Institutes of Health. The genomic data for JAX PDX tumors used in this work were generated by JAX Genome Technologies and Single Cell Biology Scientific Service. The development of PDX models and generation of data from Seoul National University, in collaboration with The Jackson Laboratory, was supported by the Korean Healthcare Technology R&D project through the Korean Health Industry Development Institute, funded by the Ministry of Health & Welfare, Republic of Korea (grant number: HI13C2148). Sample procurement and next generation sequencing at Huntsman Cancer Institute was performed at the Genomics and Bioinformatics Analysis and Biorepository and Molecular Pathology Shared Resources, respectively, supported by NCI P30CA042014. SNP arrays were performed at the University of Utah Health Sciences Center Genomics Core. We are grateful to Michael P. Klein for assistance with SNP array data. M.H.B. is funded by the National Institutes of Health under Ruth L. Kirschstein National Research Service Award T32HG008962 from the National Human Genome Research Institute. M.T.L. is supported by a P30 Cancer Center Support Grant CA125123 and a Core Facility Support Grant from the Cancer Research and Prevention Initiative of Texas RP170691. PDX generation and whole exome sequencing at the University of Texas MD Anderson Cancer Center were supported by the University of Texas MD Anderson Cancer Center Moon Shots Program, Specialized Program of Research Excellence (SPORE) grant CA070907. The development of PDX models and generation of data from Wistar Institute was supported by National Cancer Institute, National Institutes of Health (NCI R50-CA211199). PDMR data has been funded in whole or in part with federal funds from the National Cancer Institute, National Institutes of Health (Contract Number HHSN261200800001E). The content of this publication does not necessarily reflect the views or policies of the Department of Health and Human Services, nor does mention of trade names, commercial products, or organizations imply endorsement by the U.S. Government. The breast cancer PDX models from Washington University used for this study were developed in part through the support from The Breast Cancer Research Foundation and Fashion Footwear Charitable Foundation of New York, Inc.. The pancreatic cancer PDX models from Washington University used in this study were developed with the support of NCI grants P50 CA196510, P30 CA091842 and The Foundation for Barnes-Jewish Hospital’s Cancer Frontier Fund through the Siteman Cancer Center Investment Program. The data for these models was provided by U54 CA224083. Support for the EurOPDX consortium included funding provided by Fondazione AIRC under 5 per Mille 2018 - ID. 21091 program (E.M., A.B., L.T.), AIRC Investigator Grants 18532 (L.T.) and 20697 (A.B.), AIRC/CRUK/FC AECC Accelerator Award 22795 (L.T.), EU H2020 Research and Innovation Programme, grant agreement no. 731105 “EDIReX” (E.M., A.B., L.T., A.T.B., V.S., J.J.), Fondazione Piemontese per la Ricerca sul Cancro-ONLUS 5 per mille Ministero della Salute 2015 (E.M., L.T.), 2014 and 2016 (L.T.), My First AIRC Grant (MFAG) 19047 (C.I.), EU H2020 Research and Innovation Programme, Grant Agreement No. 754923 “COLOSSUS” (A.T.B., D.L., L.T.), European Research Council Consolidator Grant 724748 – BEAT (A.B.), Science Foundation Ireland under grant 13/CDA/2183 “COLOFORETELL’ (A.T.B), Irish Health Research Board grant ILP-POR-2019-066 (A.T.B.), ISCIII - Miguel Servet program CP14/00228 and GHD-Pink/FERO Foundation grant (V.S.), Netherlands Organization for Scientific Research (NWO) Vici grant 91814643 (J.J.), European Research Council (ERC) Synergy project CombatCancer (J.J.), Oncode Institute (J.J., R.d.B.) and Dutch Cancer Society (J.J., R.d.B.), NCI grant U24 CA204781 (J.H.C., T.F.M.). The authors thank the Breast Cancer Group from VHIO for providing study materials. We thank Debbie M. Krupke from The Jackson Laboratory for assistance with organizing the tumor type information.

## AUTHOR CONTRIBUTIONS

X.Y.W., C.J.B., J.J., A.T.B., L.T, J.A.M., C.I., E.M. and J.H.C. conceived and jointly supervised the study. X.Y.W. organized the study, collected and structured the data, and designed and carried out the analyses. J.G. collected and organized the EurOPDX data and carried out the analyses. X.Y.W., E.M. and J.H.C. wrote the manuscript. J.G, C.I, Z.-M.Z., A.S., and M.W.L. contributed to the refinement of the manuscript. A.S. and M.W.L developed the workflows. A.S., Z.-M.Z, M.W.L, and Y.-S.S assisted with the computational analyses. R.J., C.F., J.R., D.A.D, J.R. and B.D. assisted with the workflow development and data collection and organization on the Cancer Genomics Cloud. R.E.B. and R.d.B. contributed to sample selection and processing of EurOPDX data. C.J.B., R.P., L.C., Y.A.E., J.H.D., S.S., M.B., C.-H.Y., E.C.-S., A.L.W, B.E.W., M.T.L., Y.X., J.W., B.F., J.R., F.M.-B., J.W., A.V.K., V.R., M.H., H.S., R.J.M., S.D., L.D., F.G., A.B., L.T., A.L., A.C.O., A.T.B., E.M., D.L., R.d.B., P.t.B., J.J., V.S., E. Marangoni, H.K., J.-I.K., H.-K.Y., C.L., E.M. and J.H.C. contributed the sequencing and array data. C.J.B., E.M. and J.H.C. directed the project. The named author list describes the primary contributors of data and analysis to the project, though these studies were supported by consortium-wide activities. All members of the PDXNet and EurOPDX Consortia participated in group discussions or supportive analyses regarding the study design, data standards, sample collection, or data analysis approaches.

## ONLINE METHODS

### Experimental details for sample collection, PDX engraftment and passaging, and array or sequencing

The tumor types and patient tumor (PT) and patient derived xenograft (PDX) samples contributed by various centers are summarized in Supplementary Fig. 1-12 and Supplementary Table 1. The sample collection, PDX engraftment and passaging, and array and sequencing methodologies by the various centers are described below.

#### The Jackson Laboratory (JAX)

Patient tumor engraftment and PDX passaging of various tumor types were performed as previously described^1–3^. Detailed information of the PDX models can be found in the PDX model search form in Mouse Tumor Biology Database (MTB, http://tumor.informatics.jax.org/mtbwi/pdxSearch.do). SNP array samples were genotyped with the Affymetrix Genome-Wide Human SNP Array 6.0 as described in Woo et al^3^. Whole-exome sequencing were processed as follows: DNA was isolated from tumor and blood samples using the Wizard Genomic DNA Purification Kit (Promega) according to the manufacturer’s protocols. DNA quality was assessed using an E-Gel General Purpose Agarose Gel, 0.8% (Invitrogen) and Nanodrop 2000 spectrophotometer (Thermo Scientific). DNA concentration was determined using a Qubit dsDNA BR Assay Kit (Thermo Scientific). Libraries were prepared by the Genome Technologies core facility at The Jackson Laboratory using SureSelectXT Reagents and SureSelectXT Human All Exon V4 Target Enrichment System (Agilent Technologies), according to the manufacturer’s instructions. Briefly, the protocol entails shearing the DNA using the Covaris E220 Focused-ultrasonicator (Covaris), ligating Illumina specific adapters, and PCR amplification. Amplified DNA libraries are then hybridized to the Human All Exon probes, amplified using indexed primers, and checked for quality and concentration using the DNA High-Sensitivity LabChip assay (Agilent Technologies) and quantitative PCR (KAPA Biosystems), according to the manufacturers’ instructions. Libraries were sequenced on a HiSeq 2500 100bp paired-end flow cell using TruSeq Rapid SBS reagents (Illumina). Average coverage for normal samples was 154.38x (115.13 min – 212.31 max), and was 232.10x for tumor samples (161.48 min – 280.65 max).

#### Seoul National University-Jackson Laboratory (SNU-JAX)

Gastric cancer tissues, paired normal gastric tissues, and blood samples were obtained from individuals who underwent gastrectomies at the Hospital of Seoul National University from 2014 to 2016. All samples were obtained with informed consent at the Hospital of Seoul National University, and the institutional review board approved the study per the Declaration of Helsinki. These samples were stored into RPMI media with 1% penicillin/streptomycin immediately after resected from patients and shipped using specimen ice box to the laboratory within half an hour. Gastric cancer samples were divided into several small pieces (2mm × 2mm) and used to generate PDX models and for genomic analysis. Mice were cared for according to institutional guidelines of the Institutional Animal Care and Use Committee of the Seoul National University (no. 14-0016-C0A0). For PDX models, surgically resected tissues were minced into pieces approximately ~2 mm in size and injected into the subcutaneous area in the flanks of 6-week-old NOD/SCID/IL-2γ-receptor null female mice (NSG^TM^ mice, Jackson Laboratory, Bar Harbor, ME). The volume of tumors and body weight of mice were checked once or twice a week. The volume was calculated as (tumor length x tumor width^2^) / 2. When a tumor reached >700~1000 mm^3^, the mouse was sacrificed, and tumor tissues were stored. Tumor tissues were divided and stored for several purposes: (1) Tumor tissues were cryopreserved in liquid nitrogen and stored at −80 °C for generating next passage PDXs. (2) Tumor tissues were frozen in liquid nitrogen for genomic analysis. Whole-exome sequencing was conducted as follows: Genomic DNA (gDNA) was extracted from blood and tissues using DNeasy blood and tissue kit (QIAGEN) and checked for purity, concentration, and integrity by OD260/280 ratio using NanoDrop Instruments (NanoDrop Technologies, Wilmington, DE, USA) and agarose gel electrophoresis. DNA was sheared by fragmentation by Bioruptor (Diagenode, Inc., Denville, NJ, USA) and purified using Agencourt AMPure XP beads (Beckman Coulter, Fullerton, CA, USA). DNA samples were then tested for size distribution and concentration using an Agilent Bioanalyzer 2100. Standard protocols were utilized for adaptor ligation, indexing, high-fidelity PCR amplification. Subsequently, exome enrichment was performed by hybrid capture with the All Exon v5 capture library. Capture libraries were amplified, pooled, and submitted to the commercial sequencing company (Macrogen) for 100bp paired-end, multiplex sequencing on a HiSeq 2000 sequencing system. Average coverage for normal samples was 62.67× (38.97 min – 108.77 max), and was 102.35x for tumor samples (36.02 min – 150.49 max). RNA-Sequencing data was generated as follows: RNA was extracted from tissues using the RNeasy Mini Kit (Qiagen, Valencia, CA, USA). RNA-Sequencing libraries were prepared from 1 μg total RNA using the TruSeq RNA Sample Preparation v2 Kit (Illumina, San Diego, CA) according to the manufacturer’s protocol. Libraries were submitted to the commercial sequencing company (Macrogen) for 100bp paired-end, multiplex sequencing on a HiSeq 2000 sequencer.

#### Huntsman Cancer Institute (HCI)

Patient tumor engraftment and PDX passaging of breast cancer samples were performed as previously described^4,5^. SNP array samples were genotyped by the Affymetrix SNP 6.0 array for profiling. These samples were processed, according to DeRose et al^5^. Additionally, some samples, were also processed using the Illumina Infinium Omni 2.5 Exome-8 v1.3 Beadchip array. Hybridized arrays were scanned using an Illumina iScan instrument following the Illumina Infinium LCG Assay Manual Protocol and processed using GenomeStudio. When samples had both Affymetrix and Illumina chips, we deferred to Illumina intensity values for copy number calling. Whole-exome sequencing was conducted as follows: Agilent SureSelectXT Human All Exon V6+COSMIC or Agilent Human All Exon 50Mb library preparation protocols were used with inputs of 100-3000ng sheared genomic DNA (Covaris). Library construction was performed using the Agilent Technologies SureSelectXT Reagent Kit. The concentration of the amplified library was measured using a Qubit dsDNA HS Assay Kit (ThermoFisher Scientific). Amplified libraries (750 ng) were enriched for exonic regions using either the Agilent Technologies SureSelectXT Human All Exon v6+COSMIC or Agilent Human All Exon 50Mb kits and PCR amplified. Enriched libraries were qualified on an Agilent Technologies 2200 TapeStation using a High Sensitivity D1000 ScreenTape assay and the molarity of adapter-modified molecules was defined by quantitative PCR using the Kapa Biosystems Kapa Library Quant Kit. The molarity of individual libraries was normalized to 5 nM, and equal volumes were pooled in preparation for Illumina sequence analysis. Sequencing libraries (25 pM) were chemically denatured and applied to an Illumina HiSeq v4 paired-end flow cell using an Illumina cBot. Hybridized molecules were clonally amplified and annealed to sequencing primers with reagents from an Illumina HiSeq PE Cluster Kit v4-cBot (PE-401-4001). Following the transfer of the flowcell to an Illumina HiSeq 2500 instrument (HCS v2.2.38 and RTA v1.18.61), a 125-cycle paired-end sequence run was performed using HiSeq SBS Kit v4 sequencing reagents (FC-401-4003). Average coverage for normal samples was 90.22× (15.28 min – 131.69 max), and was 96.66x for tumor samples (10.65 min – 166.06 max).

#### Baylor College of Medicine (BCM)

Patient tumor engraftment and PDX passaging of breast cancer samples were performed as previously described^6,7^. SNP array samples were genotyped at Huntsman Cancer Institute using the Illumina Infinium Omni 2.5Exome-8 v1.4 Beadchip array by the procedures provided in the HCI section above.

#### The University of Texas MD Anderson Cancer Center (MDACC)

Fresh non-small-cell lung carcinoma tumor samples were collected from surgically resected specimens with the informed consent of the patients. Generation and passaging of PDXs, and histological analysis and DNA fingerprint assay for PDXs and their primary tumor tissues were performed as previously described^8^. The protocols for the use of clinical specimens and data in this study were approved by the Institutional Review Board at The University of Texas MD Anderson Cancer Center. All animal studies were carried out in accordance with the Guidelines for the Care and Use of Laboratory Animals (National Institutes of Health Publication 85-23) and the institutional guidelines of MDACC. Whole-exome sequencing was conducted at the Sequencing and Microarray Core Facility at MD Anderson Cancer Center as follows: Genomic DNA was quantified and quality was assessed using Picogreen (Invitrogen) and Genomic DNA Tape for the 2200 Tapestation (Agilent), respectively. DNA from each sample (100-500 ng of genomic DNA) was sheared by sonication and then used for library preparation by using KAPA library preparation kit (KAPA) following manufacturer’s instruction. Equimolar amounts of DNA were pooled (2-6 samples per pool) and whole exome regions were captured by using biotin labeled probes from Roche Nimblegen (Exome V3) followed manufacture’s protocol. The captured libraries were sequenced on a HiSeq 2000 with 100bp paired-end (Illumina Inc., San Diego, CA, USA) on a paired-end flowcell. Average coverage for normal samples was 85.61x (40.80 min – 228.41 max), and was 125.79x for tumor samples (25.12 min – 251.53 max).

#### The WISTAR Institute (WISTAR)

Tumor biopsy samples were collected according to IRB-approved protocol with the informed written consent of the patients. Collected fresh tumor pieces were snap frozen and stored at −80 °C. Subcutaneous implantation into NSG SCID mice were used to create PDX models. BRAF inhibitor treatment (PLX) was administered as PLX4720 200ppm chemical additive diet chow (Research Diets, New Brunswick, NJ). Whole exome sequencing was conducted as follows: Genome DNA extraction was done using Qiagen DNeasy Blood & Tissue Kit, and libraries for whole exome sequencing were performed using Nextera DNA exome kit. Capture libraries were amplified, pooled, and then sequenced on an Illumina HiSeq 2500 76bp paired-end run. Average coverage for normal samples was 97.50× (71.46 min – 124.64 max), and was 208.27x for tumor samples (146.88 min – 281.20 max).

#### National Cancer Institute Patient-Derived Models Repository (PDMR)

For engraftments, tumor material plus a drop of Matrigel (BD BioSciences, Bedford, MA) were implanted subcutaneously in NSG^TM^ mouse model NOD.Cg-Prkdc^scid^ Il2rg^tm1Wjl^/SzJ. Mice were housed in sterile, filter-capped polycarbonate cages, maintained in a barrier facility on a 12-hour light/dark cycle, and were provided sterilized food and water, ad libitum. Animals were monitored weekly for tumor growth. The initial passage of material was grown to approximately 1000-2000mm^3^ calculated using the following formula: weight (mg) = (tumor length x [tumor width]^2^) / 2. Tumor material was then harvested, a portion cryopreserved, and the remainder implanted into NSG host mice. Every PDX tumor harvested and cryopreserved also has 2-3 fragments snap frozen for next generation sequence analysis and short tandem repeat validation and a piece is fixed in neutral buffered formalin and then embedded in paraffin for histological assessment. Related patient data, clinical history, representative histology and short-tandem repeat profiles for the PDX models can be found at https://pdmr.cancer.gov. Full PDMR standard operating procedures for tumor engraftment and PDX passaging are available at https://pdmr.cancer.gov/sops. Whole-exome sequencing data were generated with the Agilent SureSelect capture kit, and sequenced with 125bp pair-end Illumina HiSeq 2500 runs following standard operating procedures available here: https://pdmr.cancer.gov/sops. Average coverage for normal samples was 148.47x (50.95 min – 242.24 max), and was 174.77x for tumor samples (81.41 min – 403.22 max).

#### Washington University in St. Louis (WUSTL)

All human tissues acquired for these experiments were processed in compliance with NIH regulations and institutional guidelines, approved by the Institutional Review Board at Washington University. Tumors from all patients were obtained via core needle biopsy, skin punch biopsy, or surgical resection after informed consent. All animal procedures were reviewed and approved by the Institutional Animal Care and Use Committee at Washington University in St. Louis. Pancreatic cancer models were derived from tissue fragments implanted subcutaneously into dorsal flank regions of non-humanized, female NOD/SCID/γ mice (Jackson Laboratory, Bar Harbor, ME) using Matrigel. The sample tissues for these PDX models were obtained from archived, cryopreserved PDX harvests. Final tumor passages in mice were kept cold and harvested into RPMI-1640 with antibiotic and antimycotic additives. Pieces of each tumor were processed into the following: flash frozen tissue fragments, OCT blocks and matched Haemotoxylin and Eosin (H&E) slides, formalin fixed paraffin blocks and matched H&E slides, RNAlater tissue storage, and cryopreserved fragments (FBS + 10% DMSO). A minimum of 250 mg of flash frozen material was submitted to the Siteman Cancer Center’s Proteomics Core. The tissues were cryo-pulverized and subsequently divided for DNA and RNA preparation, and long-term storage. Patient tumors were obtained directly from operating rooms and placed into sterile collection media (RPMI-1640 with antibiotic and antimycotic additives). Pieces of each tumor were processed into the following: flash frozen tissue fragments, OCT blocks and matched H&E slides, formalin fixed paraffin blocks and matched H&E slides, and cryopreserved fragments (FBS + 10% DMSO). Parental genomic DNA was prepared from OCT blocks if available, and if not available, paraffin blocks were utilized. In addition, genomic DNA for sequencing control was prepped from peripheral blood mononuclear cells that were both procured and processed at time of surgery. Breast cancer models were derived from tissue fragments implanted subcutaneously into dorsal flank regions of non-humanized, NOD/SCID/γ mice (Jackson Laboratories, Bar Harbor, ME) as previously described^7,9^. Whole-exome sequencing was conducted as follows: Libraries were constructed using unamplified genomic DNA (minimum 100 ng) from blood (normal), tumor, and xenograft samples. Exons were captured via IDT Exome library kit followed by high-throughput sequencing on an Illumina NovaSeq S4 platform (Illumina Inc., San Diego, CA) using 150bp paired-end reads. Details of whole exome library construction have been given elsewhere (Fisher, Barry et al. 2011). Average coverage for normal pancreatic cancer samples was 85.73x (55.65 min – 108.91 max), and was 124.01x (49.68 min – 242.35 max) for tumor pancreatic cancer samples. Average coverage for normal breast cancer samples was 58.33x (45.37 min – 70.30 max), and was 89.90x (17.24 min – 149.53 max) for tumor breast cancer samples.

#### Shanghai Institute for Biological Sciences (SIBS)

Gene expression and copy number data, generated by the Affymetrix Human Genome U133 Plus 2.0 Array and Affymetrix Human SNP 6.0 platforms respectively, of hepatocellular carcinoma (HCC) PDX models were retrieved from the Gene Expression Omnibus (GEO) accession ID GSE90653^10^. Expression microarray data generated by the Affymetrix Human Genome U133 Plus 2.0 Array for normal liver were downloaded from GEO and ArrayExpress: GSE3526^11^, GSE33006^12^ and E-MTAB-1503-3^13^.

#### EurOPDX colorectal cancer (EuroPDX CRC)

Liver-metastatic colorectal cancer samples were obtained from surgical resection of liver metastases at the Candiolo Cancer Institute, the Mauriziano Umberto I Hospital, and the San Giovanni Battista Hospital. Informed consent for research use was obtained from all patients at the enrolling institution before tissue banking, and study approval was obtained from the ethics committees of the three centers. Tissue from hepatic metastasectomy in affected individuals was fragmented and either frozen or prepared for implantation as described previously^14,15^. Non-obese diabetic/severe combined immunodeficient (NOD/SCID) female mice (4–6 weeks old) were used for tumor implantation. Snap-frozen aliquots were obtained from surgical specimens and corresponding tumor grafts at different passages. Whole genome sequencing was conducted as follows: DNA was extracted using Maxwell RSC Blood DNA kit (Promega AS1400) from colorectal cancer liver metastasis and corresponding tumor grafts at different passages. Genomic DNA was fragmented and used for Illumina TruSeq library construction (Illumina) according to the manufacturer’s instructions. Libraries were then purified with Qiagen MinElute column purification kit and eluted in 17 µl of 70°C EB to obtain 15 µl of DNA library. The libraries were sequenced on HiSeq4000 (Illumina) with single-end reads of 51bp at low coverage (~0.1x genome coverage on average).

#### EurOPDX breast cancer (EuroPDX BRCA)

Human breast tumors were obtained from surgical resections at the Netherland Cancer Institute (NKI), Institut Curie (IC) and Vall d’Hebron Institute of Oncology (VHIO). Engraftment was conducted with different procedures at each center. <u>NKI</u>: Small tumor fragments (2mm diameter) were implanted into the 4th mammary fat pad of 8-week-old Swiss female nude mice. Mice were checked for tumor appearance once a week, and supplemented with estrogen, if the tumor was ER positive. After palpable tumor detection, tumor size was measured twice a week. When tumors reached a size of 700-1000 mm^3^, animals were sacrificed and tumors were explanted and subdivided in fragments for serial transplantation as described above, or for frozen vital storage in liquid nitrogen. <u>IC</u>: Breast cancer fragments were obtained from patients at the time of surgery, with informed written patient consent. Fragments of 30 to 60 mm^3^ were grafted into the interscapular fat pad of 8 to 12-week-old female Swiss nude mice. Mice were supplemented with estrogen. Xenografts appeared at the graft site 2 to 8 months after grafting. When tumors were close to 1500 mm^3^, they were subsequently transplanted from mouse to mouse and stocked frozen in DMSO-fetal calf serum (FCS) solution or frozen dried in nitrogen. Fragment fixed tissues in phosphate buffered saline (PBS) 10% formol for histologic studies were also stored. The experimental protocol and animal housing were in accordance with institutional guidelines as proposed by the French Ethics Committee (Agreement B75-05-18, France). <u>VHIO</u>: Fresh tumor samples from patients with breast cancer were collected for implantation following an institutional IRB-approved protocol and the associated informed consent, or by the National Research Ethics Service, Cambridgeshire 2 REC (REC reference number: 08/H0308/178). Experiments were conducted following the European Union’s animal care directive (2010/63/EU) and were approved by the Ethical Committee of Animal Experimentation of the Vall d’Hebron Research Institute. Surgical or biopsy specimens from primary tumors or metastatic lesions were immediately implanted in mice. Fragments of 30 to 60 mm^3^ were implanted into the mammary fat pad (surgery samples) or the lower flank (metastatic samples) of 6-week-old female athymic HsdCpb:NMRI-Foxn1nu mice (Harlan Laboratories). Animals were continuously supplemented with estradiol. Upon growth of the engrafted tumors, the model was perpetuated by serial transplantation onto the lower flank. Tumor growth was measured with caliper bi-weekly. In all experiments, mouse weight was recorded twice weekly. When tumors reached 1500 mm3, mice were euthanized and tumors were explanted. Whole genome sequencing was conducted as follows: genomic DNA was extracted from breast cancers and corresponding PDXs using (i) QIAamp DNA Mini Kit s(50) (#51304, Qiagen) (IC) or (ii) according to Laird PW’s protocol^16^ (NKI and VHIO). The amount of double stranded DNA in the genomic DNA samples was quantified by using the Qubit® dsDNA HS Assay Kit (Invitrogen, cat no Q32851). Up to 2000 ng of double stranded genomic DNA were fragmented by Covaris shearing to obtain fragment sizes of 160-180bp. Samples were purified using 1.6X Agencourt AMPure XP PCR Purification beads according to manufacturer’s instructions (Beckman Coulter, cat no A63881). The sheared DNA samples were quantified and qualified on a BioAnalyzer system using the DNA7500 assay kit (Agilent Technologies cat no. 5067-1506). With an input of maximum 1 µg sheared DNA, library preparation for Illumina sequencing was performed using the KAPA HTP Library Preparation Kit (KAPA Biosystems, KK8234). During library enrichment, 4-6 PCR cycles were used to obtain enough yield for sequencing. After library preparation the libraries were cleaned up using 1X AMPure XP beads. All DNA libraries were analyzed on the GX Caliper (a PerkinElmer company) using the HT DNA High Sensitivity LabChip, for determining the molarity. Up to two pools of 24 uniquely indexed samples and one pool of 81 uniquely indexed samples were mixed together by equimolar pooling in a final concentration of 10nM, and subjected to sequencing on an Illlumina HiSeq2500 machine in a total of 12 lanes of a single read 65bp run at low coverage (~0.4x genome coverage on average), according to manufacturer’s instructions.

### Consolidating tumor types from different datasets

As the terminology of tumor types/subtypes by the different contributing centers were not consistent, we used the Disease Ontology database^17^ (http://disease-ontology.org/), cancer types listed in NCI website (https://www.cancer.gov/types) and in TCGA publications^18,19^ to unify and group the tumor types/subtypes under broader terms as shown in Fig.1 and Supplementary Table 2.

### Copy number alteration (CNA) estimation methods

#### SNP array

The estimation of CNA profiles from SNP array were detailed previously^3^. In short, for Affymetrix Human SNP 6.0 arrays, PennCNV-Affy and Affymetrix Power Tools^20^ were used to extract the B-allele frequency (BAF) and Log R Ratio (LRR) from the CEL files. Due to the absence of paired-normal samples, the allele-specific signal intensity for each PDX tumor were normalized relative to 300 randomly selected sex-matched Affymetrix Human SNP 6.0 array CEL files obtained from the International HapMap project^21^. For Illumina Infinium Omni2.5Exome-8 SNP arrays (v1.3 and v1.4 kit), the Illumina GenomeStudio software was used to extract the B-allele frequency (BAF) and Log R Ratio (LRR) from the signal intensity of each probe. The single sample mode of the Illumina GenomeStudio was used, which normalizes the signal intensities of the probes with an Illumina in-house dataset. The single tumor version of ASCAT^22^ (v2.4.3 for JAX SNP data, v2.5.1 for SIBS SNP data) was used for GC correction, predictions of the heterozygous germline SNPs based on the SNP array platform, and estimation of ploidy, tumor content and allele-specific copy number segments. The resultant copy number segments were annotated with log_2_ ratio of total copy number relative to predicted ploidy from ASCAT.

#### Whole-exome sequencing (WES) data

All the samples were subjected to quality control (filtering and trimming of poor-quality reads and bases) using in-house QC script with the cut-off that half of the read length should be ≥20 in base quality at phred scale. We further removed the known adaptors using cut-adapt^23^ v1.15 11 at -m 36. Afterward, we aligned the reads to the human genome (GRCh38.p5) using bwakit^24^ v0.7.15. Engrafted tumor samples were subjected to the additional step of mouse read removal using Xenome^25^ v1.0.0, with default parameters. The alignment was converted to BAM format using Picard SortSam v2.8.1 (https://broadinstitute.github.io/picard/), and duplicates were removed by Picard MarkDuplicates utility. BaseRecalibrator from the Genome Analysis Tool Kit^26,27^ (GATK) v4.0.5.1 was used to adjust the quality of raw reads. Training files for the base quality scale recalibration were Mills_and_1000G_gold_standard.indels.hg38.vcf.gz, Homo_sapiens_assembly38.known_indels.vcf.gz, and dbSNP v151. Mean target coverage was determined for each sample by Picard CollectHsMetrics. Aligned bams were subset to target region by GATK and SAMTools^28^ v0.1.18 was used to generate the pileup for each sample. Pileup data were used for CNA estimation as calculated with Sequenza^29^ v2.1.2. Both tumor and normal data, that utilized the same capture array, were used as input. pileup2seqz and GC-windows (-w 50) modules from sequenza-utils.py utility were used to create the native seqz format file for Sequenza and compute the average GC content in sliding windows from hg38 genome, respectively. Finally, we ran the three Sequenza modules with these modified parameters (sequenza.extract: assembly = “hg38”, sequenza.fit: chromosome.list = 1:23, and sequenza.results: chromosome.list = 1:23) to estimate the segments of copy number gains/losses. Finally, segments lacking read counts, in which ≥50% of the segment with zero read coverage, were removed. A reference implementation of this workflow (Supplementary Fig. 77) is developed and deployed in the cancer genomics cloud at SevenBridges (https://cgc.sbgenomics.com/public/apps#pdxnet/pdx-wf-commit2/wes-cnv-tumor-normal-workflow/, https://cgc.sbgenomics.com/public/apps#pdxnet/pdx-wf-commit2/pdx-wes-cnv-xenome-tumor-normal-workflow/).

#### Low-pass whole-genome sequencing (WGS) data

Whole-genome sequence reads from EuroPDX CRC liver metastasis and corresponding tumor grafts at different passages were mapped to the reference human genome (GRCh37) using Burrows-Wheeler Aligner^24^ (BWA) v0.7.12. SAMTools^28^ v0.1.18 was used to convert SAM files into BAM files and Picard v1.43 to remove PCR duplicates (http://broadinstitute.github.io/picard/). Raw copy number profiles for each sample were estimated by QDNAseq^30^ R package v1.20 by dividing the human reference genome in non-overlapping 50 kb windows and counting the number of reads in each bin. Bins in problematic regions were removed^31^. Read counts were corrected for GC content and mappability by a LOESS regression, median-normalized and log_2_-transformed. Values below –1000 in each chromosome were floored to the first value greater than –1000 in the same chromosome. Raw log_2_ ratio values were then segmented using the ASCAT^22^ algorithm implemented in the ASCAT R package v2.0.7. Whole-genome sequence reads from EuroPDX BRCA tumors and corresponding tumor grafts at different passages were mapped to the reference human genome (GRCh38) and mouse genome (GRCm38/mm10, Ensembl 76) using Burrows-Wheeler Aligner (BWA) v0.7.15. Subsequently, mouse reads were excluded with XenofilteR^32^. Other processing steps are similar as described above. Raw copy number profiles were estimated for each sample by dividing the human reference genome in non-overlapping 20 kb windows and counting the number of reads in each bin. Only reads with at least mapping quality 37 were considered. Bins within problematic regions (i.e. multimapper regions) were excluded. Downstream analysis to estimate copy number was conducted as described above.

#### RNA-sequencing (RNA-Seq) and gene expression microarray (EXPARR) data

For SNU-JAX RNA-Seq data, Simultaneous read alignment was performed to both mouse (mm10) and human genome (GRCh38.p5) and only human specific reads were used for the expression quantification. Expression of mRNA was quantified as Transcripts Per Million (TPM) for downstream analysis using RNA-Seq by Expectation Maximization^33^ (RSEM) with ensemble GTF reference GRCh38.92. Gene expression microarray data for SIBS HCC and normal liver samples from GEO and ArrayExpress databases were profiled as follows. After initial quality control and outlier removal, CEL files were normalized according to RMA algorithm and probesets were annotated according to Affymetrix annotation file for HG-U133 Plus 2, released on 2016-03-15 build 36. For expression-based copy number inference, we referred to the previous protocols for e-karyotyping and CGH-Explorer^34–37^. For each cancer type, expression values of tumor and corresponding normal samples were merged in a single table, and gene identifiers were annotated with chromosomal nucleotide positions. Genes located on sex chromosomes were excluded. Genes which values below 1 TPM (RNAseq) or probeset log_2_-values below 6 (microarray) in more than 20% of the analyzed dataset were removed. Remaining gene expression values below the thresholds were respectively raised to 1 TPM or log_2_-value of 6. In the case of multiple transcripts (RNA-seq) or probesets (microarray) per gene, the one with the highest median value across the entire dataset was selected. According to the e-karyotyping protocol, the sum of squares of the expression values relative to their median expression across all samples was calculated for each gene, and 10% most highly variable genes were removed. For each gene, the median log_2_ expression value in normal samples was subtracted from the log2 expression value in each tumor sample and subsequently input in CGH-explorer. For tumor-only datasets, the median log_2_ expression value in the same set of tumor samples was instead subtracted. The preprocessed expression profiles of each sample were individually analyzed using CGH-Explorer (http://heim.ifi.uio.no/bioinf/Projects/CGHExplorer/). CGH-PCF analysis was carried out to call copy number according to parameters previously reported^38^: least allowed deviation = 0.25; least allowed aberration size = 30; winsorize at quantile = 0.001; penalty = 12; threshold = 0.01.

### Filtering and gene annotation of copy number segments

Copy number (CN) segments with log_2_ copy number ratio estimated from the various platforms were processed in the following steps (Supplementary Fig. 26). Segments <1kb were filtered based on the definition of CNA^39^. In addition, SNP array segments had to be covered by >10 probes, with an average probe density of 1 probe per 5kb. The copy number segments were then binned into 10kb windows to derive the median log_2_(CN ratio), which was subsequently used to re-center the copy number segments. Median-centered copy number segments were visualized using IGV^40^ v2.4.13 and GenVisR^41^ v1.16.1. Median-centered copy number of genes were calculated by intersecting the genome coordinates of copy number segments with the genome coordinates of genes (Ensembl Genes 93 for human genome assembly GRCh38, Ensembl Genes 96 for human genome assembly GRCh37). In the case where a gene overlaps multiple segments, the most conservative (lowest) estimate of copy number was used to represent the copy number of the entire intact gene.

### Comparison of CN gains and losses

For the comparison of resolution, range of CN values and frequency of gains and losses between different platforms and analysis methods, we defined copy number gain or loss segments as – Gain: log_2_(CNratio) > 0.1; Loss: log_2_(CN ratio) < -0.1.

### Correlation of CNA profiles

The overall workflow to compare CNA profiles is shown in Supplementary Fig. 26. PDX samples without passage information were omitted in the following downstream analysis. The copy number segments were binned into 10kb-windows or smaller using Bedtools^42^ v2.26.0, and the variance of log_2_(CN ratio) and range (difference) of log_2_(CN ratio) between 5^th^ to 95^th^ percentile across all the bins were calculated as a measure of degree of aberration for each CNA profile. A non-aberrant profile results in a low variance or range. While variance can be biased for CNA profiles with small segments of extreme gains or losses, we preferred the use of 5^th^ to 95^th^ percentile range to identify samples with low degree of aberration, such that a narrow range indicates ≥90% of the genome has very low-level gains and losses. The similarity of two CNA profiles is quantified by the Pearson correlation coefficient of log_2_(CN ratio) of 100kb-windows binned from segments or genes between 2 samples. Gene-based and segment-based (100kb windows) correlations were highly similar (data not shown). Using correlation avoided the issue of making copy number gain and loss calls based on thresholds, though it can be inconsistent due to different baseline and range in copy number values. Such variations are impacted by sample-specific variation in human stromal contamination or sensitivity copy number detection by different platforms.

#### Comparison of CNA profiles between different platforms

The copy number segments of each pair of data were intersected and binned into 100kb-windows or smaller using Bedtools. The Pearson correlation coefficient and linear regression model was calculated for the log_2_(CN ratio) of the windows. Windows with discrepant copy number were identified by outliers of the linear regression model defined by |studentized residual| > 3. These outlier windows were mapped to their corresponding segments to identify the size of CNA events that were discordant between the different copy number estimation methods. The proportion of the genome discordant CNA was calculated from the summation of the outlier windows.

#### Identification of genes with CNA between different samples of the same model

To compare the CNA profiles between different samples (PT or PDX) of the same model, the Pearson correlation coefficient and linear regression model was calculated for the log_2_(CN ratio) of the genes for each pair of data. Prior to that, deleted genes with log_2_(CN ratio) < -3 were rescaled to -3 to avoid large shifts in the correlation coefficient and linear regression model due to extremely negative values on the log scale. Extreme outliers of the linear regression model defined by |studentized residual| > 3 were removed to derive an improved linear regression model^43^ not biased by few extreme values. Genes with copy number changes between the samples were identified by the difference in log_2_(CN ratio) relative to the improved linear regression model of |standard residual| < 0.5. We also removed some samples with low correlation due to sample mislabeling as they displayed high correlation with samples from other models. We also omit samples with low correlation values (<0.6) which resulted from non-aberrant CNA profiles in genomically stable tumors (5^th^ to 95^th^ percentile range < 0.3, Supplementary Fig. 64).

#### Identification of aberrant sample pairs with highly discordant CNA profiles

Aberrant CNA profiles were identified based on the 100kb-window copy number range (5th to 95th percentile) >0.5, for both samples. Sample pairs with Pearson correlation <0.6 were selected as highly discordant CNA profiles between them.

### Annotation with gene sets with known cancer or treatment-related functions

Copy number altered genes (|residual| < 0.5) were annotated by various gene sets with cancer or treatment-related functions gathered from various databases and publications (Supplementary Fig. 26):

1. Genes in 10 oncogenic signaling pathways curated by TCGA and were found to be frequently altered in different cancer types^44^.
2. Genes with gain in copy number or expression, or loss in copy number or expression that conferred therapeutic sensitivity, resistance or increase/decrease in drug response from the JAX Clinical Knowledgebase^45,46^ (JAX-CKB) based on literature curation (https://ckbhome.jax.org/, as of 06-18-2019).
3. Genes with evidence of promoting oncogenic transformation by amplification or deletion from the Cancer Gene Census^47^ (COSMIC v89).
4. Significantly amplified or deleted genes in TCGA cohorts of breast cancer^48^, colorectal cancer^49^, lung adenocarcinoma^50^ and lung squamous cell carcinoma^51^ by GISTIC analysis.

### Identification of genes with recurrent copy number changes

Genes with a more stringent threshold of |residual| > 1.0 with respect to the improved regression linear model (without discriminating gain or loss) were selected for each pairwise comparison between different samples of the same model. Pairwise cases in which genes are deleted in both samples (log_2_(CN ratio) ≤ −3) are omitted. Recurrent frequency for each gene across all models was calculated on a model basis such that genes with copy number between multiple pairs of the same model was counted as once. This avoided the bias towards models with many samples of similar copy number changes between the different pairs.

### Drug response analysis using CCLE data

We developed a pipeline to evaluate gene copy number effects on drug sensitivity^52,53^ by using the Cancer Cell Line Encyclopedia^54,55^ (CCLE) cell line genomic and drug response data (CTRP v2). We downloaded the CCLE drug response data from Cancer Therapeutics Response Portal (www.broadinstitute.org/ctrp), and CCLE gene-level CNA and gene expression data from depMap data portal (‘public_19Q1_gene_cn.csv’ and ‘CCLE_depMap_19Q1_TPM.csv’, https://depmap.org/portal/download/). For CCLE drug response data, we used the area-under-concentration-response curve (AUC) sensitivity scores for each cancer cell line and each drug. In total, we collected gene-level log_2_ copy number ratio data derived from the Affymetrix SNP 6.0 platform from 668 pan-cancer CCLE cell lines, with a total of 545 cancer drugs tested. With the CCLE gene-level CNA and AUC drug sensitivity scores, we performed gene-drug response association analyses for genes with recurrent copy number changes. Pearson correlation p-values between each gene’s log_2_ (CN ratio) and each drug’s AUC score across all cell lines were calculated, and q-values were calculated by multiple testing Bonferroni correction. Significant gene-CNA and drug associations were kept (q-value < 0.1) to further evaluate gene-expression and drug response associations. If a gene’s expression was also significantly correlated with AUC drug sensitivity scores, particularly in the same direction (either positively or negatively correlated) as the gene-CNA and drug association, that gene would be considered as significantly correlated with drug response based on both its CNA and gene expression.

### GISTIC analysis of WGS data

To obtain perfectly matching and comparable PT–PDX cohorts, for GISTIC analysis, CRC trios in which at least one sample did not display significant CNAs were excluded from the analysis resulting in a total of 87 triplets. The GISTIC^56^ algorithm (GISTIC 2 v6.15.28) was applied on the segmented profiles using the GISTIC GenePattern module (https://cloud.genepattern.org/), with default parameters and genome reference files Human_Hg19.mat for EuroPDX CRC data and hg38.UCSC.add_miR.160920.refgene.mat for EuroPDX BRCA data. For each dataset, GISTIC provides separate results (including segments, G-scores and FDR q-values) separately for recurrent amplifications and recurrent deletions. Deletion G-scores were assigned negative values for visualization. We observed that the G-Score range was systematically lower in PT cohorts, which is likely the result of the dilution of CNA by normal stromal DNA. In contrast, human stromal DNA in PDX samples were lower or negligible. To account for this difference in gene-level G-scores, PDXs at early and late passages were scaled with respect to PT gene-level G-score values using global linear regression, separately for amplification and deletion outputs.

### Gene set enrichment analysis (GSEA) of WGS data

To assess the biological functions associated with the recurrent alterations detected by the GISTIC analysis, we performed GSEAPreranked analysis^57,58^ on gene-level GISTIC G-score profiles, for both amplifications and deletions. In particular, we applied the algorithm with 1000 permutations on various gene set collections from the Molecular Signatures Database^59,60^ (MSigDB): H (Hallmark), C2 (Curated : CGP chemical and genetic perturbations, CP canonical pathways), C5 (Gene Ontology: BP biological process, MF molecular function, CC cellular component) and C6 (Oncogenic Signatures) composed of 50, 4762, 5917 and 189 gene sets respectively. We also included gene sets with known cancer or treatment-related functions described in an earlier section. We noted that multiple genes with contiguous chromosomal locations, typically in recurrent amplicons, generated spurious enrichment for gene sets which consists of multiple genes of adjacent positions, while very few or none of them had a significant GISTIC G-score. To avoid this confounding issue, we only considered the “leading edge genes”, i.e. those genes with increasing Normalized Enrichment Score (NES) up to its maximum value, that contribute to the GSEA significance for a given gene set. The leading-edge subset can be interpreted as the core that accounts for the gene set’s enrichment signal (http://software.broadinstitute.org/gsea). We included a requirement that the leading edge genes passing the GISTIC G-score significant thresholds based on GISTIC q-value 0.25 (Supplementary Table 8 and Fig. 73) make up at least 20% of the gene set. This 20% threshold was chosen as the minimal threshold at which gene sets assembled from TCGA-generated lists of genes with recurrent CNA in CRC or BRCA were identified as significant in GSEA (see Supplementary Table 9). Finally, gene sets with a NES greater than 1.5 and a FDR q-value of less than 0.05, which passed the leading edge criteria, were considered significantly enriched in genes affected by recurrent CNAs.

