## Supplementary Figures and Tables 1-4 for "Conservation of copy number profiles during engraftment and passaging of patient-derived cancer xenografts"

Supplementary Figure 1: Summary of PDX models collected from various centers in the PDXNET and EuroPDX consortium, and publicly available data.

Supplementary Figure 2: Number of patient tumor and PDX passages per model in the JAX PDX resource consisting of various tumor types assayed by Affymetrix SNP 6.0 array and whole exome sequencing (P: Primary malignancy, M: Metastatic, R: Recurrent/Relapse, NS: Not specified).

Supplementary Figure 3: Number of patient tumor and PDX passages per model in SNU-JAX gastric cancer dataset assayed by whole-exome sequencing (WES) and RNA sequencing (RNASEQ). Models labeled with ”LN” are lymph node metastatic tumors for the same patient. (*: Multiple patient tumors available for the same patient, different relapse time points or different metastatic sites)

Supplementary Figure 4: Number of patient tumor and PDX passages per model in the HCI breast cancer dataset assayed by whole-exome sequencing (WES), Affymetrix SNP 6.0 array and Illumina Infinium Omni2.5Exome8 (v1.3) SNP array. HCI-007 is relapse tumor from the patient of HCI-005, HCI-024 is a skin metastasis tumor from the patient of HCI-023. Samples labeled with “PDX” indicates passage number is unknown. (*: Multiple tumors available for the same patient, different relapse time points or different metastatic sites)

Supplementary Figure 5: Number of PDX passages per model in the BCM breast cancer dataset assayed by Illumina Infinium Omni2.5Exome8 (v1.4) SNP array.

Supplementary Figure 6: Number of patient tumor and PDX passages per model in the MDACC lung cancer dataset assayed by whole-exome sequencing (WES).

Supplementary Figure 7: Number of patient tumor and PDX passages per model in Wistar skin cutaneous melanoma dataset assayed by whole-exome sequencing (WES). PDX samples labeled with ”PLX” are BRAF inhibitor (PLX) treated. (*: Multiple tumors available for the same patient, different relapse time points or different metastatic sites)

Supplementary Figure 8: Number of patient tumor and PDX passages per model in the NCI PDMR resource consisting of various tumor types assayed by whole exome sequencing.

Supplementary Figure 9: Number of patient tumor and PDX passages per model in the WUSTL breast cancer and pancreatic cancer datasets assayed by whole-exome sequencing (WES).

Supplementary Figure 10: Number of patient tumor and PDX passages per model in the SIBS hepatocellular carcinoma dataset assayed by Affymetrix SNP 6.0 array and Affymetrix gene expression array (EXPARR).

Supplementary Figure 11: Number of patient tumor and PDX passages per model in the EuroPDX colorectal cancer (CRC) liver metastasis dataset assayed by whole-genome sequencing.

Supplementary Figure 12: Number of patient tumor and PDX passages per model in the EuroPDX breast cancer (BRCA) dataset assayed by whole-genome sequencing.

Supplementary Figure 13: CNA profiles for matched patient tumor samples estimated from SNP array and WES for ”SNP vs WES” benchmarking (see Supplementary Table 3).

Supplementary Figure 14: CNA profiles for (**a**) patient tumor and (**b**) PDX samples estimated from WES used for ”WES vs RNASEQ (NORM/TUM)” benchmarking (see Supplementary Table 3).

Supplementary Figure 15: CNA profiles for (**a**) patient tumor and (**b**) PDX samples estimated from RNA-Seq, normalized by median expression of normal samples of the same tumor type, used for ”WES vs RNASEQ (NORM)” and “RNASEQ NORM vs TUM” benchmarking. CNA profiles for (**c**) patient tumor and (**d**) PDX samples estimated from RNA-Seq, normalized by median expression of same set of patient tumors, used for ”WES vs RNASEQ (TUM)” and “RNASEQ NORM vs TUM” benchmarking (see Supplementary Table 3).

Supplementary Figure 16: CNA profiles for patient tumor and PDX samples estimated from SNP array used for ”SNP vs EXPARR (NORM/TUM)” benchmarking (see Supplementary Table 3).

Supplementary Figure 17: CNA profiles for patient tumor and PDX samples estimated from gene expression array, normalized by (**a**) median expression of normal samples of the same tumor type and (**b**) median expression of same set of patient tumors, used for ”SNP vs EXPARR (NORM/TUM)” and “EXPARR NORM vs TUM” benchmarking (see Supplementary Table 3).

Supplementary Figure 18: Heatmap representing the Pearson correlation coefficients of the $\log_2$(CN ratio) of 100kb-windows binned from copy number segments of CNA profiles between matched samples estimated from SNP array and WES. The variance and range (5 – 95 percentile) values were calculated from the $\log_2$(copy number ratio) across all 100kb-windows per sample

Supplementary Figure 19: (**a**) Pearson correlation and linear regression of the $\log_2$(CN ratio) of 100kb-windows binned from copy number segments, of CNA profiles between matched patient tumor samples estimated from SNP array and WES. Outliers of the linear regression (red points) are identified by studentized residuals > 3 and < -3. (**b**) Comparison of segment sizes between the combined outlier and non-outliers in (**a**).

Supplementary Figure 20: Frequencies of copy number gains (log2(CN ratio) > 0.1) and losses (log2(CN ratio) < –0.1) estimated from RNA-Seq and gene expression array normalized by median expression of normal samples of the same tumor type (RNASEQ NORM, EXPARR NORM) or median expression of same set of patient tumors (RNASEQ TUM, EXPARR TUM) (see Supplementary Table 3).

Supplementary Figure 21: Heatmap representing the Pearson correlation coefficients of the $\log_2$(CN ratio) of 100kb-windows binned from copy number segments of CNA profiles estimated from (**a**) RNA-Seq (RNASEQ NORM vs TUM) and (**b**) gene expression array (EXPARR NORM vs TUM), between matched samples normalized by median expression of normal samples of the same tumor type and median expression of same set of patient tumors. The variance and range (5 – 95 percentile) values were calculated from the $\log_2$(copy number ratio) across all 100kb-windows per sample.

Supplementary Figure 22: Heatmap representing the Pearson correlation coefficients of the $\log_2$(CN ratio) of 100kb-windows binned from copy number segments of CNA profiles between matched samples estimated from WES and RNA-Seq, (**a**) normalized by median expression of normal samples of the same tumor type “WES vs RNASEQ (NORM)” or (**b**) median expression of same set of patient tumors “WES vs RNASEQ (TUM)” . The variance and range (5 – 95 percentile) values were calculated from the $\log_2$(copy number ratio) across all 100kb-windows per sample.

Supplementary Figure 23: Heatmap representing the Pearson correlation coefficients of the $\log_2$(CN ratio) of 100kb-windows binned from copy number segments of CNA profiles between matched samples estimated from SNP array and gene expression microarray, (**a**) normalized by median expression of normal samples of the same tumor type ”SNP vs EXPARR (NORM)” or (**b**) median expression of same set of patient tumors ”SNP vs EXPARR (TUM)”. The variance and range (5 – 95 percentile) values were calculated from the $\log_2$(copy number ratio) across all 100kb-windows per sample.

Supplementary Figure 24: Comparison of segment sizes between the combined outlier and non-outliers in (**a**) WES vs RNASEQ (NORM), (**b**) WES vs RNASEQ (TUM), (**c**) SNP vs EXPARR (NORM), and (**d**) SNP vs EXPARR (TUM) (See Supplementary Table 3).

Supplementary Figure 25: Pearson correlation and linear regression of the $\log_2$(CN ratio) of 100kb-windows binned from copy number segments, of CNA profiles between matched patient tumor samples estimated from different platforms and analysis methods for examples shown in Fig. 2d. Outliers of the linear regression (red points) are identified by studentized residuals > 3 and < -3.

Supplementary Figure 26: A correlation and robust regression approach to quantify similarity of CNA profiles and identify genes with copy number changes between two samples.

Supplementary Figure 27: CNA profiles (IGV heatmap) and correlation heatmap of gene-based copy number (log2(CN ratio), median centered) of samples from JAX SNP array bladder cancer dataset.

Supplementary Figure 28: CNA profiles (IGV heatmap) and correlation heatmap of gene-based copy number (log2(CN ratio), median centered) of samples from JAX SNP array breast cancer dataset.

Supplementary Figure 29: CNA profiles (IGV heatmap) and correlation heatmap of gene-based copy number (log2(CN ratio), median centered) of samples from JAX SNP array colorectal cancer dataset.

Supplementary Figure 30: CNA profiles (IGV heatmap) and correlation heatmap of gene-based copy number (log2(CN ratio), median centered) of samples from JAX SNP array glioblastoma multiforme (GBM) dataset.

Supplementary Figure 31: CNA profiles (IGV heatmap) and correlation heatmap of gene-based copy number (log2(CN ratio), median centered) of samples from JAX SNP array lung adenocarcinoma (LUAD) dataset.

Supplementary Figure 32: CNA profiles (IGV heatmap) and correlation heatmap of gene-based copy number (log2(CN ratio), median centered) of samples from JAX SNP array lung squamous cell carcinoma (LUSC) dataset.

Supplementary Figure 33: CNA profiles (IGV heatmap) and correlation heatmap of gene-based copy number (log2(CN ratio), median centered) of samples from JAX SNP array other lung cancer subtypes dataset.

Supplementary Figure 34: CNA profiles (IGV heatmap) and correlation heatmap of gene-based copy number (log2(CN ratio), median centered) of samples from JAX SNP array skin melanoma dataset.

Supplementary Figure 35: CNA profiles (IGV heatmap) and correlation heatmap of gene-based copy number (log2(CN ratio), median centered) of samples from JAX SNP array ovarian cancer dataset.

Supplementary Figure 36: CNA profiles (IGV heatmap) and correlation heatmap of gene-based copy number (log2(CN ratio), median centered) of samples from JAX SNP array sarcoma dataset.

Supplementary Figure 37: CNA profiles (IGV heatmap) and correlation heatmap of gene-based copy number (log2(CN ratio), median centered) of samples from JAX SNP array other cancers dataset.

Supplementary Figure 38: CNA profiles (IGV heatmap) and correlation heatmap of gene-based copy number (log2(CN ratio), median centered) of samples from BCM SNP array breast cancer dataset.

Supplementary Figure 39: CNA profiles (IGV heatmap) and correlation heatmap of gene-based copy number (log2(CN ratio), median centered) of samples from SIBS SNP array hepatocellular carcinoma (HCC) dataset.

Supplementary Figure 40: CNA profiles (IGV heatmap) and correlation heatmap of gene-based copy number (log2(CN ratio), median centered) of samples from SIBS gene expression array (normalized by median expression of normal liver tissue samples) hepatocellular carcinoma (HCC) dataset.

Supplementary Figure 41: CNA profiles (IGV heatmap) and correlation heatmap of gene-based copy number (log2(CN ratio), median centered) of samples from SIBS gene expression array (normalized by median expression of tumor samples of the same dataset) hepatocellular carcinoma (HCC) dataset.

Supplementary Figure 42: CNA profiles (IGV heatmap) and correlation heatmap of gene-based copy number (log2(CN ratio), median centered) of samples from HCI SNP array breast cancer dataset.

Supplementary Figure 43: CNA profiles (IGV heatmap) and correlation heatmap of gene-based copy number (log2(CN ratio), median centered) of samples from HCI WES breast cancer dataset.

Supplementary Figure 44: CNA profiles (IGV heatmap) and correlation heatmap of gene-based copy number (log2(CN ratio), median centered) of samples from SNU-JAX WES gastric cancer dataset.

Supplementary Figure 45: CNA profiles (IGV heatmap) and correlation heatmap of gene-based copy number (log2(CN ratio), median centered) of samples from SNU-JAX RNA-Seq (normalized by median expression of normal gastric tissue samples from the same patients) gastric cancer dataset.

Supplementary Figure 46: CNA profiles (IGV heatmap) and correlation heatmap of gene-based copy number (log2(CN ratio), median centered) of samples from SNU-JAX RNA-Seq (normalized by median expression of tumor samples of the same dataset) gastric cancer dataset.

Supplementary Figure 47: CNA profiles (IGV heatmap) and correlation heatmap of gene-based copy number (log2(CN ratio), median centered) of samples from MDACC WES lung adenocarcinoma (LUAD) dataset.

Supplementary Figure 48: CNA profiles (IGV heatmap) and correlation heatmap of gene-based copy number (log2(CN ratio), median centered) of samples from MDACC WES lung squamous cell carcinoma (LUSC) dataset.

Supplementary Figure 49: CNA profiles (IGV heatmap) and correlation heatmap of gene-based copy number (log2(CN ratio), median centered) of samples from MDACC WES other lung cancer subtypes dataset.

Supplementary Figure 49: CNA profiles (IGV heatmap) and correlation heatmap of gene-based copy number (log2(CN ratio), median centered) of samples from MDACC WES other lung cancer subtypes dataset.

Supplementary Figure 50: CNA profiles (IGV heatmap) and correlation heatmap of gene-based copy number (log2(CN ratio), median centered) of samples from PDMR WES bladder cancer dataset.

Supplementary Figure 51: CNA profiles (IGV heatmap) and correlation heatmap of gene-based copy number (log2(CN ratio), median centered) of samples from PDMR WES colorectal cancer dataset.

Supplementary Figure 52: CNA profiles (IGV heatmap) and correlation heatmap of gene-based copy number (log2(CN ratio), median centered) of samples from PDMR WES head and neck cancer dataset.

Supplementary Figure 53: CNA profiles (IGV heatmap) and correlation heatmap of gene-based copy number (log2(CN ratio), median centered) of samples from PDMR WES lung cancer dataset.

Supplementary Figure 54: CNA profiles (IGV heatmap) and correlation heatmap of gene-based copy number (log2(CN ratio), median centered) of samples from PDMR WES pancreatic cancer dataset.

Supplementary Figure 55: CNA profiles (IGV heatmap) and correlation heatmap of gene-based copy number (log2(CN ratio), median centered) of samples from PDMR WES renal cancer dataset.

Supplementary Figure 56: CNA profiles (IGV heatmap) and correlation heatmap of gene-based copy number (log2(CN ratio), median centered) of samples from PDMR WES sarcoma dataset.

Supplementary Figure 57: CNA profiles (IGV heatmap) and correlation heatmap of gene-based copy number (log2(CN ratio), median centered) of samples from PDMR WES skin cancer dataset.

Supplementary Figure 58: CNA profiles (IGV heatmap) and correlation heatmap of gene-based copy number (log2(CN ratio), median centered) of samples from PDMR WES other cancers dataset.

Supplementary Figure 59: CNA profiles (IGV heatmap) and correlation heatmap of gene-based copy number (log2(CN ratio), median centered) of samples from WISTAR WES skin melanoma dataset.

Supplementary Figure 60: CNA profiles (IGV heatmap) and correlation heatmap of gene-based copy number (log2(CN ratio), median centered) of samples from WUSTL WES breast cancer dataset.

Supplementary Figure 61: CNA profiles (IGV heatmap) and correlation heatmap of gene-based copy number (log2(CN ratio), median centered) of samples from WUSTL WES pancreatic cancer dataset.

Supplementary Figure 62: CNA profiles (IGV heatmap) and correlation heatmap of gene-based copy number (log2(CN ratio), median centered) of samples from EuroPDX WGS colorectal cancer (liver metastases) dataset.

Supplementary Figure 63: CNA profiles (IGV heatmap) and correlation heatmap of gene-based copy number (log2(CN ratio), median centered) of samples from EuroPDX WGS breast cancer dataset.

Supplementary Figure 64: PT samples have a lower range of CNA values than PDX samples. Comparison of range of CNA in pairs of samples (PT or PDX) from the same model (left panel). Pearson correlation of the samples versus the minimum range of the two samples (right panel). Samples with lower range tend to have lower correlations with other samples. For a given sample, range is defined as $\log_2$(CN ratio) of the 95$^{th}$ percentile minus $\log_2$(CN ratio) of the 5$^{th}$ percentile value of median-centered copy number values across 100-kb windows binned from copy number segments of each sample. (**a**) All data; (**b**) After removing comparisons of low correlation (< 0.6) due to non-aberrant samples (range < 0.3). (Sample 1: PT or lower passage PDX, Sample 2: later passage PDX or same passage PDX of different lineage)

Supplementary Figure 65: (**a**) Range of CNA profiles between PT-PDX or PDX-PDX sample pairs from the same model. (**b**) Pearson correlation of the samples versus the ratio of range between two samples (PT/PDX or PDX-1/PDX-2). Samples pairs with ratio of range much greater or less than 1 i.e. one sample is much less aberrant than the other, tend to have lower correlations. For a given sample, range is defined as $\log_2$(CN ratio) of the 95$^{th}$ percentile minus $\log_2$(CN ratio) of the 5$^{th}$ percentile value of median-centered copy number values across 100-kb windows binned from copy number segments of each sample. P-values were computed by Wilcoxon rank sum test (ns: non-significant, $p > 0.05$). (PDX-1: lower passage PDX, PDX-2: later passage PDX or same passage PDX of different lineage)

Supplementary Figure 66: (a) Comparison of Pearson correlation coefficients of gene-based copy number using DNA-based (WES or SNP array) versus RNA-based (RNA-Seq or gene expression array) methods. (b) Distribution of Pearson correlation coefficients of gene-based copy number, using DNA-based (WES or SNP array) and RNA-based (RNA-Seq or gene expression array) methods, for pairs of samples with high correlation by the DNA-based method (>0.8 for SNU-JAX Gastric cancer, >0.9 for SIBS HCC). P-values were computed by Wilcoxon rank sum test.

Supplementary Figure 67: Distribution of Pearson correlation coefficients of gene-based copy number, estimated by (a) SNP array, (b) WES, (c) WGS, between different combinations of patient tumor and PDX passages of the same model. Comparisons relative to passages P1 or later (refer to Fig. 3d – f for comparisons with PT and P0).

Supplementary Figure 68: Distribution of Pearson correlation coefficients of gene-based copy number between early and very-late passages of the same model for the BCM SNP array breast cancer dataset. Correlation for “other passages” are based on models from all other SNP array datasets.

Supplementary Figure 69: Correlation and regression of gene-based copy number between early and very-late passages of the same model for the BCM SNP array breast cancer dataset. Genes with copy number changes between the passages are identified by |residual| > 0.5 (purple dots). Some genes show signs of complete deletion ($log_2$(CN ratio) < -2) but then reappear in later passages, suggesting bottlenecks from minor populations.

Supplementary Figure 70: Scatter plot of Pearson correlation between samples of PDX-early and PDX-late versus the corresponding passage difference for same lineage samples.

Supplementary Figure 71: $log_2$(CN ratio) values between each pair of samples of recurrent genes (see Supplementary Table 4). PDX-1: earlier passage, PDX-2: same passage but different lineage or later passage.

Supplementary Figure 72. GISTIC analysis of recurrent CNAs in TCGA primary tumors and EurOPDX collections of PTs and derived PDXs, at early and late passages, of (a) colorectal cancer and (b) breast cancer. For each GISTIC plot the top axis reports the G-score and the bottom axis the q-value. Red line plots: amplifications, blue line plots: deletions.

Supplementary Figure 73: Correlation heatmap of gene-based copy number (log2(CN ratio), median centered) of multi-region samples of the same tumor from TRACERx (a) lung adenocarcinoma (LUAD), (b) lung squamous cell carcinoma (LUSC) and (c) other lung cancer subtypes.

Supplementary Figure 74: Comparison of distribution of proportion of altered genes between multi-region tumor pairs from TRACERx, and PT-PDX and PDX-PDX pairs for various gene sets for LUAD and LUSC. Copy number altered genes were identified by |residual| > 0.5 from linear regression model for each pairwise comparison. P-values were computed by Wilcoxon rank sum test. Protein-coding: protein-coding genes annotated by Ensembl; Oncogenic pathways: genes in oncogenic signaling pathways identified by TCGA; Census Amp Del: genes with frequent amplifications or deletions annotated in the Cancer Gene Census

Supplementary Figure 75: Fraction of genes of different gene sets (see Fig. 4) with copy number changes (|residual| > 0.5) between early and late passages of each breast cancer model from the BCM breast cancer dataset.

Supplementary Figure 76: Window-based and segmented copy number estimated as (**a**) depth ratio by Sequenza for WES, and (**b**) $log_2$ (ratio) by ASCAT for WGS, for same lineage samples identified to be aberrant (range > 0.5 for both samples) and low correlation (Pearson correlation coefficient < 0.6) between the sample pairs

Supplementary Figure 77: Workflow for copy number estimation by Sequenza from whole-exome sequencing data with paired-normal for (A) patient tumor (without Xenome) and (B) PDX tumor (with Xenome for mouse reads removal).

### Supplementary Tables

Supplementary Table 1: Summary of datasets collected from various centers in the PDXNET consortium, EuroPDX consortium and published datasets.

Supplementary Table 2: Summary of datasets by tumor type.

Supplementary Table 3: Benchmarking dataset which comprises copy number alteration profiles estimated for matched samples assayed across multiple platforms

Supplementary Table 4: Recurrent frequency (based on models) of genes with >5% recurrence with large copy number deviation (|residual| > 1) from linear regression model for PT-PDX (279 models) and PDX-PDX (306 models) comparisons.

Supplementary Table 5: Association between copy number status of cell lines (n=668) and drug response (n=545) from CCLE database for recurrent genes in Supplementary Table 4. (Attached as separate Excel spreadsheet)

Supplementary Table 6: Segment-level GISTIC scores for CRC and BRCA WGS cohorts, subdivided into PT, PDX-early and PDX-late. (Attached as separate Excel spreadsheet)

Supplementary Table 7: Gene-level GISTIC scores for CRC and BRCA WGS cohorts, subdivided into PT, PDX early and PDX late.

Supplementary Table 8: Significance thresholds for G-score in the various CRC and BRCA WGS cohorts. (Attached as separate Excel spreadsheet)

Supplementary Table 9: Results of GSEA analysis. Gene sets found enriched in genes with high or low G-Score in at least one of the PT / PDX-early / PDX-late cohorts of the CRC and BRCA WGS datasets. (Attached as separate Excel spreadsheet)

### Supplementary Data

Supplementary Dataset 1: CNA segments with median-centered log2(CN ratio) of all datasets (Attached as separate Excel spreadsheet)

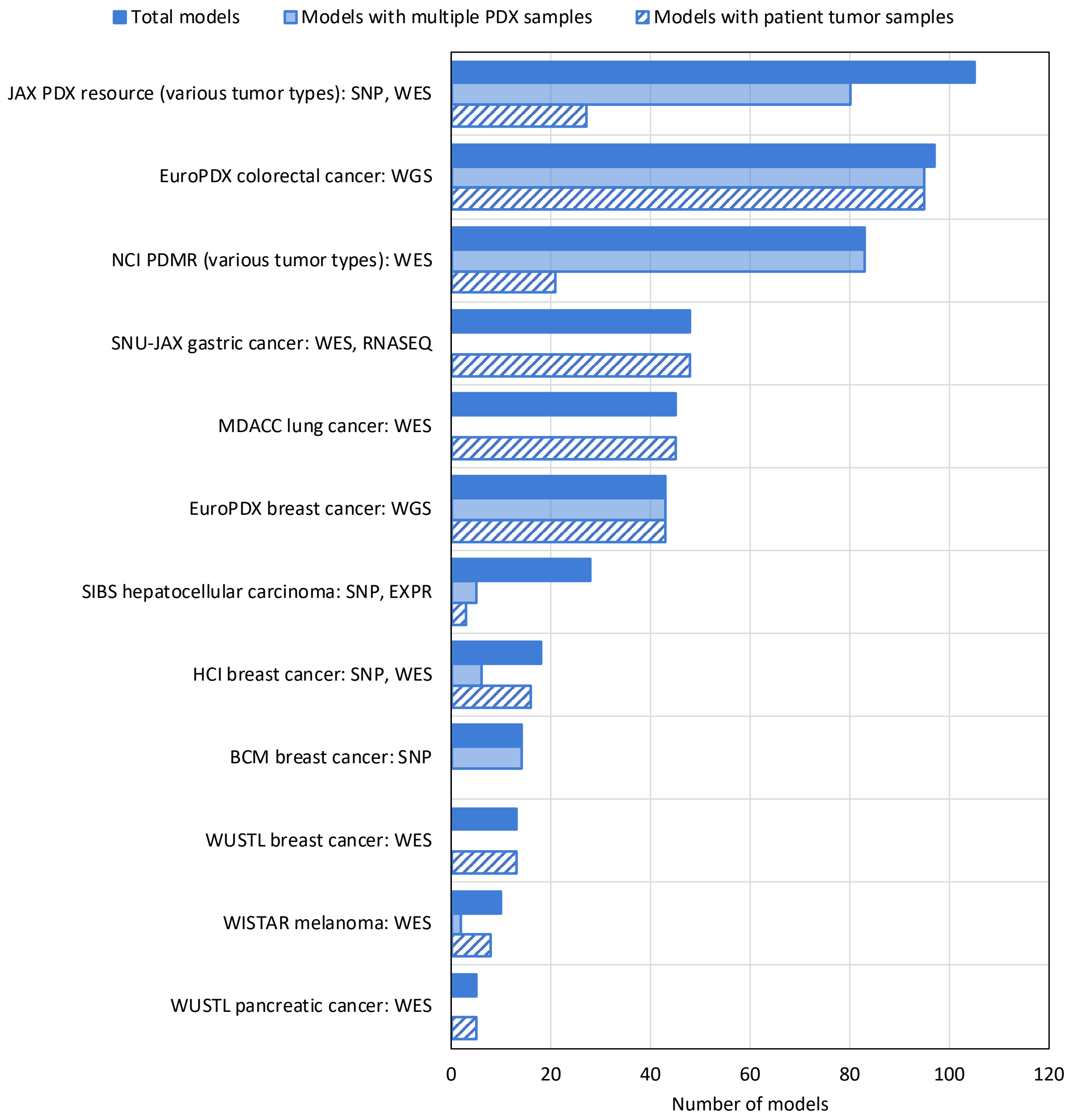

Supplementary Figure 1: Summary of PDX models collected from various centers in the PDXNET and EuroPDX consortium, and publicly available data.

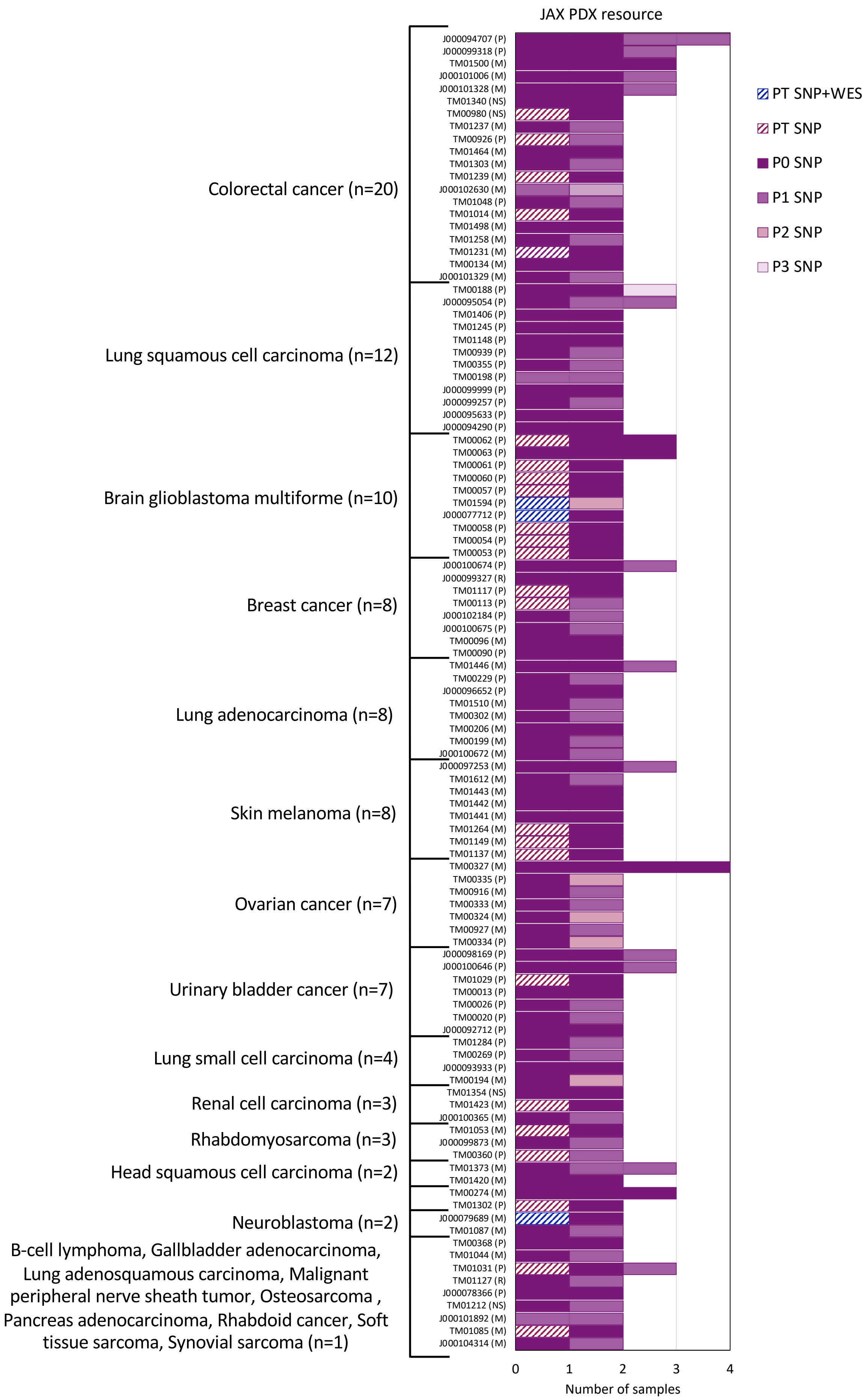

Supplementary Figure 2: Number of patient tumor and PDX passages per model in the JAX PDX resource consisting of various tumor types assayed by Affymetrix SNP 6.0 array and whole exome sequencing (P: Primary malignancy, M: Metastatic, R: Recurrent/Relapse, NS: Not specified).

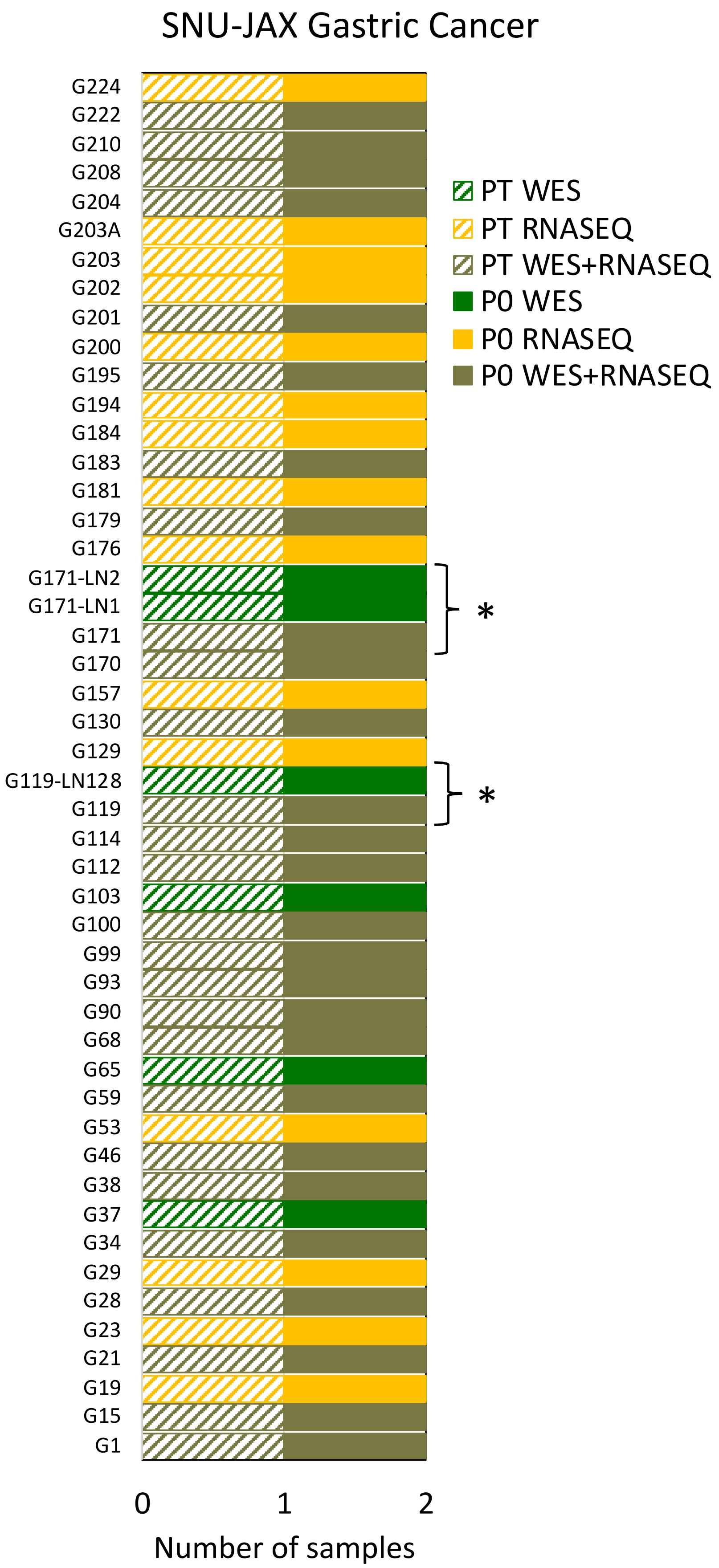

Supplementary Figure 3: Number of patient tumor and PDX passages per model in SNU-JAX gastric cancer dataset assayed by whole-exome sequencing (WES) and RNA sequencing (RNASEQ). Models labeled with "LN" are lymph node metastatic tumors for the same patient. (*: Multiple patient tumors available for the same patient, different relapse time points or different metastatic sites)

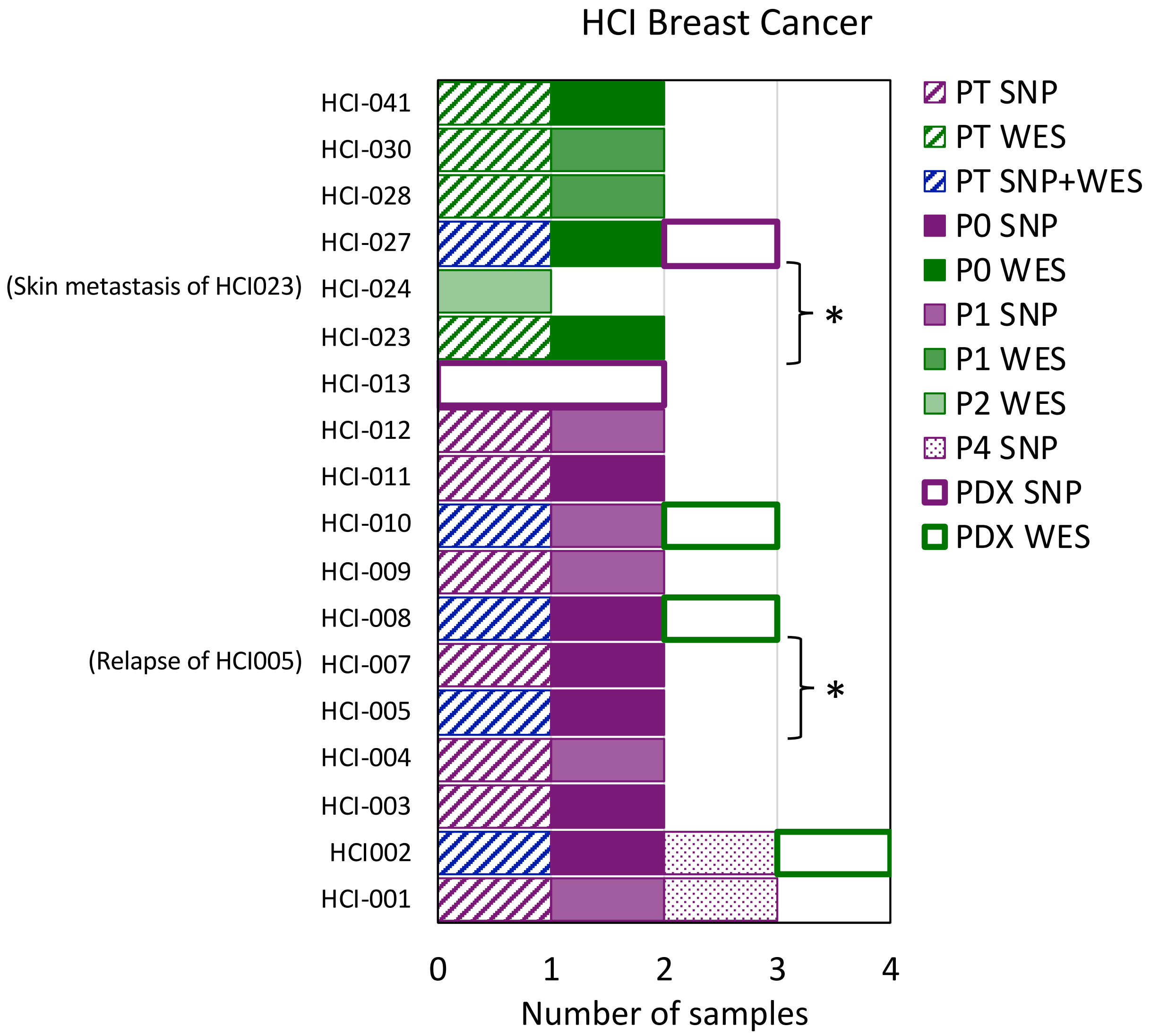

Supplementary Figure 4: Number of patient tumor and PDX passages per model in the HCI breast cancer dataset assayed by whole-exome sequencing (WES), Affymetrix SNP 6.0 array and Illumina Infinium Omni2.5Exome8 (v1.3) SNP array. HCI-007 is relapse tumor from the patient of HCI-005, HCI-024 is a skin metastasis tumor from the patient of HCI-023. Samples labeled with “PDX” indicates passage number is unknown. (*: Multiple tumors available for the same patient, different relapse time points or different metastatic sites)

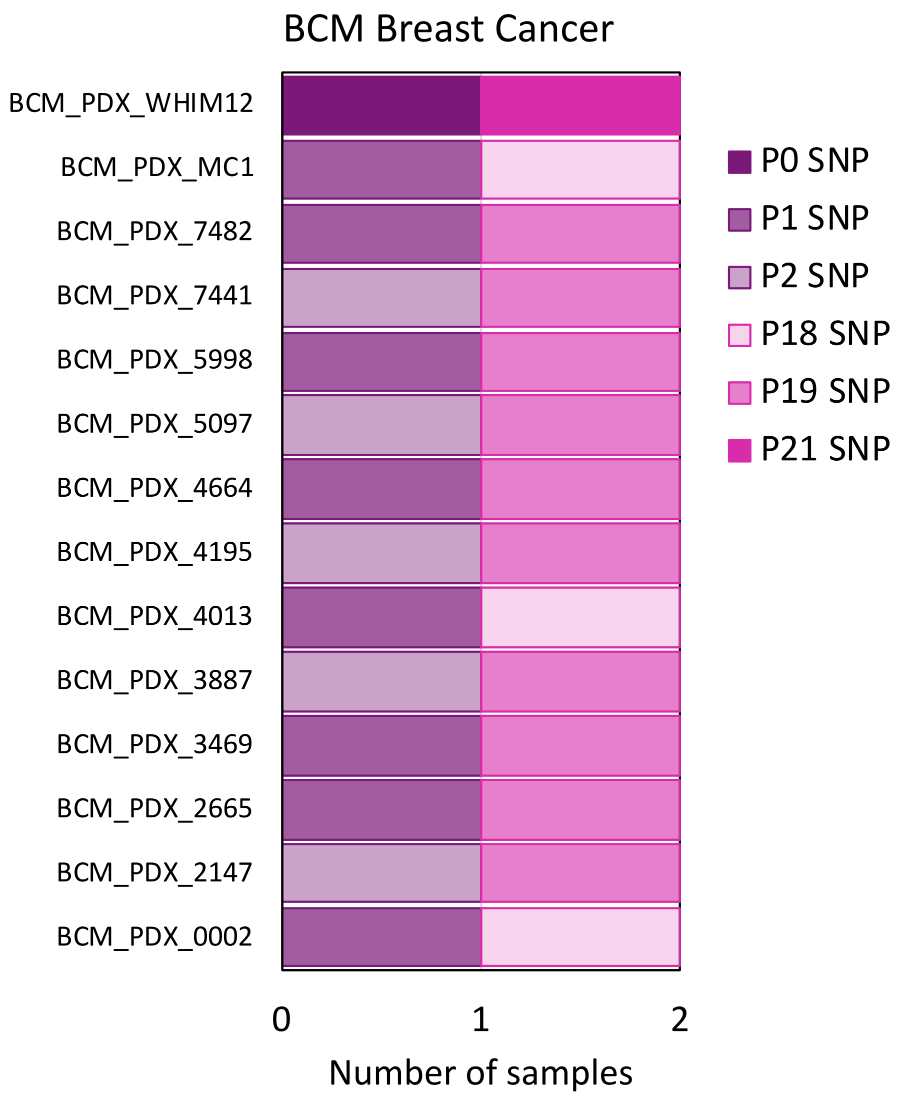

Supplementary Figure 5: Number of PDX passages per model in the BCM breast cancer dataset assayed by Illumina Infinium Omni2.5Exome8 (v1.4) SNP array.

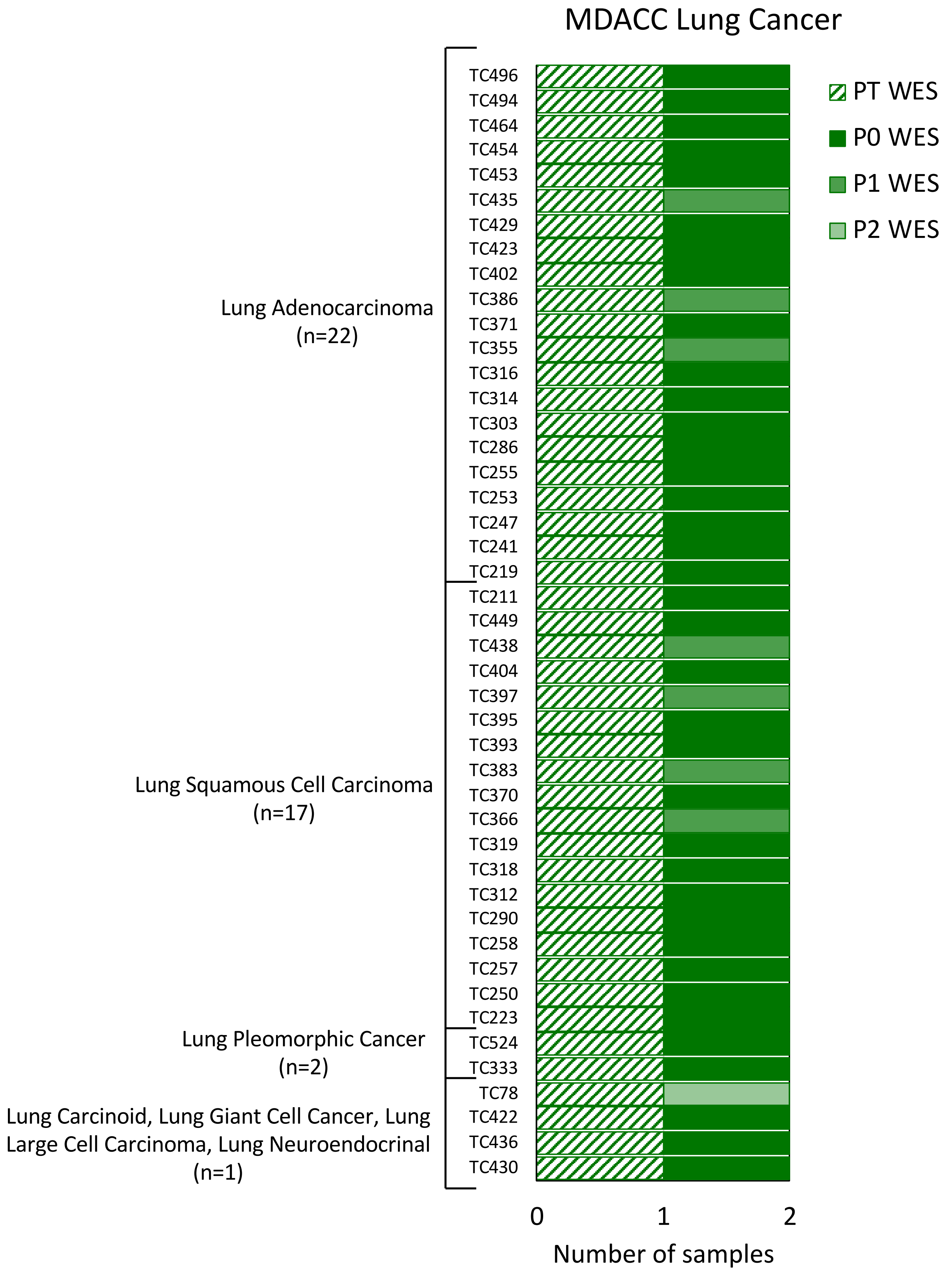

Supplementary Figure 6: Number of patient tumor and PDX passages per model in the MDACC lung cancer dataset assayed by whole-exome sequencing (WES).

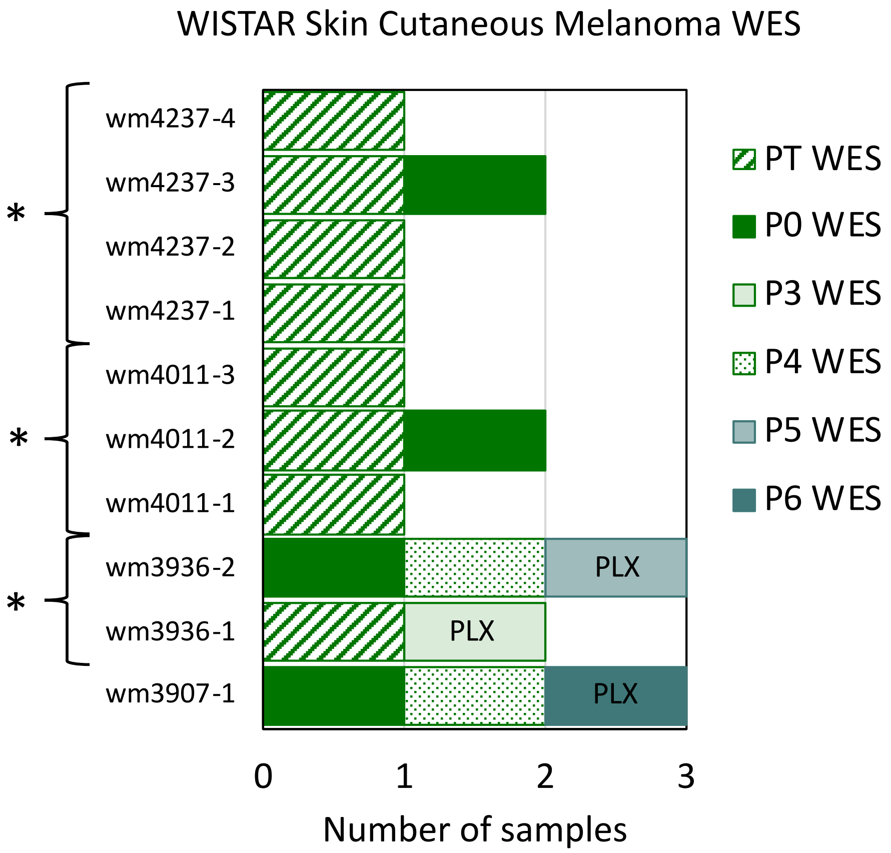

Supplementary Figure 7: Number of patient tumor and PDX passages per model in Wistar skin cutaneous melanoma dataset assayed by whole-exome sequencing (WES). PDX samples labeled with ”PLX” are BRAF inhibitor (PLX) treated. (*: Multiple tumors available for the same patient, different relapse time points or different metastatic sites)

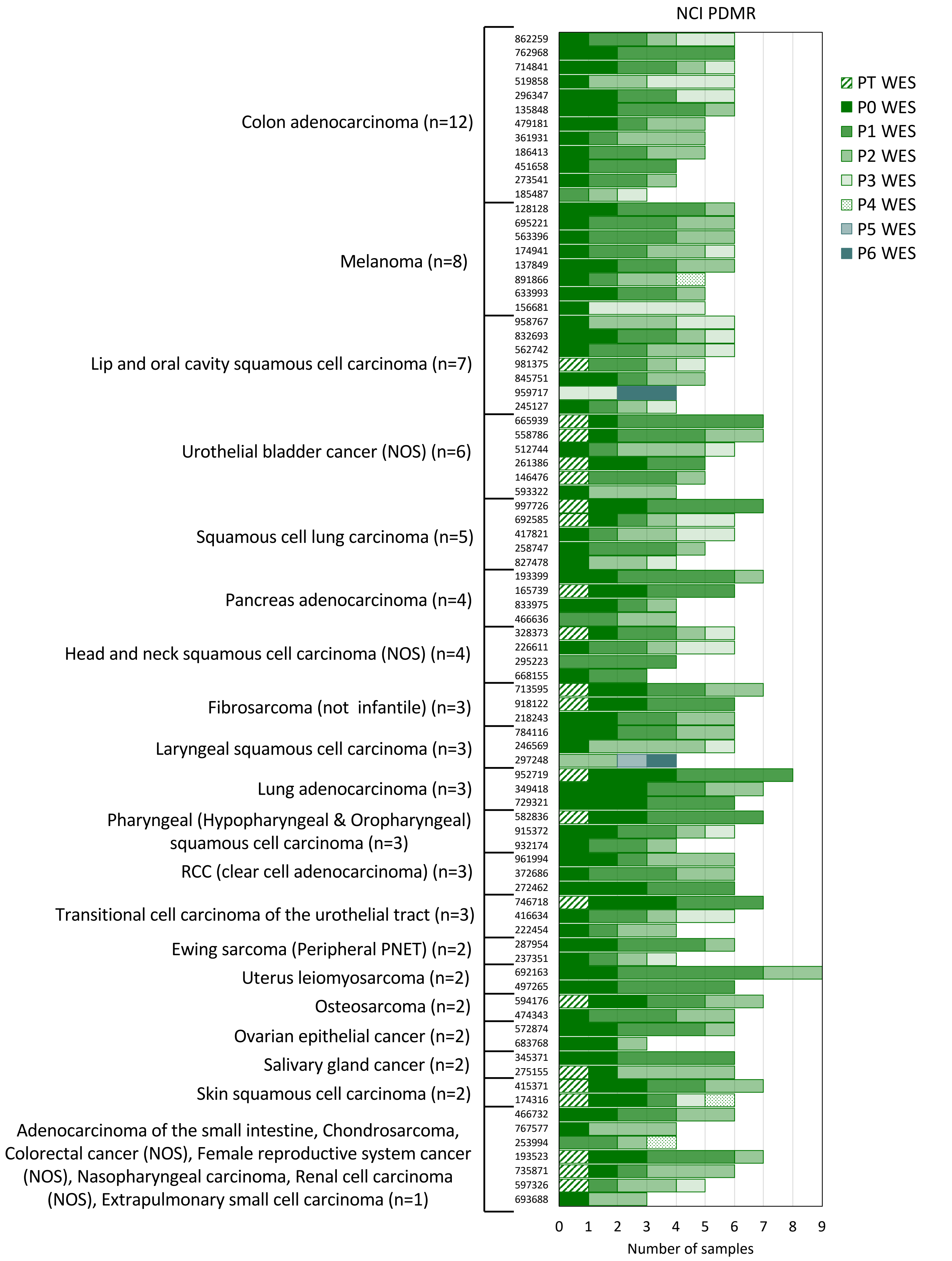

Supplementary Figure 8: Number of patient tumor and PDX passages per model in the NCI PDMR resource consisting of various tumor types assayed by whole exome sequencing.

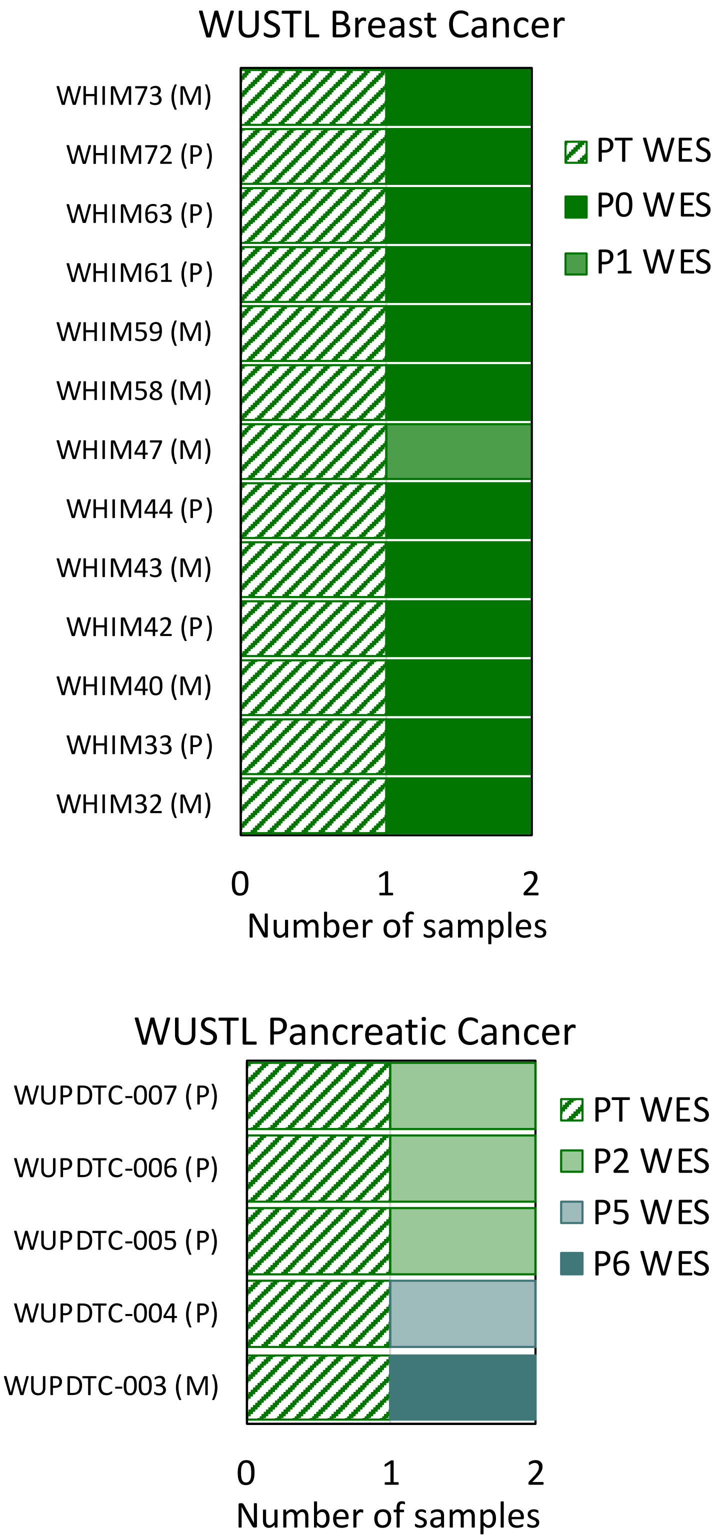

Supplementary Figure 9: Number of patient tumor and PDX passages per model in the WUSTL breast cancer and pancreatic cancer datasets assayed by whole-exome sequencing (WES).

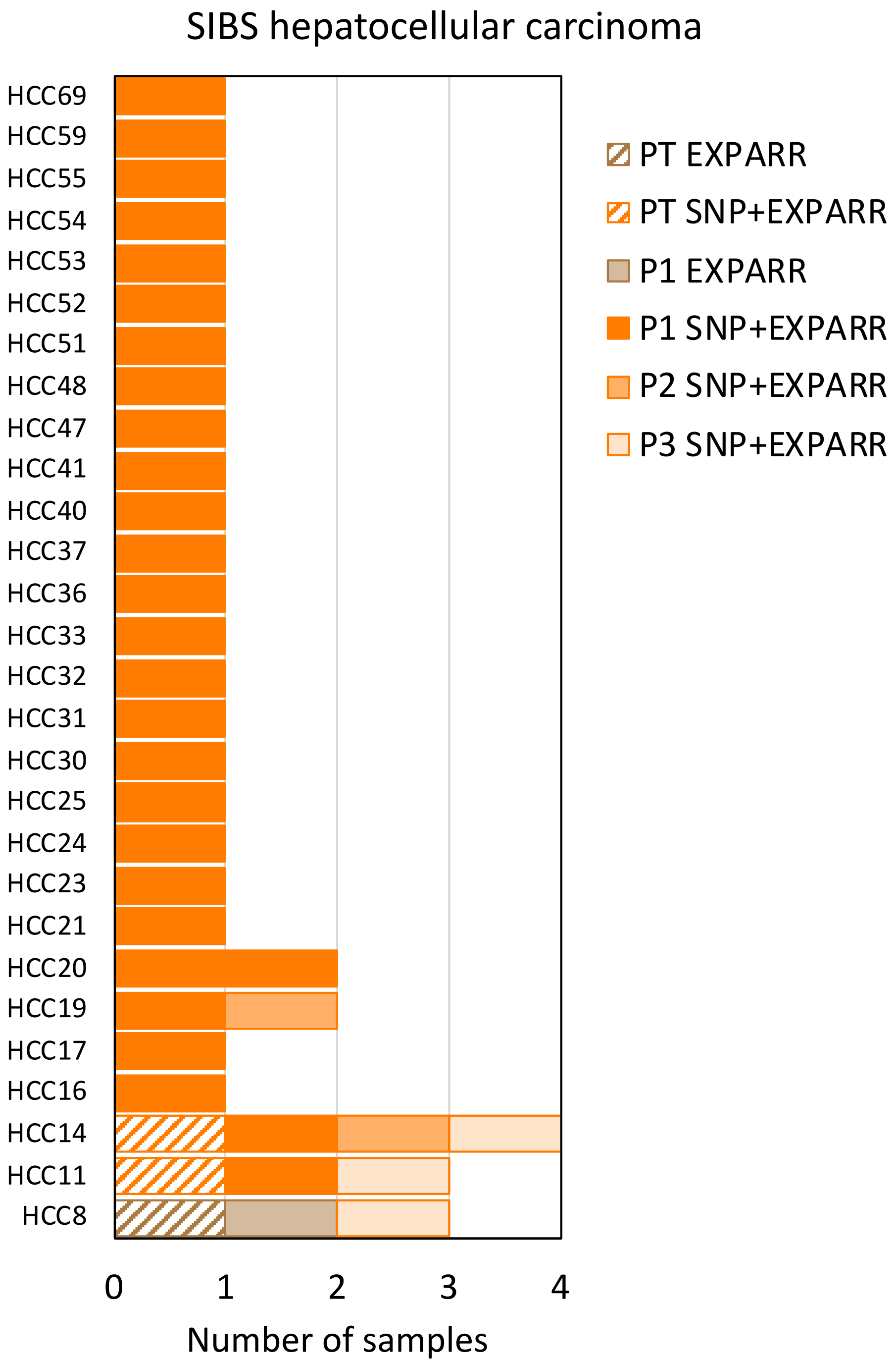

Supplementary Figure 10: Number of patient tumor and PDX passages per model in the SIBS hepatocellular carcinoma dataset assayed by Affymetrix SNP 6.0 array and Affymetrix gene expression array (EXPARR).

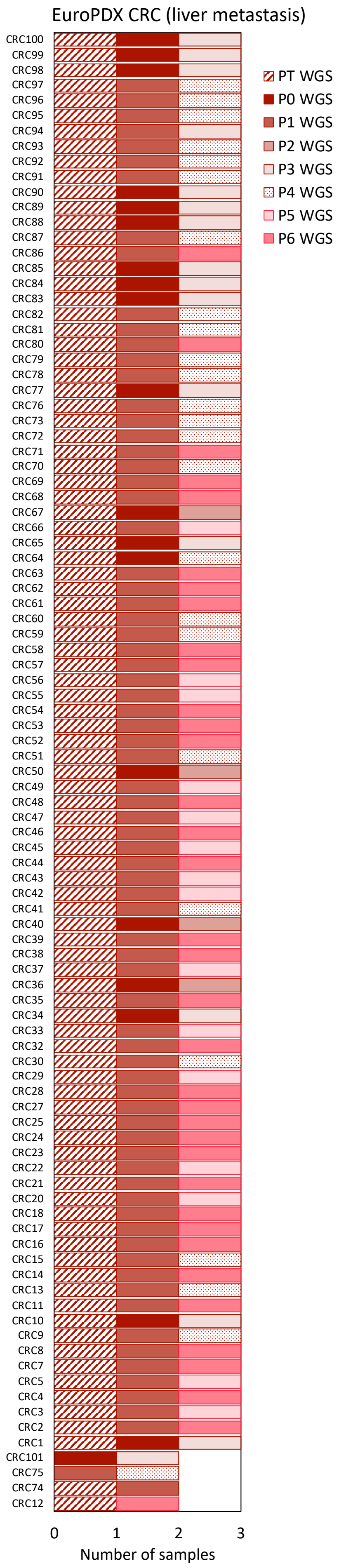

Supplementary Figure 11: Number of patient tumor and PDX passages per model in the EuroPDX colorectal cancer (CRC) liver metastasis dataset assayed by whole-genome sequencing.

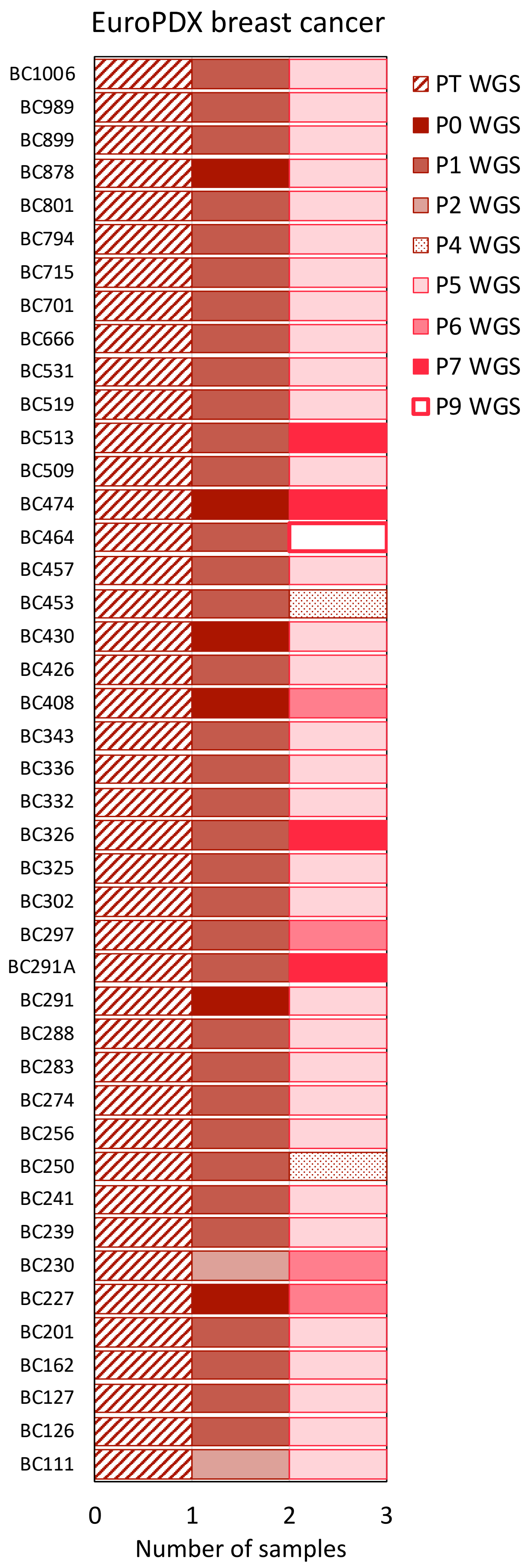

Supplementary Figure 12: Number of patient tumor and PDX passages per model in the EuroPDX breast cancer (BRCA) dataset assayed by whole-genome sequencing.

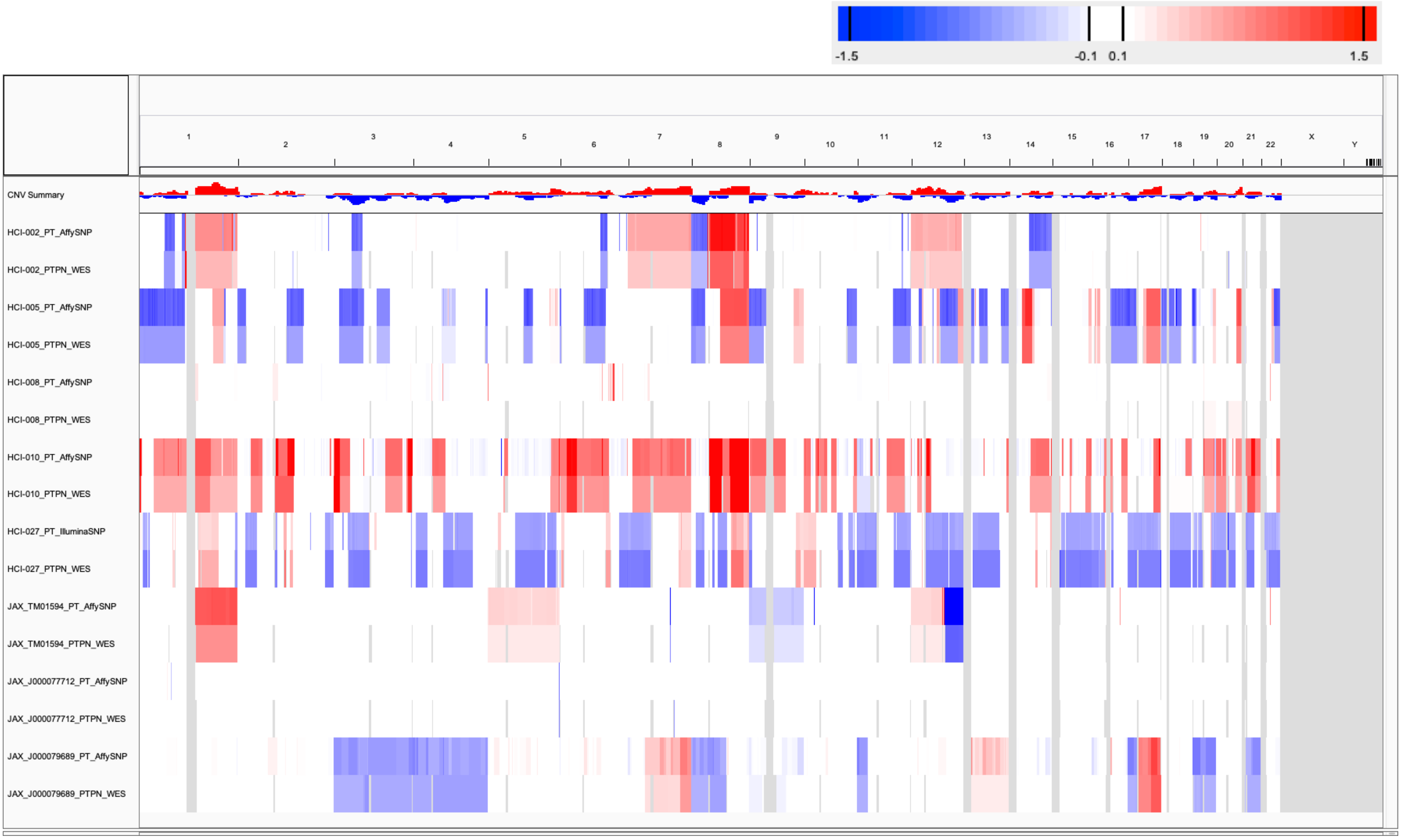

Supplementary Figure 13: CNA profiles for matched patient tumor samples estimated from SNP array and WES for ”SNP vs WES” benchmarking (see Supplementary Table 3).

a

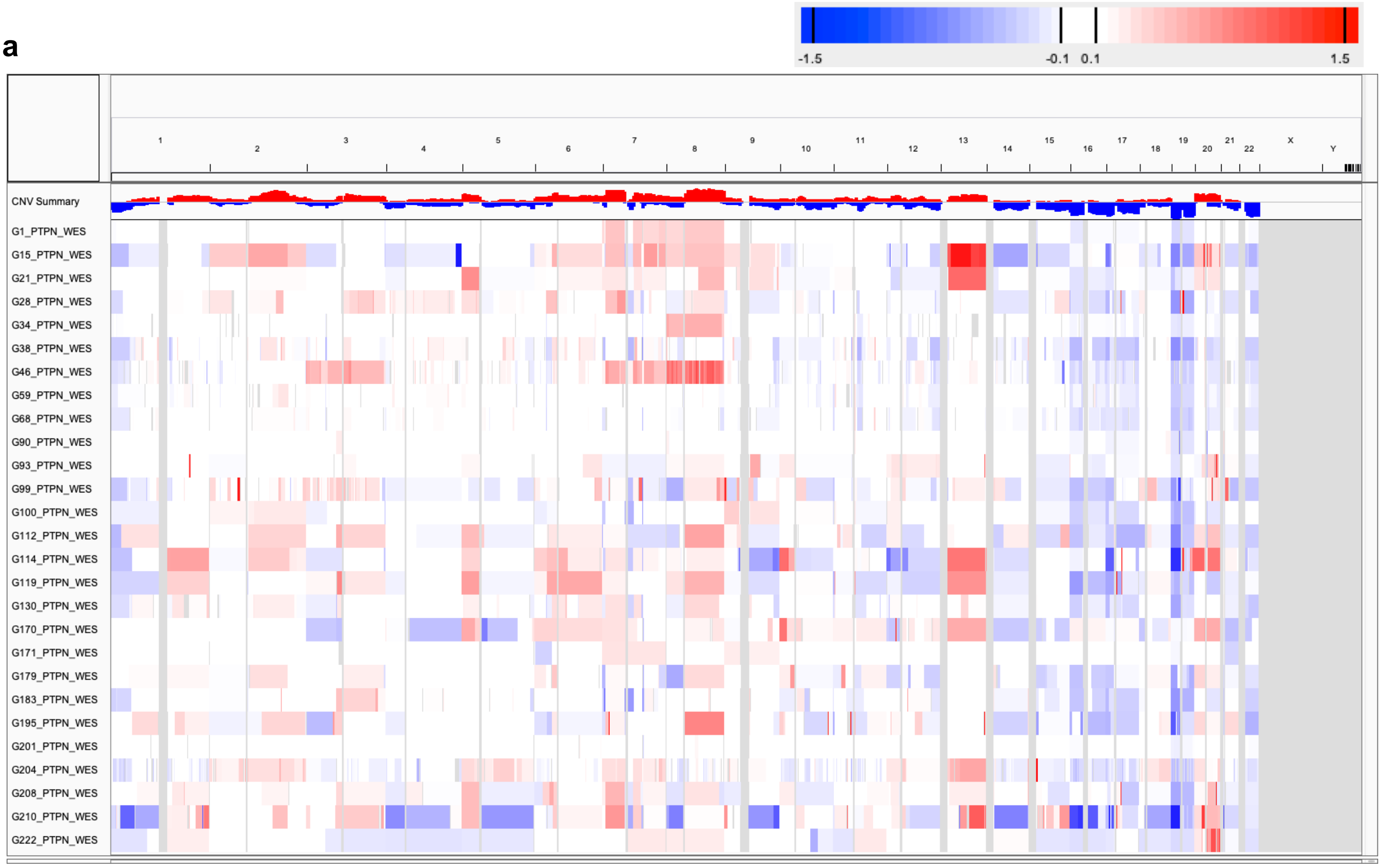

b

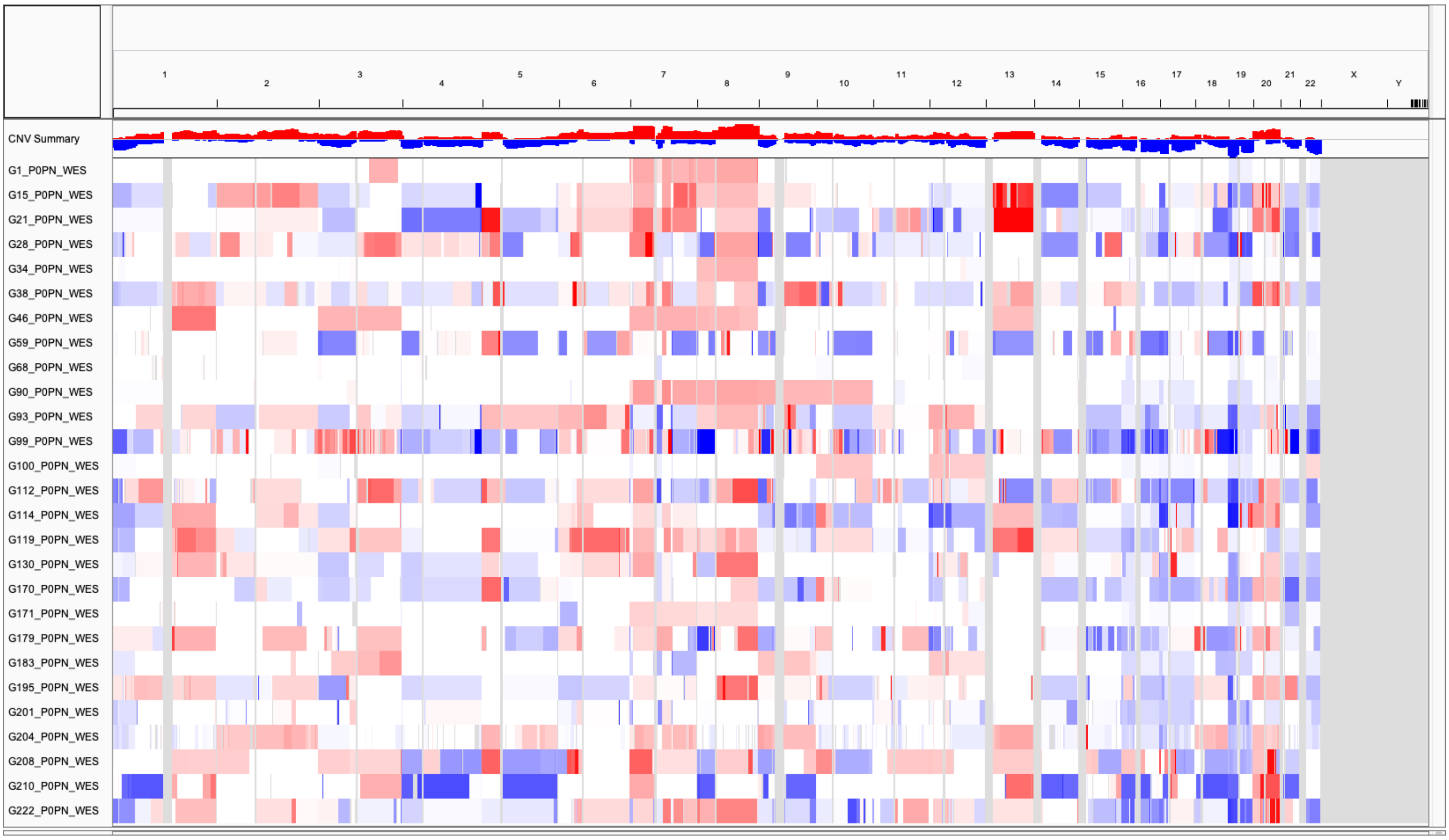

Supplementary Figure 14: CNA profiles for (**a**) patient tumor and (**b**) PDX samples estimated from WES used for ”WES vs RNASEQ (NORM/TUM)” benchmarking (see Supplementary Table 3).

a

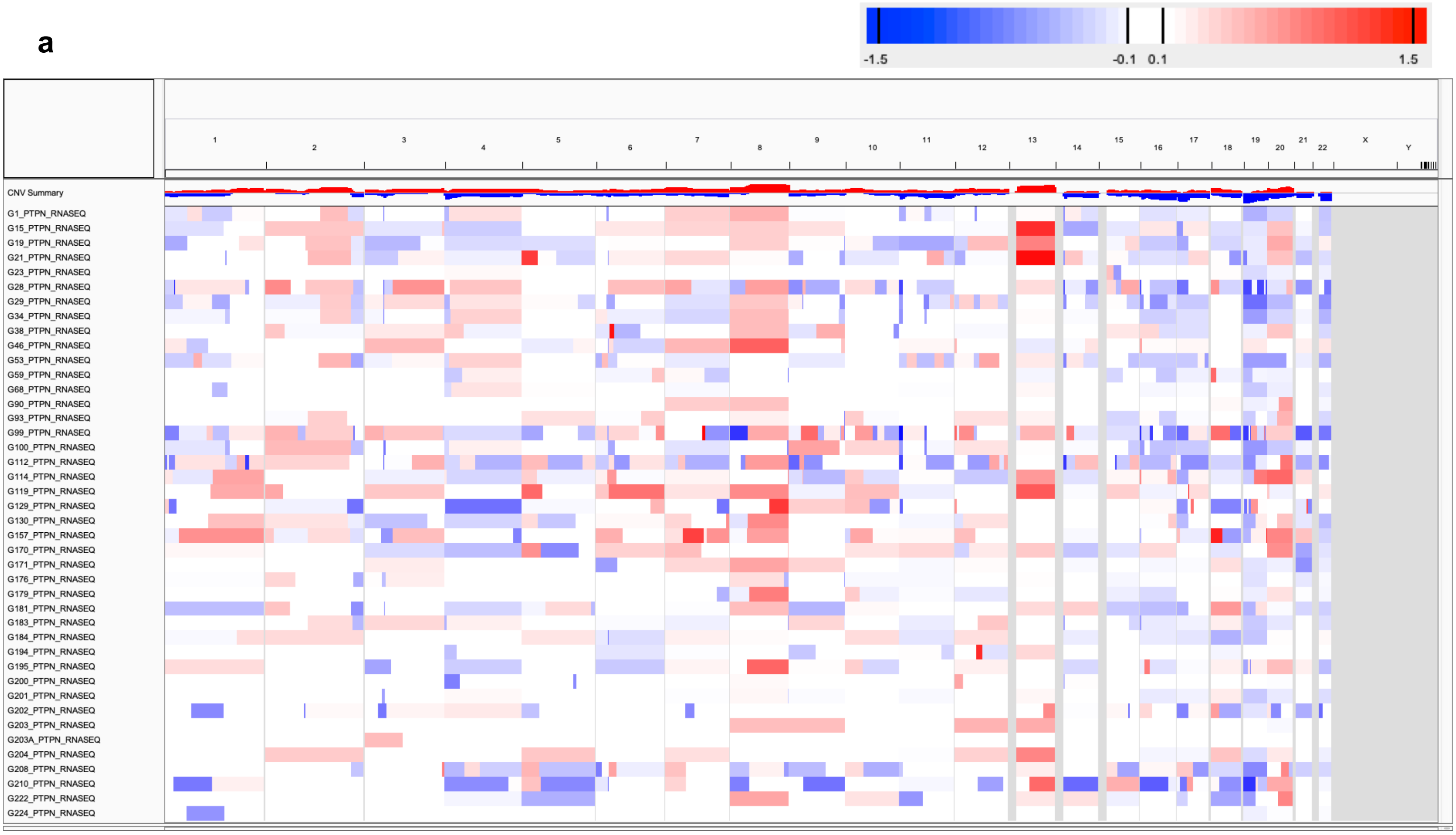

b

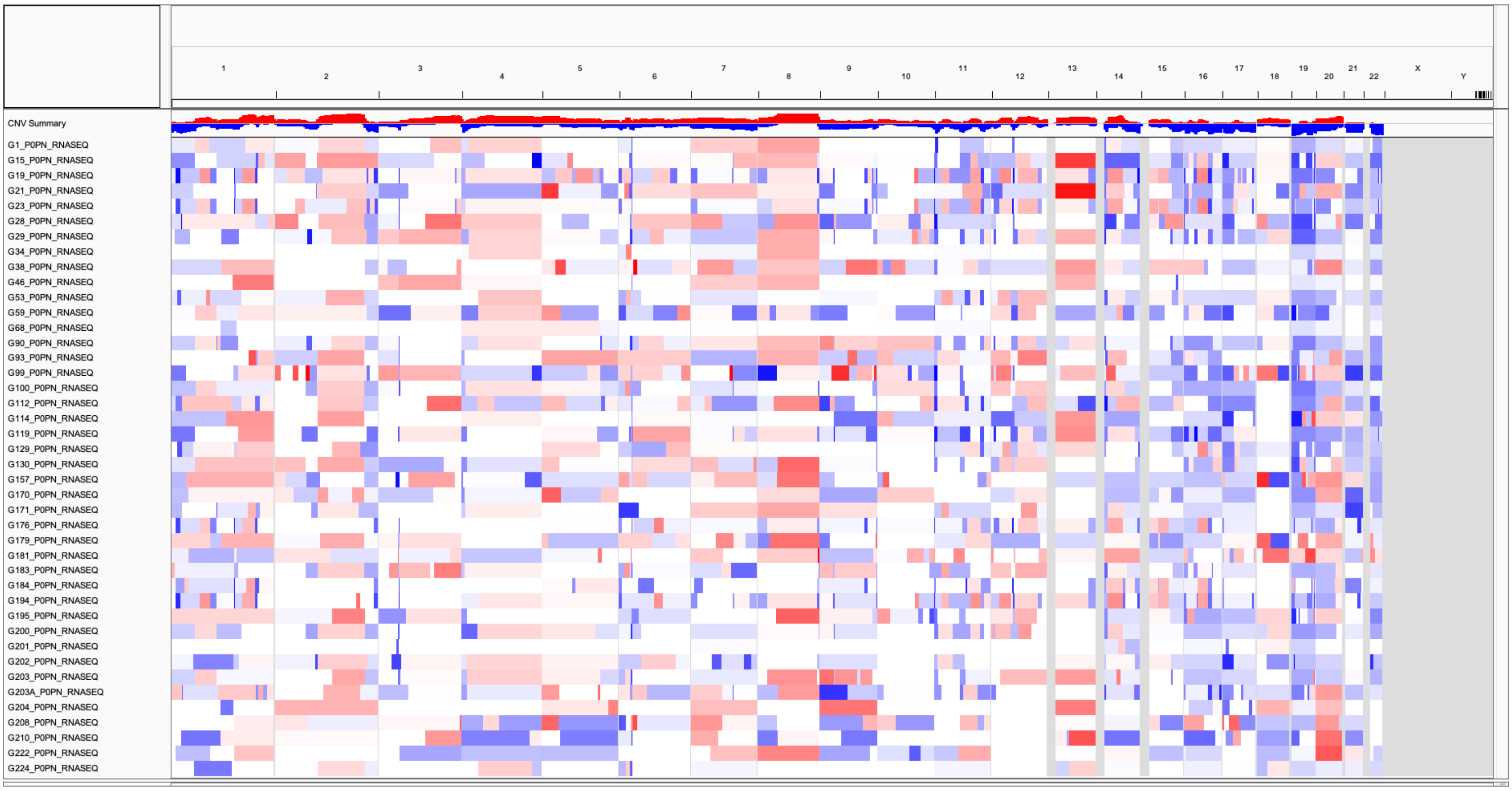

Supplementary Figure 15: (Continue next page)

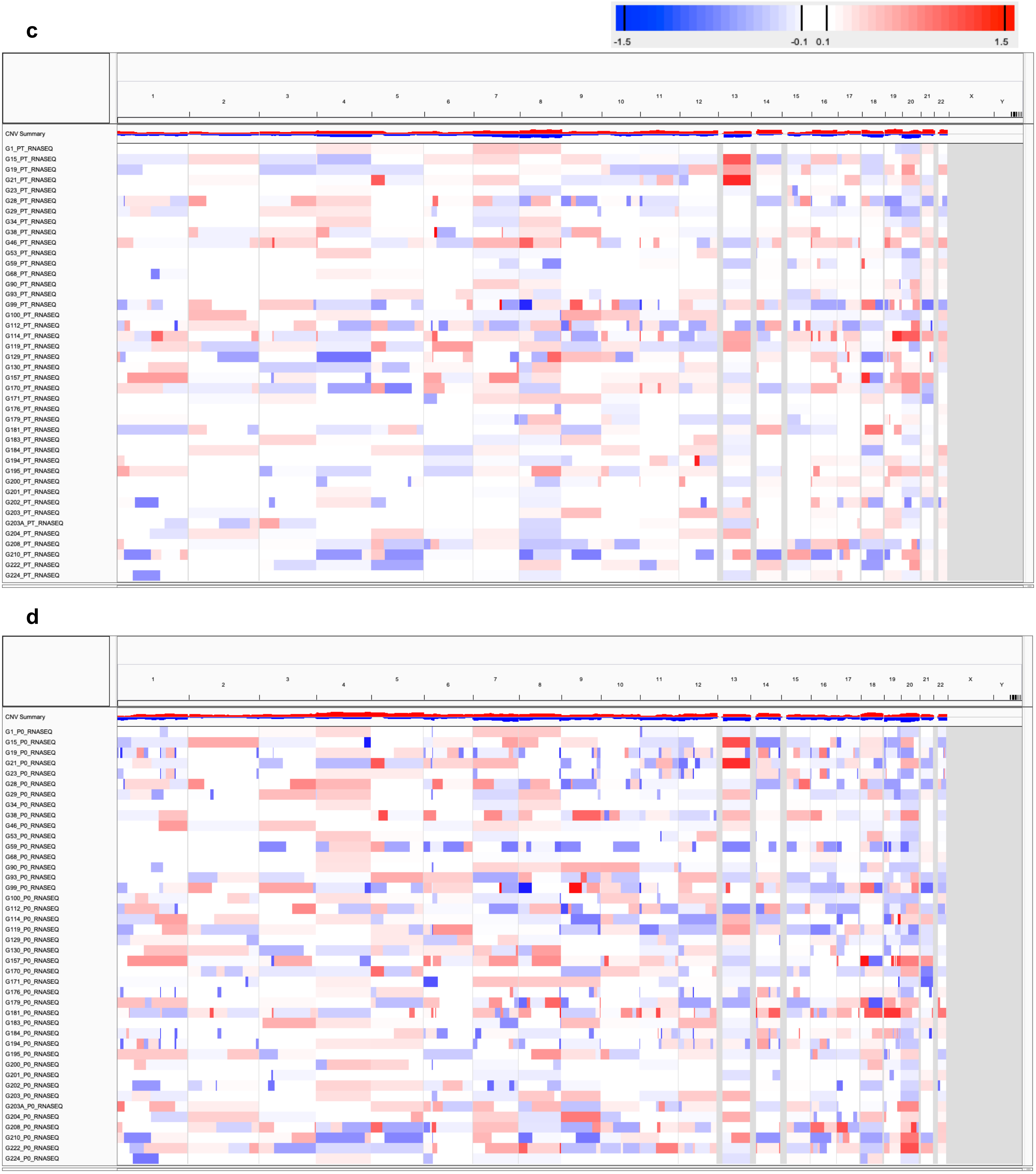

Supplementary Figure 15: CNA profiles for (**a**) patient tumor and (**b**) PDX samples estimated from RNA-Seq, normalized by median expression of normal samples of the same tumor type, used for ”WES vs RNASEQ (NORM)” and “RNASEQ NORM vs TUM” benchmarking. CNA profiles for (**c**) patient tumor and (**d**) PDX samples estimated from RNA-Seq, normalized by median expression of same set of patient tumors, used for ”WES vs RNASEQ (TUM)” and “RNASEQ NORM vs TUM” benchmarking (see Supplementary Table 3).

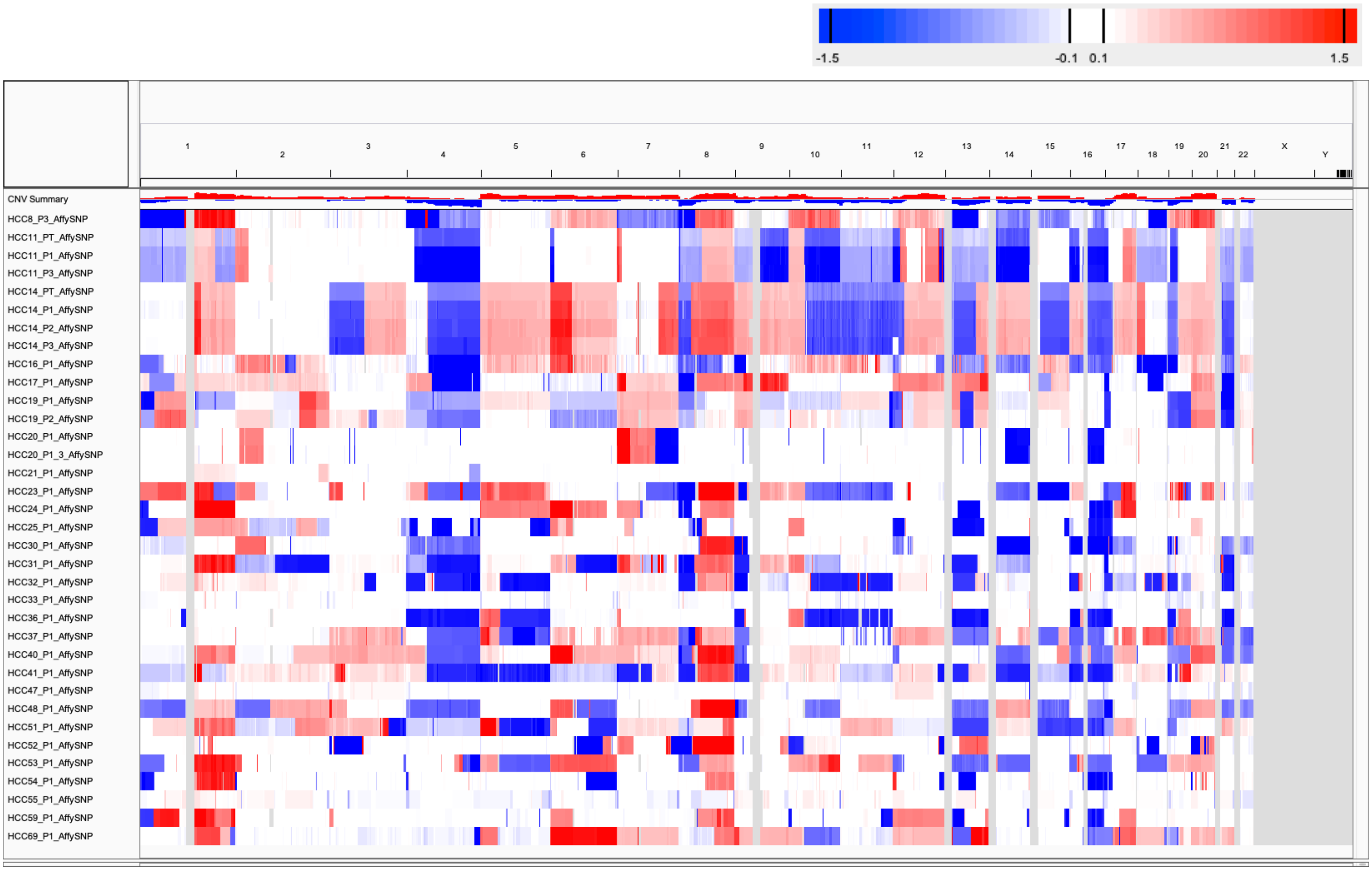

Supplementary Figure 16: CNA profiles for patient tumor and PDX samples estimated from SNP array used for ”SNP vs EXPARR (NORM/TUM)” benchmarking (see Supplementary Table 3).

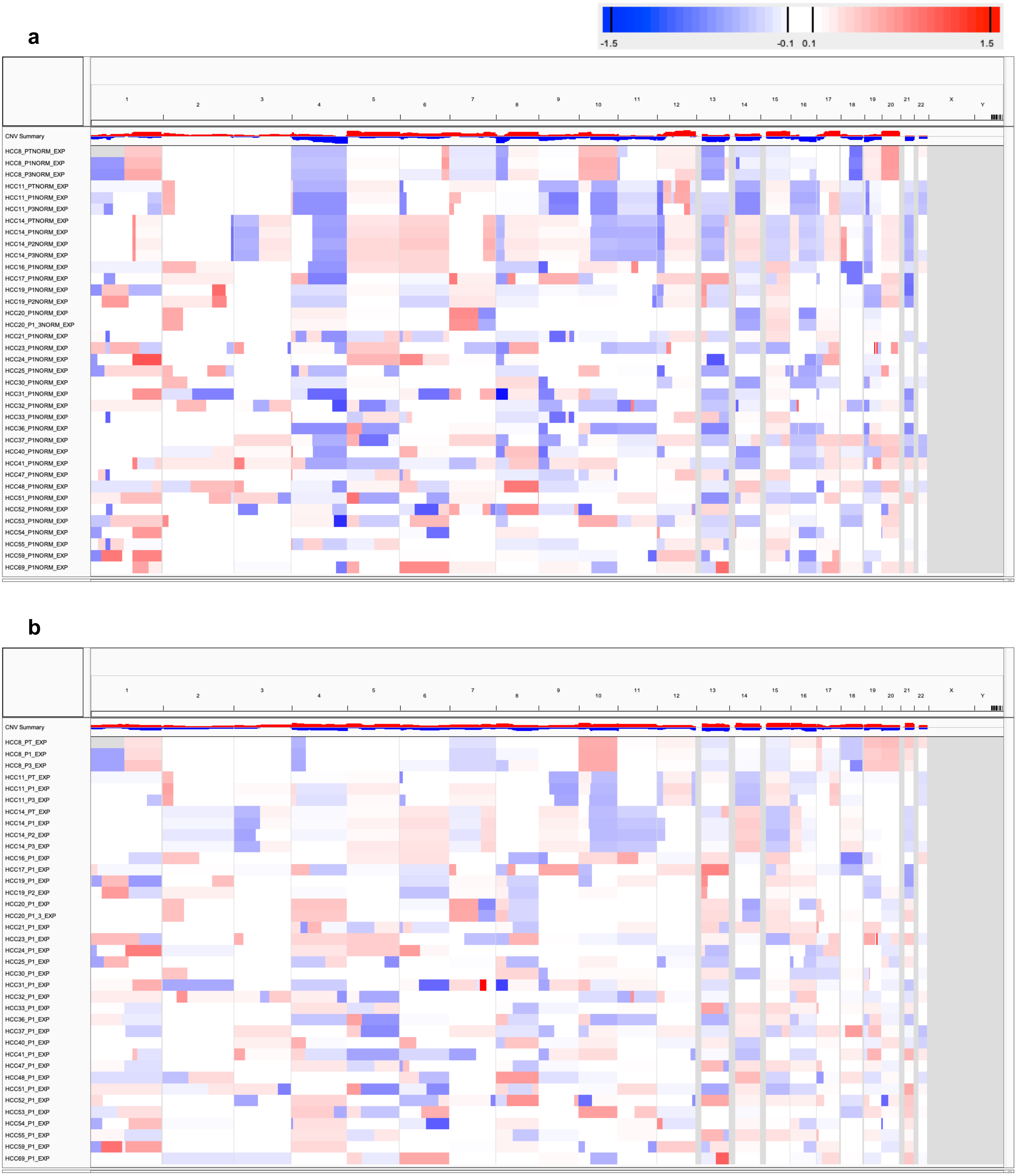

Supplementary Figure 17: CNA profiles for patient tumor and PDX samples estimated from gene expression array, normalized by (**a**) median expression of normal samples of the same tumor type and (**b**) median expression of same set of patient tumors, used for ”SNP vs EXPARR (NORM/TUM)” and “EXPARR NORM vs TUM” benchmarking (see Supplementary Table 3).

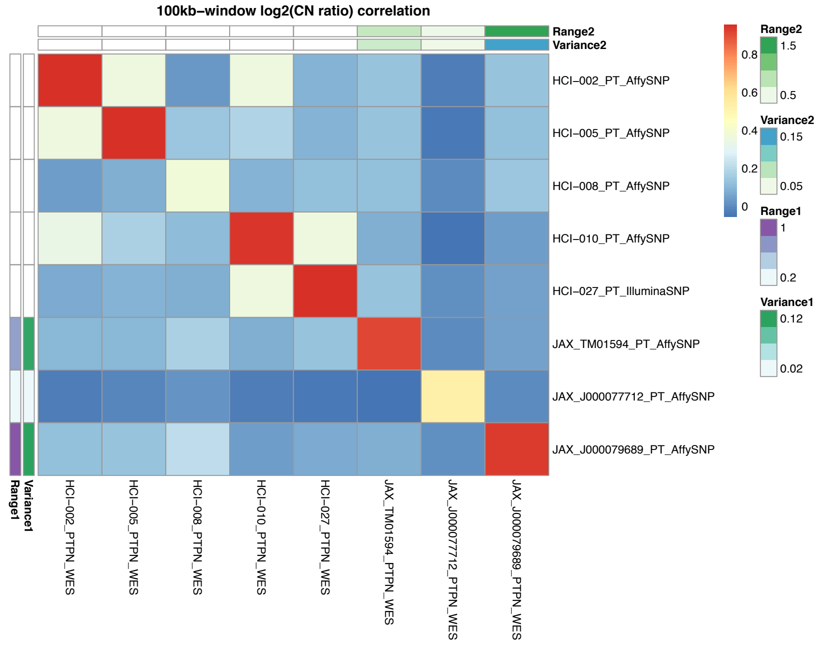

Supplementary Figure 18: Heatmap representing the Pearson correlation coefficients of the $\log_2$(CN ratio) of 100kb-windows binned from copy number segments of CNA profiles between matched samples estimated from SNP array and WES. The variance and range (5 – 95 percentile) values were calculated from the $\log_2$(copy number ratio) across all 100kb-windows per sample.

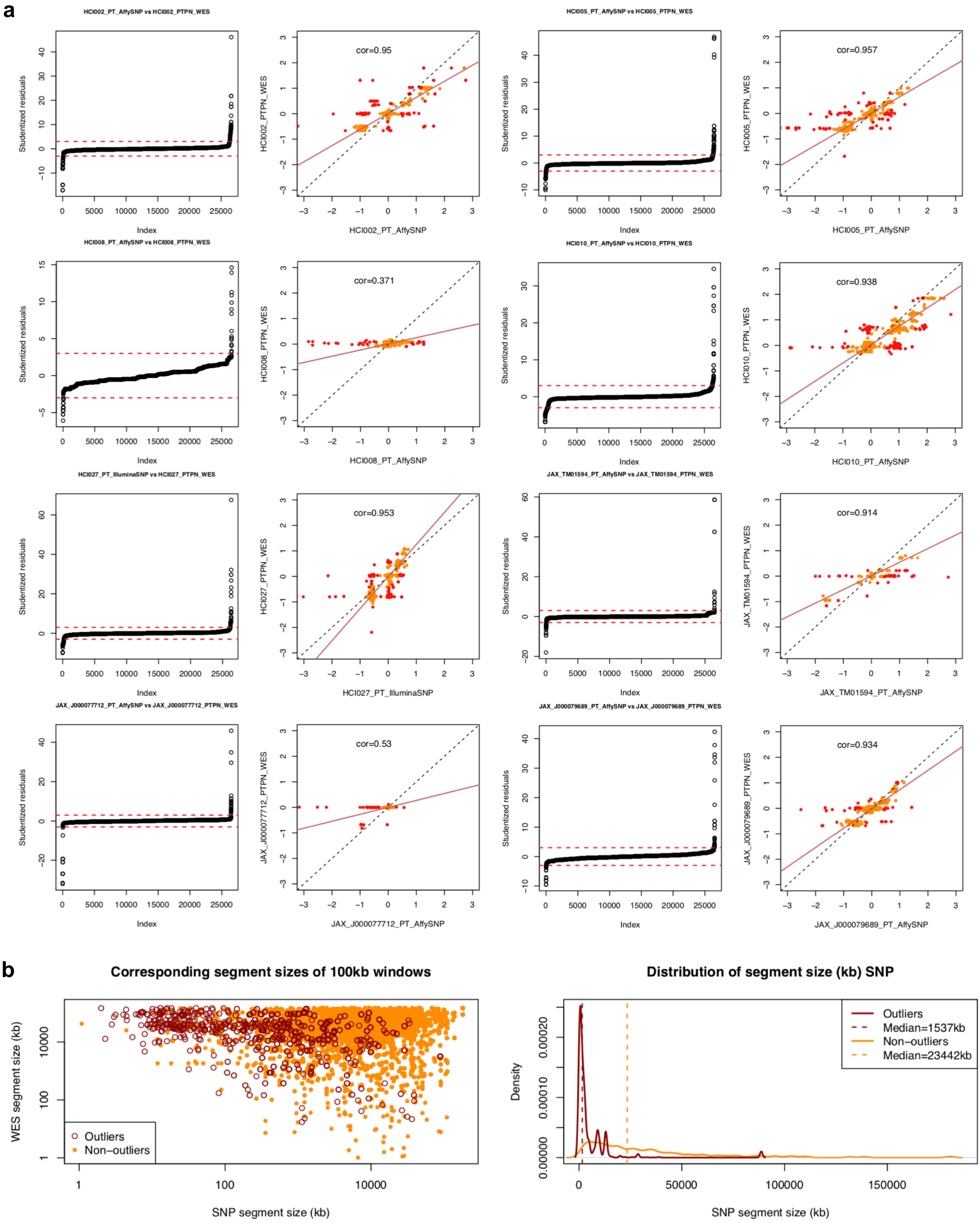

Supplementary Figure 19: (**a**) Pearson correlation and linear regression of the $\log_2$(CN ratio) of 100kb-windows binned from copy number segments, of CNA profiles between matched patient tumor samples estimated from SNP array and WES. Outliers of the linear regression (red points) are identified by studentized residuals > 3 and < -3. (**b**) Comparison of segment sizes between the combined outlier and non-outliers in (**a**).

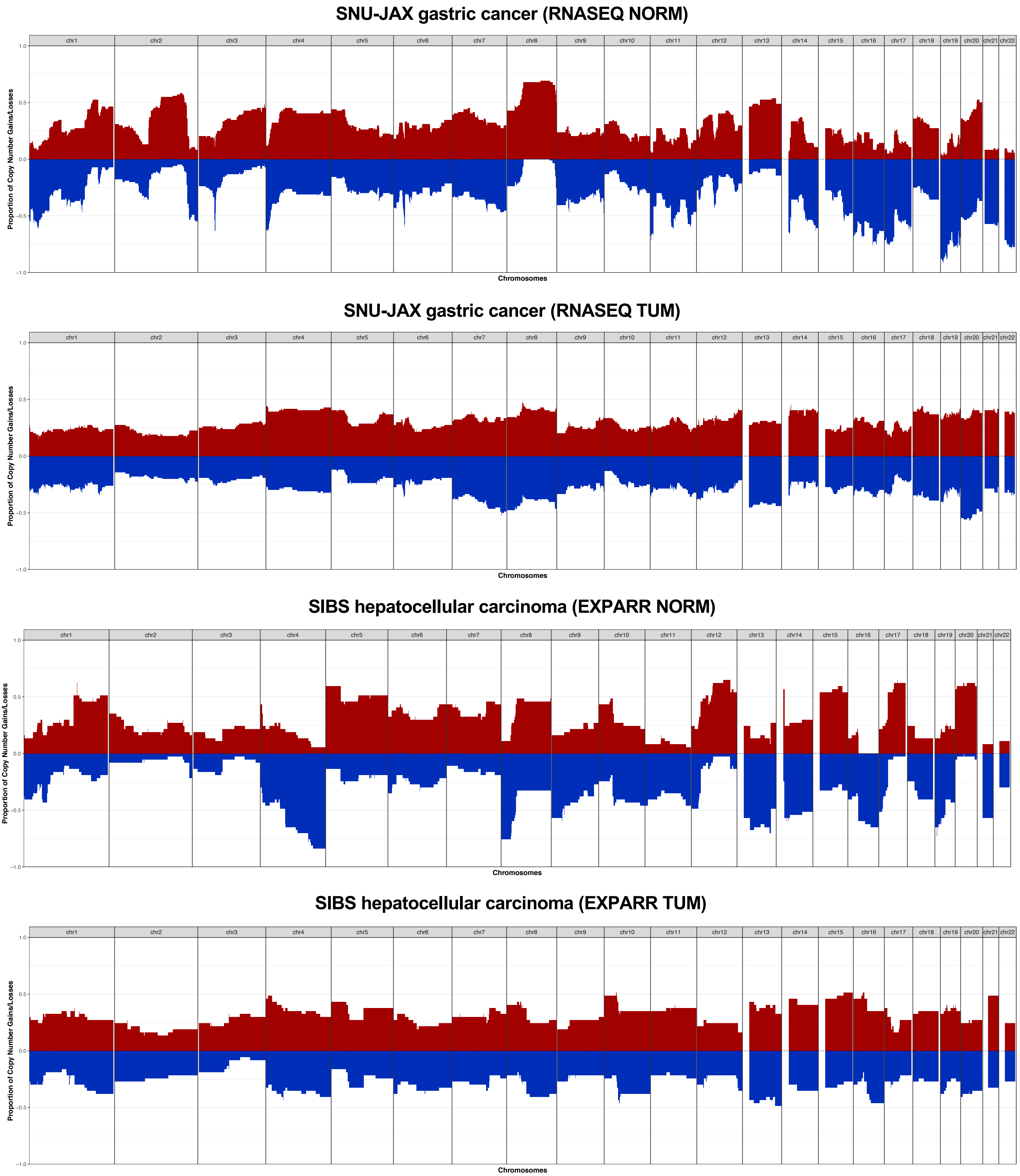

Supplementary Figure 20: Frequencies of copy number gains (log2(CN ratio) > 0.1) and losses (log2(CN ratio) < –0.1) estimated from RNA-Seq and gene expression array normalized by median expression of normal samples of the same tumor type (RNASEQ NORM, EXPARR NORM) or median expression of same set of patient tumors (RNASEQ TUM, EXPARR TUM) (see Supplementary Table 3).

**a**

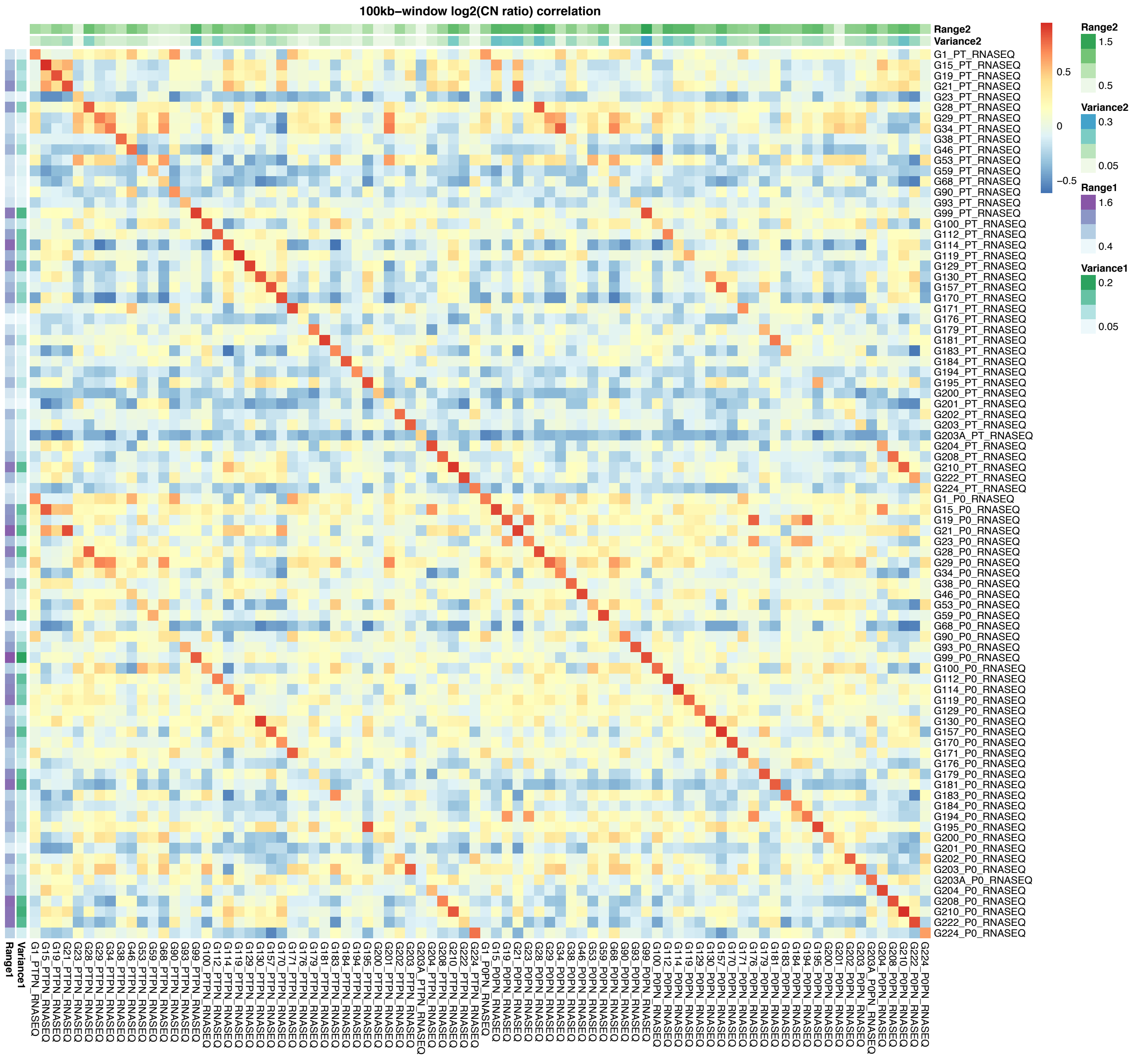

Supplementary Figure 21: (Continue next page)

**b**

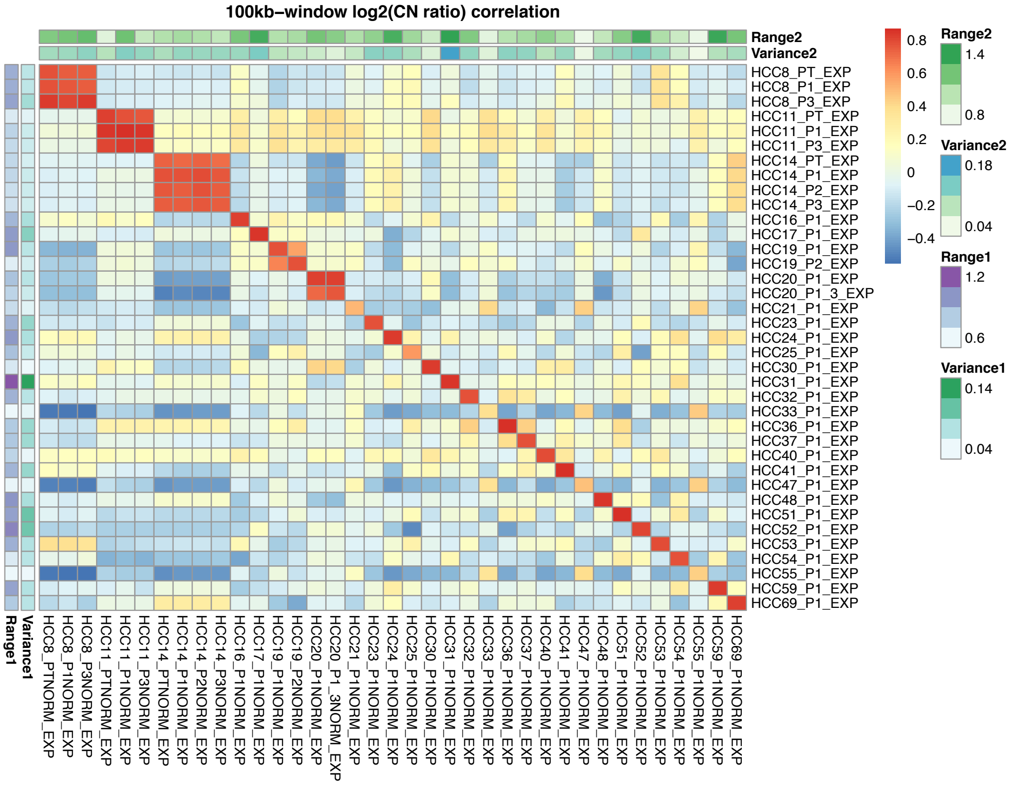

Supplementary Figure 21: Heatmap representing the Pearson correlation coefficients of the $log_2$(CN ratio) of 100kb-windows binned from copy number segments of CNA profiles estimated from (**a**) RNA-Seq (RNASEQ NORM vs TUM) and (**b**) gene expression array (EXPARR NORM vs TUM), between matched samples normalized by median expression of normal samples of the same tumor type and median expression of same set of patient tumors. The variance and range (5 – 95 percentile) values were calculated from the $log_2$(copy number ratio) across all 100kb-windows per sample.

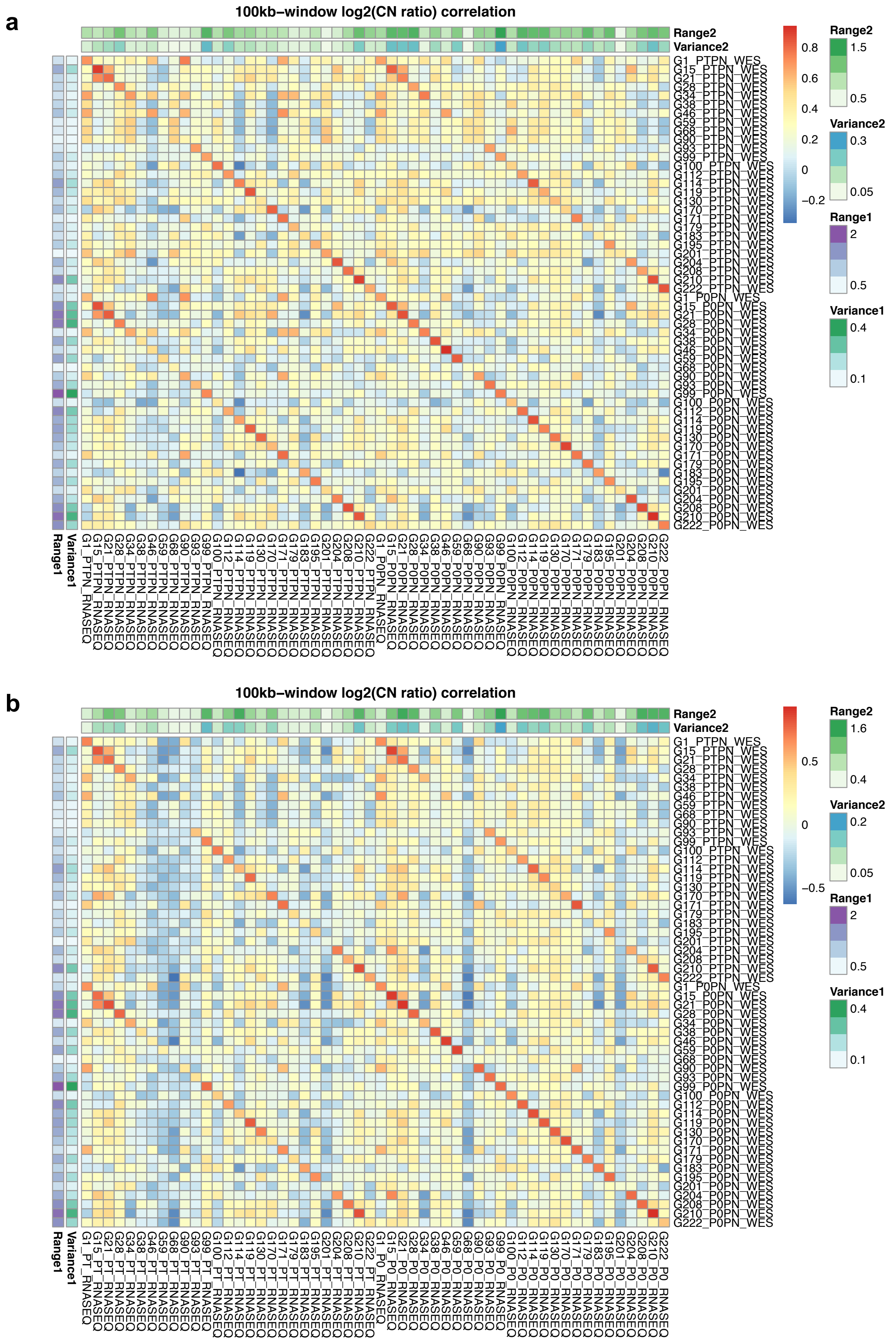

Supplementary Figure 22: Heatmap representing the Pearson correlation coefficients of the $log_2$(CN ratio) of 100kb-windows binned from copy number segments of CNA profiles between matched samples estimated from WES and RNA-Seq, (**a**) normalized by median expression of normal samples of the same tumor type “WES vs RNASEQ (NORM)” or (**b**) median expression of same set of patient tumors “WES vs RNASEQ (TUM)” . The variance and range (5 – 95 percentile) values were calculated from the $log_2$(copy number ratio) across all 100kb-windows per sample.

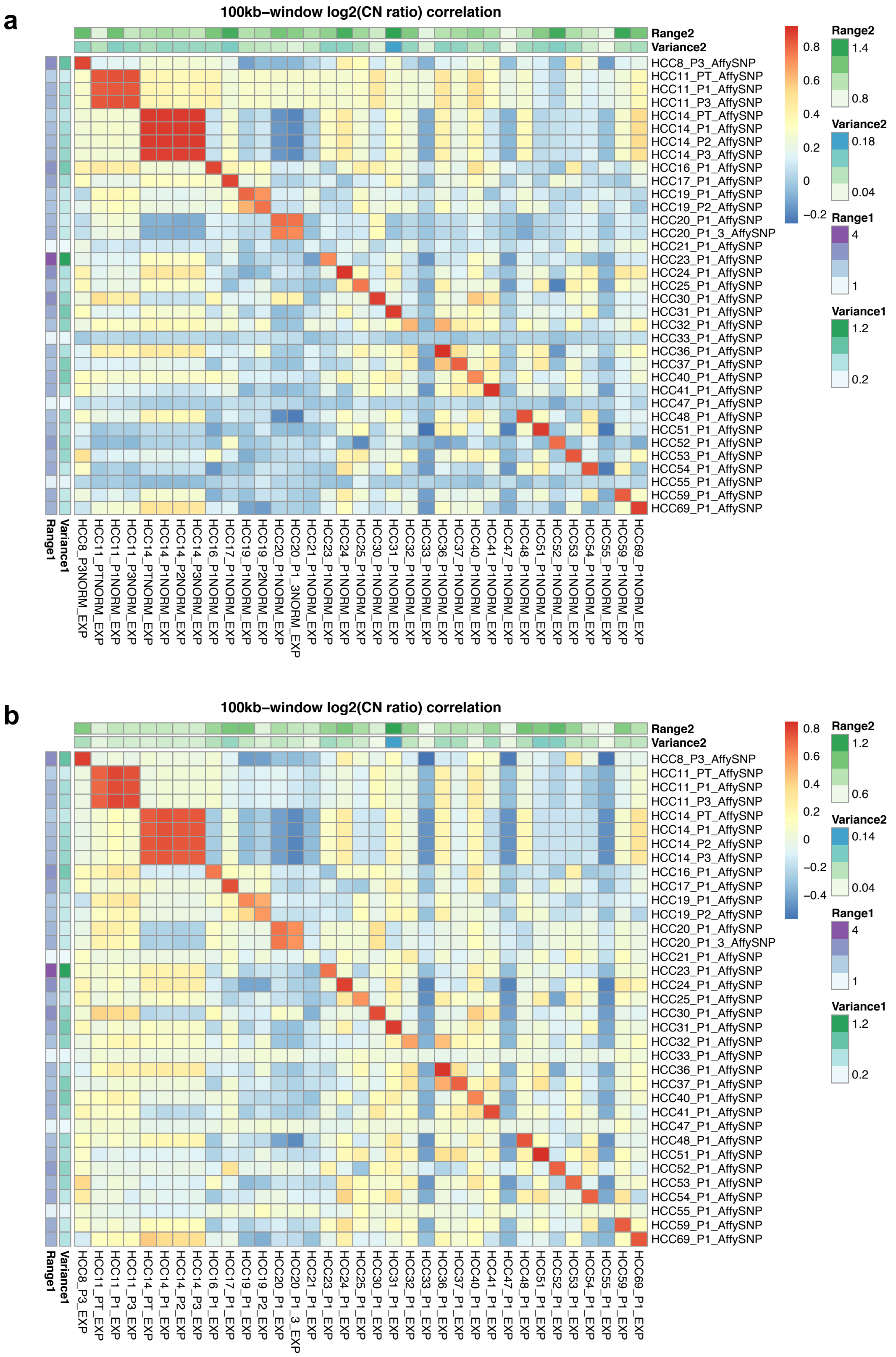

Supplementary Figure 23: Heatmap representing the Pearson correlation coefficients of the $\log_2$(CN ratio) of 100kb-windows binned from copy number segments of CNA profiles between matched samples estimated from SNP array and gene expression microarray, (**a**) normalized by median expression of normal samples of the same tumor type ”SNP vs EXPARR (NORM)” or (**b**) median expression of same set of patient tumors ”SNP vs EXPARR (TUM)”. The variance and range (5 – 95 percentile) values were calculated from the $\log_2$(copy number ratio) across all 100kb-windows per sample.

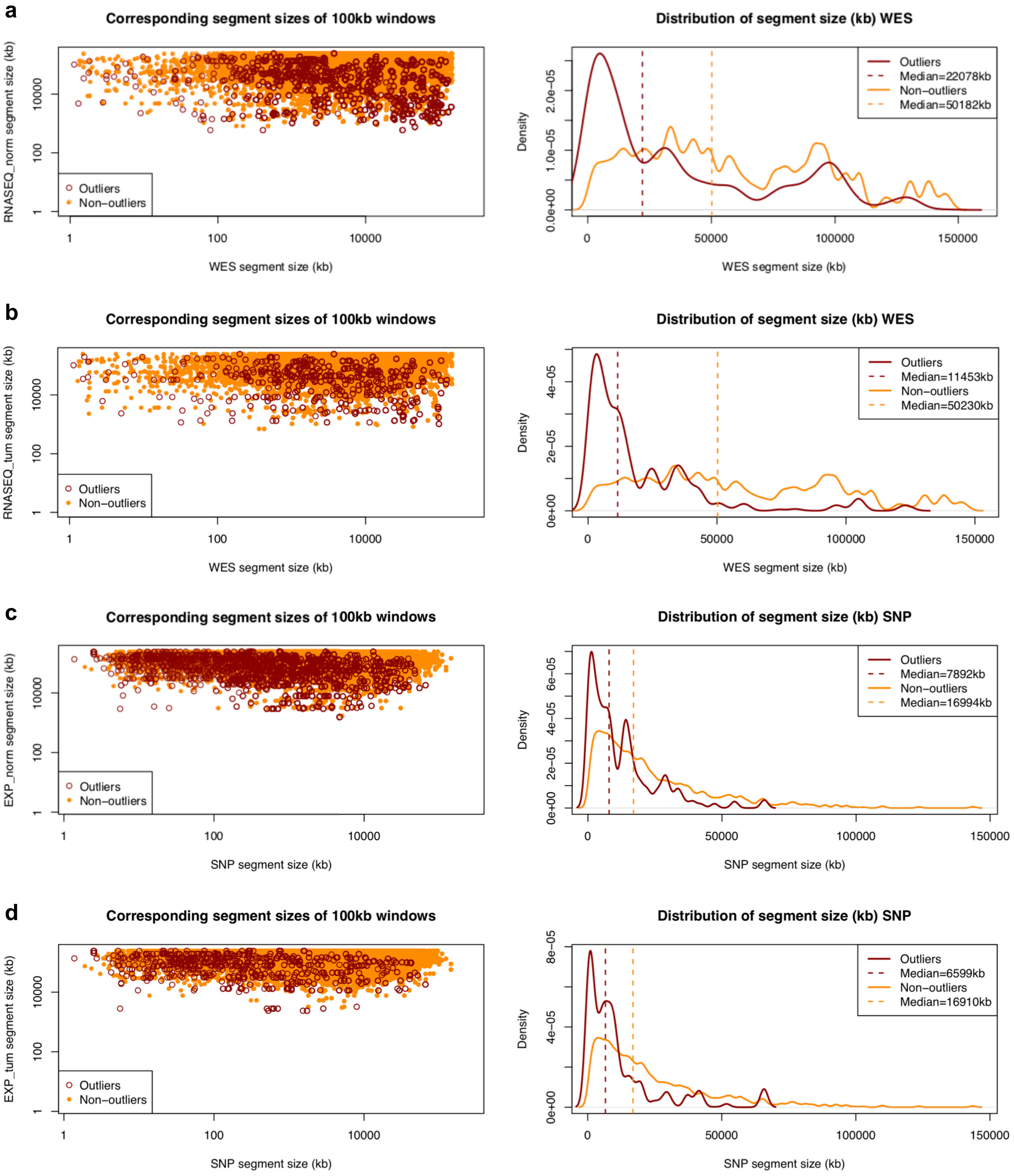

Supplementary Figure 24: Comparison of segment sizes between the combined outlier and non-outliers in (**a**) WES vs RNASEQ (NORM), (**b**) WES vs RNASEQ (TUM), (**c**) SNP vs EXPARR (NORM), and (**d**) SNP vs EXPARR (TUM) (See Supplementary Table 3).

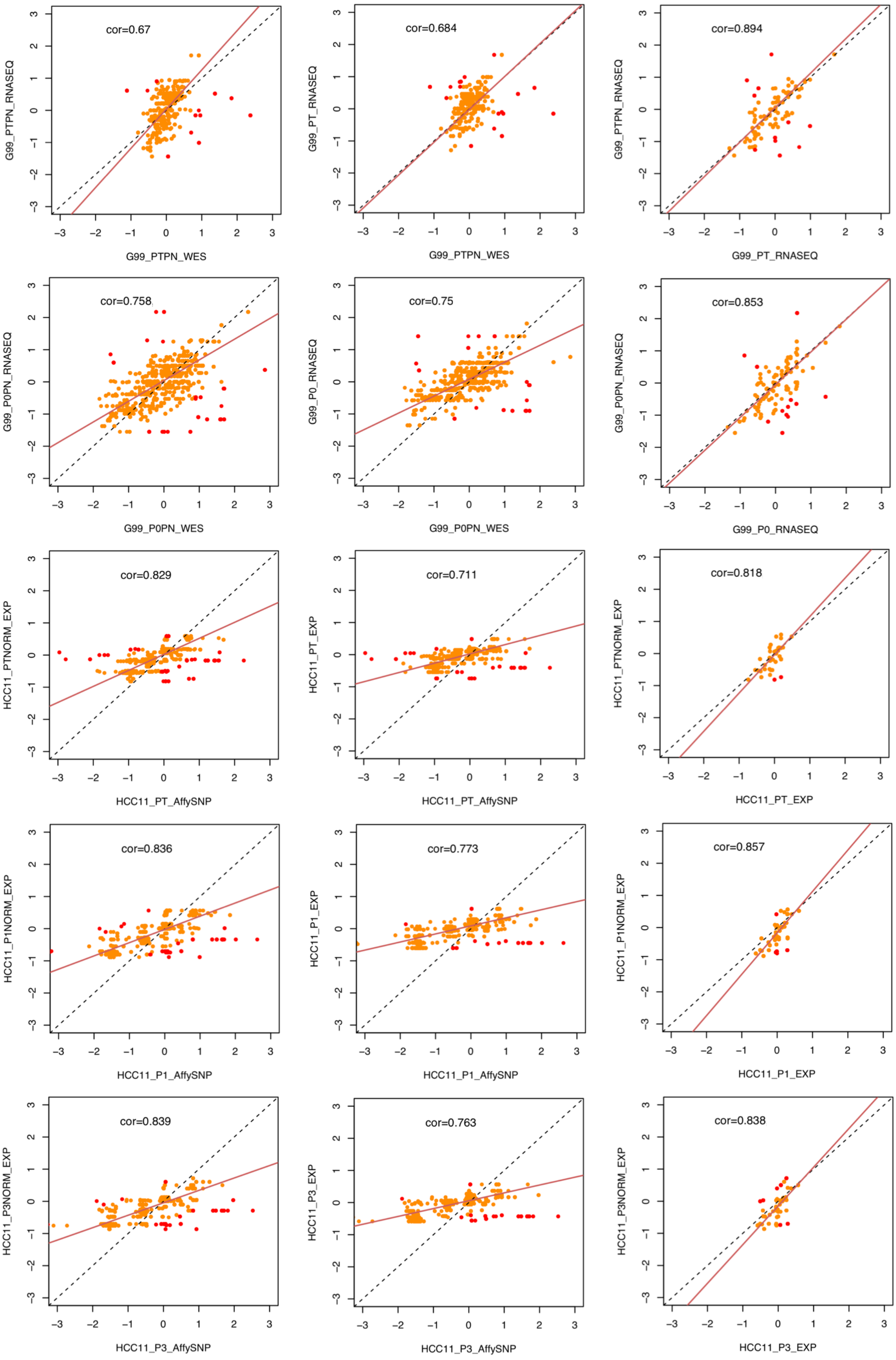

Supplementary Figure 25: Pearson correlation and linear regression of the $log_2$(CN ratio) of 100kb-windows binned from copy number segments, of CNA profiles between matched patient tumor samples estimated from different platforms and analysis methods for examples shown in Fig. 2d. Outliers of the linear regression (red points) are identified by studentized residuals > 3 and < -3.

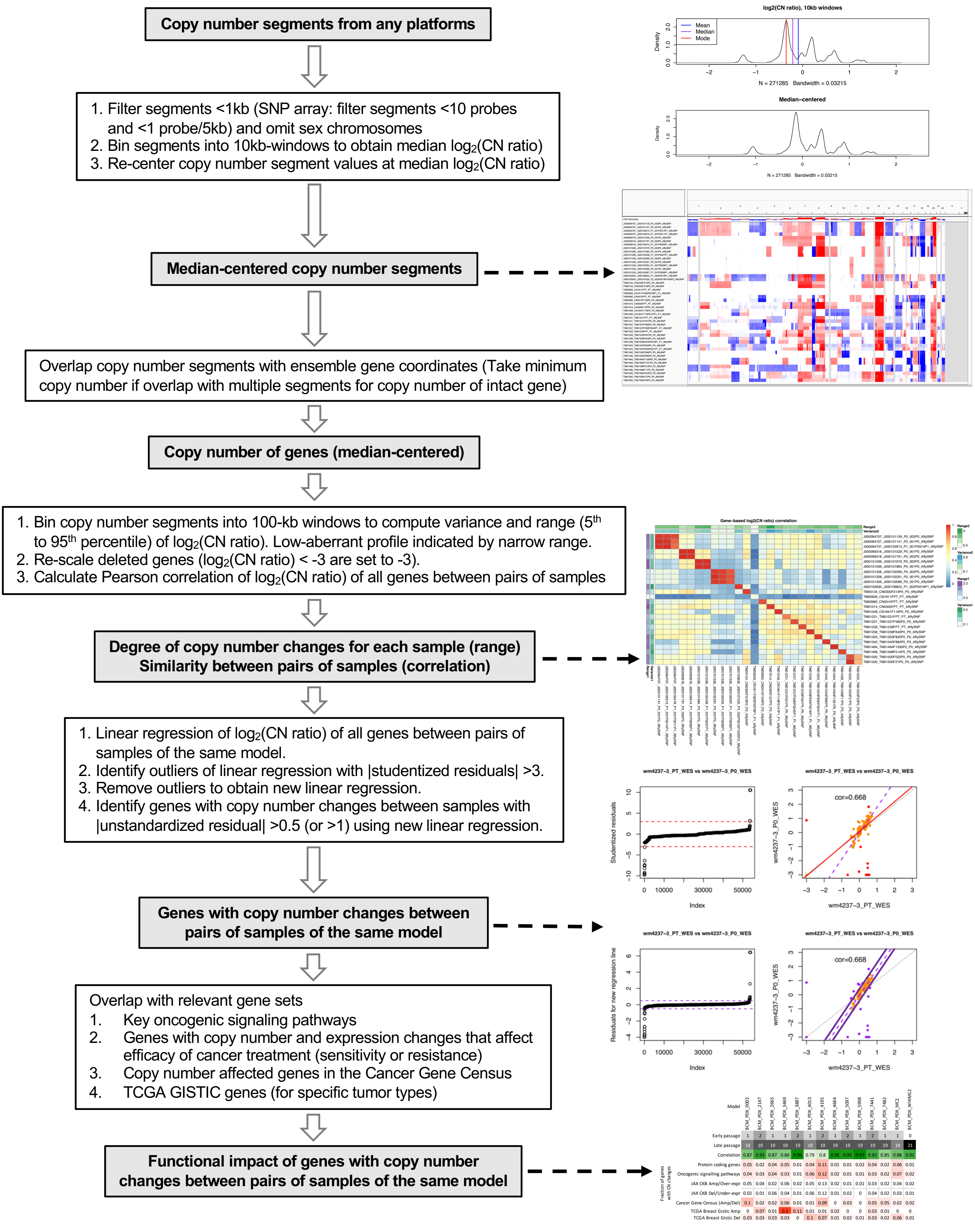

Supplementary Figure 26: A correlation and robust regression approach to quantify similarity of CNA profiles and identify genes with copy number changes between two samples.

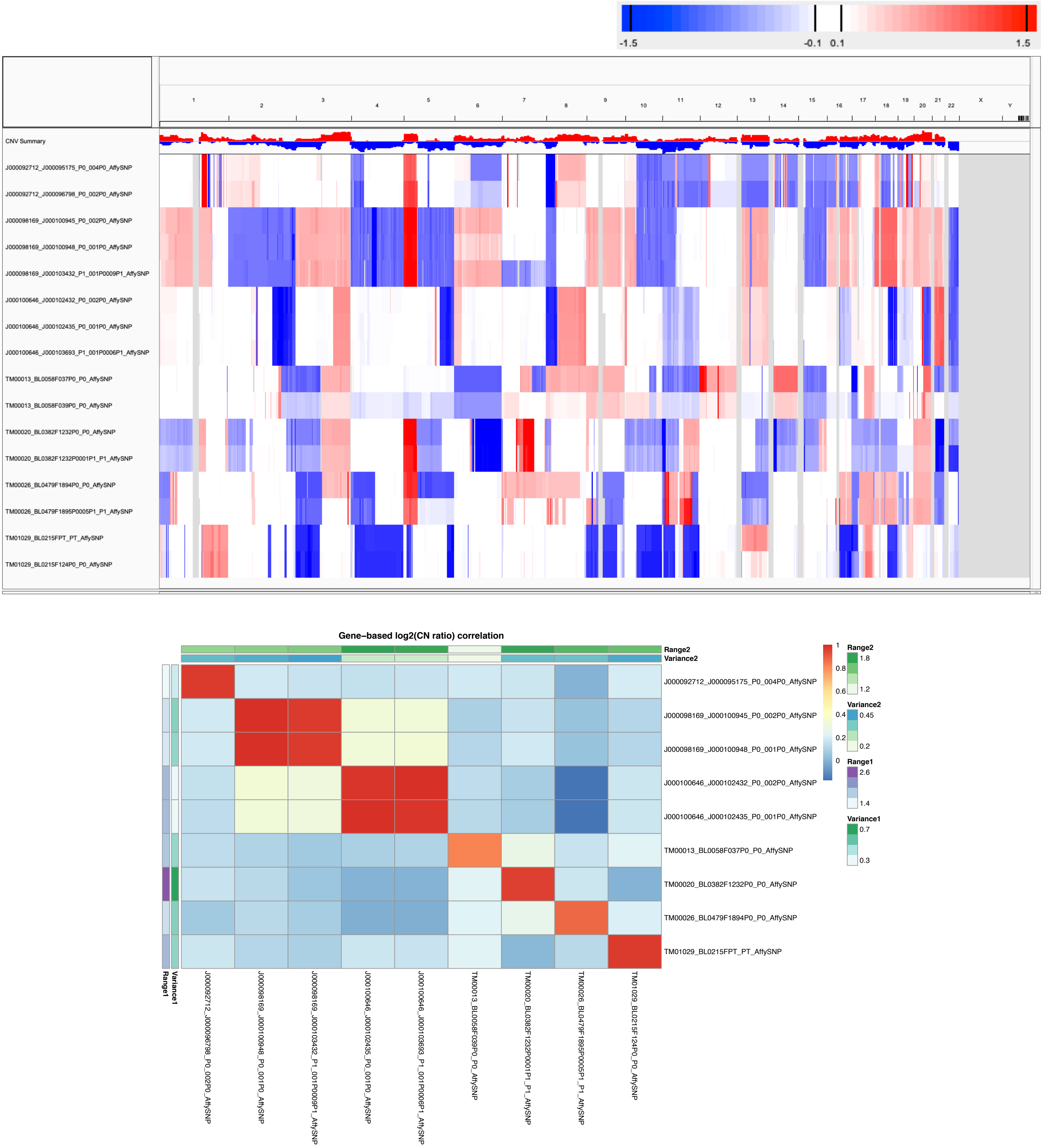

Supplementary Figure 27: CNA profiles (IGV heatmap) and correlation heatmap of gene-based copy number (log2(CN ratio), median centered) of samples from JAX SNP array bladder cancer dataset.

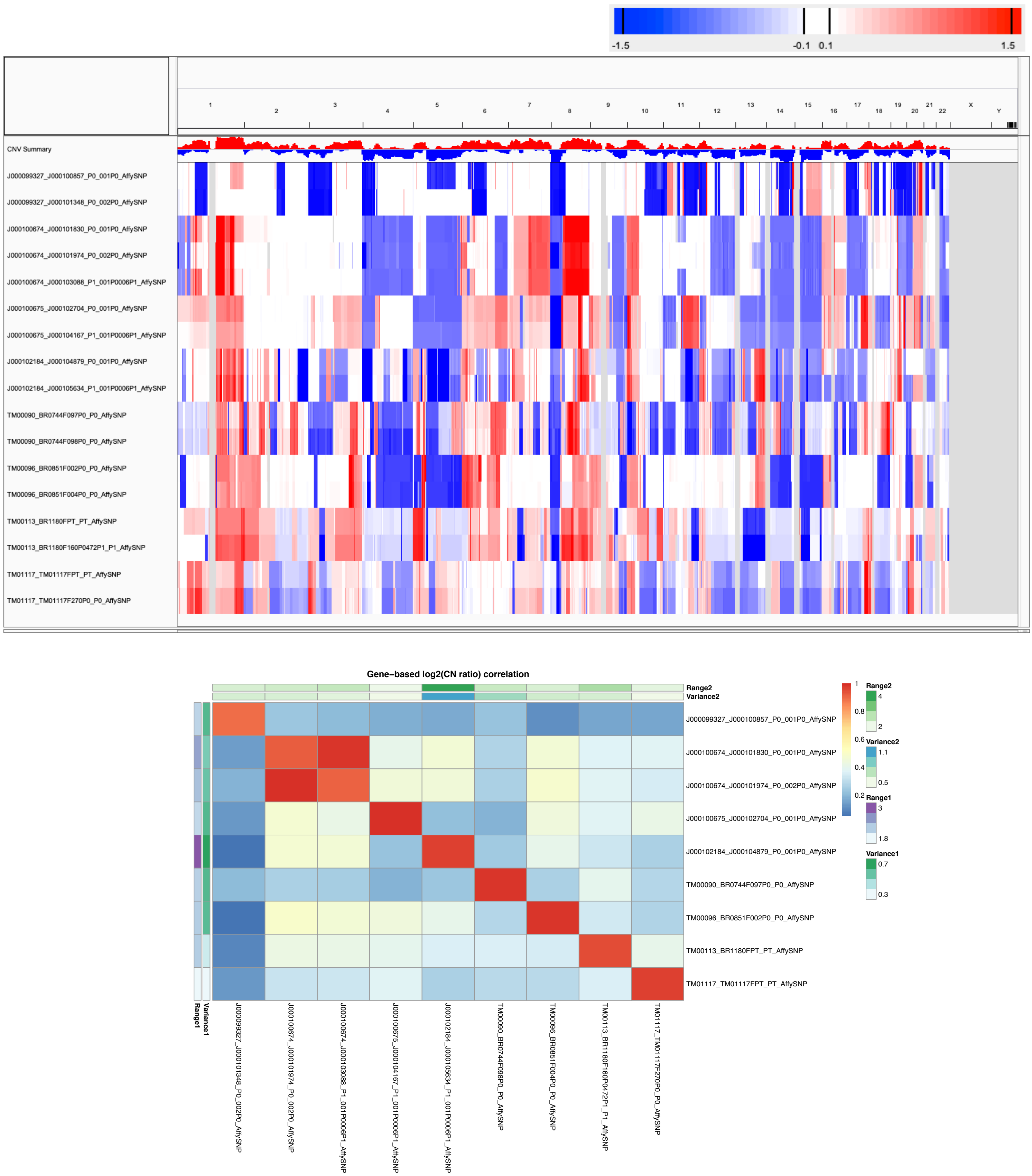

Supplementary Figure 28: CNA profiles (IGV heatmap) and correlation heatmap of gene-based copy number (log2(CN ratio), median centered) of samples from JAX SNP array breast cancer dataset.

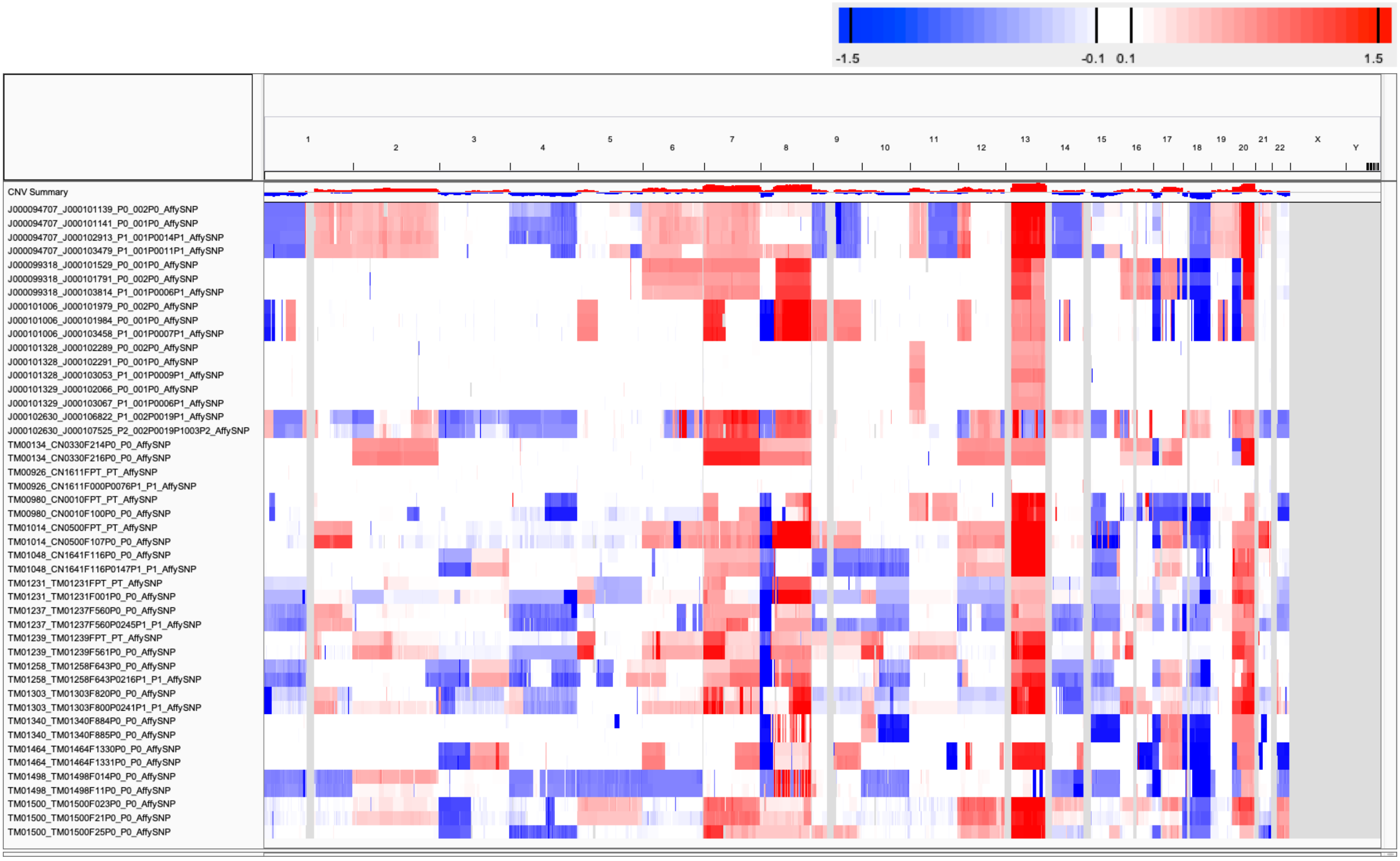

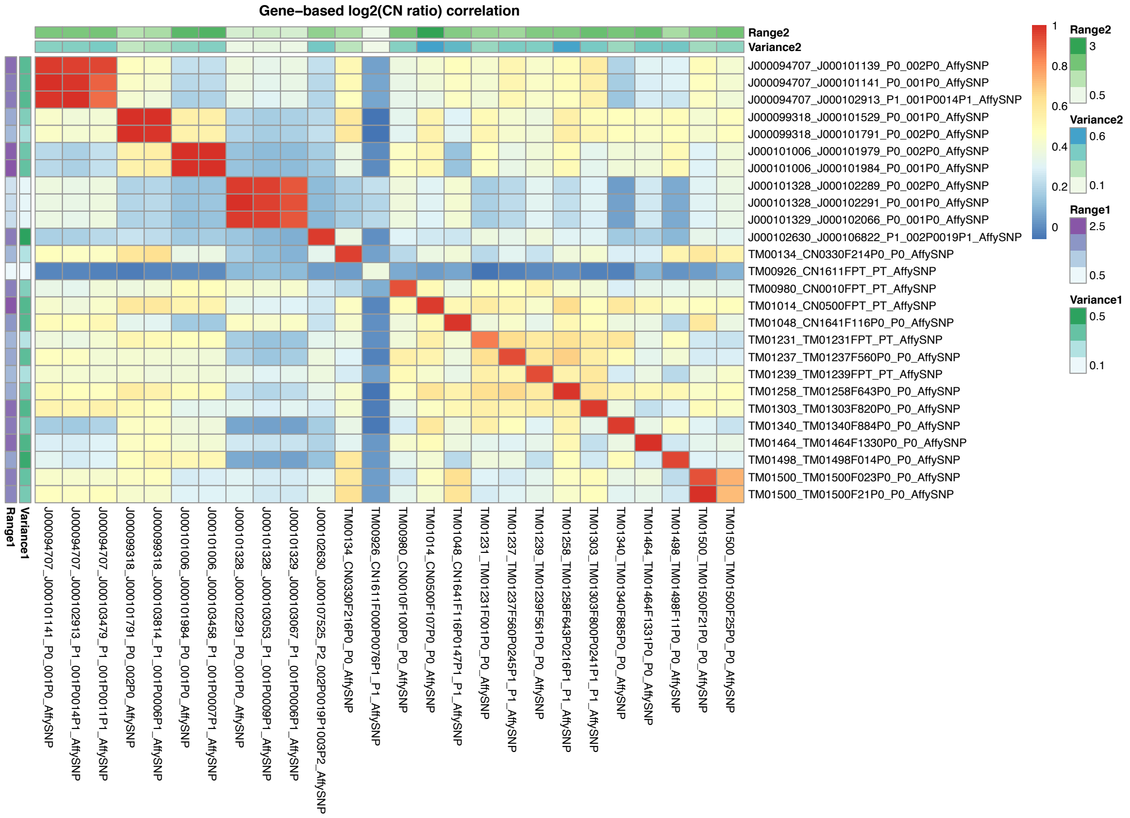

Supplementary Figure 29: CNA profiles (IGV heatmap) and correlation heatmap of gene-based copy number (log2(CN ratio), median centered) of samples from JAX SNP array colorectal cancer dataset.

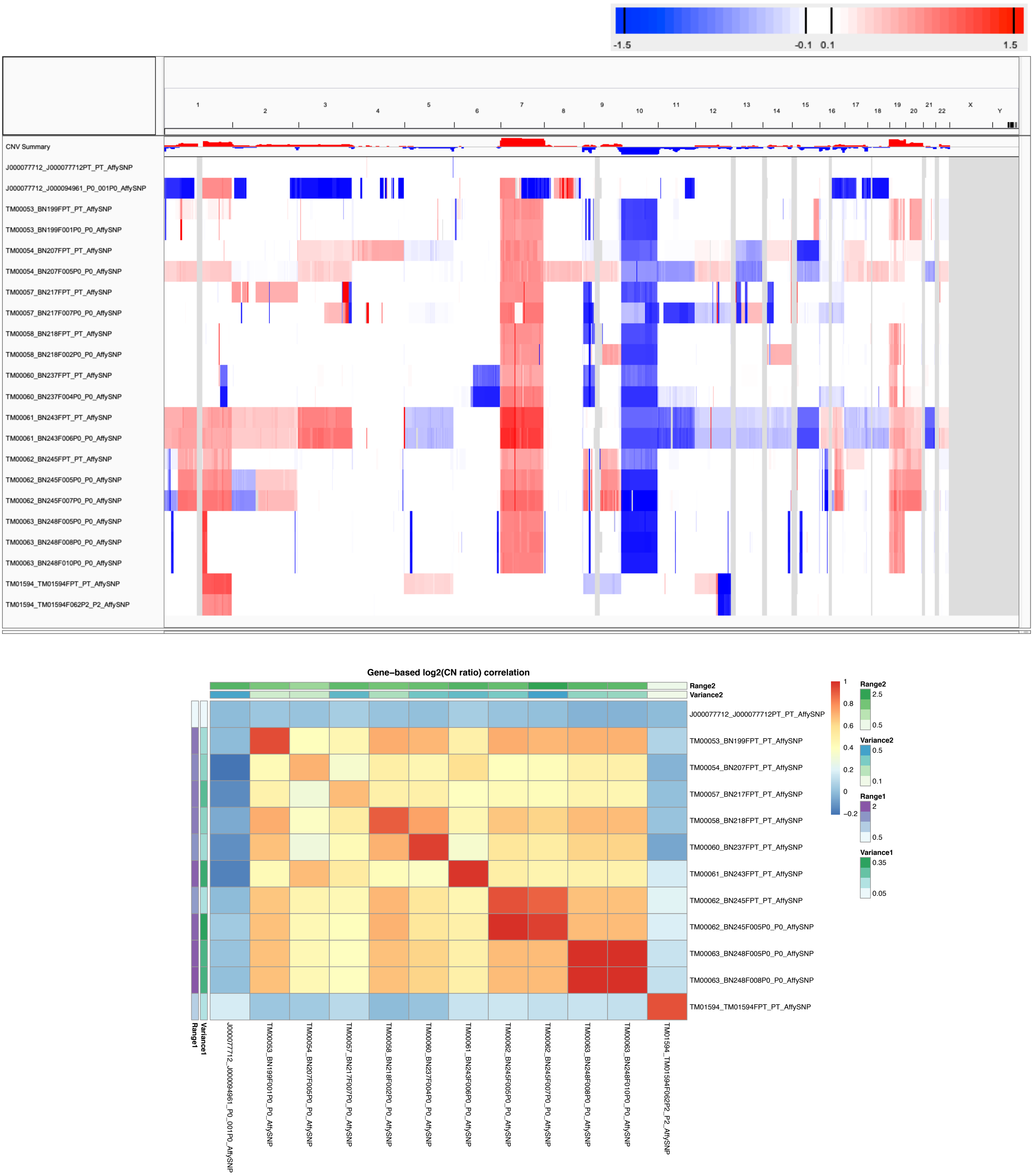

Supplementary Figure 30: CNA profiles (IGV heatmap) and correlation heatmap of gene-based copy number (log2(CN ratio), median centered) of samples from JAX SNP array glioblastoma multiforme (GBM) dataset.

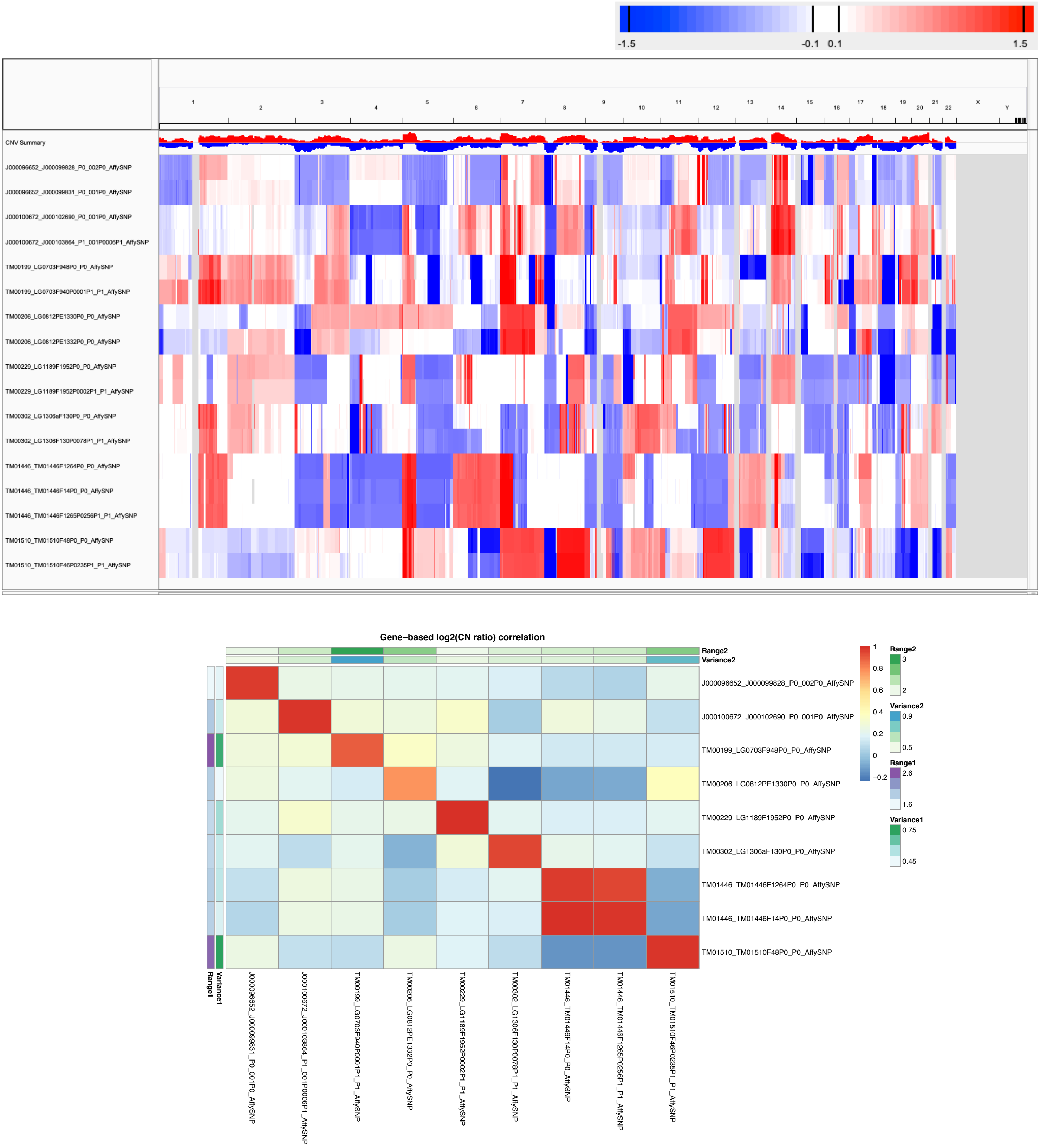

Supplementary Figure 31: CNA profiles (IGV heatmap) and correlation heatmap of gene-based copy number (log2(CN ratio), median centered) of samples from JAX SNP array lung adenocarcinoma (LUAD) dataset.

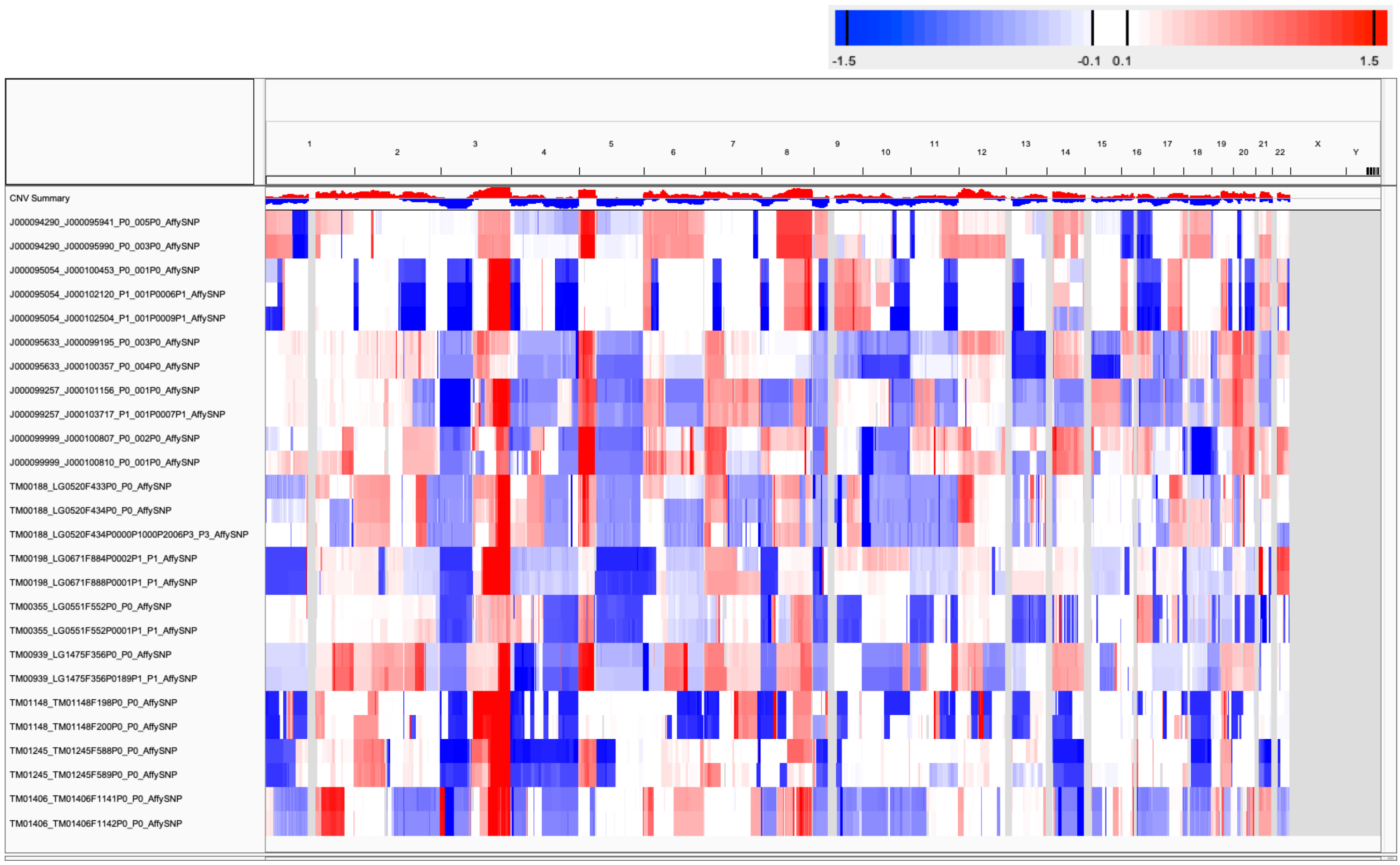

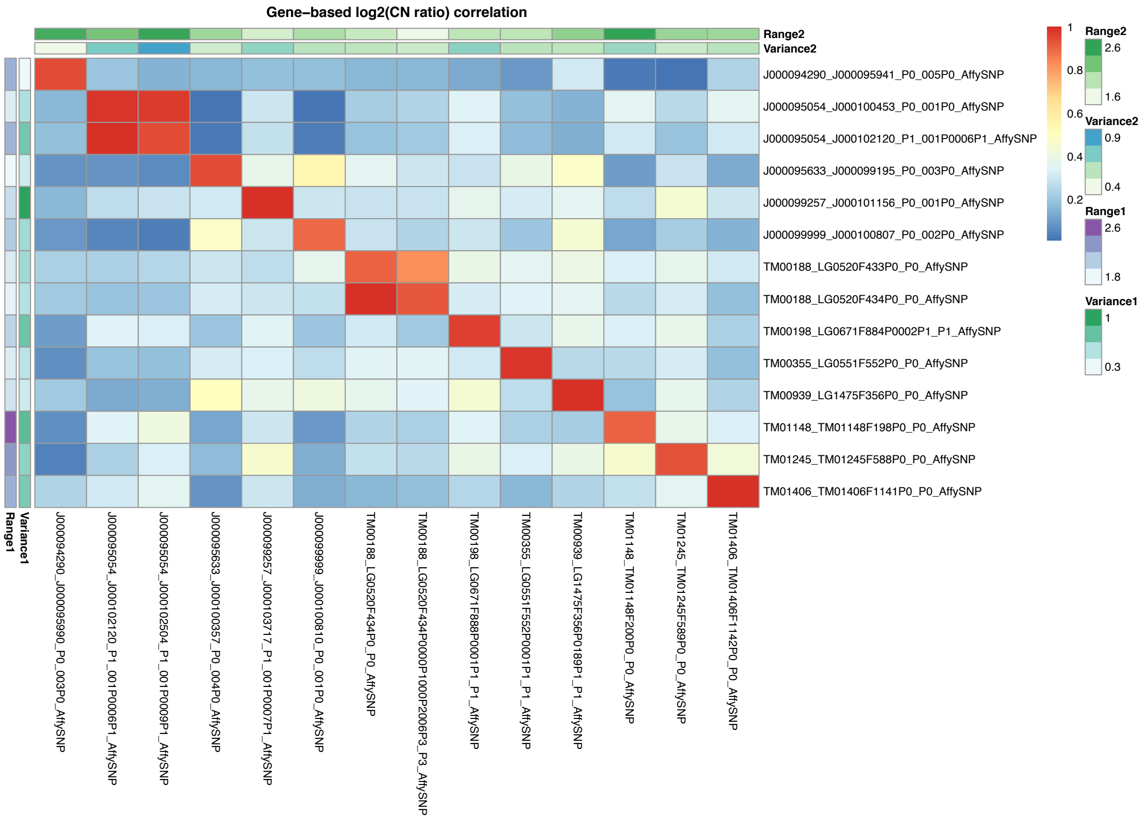

Supplementary Figure 32: CNA profiles (IGV heatmap) and correlation heatmap of gene-based copy number (log2(CN ratio), median centered) of samples from JAX SNP array lung squamous cell carcinoma (LUSC) dataset.

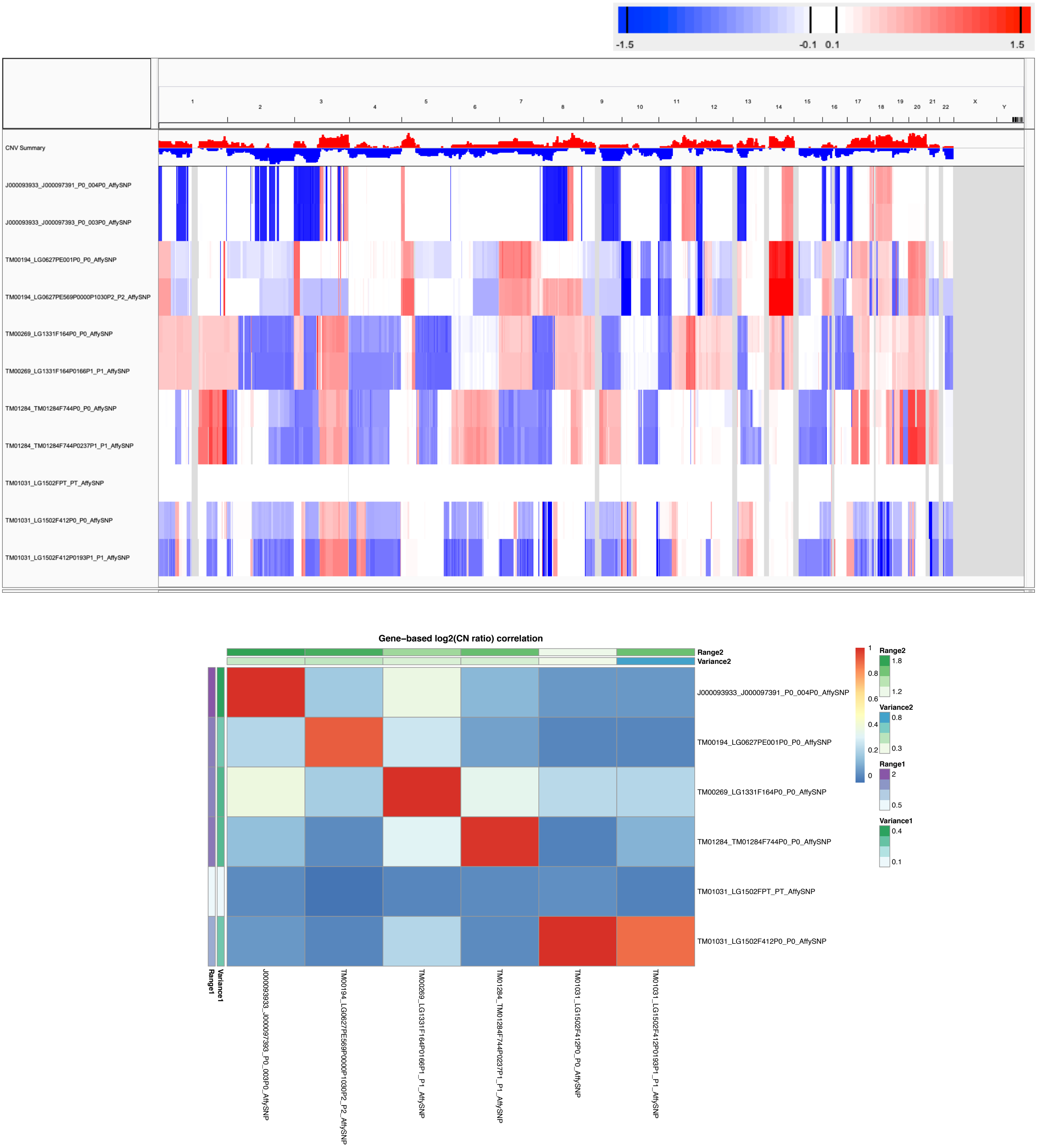

Supplementary Figure 33: CNA profiles (IGV heatmap) and correlation heatmap of gene-based copy number (log2(CN ratio), median centered) of samples from JAX SNP array other lung cancer subtypes dataset.

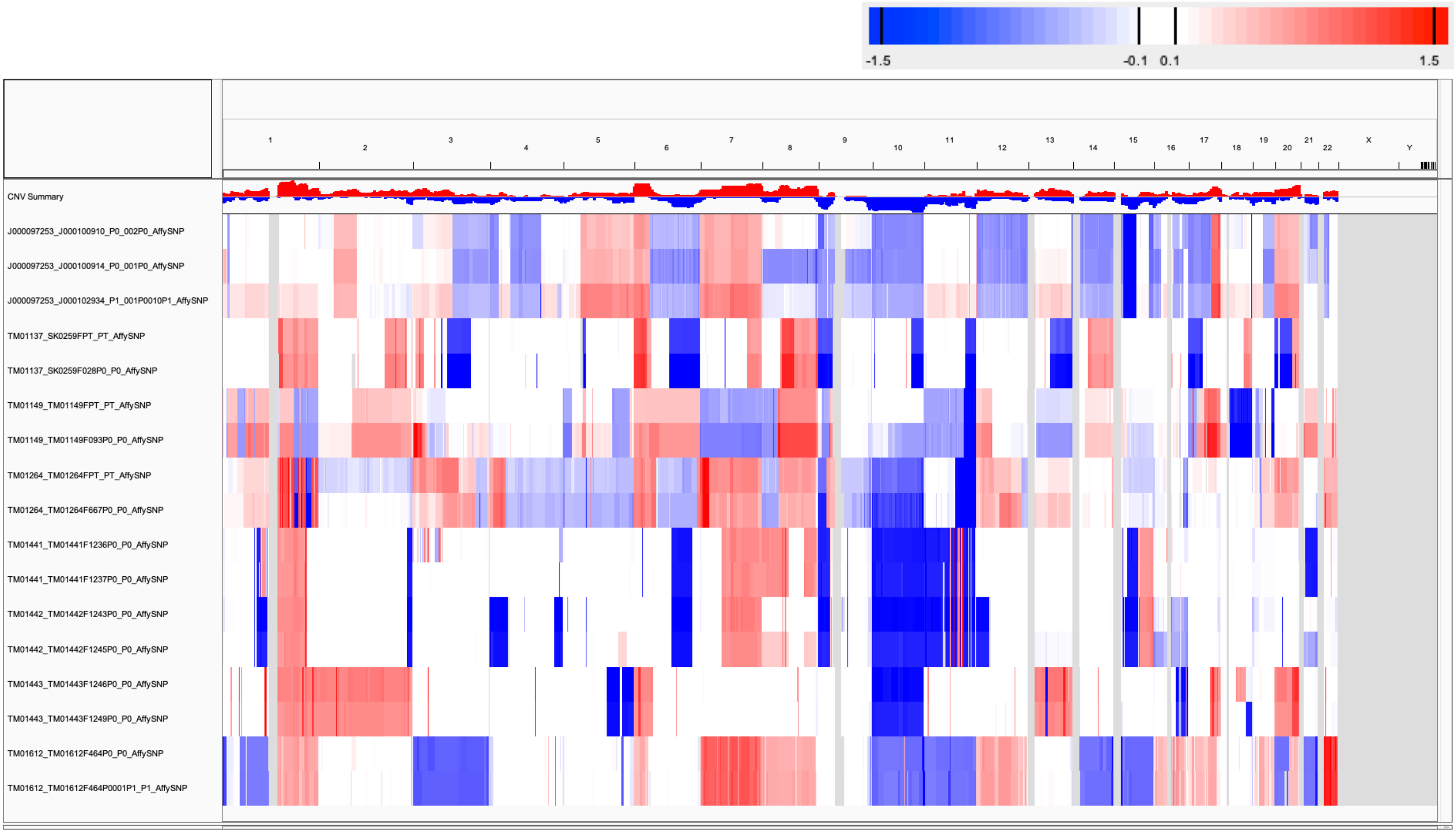

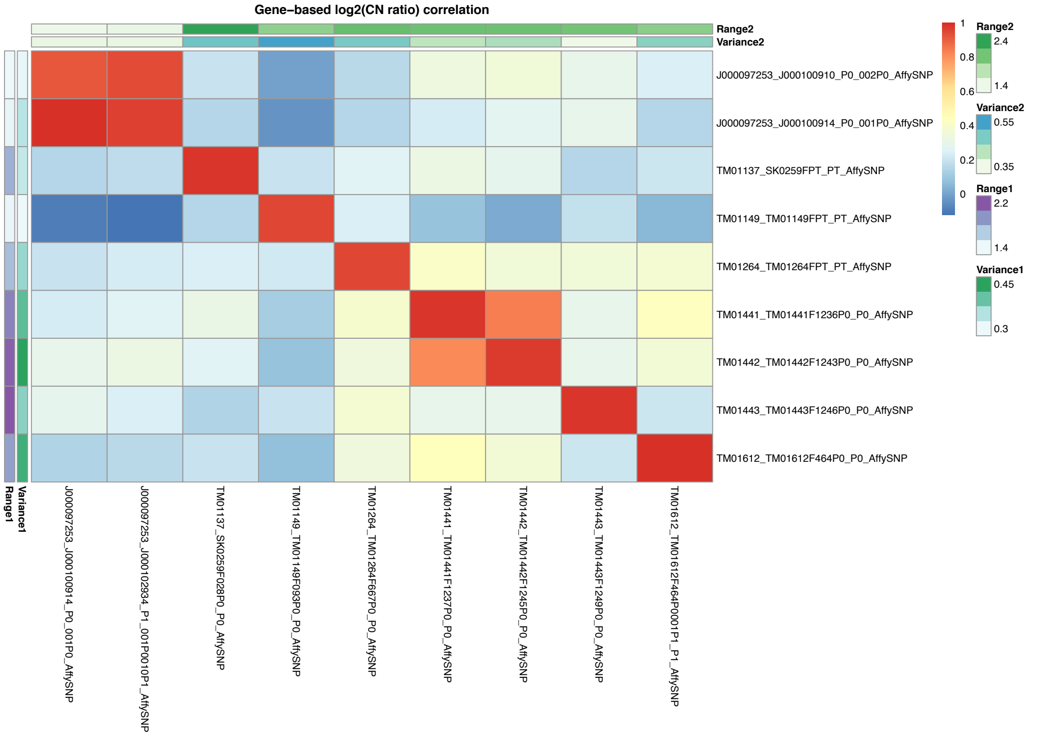

Supplementary Figure 34: CNA profiles (IGV heatmap) and correlation heatmap of gene-based copy number (log2(CN ratio), median centered) of samples from JAX SNP array skin melanoma dataset.

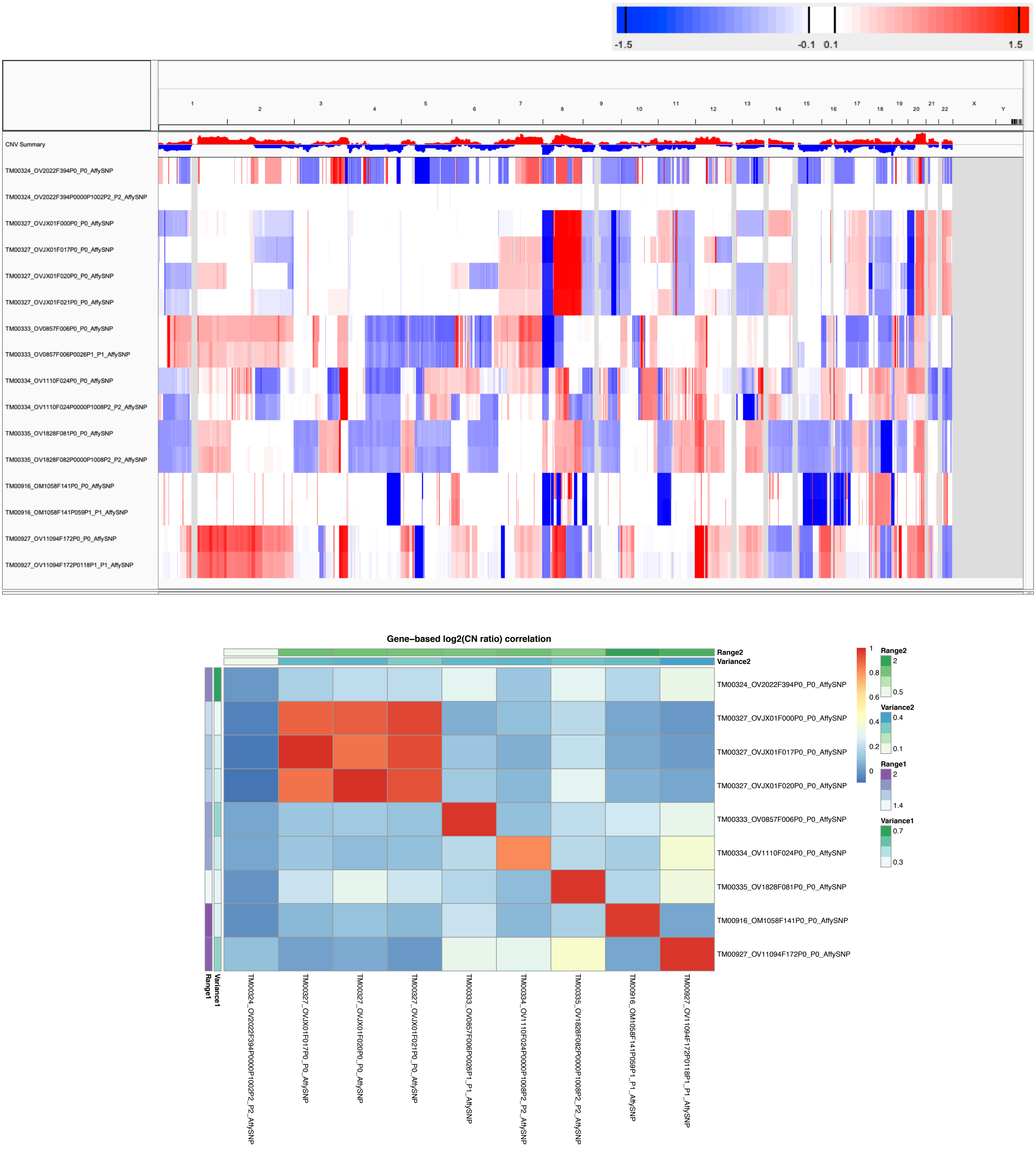

Supplementary Figure 35: CNA profiles (IGV heatmap) and correlation heatmap of gene-based copy number (log2(CN ratio), median centered) of samples from JAX SNP array ovarian cancer dataset.

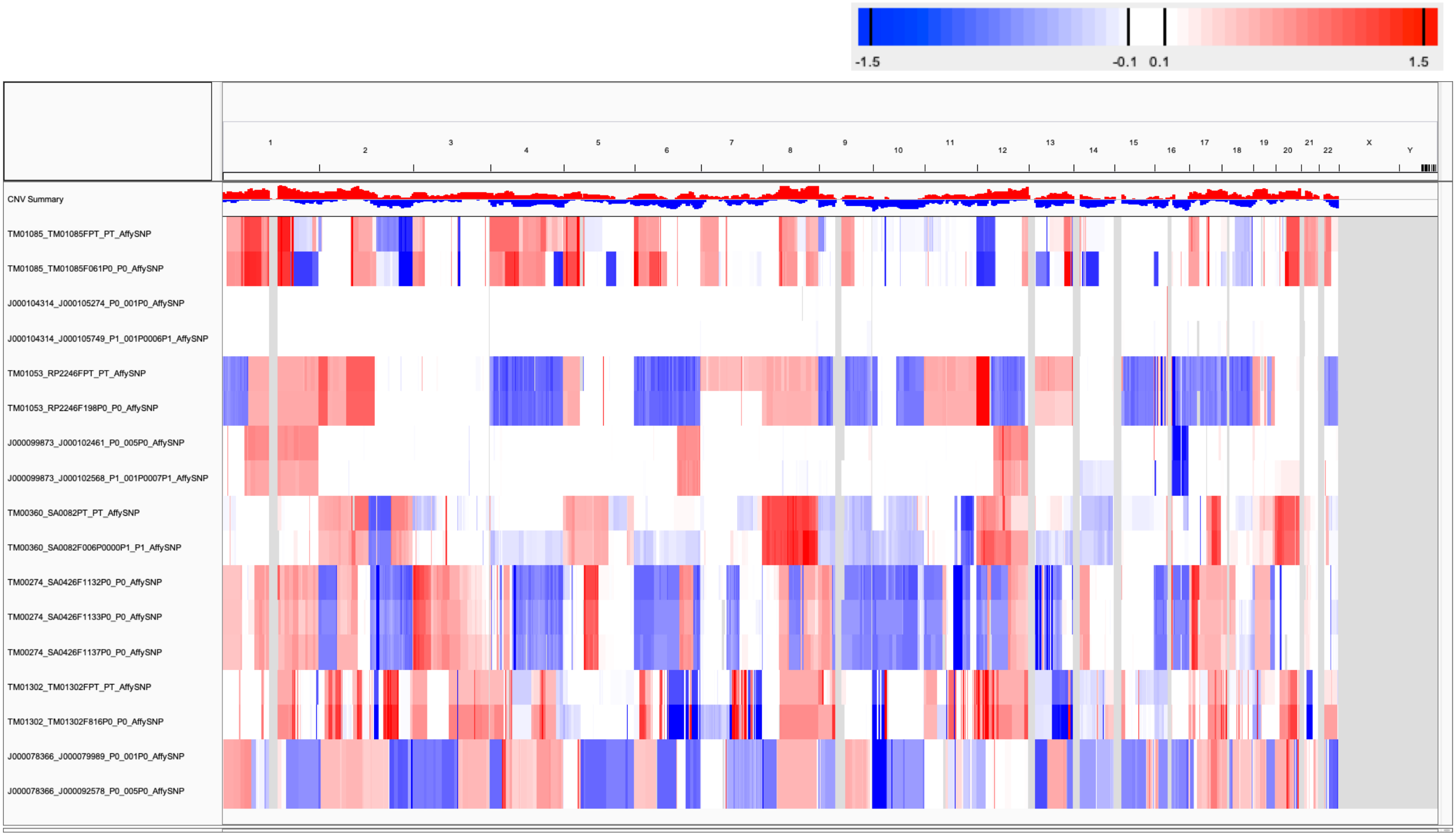

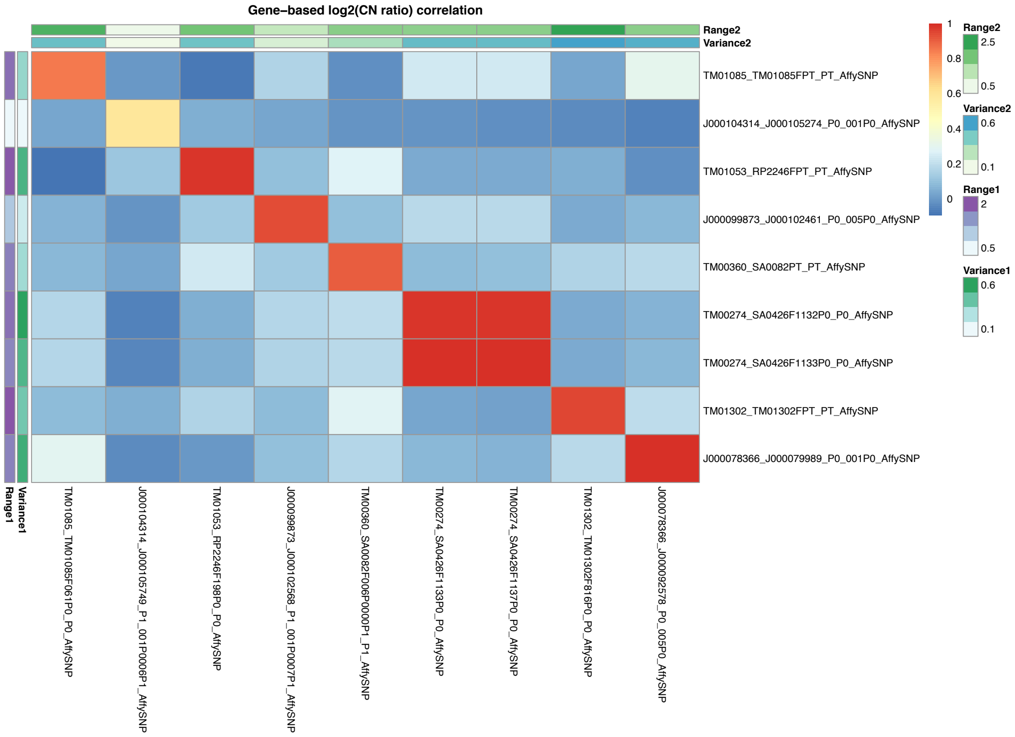

Supplementary Figure 36: CNA profiles (IGV heatmap) and correlation heatmap of gene-based copy number (log2(CN ratio), median centered) of samples from JAX SNP array sarcoma dataset.

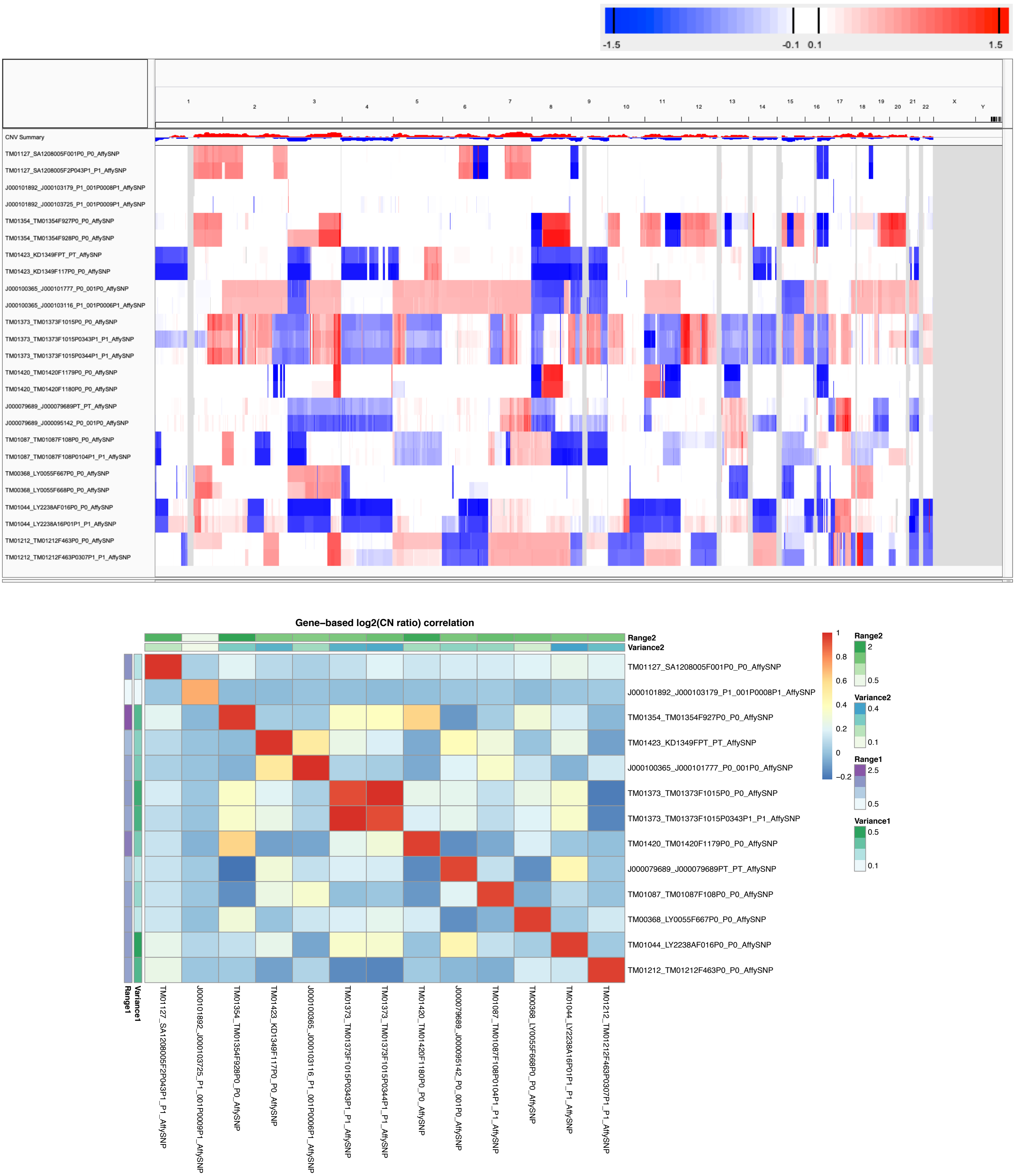

Supplementary Figure 37: CNA profiles (IGV heatmap) and correlation heatmap of gene-based copy number (log2(CN ratio), median centered) of samples from JAX SNP array other cancers dataset.

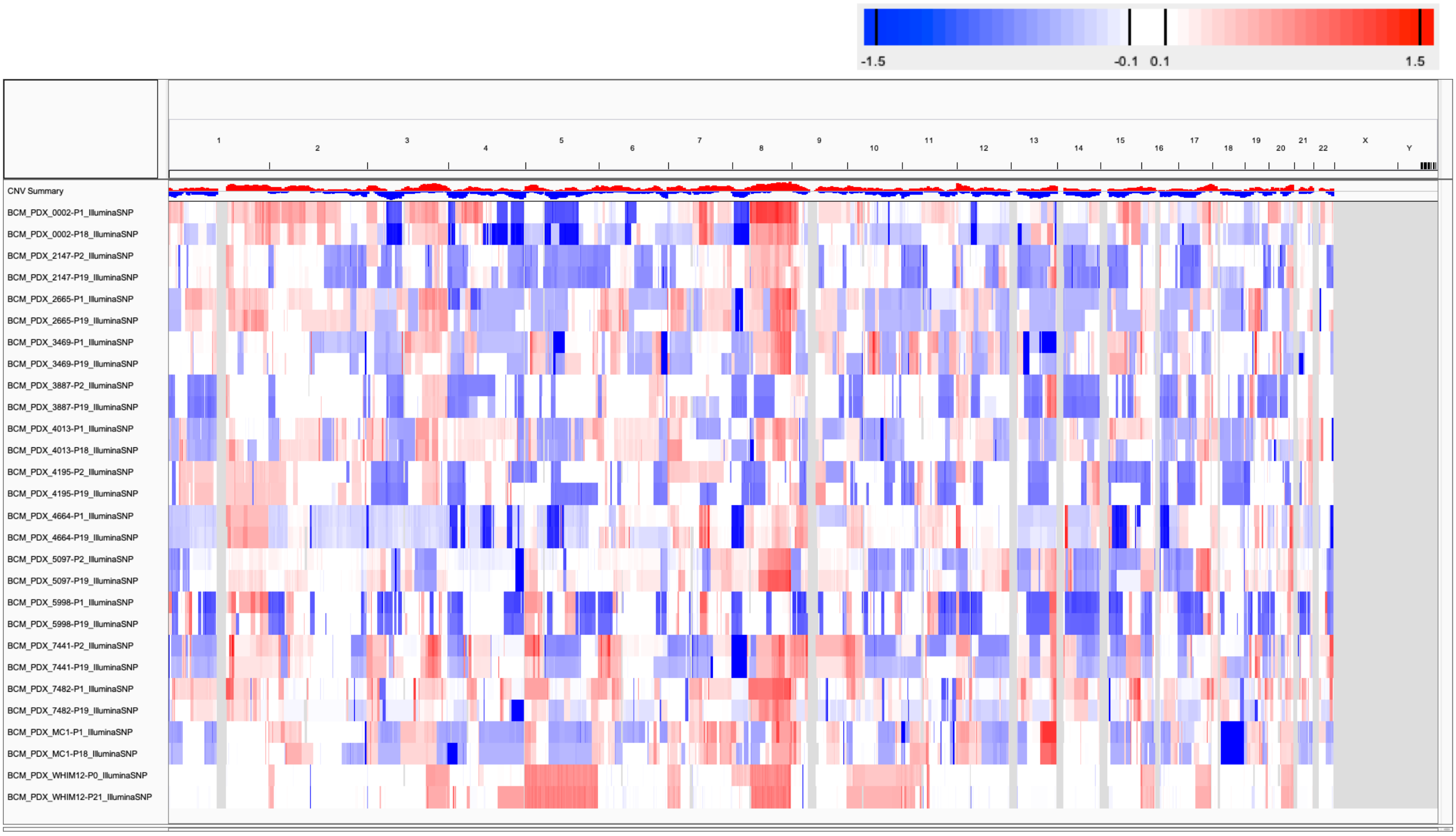

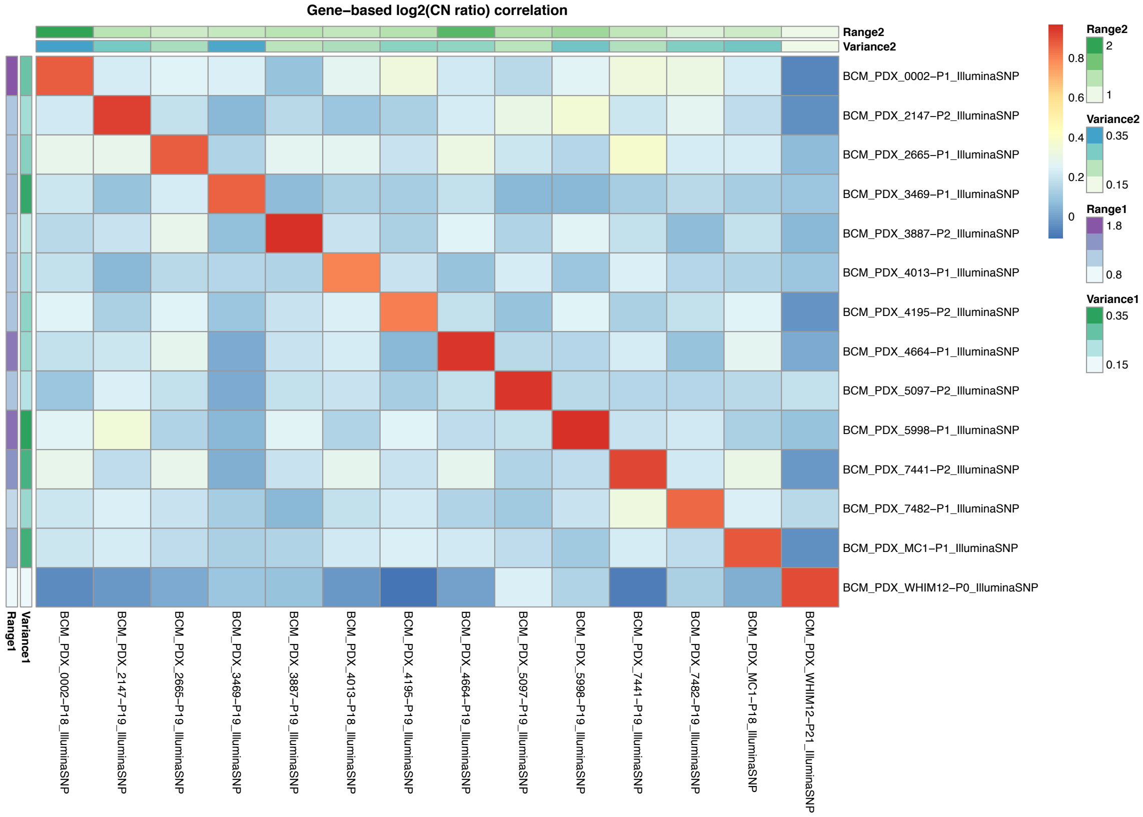

Supplementary Figure 38: CNA profiles (IGV heatmap) and correlation heatmap of gene-based copy number (log2(CN ratio), median centered) of samples from BCM SNP array breast cancer dataset.

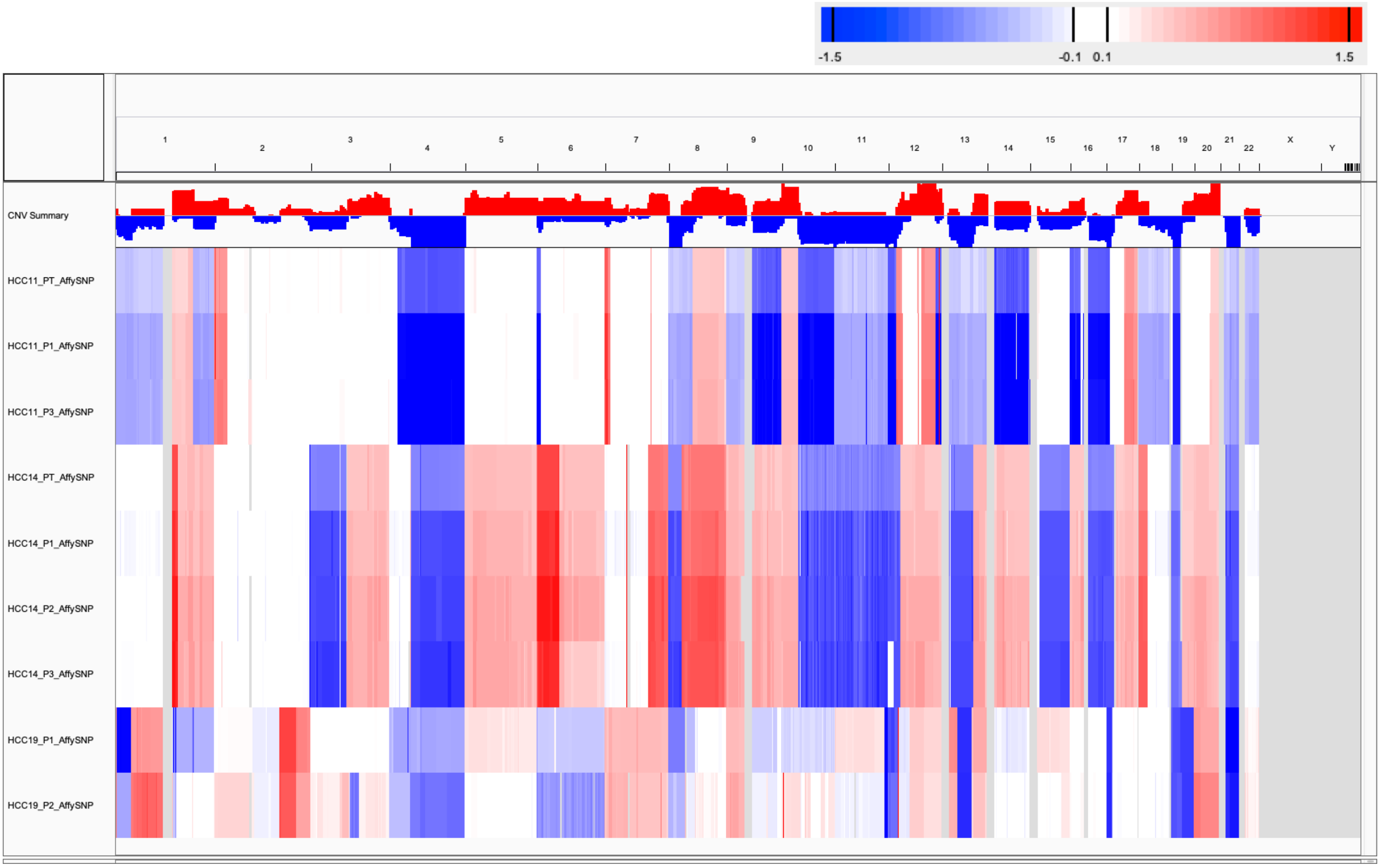

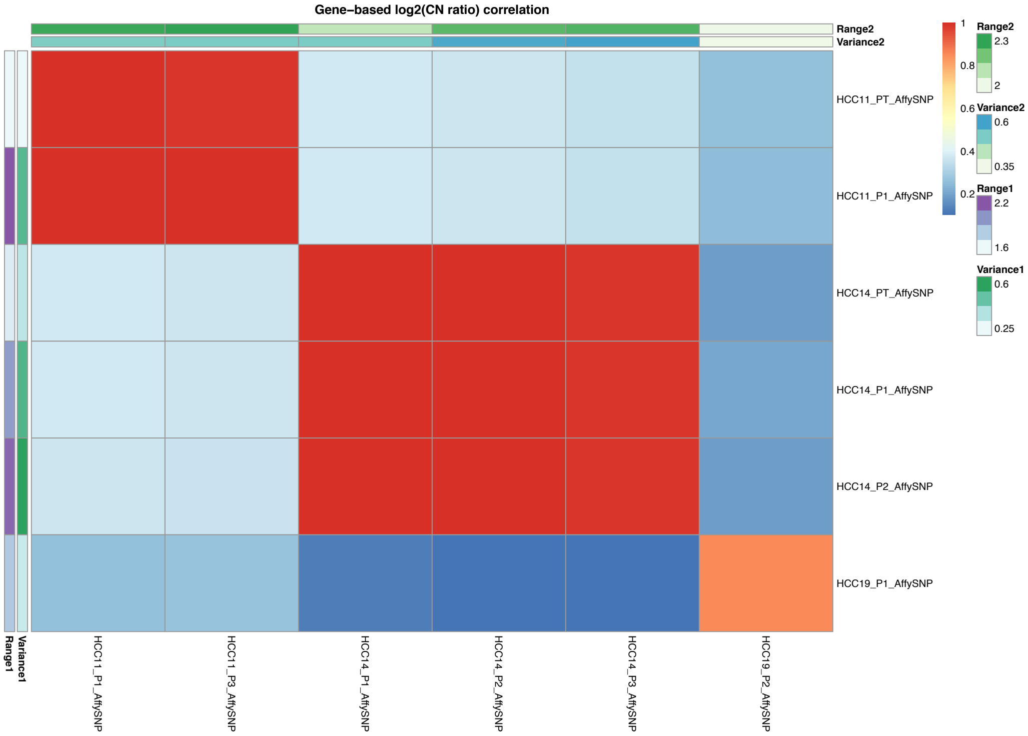

Supplementary Figure 39: CNA profiles (IGV heatmap) and correlation heatmap of gene-based copy number (log2(CN ratio), median centered) of samples from SIBS SNP array hepatocellular carcinoma (HCC) dataset.

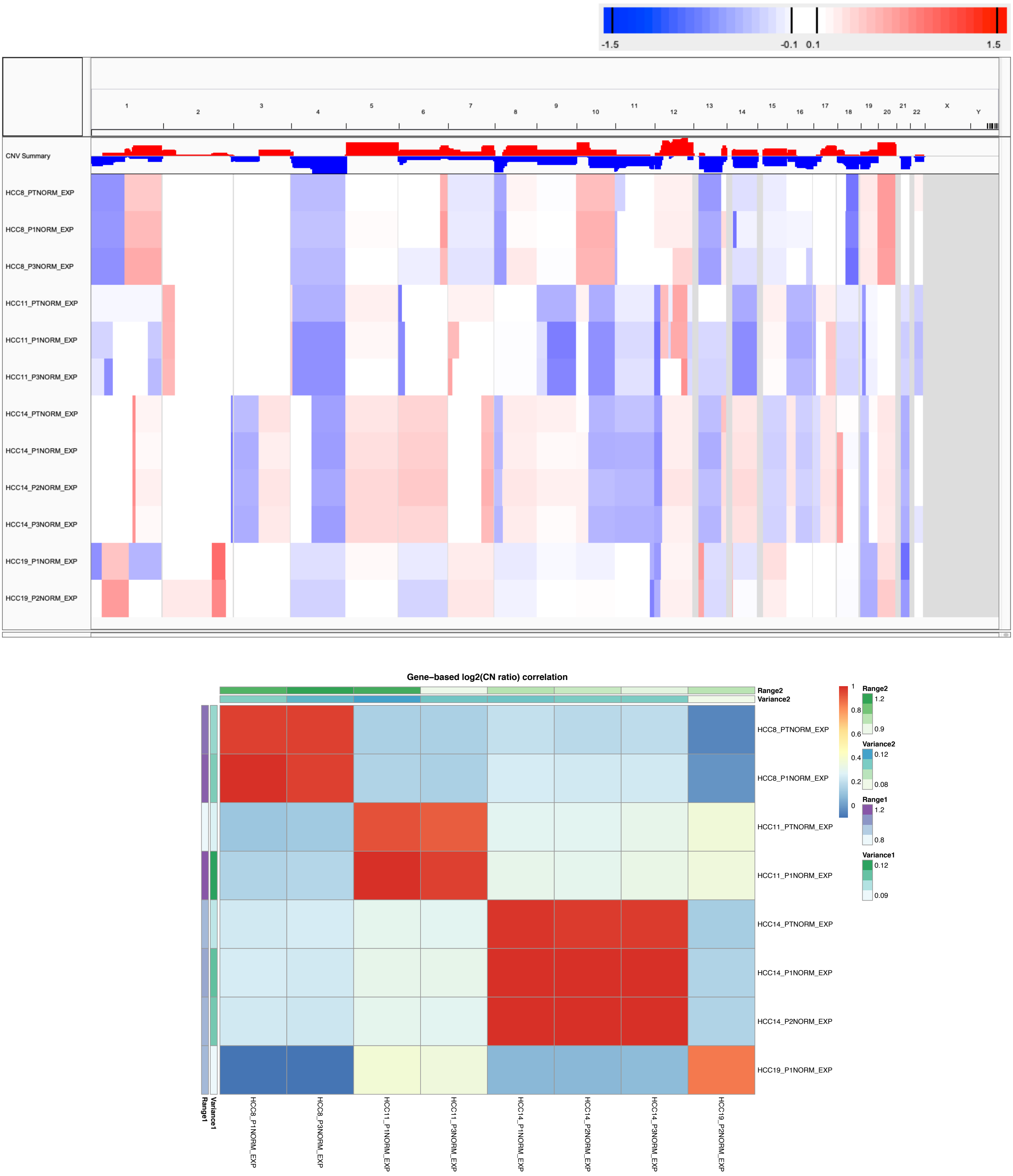

Supplementary Figure 40: CNA profiles (IGV heatmap) and correlation heatmap of gene-based copy number (log2(CN ratio), median centered) of samples from SIBS gene expression array (normalized by median expression of normal liver tissue samples) hepatocellular carcinoma (HCC) dataset.

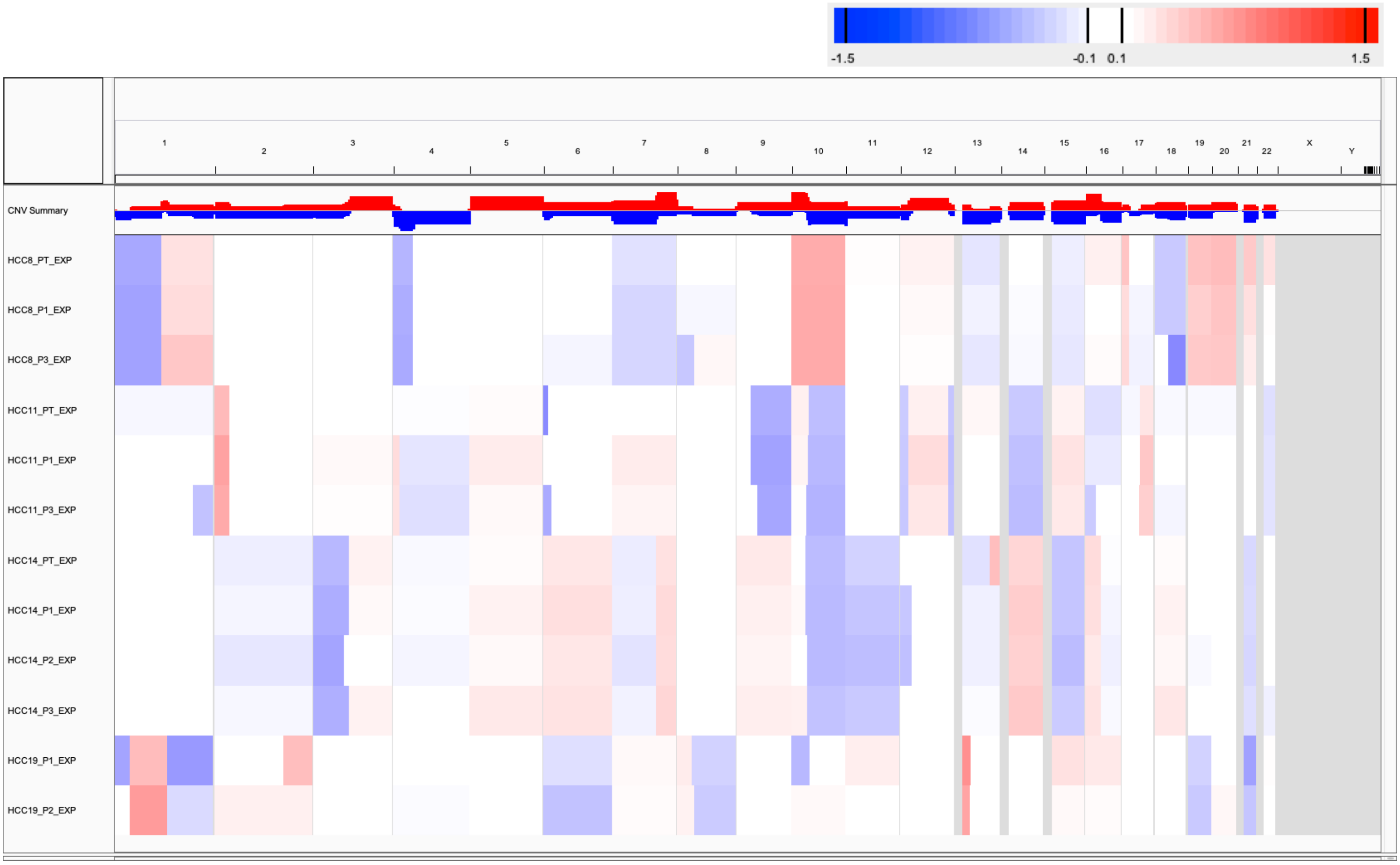

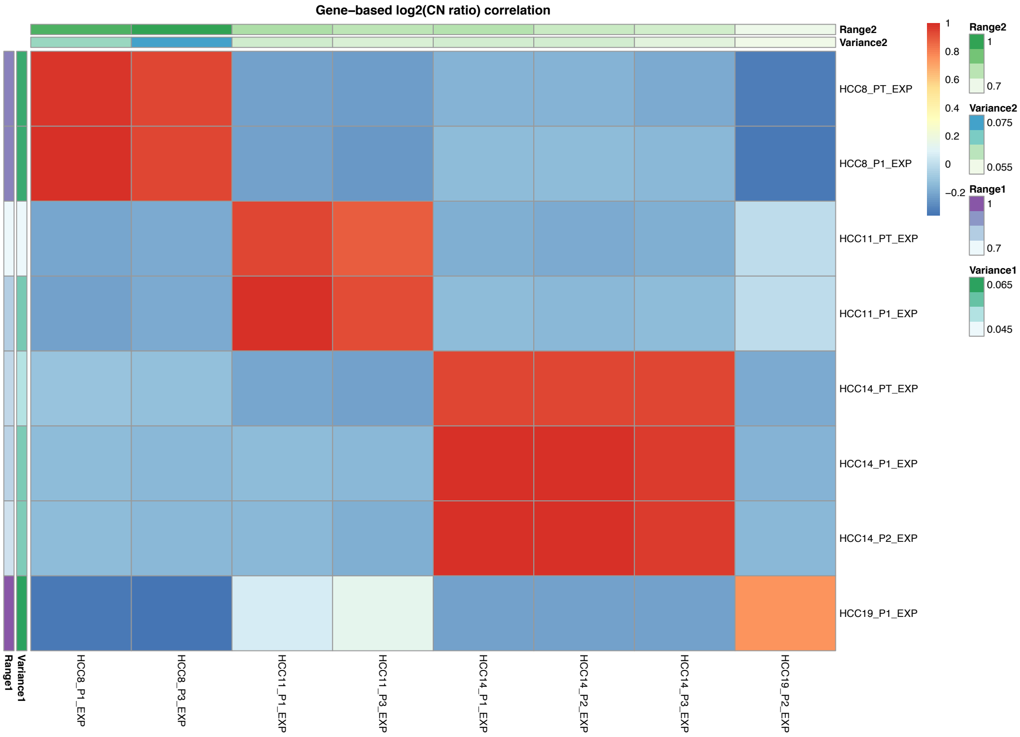

Supplementary Figure 41: CNA profiles (IGV heatmap) and correlation heatmap of gene-based copy number (log2(CN ratio), median centered) of samples from SIBS gene expression array (normalized by median expression of tumor samples of the same dataset) hepatocellular carcinoma (HCC) dataset.

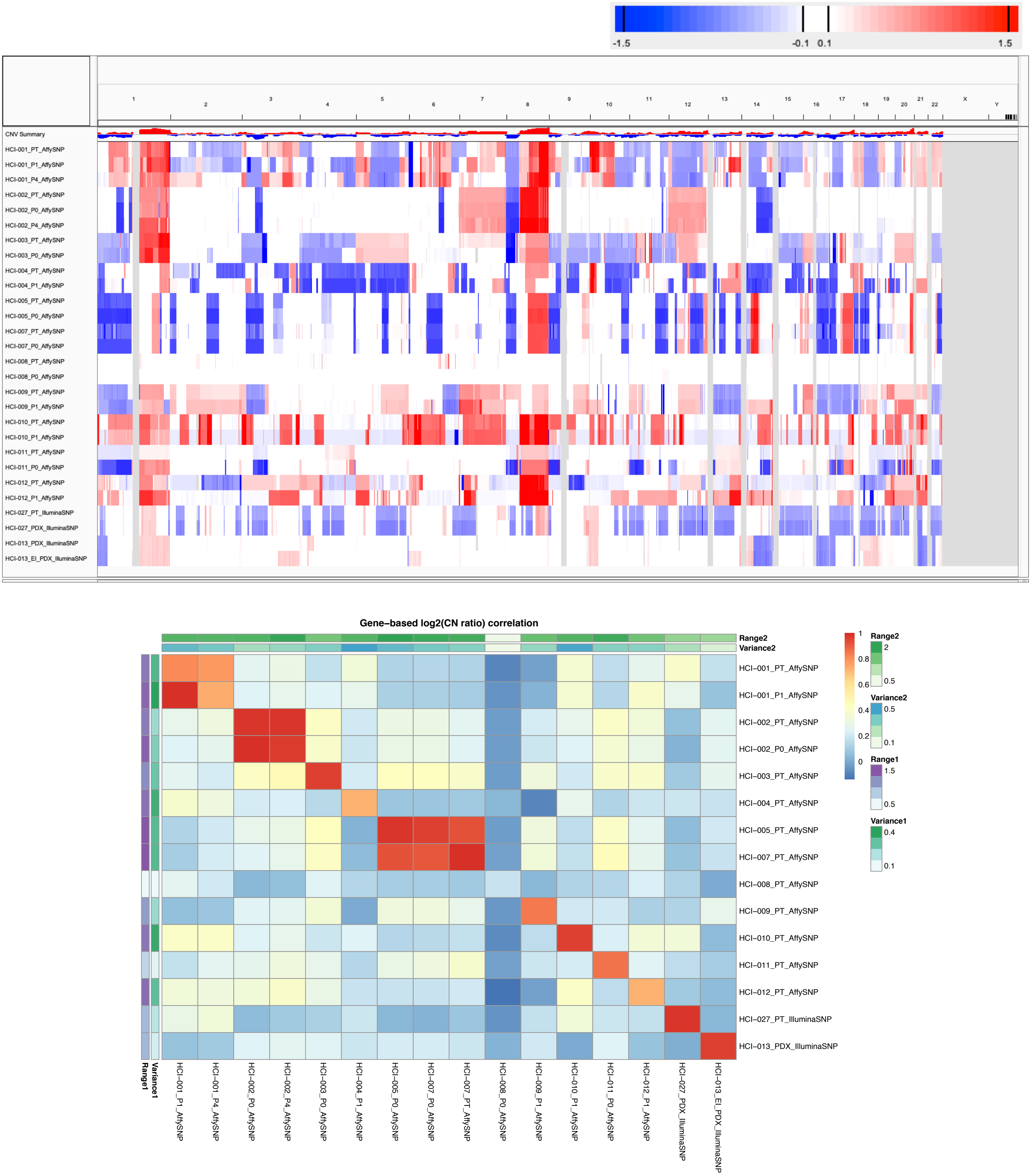

Supplementary Figure 42: CNA profiles (IGV heatmap) and correlation heatmap of gene-based copy number (log2(CN ratio), median centered) of samples from HCI SNP array breast cancer dataset.

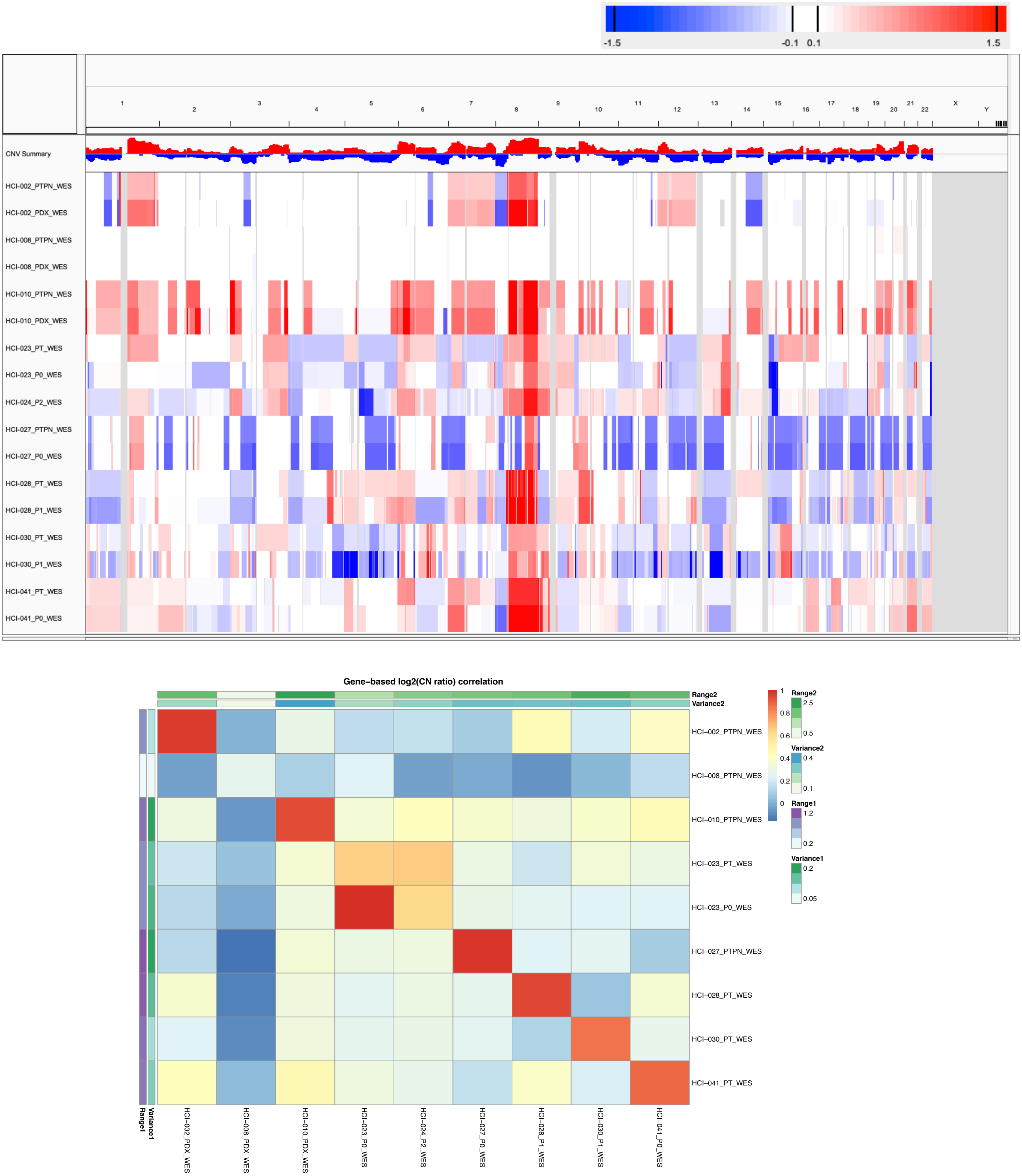

Supplementary Figure 43: CNA profiles (IGV heatmap) and correlation heatmap of gene-based copy number (log2(CN ratio), median centered) of samples from HCI WES breast cancer dataset.

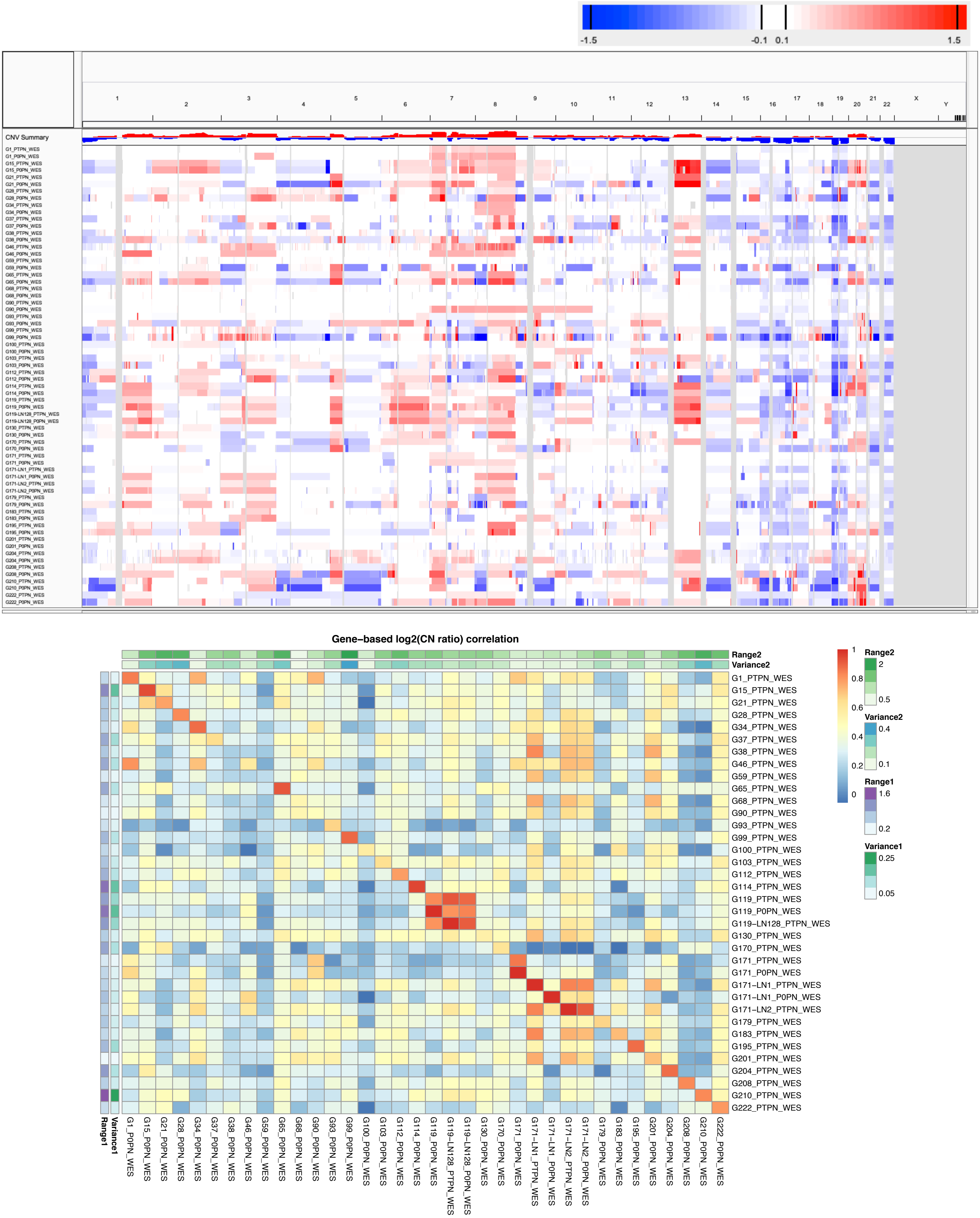

Supplementary Figure 44: CNA profiles (IGV heatmap) and correlation heatmap of gene-based copy number (log2(CN ratio), median centered) of samples from SNU-JAX WES gastric cancer dataset.

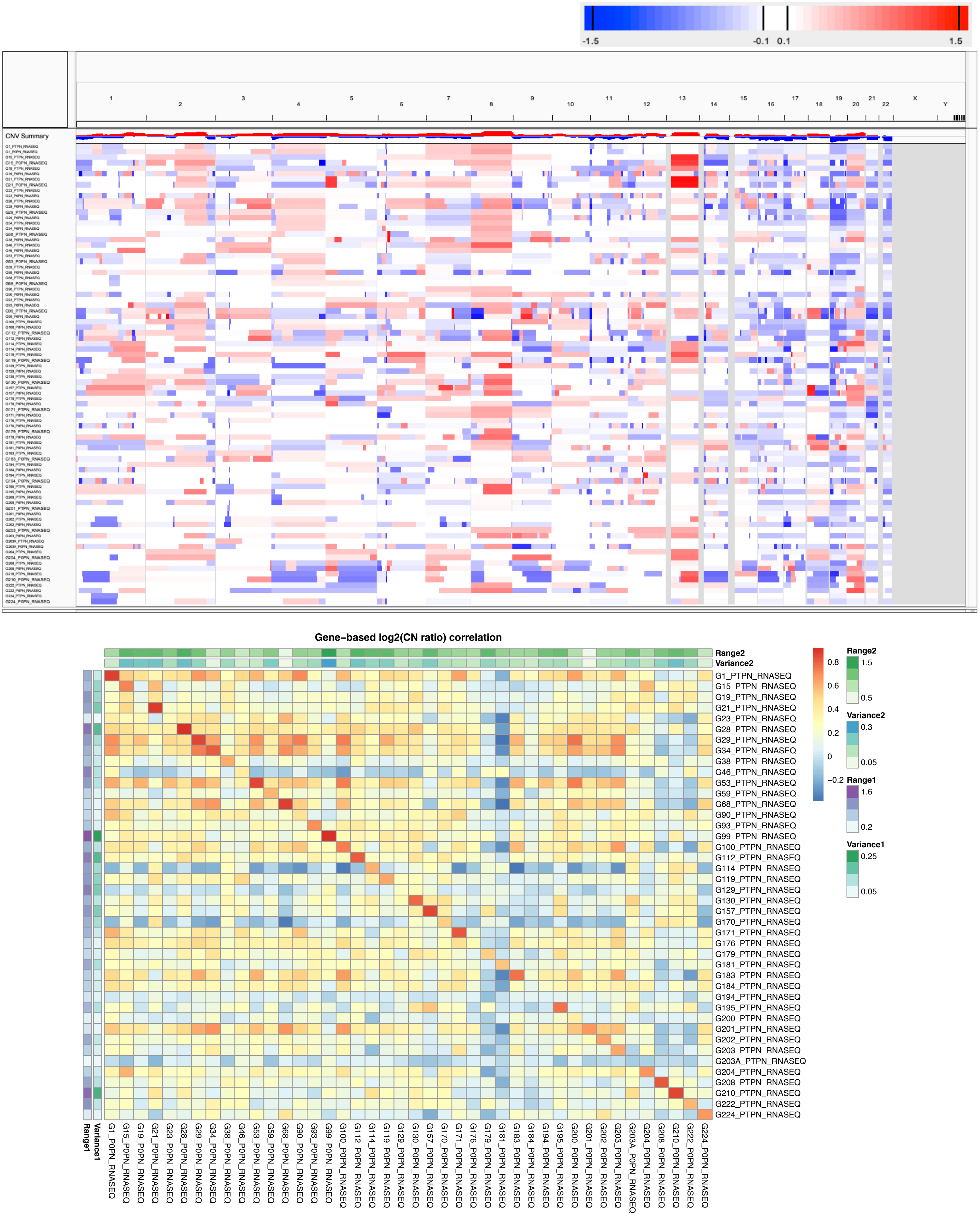

Supplementary Figure 45: CNA profiles (IGV heatmap) and correlation heatmap of gene-based copy number (log2(CN ratio), median centered) of samples from SNU-JAX RNA-Seq (normalized by median expression of normal gastric tissue samples from the same patients) gastric cancer dataset.

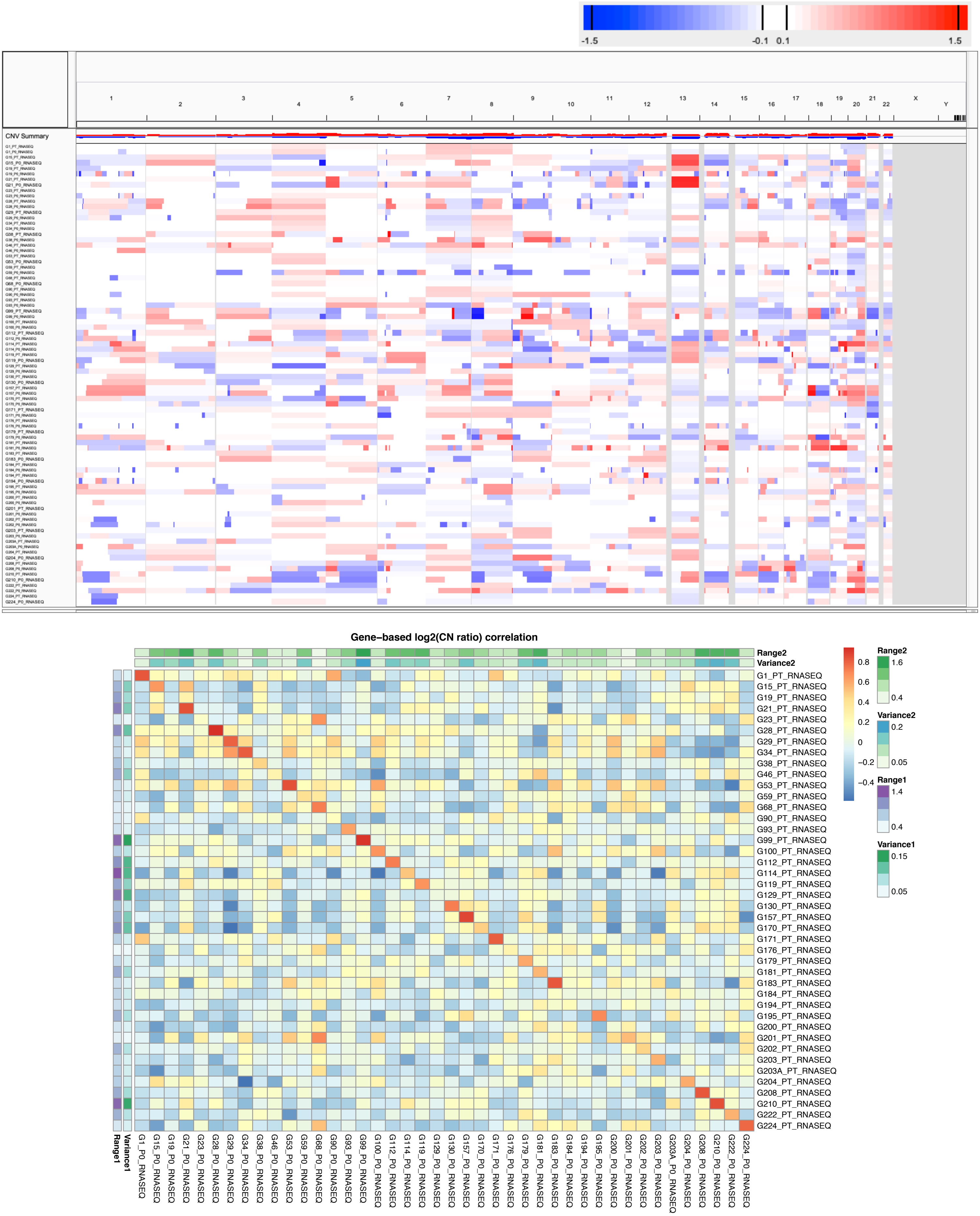

Supplementary Figure 46: CNA profiles (IGV heatmap) and correlation heatmap of gene-based copy number (log2(CN ratio), median centered) of samples from SNU-JAX RNA-Seq (normalized by median expression of tumor samples of the same dataset) gastric cancer dataset.

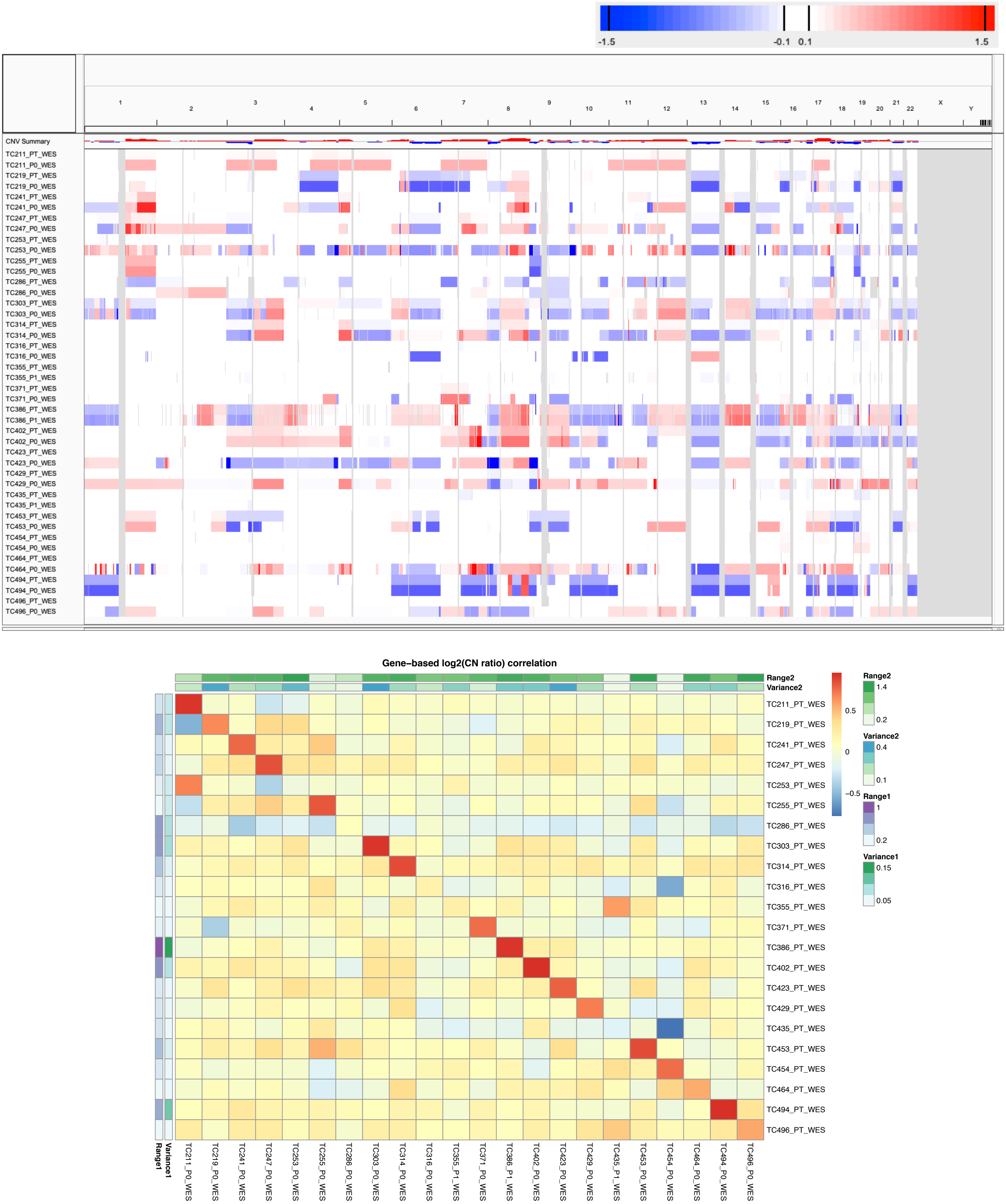

Supplementary Figure 47: CNA profiles (IGV heatmap) and correlation heatmap of gene-based copy number (log2(CN ratio), median centered) of samples from MDACC WES lung adenocarcinoma (LUAD) dataset.

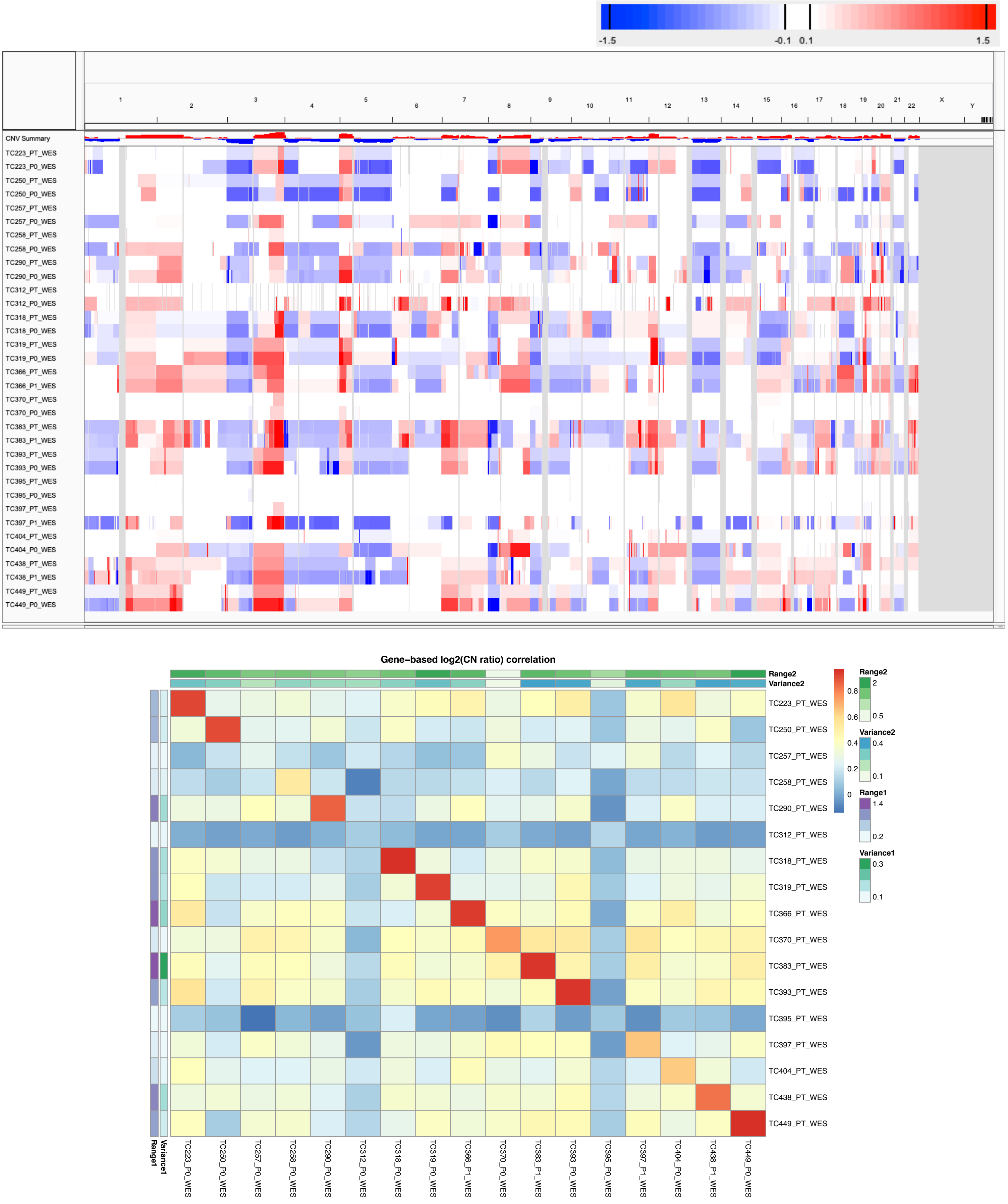

Supplementary Figure 48: CNA profiles (IGV heatmap) and correlation heatmap of gene-based copy number (log2(CN ratio), median centered) of samples from MDACC WES lung squamous cell carcinoma (LUSC) dataset.

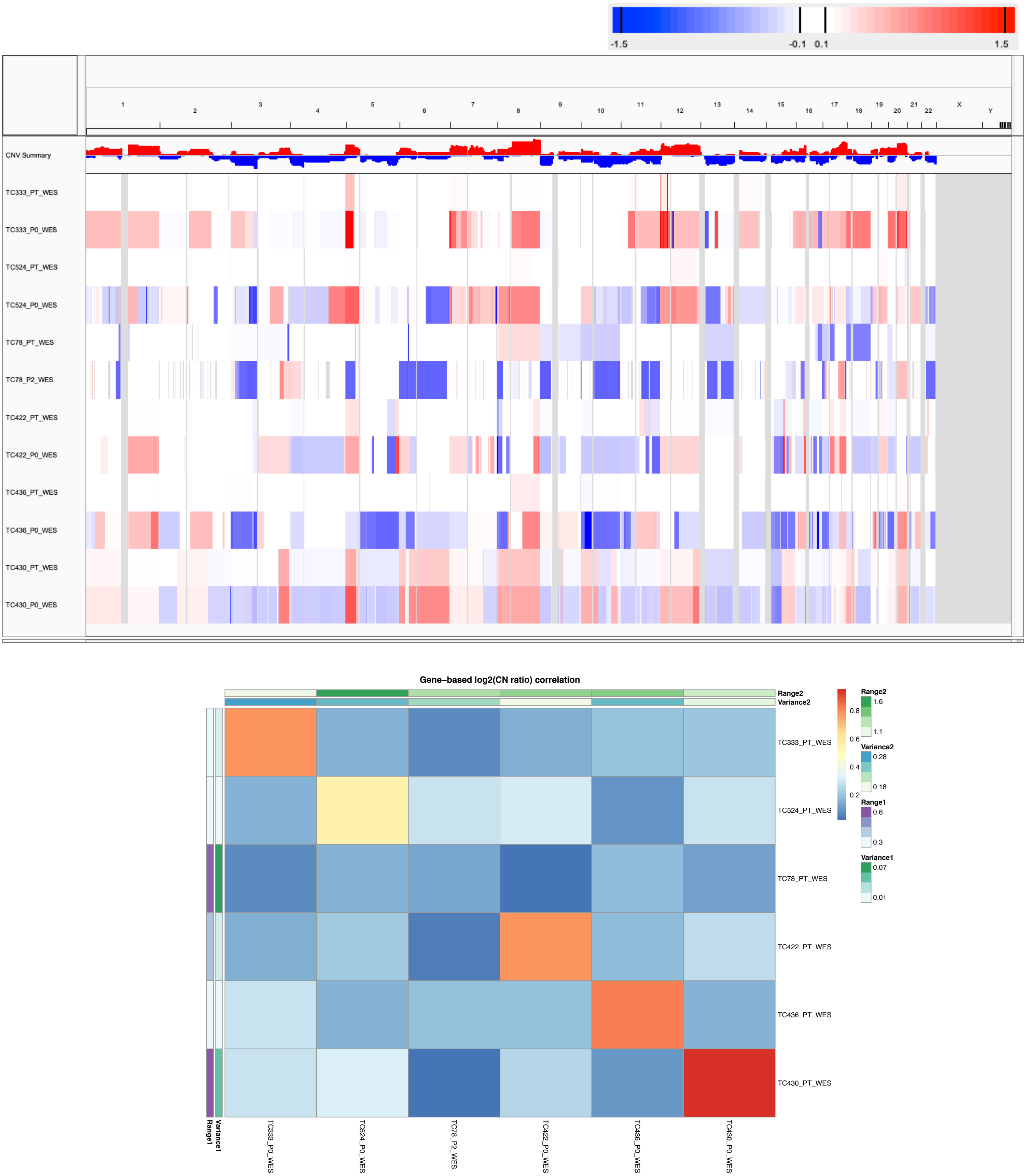

Supplementary Figure 49: CNA profiles (IGV heatmap) and correlation heatmap of gene-based copy number (log2(CN ratio), median centered) of samples from MDACC WES other lung cancer subtypes dataset.

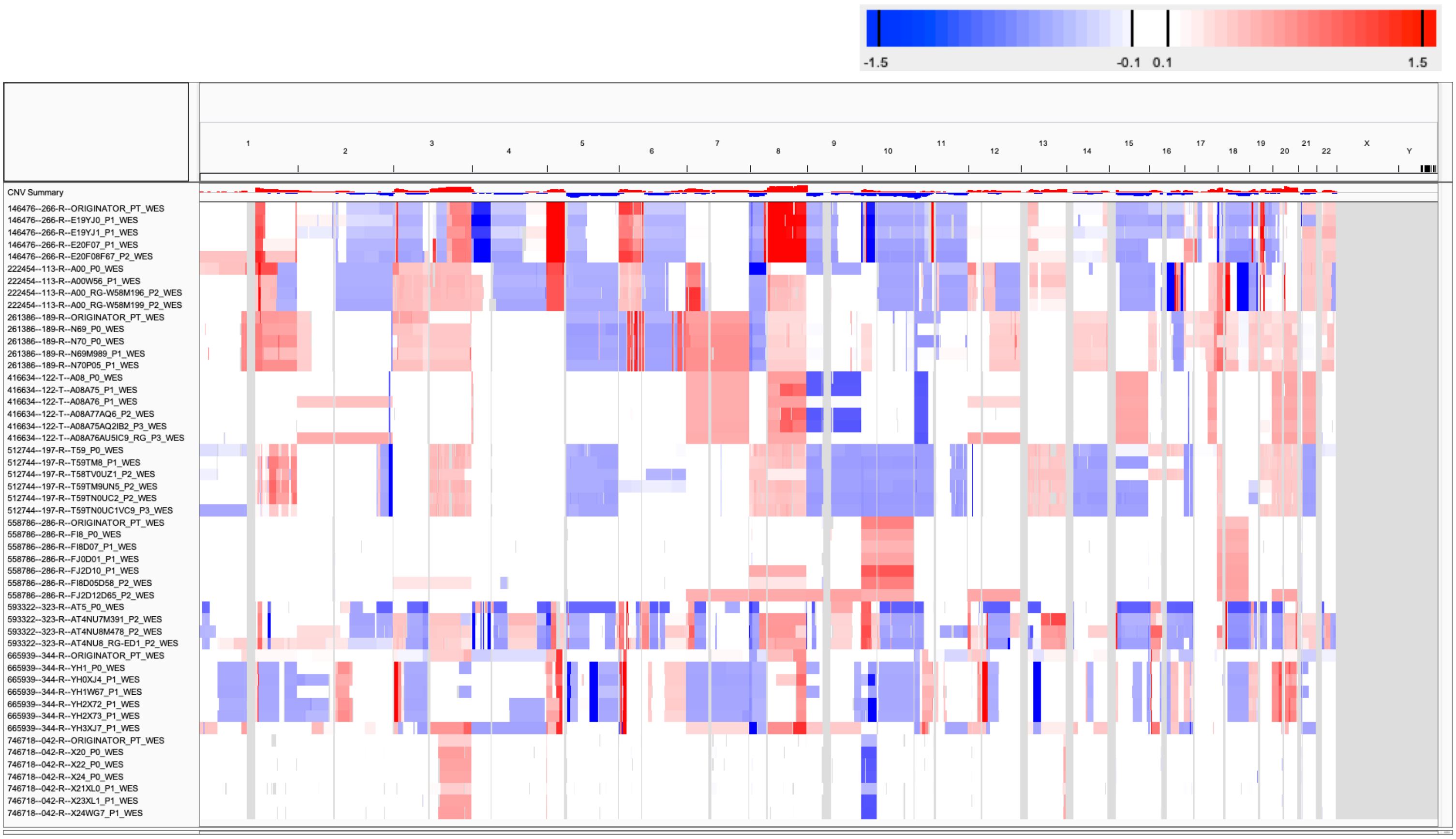

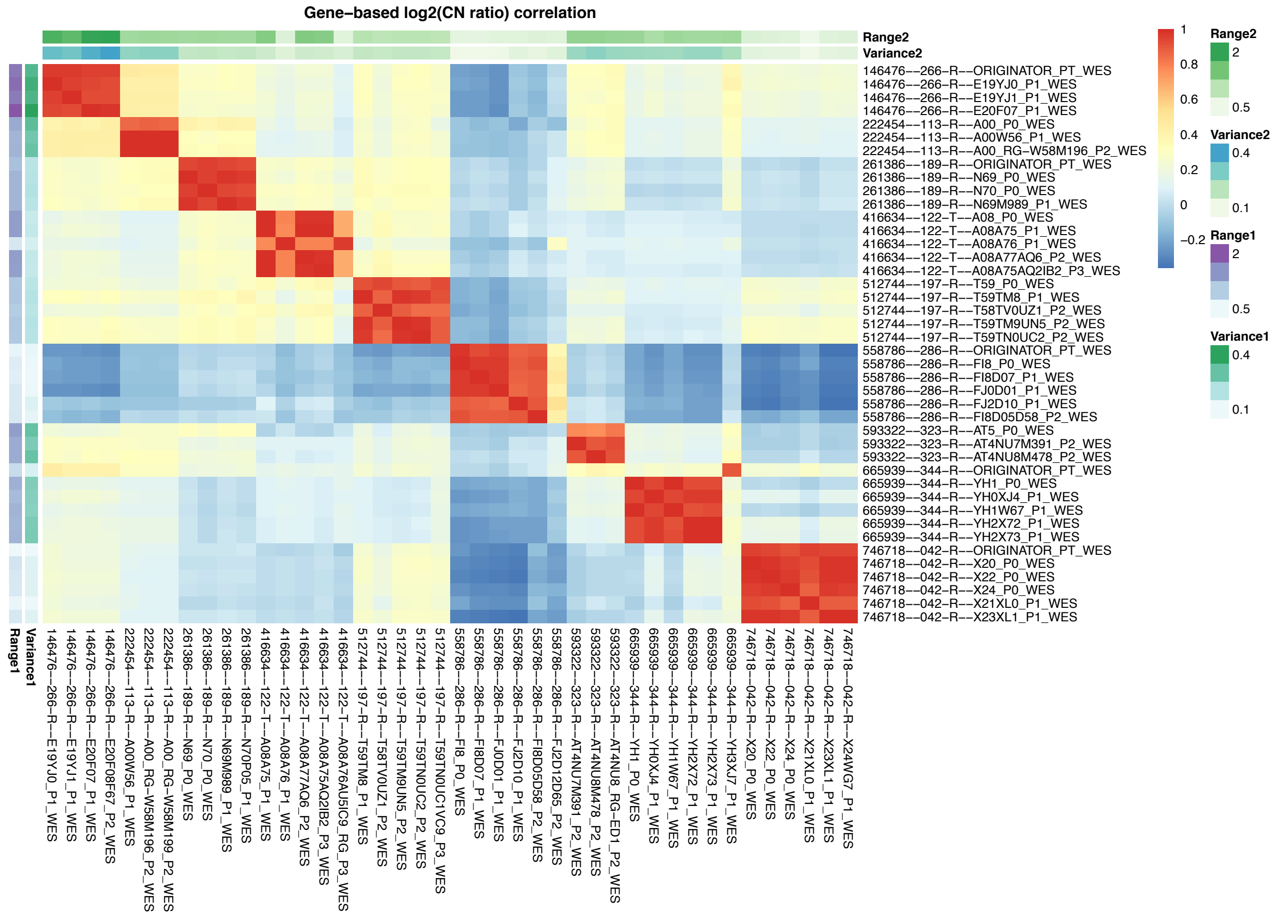

Supplementary Figure 50: CNA profiles (IGV heatmap) and correlation heatmap of gene-based copy number (log2(CN ratio), median centered) of samples from PDMR WES bladder cancer dataset.

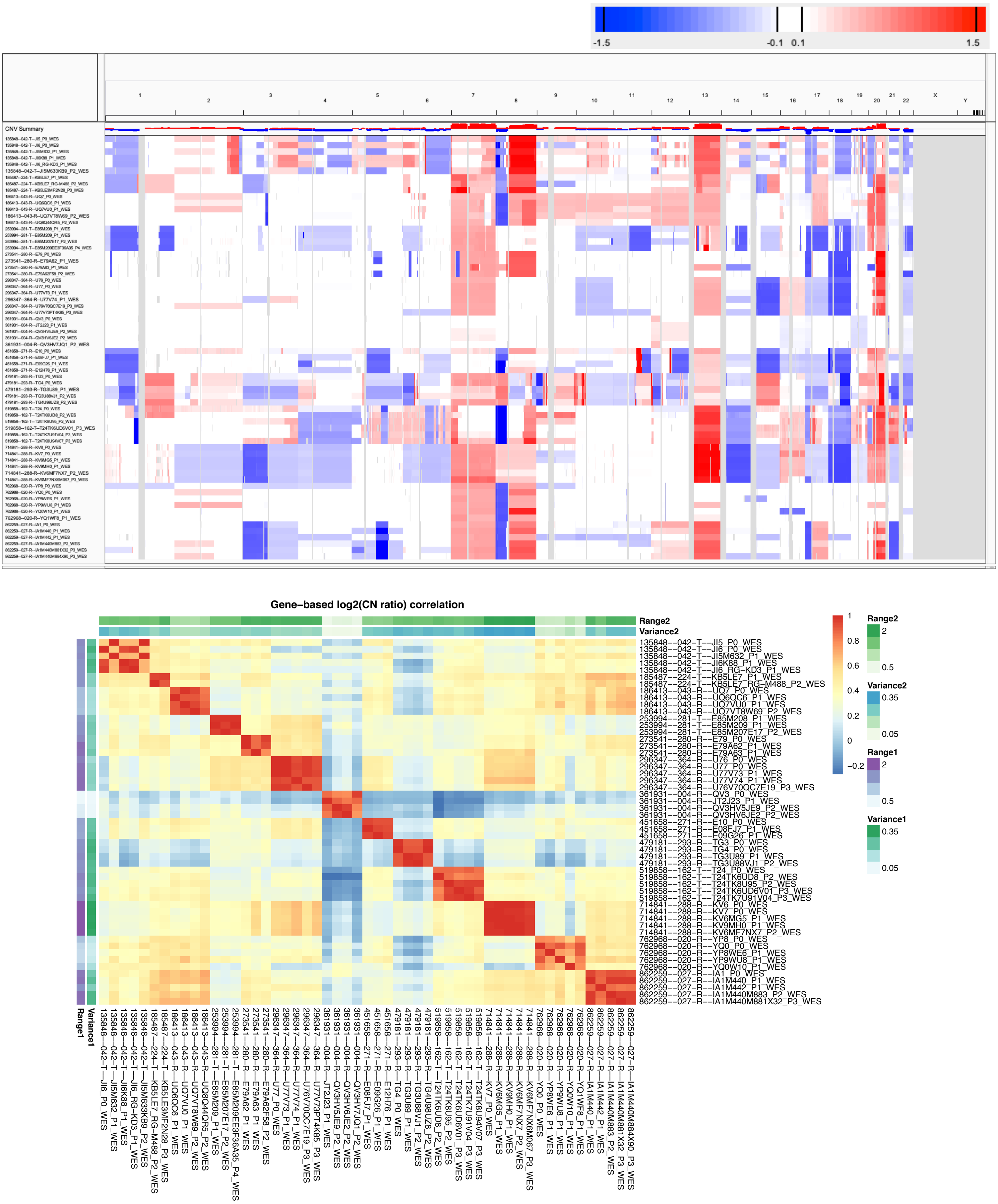

Supplementary Figure 51: CNA profiles (IGV heatmap) and correlation heatmap of gene-based copy number (log2(CN ratio), median centered) of samples from PDMR WES colorectal cancer dataset.

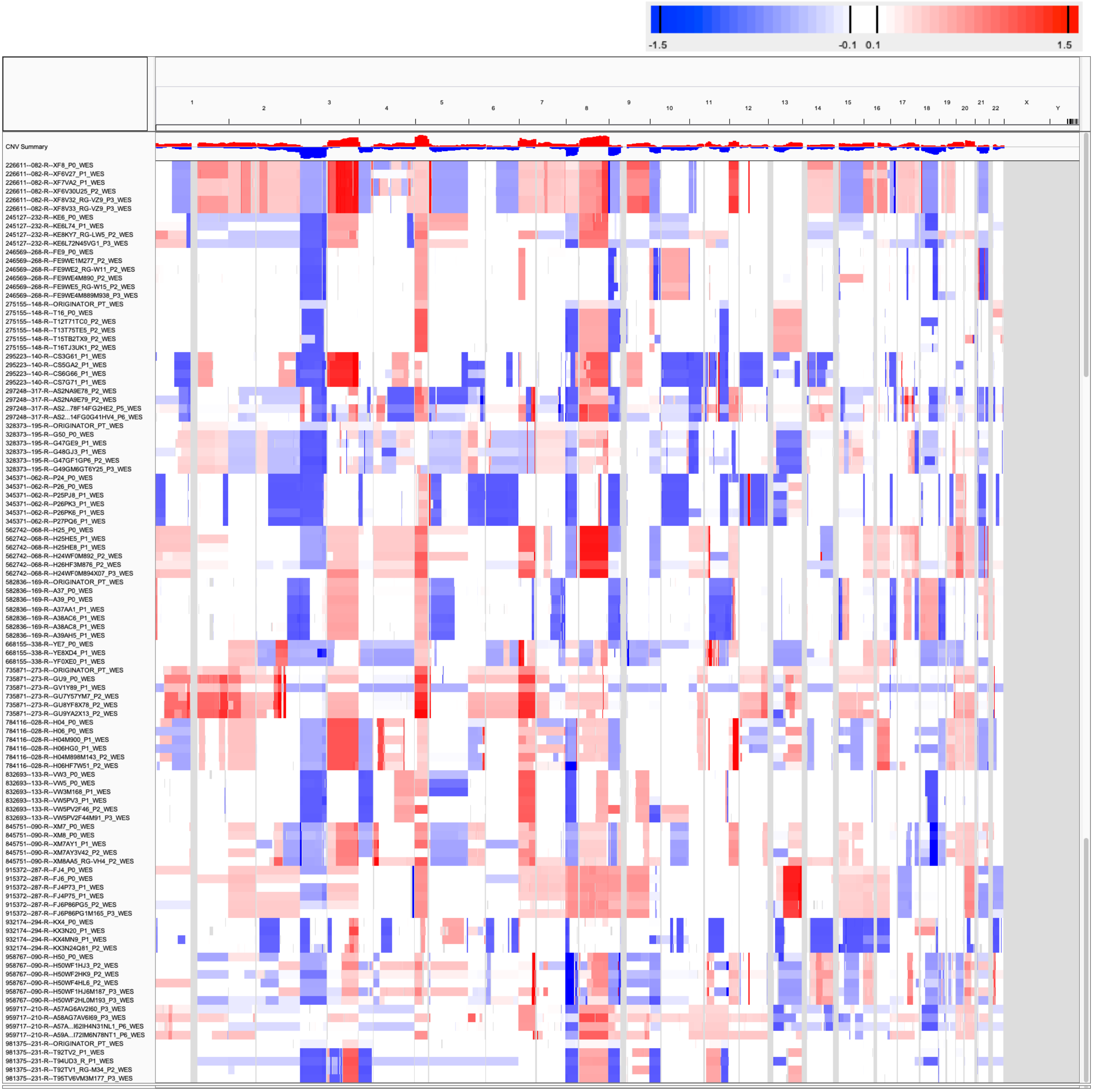
-1.5
-0.1
0.1
1.5
X
Y
CNV Summary
226611--082-R--XF8_P0_WES
226611--082-R--XF6V27_P1_WES
226611--082-R--XF7VA2_P1_WES
226611--082-R--XF6V30U25_P2_WES
226611--082-R--XF8V32_RG-VZ9_P3_WES
226611--082-R--XF8V33_RG-VZ9_P3_WES
245127--232-R--KE6_P0_WES
245127--232-R--KE6L74_P1_WES
245127--232-R--KE8KY7_RG-LW5_P2_WES
245127--232-R--KE6L72N45VG1_P3_WES
246569--268-R--FE9_P0_WES
246569--268-R--FE9WE1M277_P2_WES
246569--268-R--FE9WE2_RG-W11_P2_WES
246569--268-R--FE9WE4M890_P2_WES
246569--268-R--FE9WE5_RG-W15_P2_WES
246569--268-R--FE9WE4M889M938_P3_WES
275155--148-R--ORIGINATOR_PT_WES
275155--148-R--T16_P0_WES
275155--148-R--T12T71TC0_P2_WES
275155--148-R--T13T75TE5_P2_WES
275155--148-R--T15TB2TX9_P2_WES
275155--148-R--T16TJ3UK1_P2_WES
295223--140-R--CS3G61_P1_WES
295223--140-R--CS5GA2_P1_WES
295223--140-R--CS6G66_P1_WES
295223--140-R--CS7G71_P1_WES
297248--317-R--AS2NA9E78_P2_WES
297248--317-R--AS2NA9E79_P2_WES
297248--317-R--AS2...78F14FG2HE2_P5_WES
297248--317-R--AS2...14FG0G41HV4_P6_WES
328373--195-R--ORIGINATOR_PT_WES
328373--195-R--G50_P0_WES
328373--195-R--G47GE9_P1_WES
328373--195-R--G48GJ3_P1_WES
328373--195-R--G47GF1GP6_P2_WES
328373--195-R--G49GM6GT6Y25_P3_WES
345371--062-R--P24_P0_WES
345371--062-R--P26_P0_WES
345371--062-R--P25PJ8_P1_WES
345371--062-R--P26PK3_P1_WES
345371--062-R--P26PK6_P1_WES
345371--062-R--P27PQ6_P1_WES
562742--068-R--H25_P0_WES
562742--068-R--H25HE5_P1_WES
562742--068-R--H25HE8_P1_WES
562742--068-R--H24WF0M892_P2_WES
562742--068-R--H26HF3M876_P2_WES
562742--068-R--H24WF0M894X07_P3_WES
582836--169-R--ORIGINATOR_PT_WES
582836--169-R--A37_P0_WES
582836--169-R--A39_P0_WES
582836--169-R--A37AA1_P1_WES
582836--169-R--A38AC6_P1_WES
582836--169-R--A38AC8_P1_WES
582836--169-R--A39AH5_P1_WES
668155--338-R--YE7_P0_WES
668155--338-R--YE8XD4_P1_WES
668155--338-R--YF0XE0_P1_WES
735871--273-R--ORIGINATOR_PT_WES
735871--273-R--GU9_P0_WES
735871--273-R--GV1Y89_P1_WES
735871--273-R--GU7Y57YM7_P2_WES
735871--273-R--GU8YF8X78_P2_WES
735871--273-R--GU9YA2X13_P2_WES
784116--028-R--H04_P0_WES
784116--028-R--H06_P0_WES
784116--028-R--H04M900_P1_WES
784116--028-R--H06HG0_P1_WES
784116--028-R--H04M898M143_P2_WES
784116--028-R--H06HF7W51_P2_WES
832693--133-R--VW3_P0_WES
832693--133-R--VW5_P0_WES
832693--133-R--VW3M168_P1_WES
832693--133-R--VW5PV3_P1_WES
832693--133-R--VW5PV2F46_P2_WES
832693--133-R--VW5PV2F44M91_P3_WES
845751--090-R--XM7_P0_WES
845751--090-R--XM8_P0_WES
845751--090-R--XM7AY1_P1_WES
845751--090-R--XM7AY3V42_P2_WES
845751--090-R--XM8AA5_RG-VH4_P2_WES
915372--287-R--FJ4_P0_WES
915372--287-R--FJ6_P0_WES
915372--287-R--FJ4P73_P1_WES
915372--287-R--FJ4P75_P1_WES
915372--287-R--FJ6P86PG5_P2_WES
915372--287-R--FJ6P86PG1M165_P3_WES
932174--294-R--KX4_P0_WES
932174--294-R--KX3N20_P1_WES
932174--294-R--KX4MN9_P1_WES
932174--294-R--KX3N24Q81_P2_WES
958767--090-R--H50_P0_WES
958767--090-R--H50WF1HJ3_P2_WES
958767--090-R--H50WF2HK9_P2_WES
958767--090-R--H50WF4HL6_P2_WES
958767--090-R--H50WF1HJ6M187_P3_WES
958767--090-R--H50WF2HL0M193_P3_WES
959717--210-R--A57AG6AV2I60_P3_WES
959717--210-R--A58AG7AV6I69_P3_WES
959717--210-R--A57A...I62IH4N31NL1_P6_WES
959717--210-R--A59A...I72IM6N78NT1_P6_WES
981375--231-R--ORIGINATOR_PT_WES
981375--231-R--T92TV2_P1_WES
981375--231-R--T94UD3_R_P1_WES
981375--231-R--T92TV1_RG-M34_P2_WES
981375--231-R--T95TV6VM3M177_P3_WES

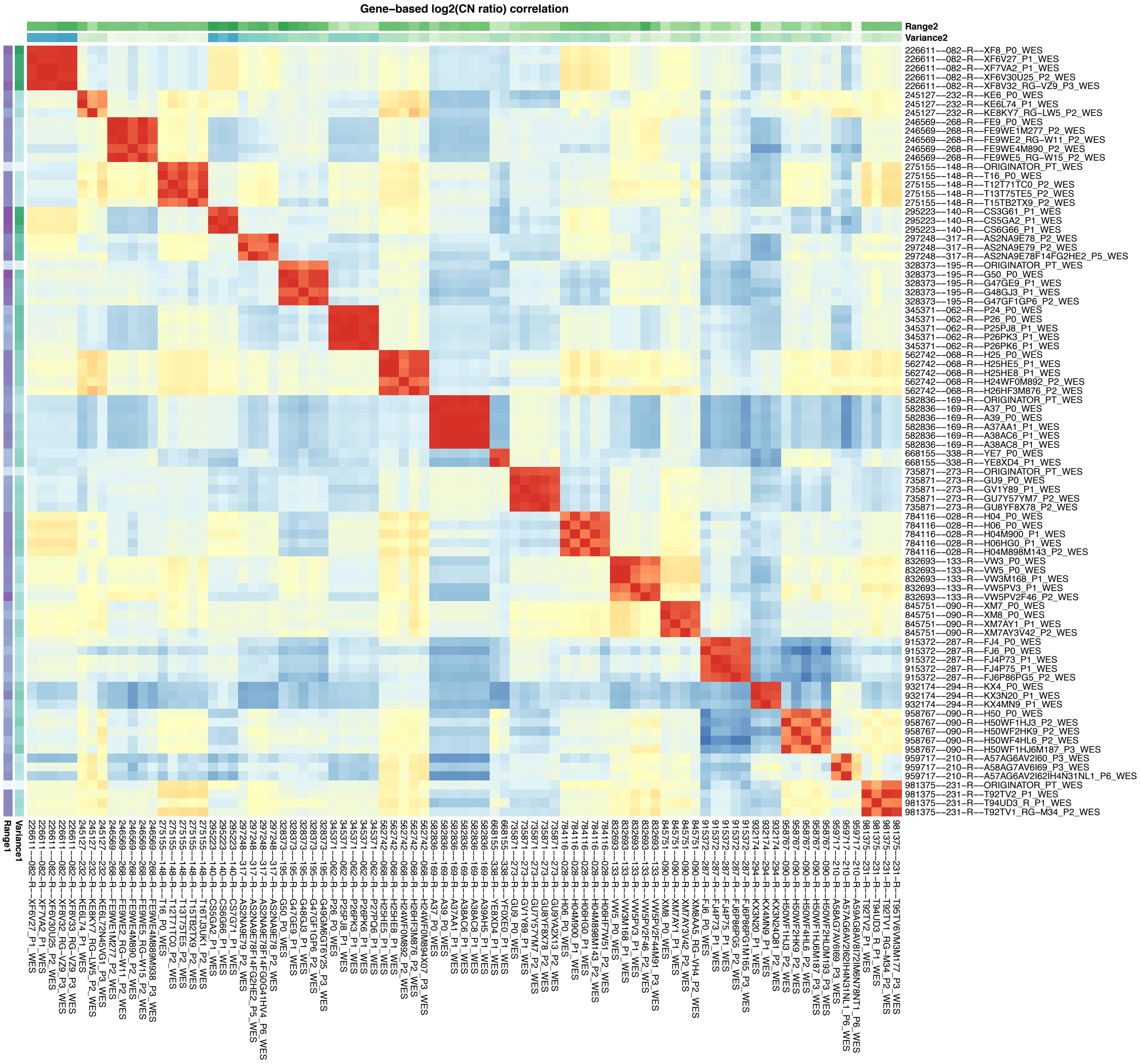

Supplementary Figure 52: CNA profiles (IGV heatmap) and correlation heatmap of gene-based copy number (log2(CN ratio), median centered) of samples from PDMR WES head and neck cancer dataset.

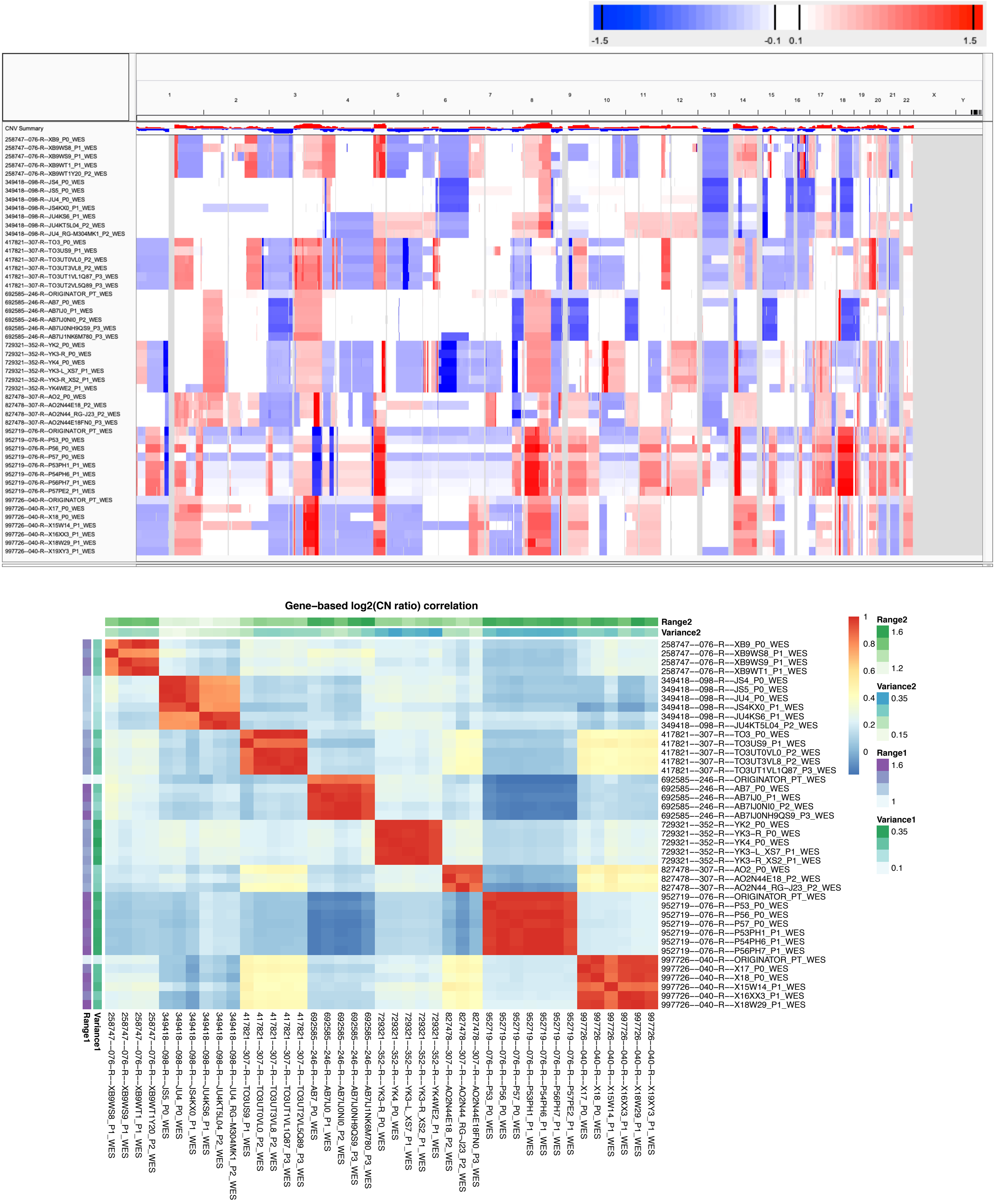

Supplementary Figure 53: CNA profiles (IGV heatmap) and correlation heatmap of gene-based copy number (log2(CN ratio), median centered) of samples from PDMR WES lung cancer dataset.

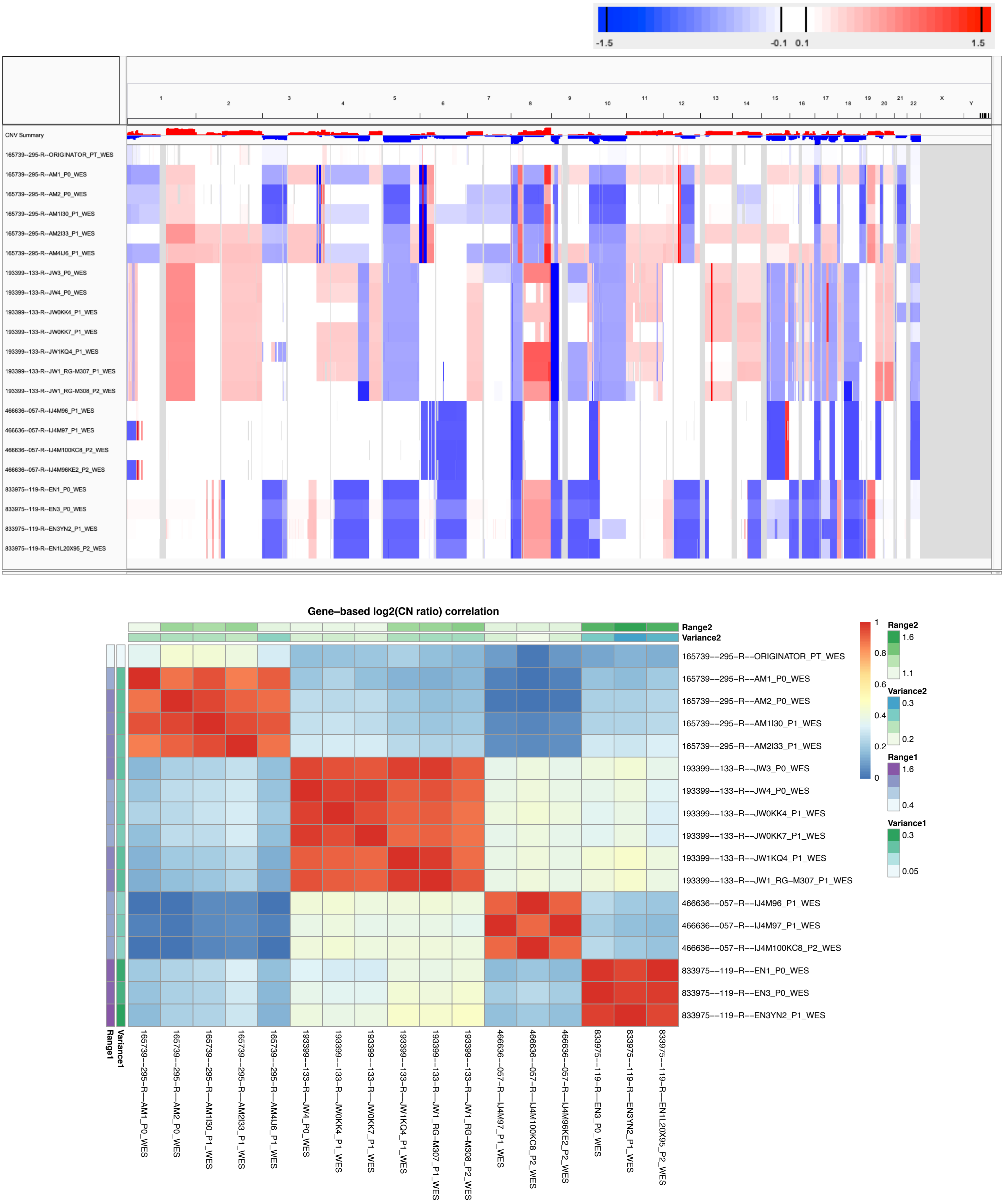

Supplementary Figure 54: CNA profiles (IGV heatmap) and correlation heatmap of gene-based copy number (log2(CN ratio), median centered) of samples from PDMR WES pancreatic cancer dataset.

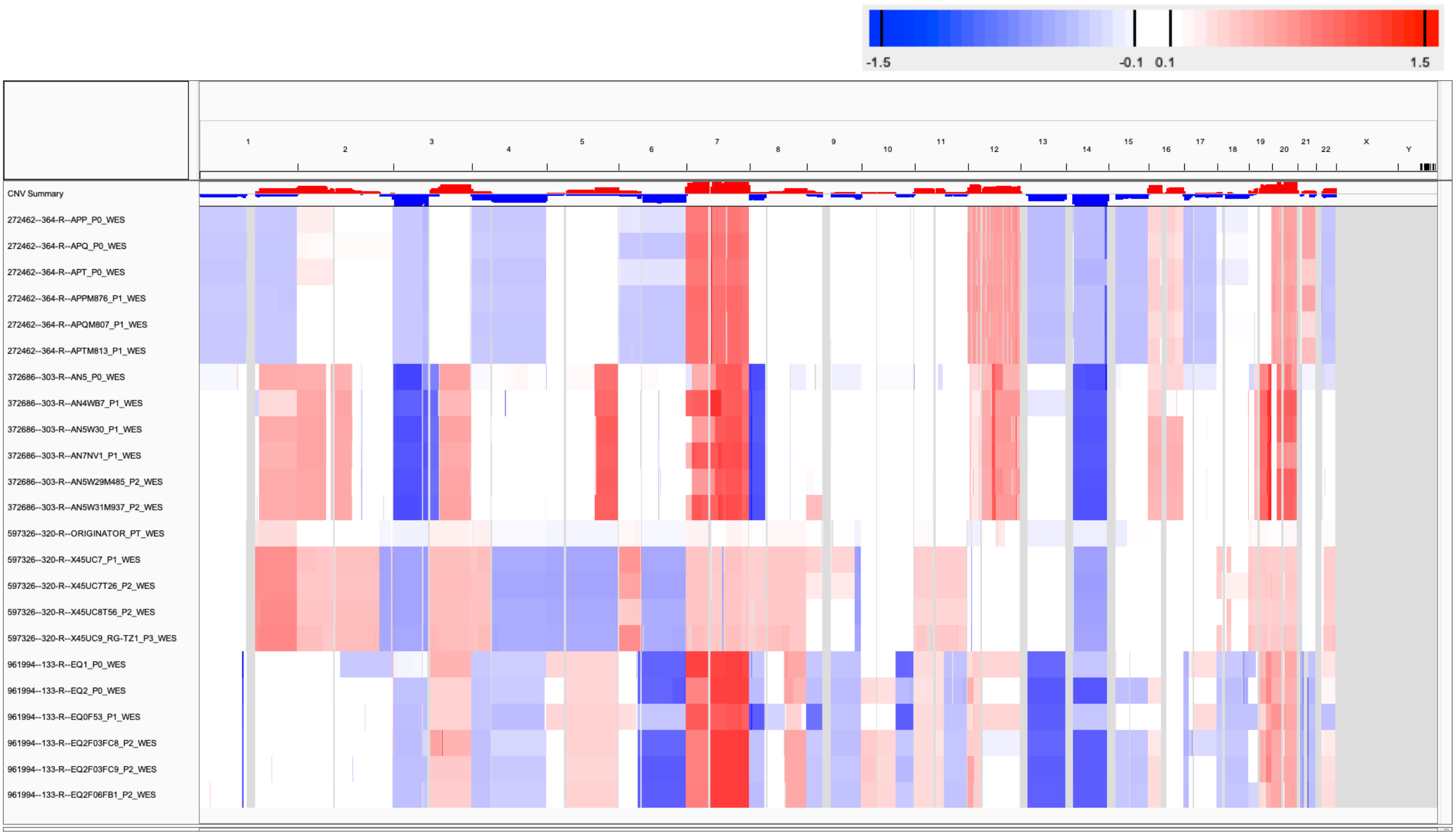

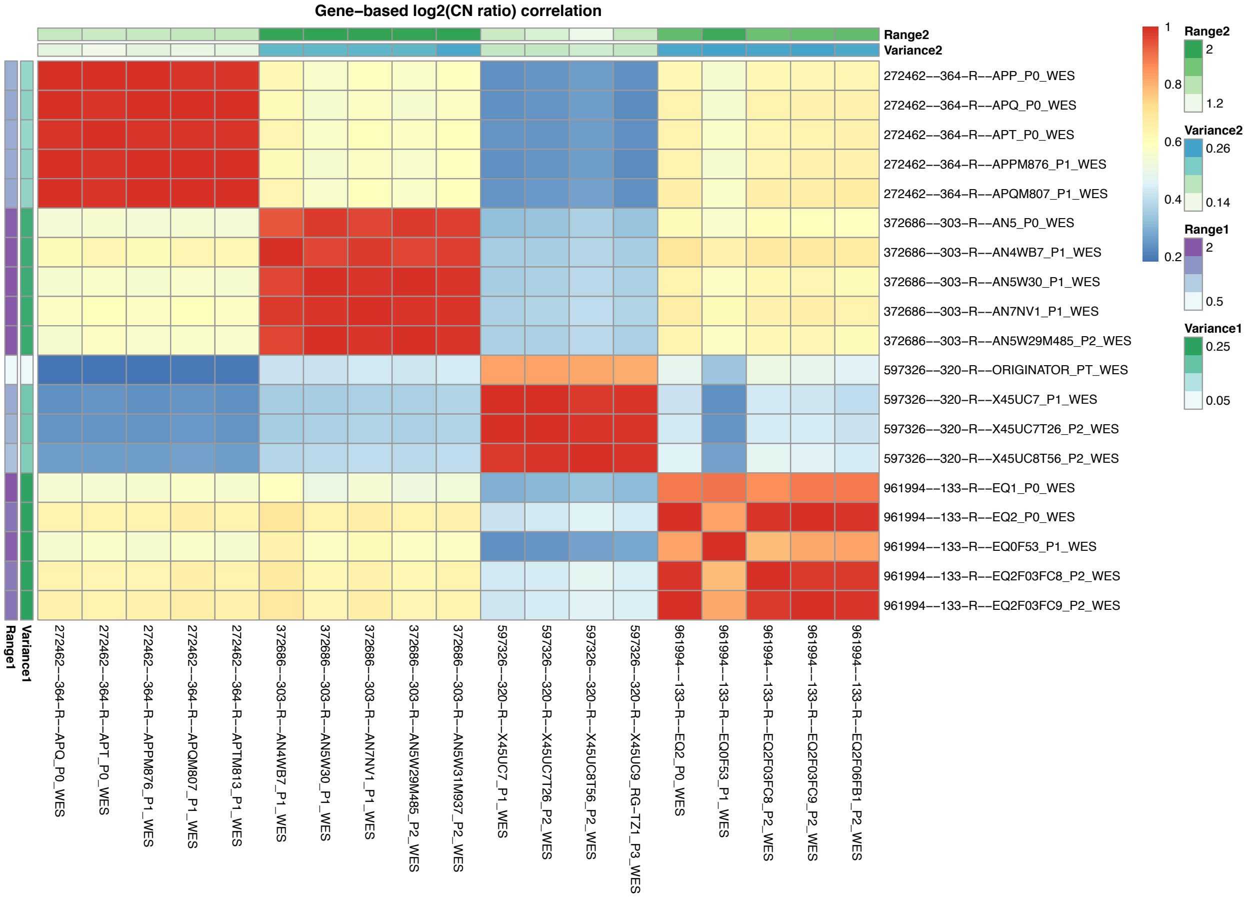

Supplementary Figure 55: CNA profiles (IGV heatmap) and correlation heatmap of gene-based copy number (log2(CN ratio), median centered) of samples from PDMR WES renal cancer dataset.

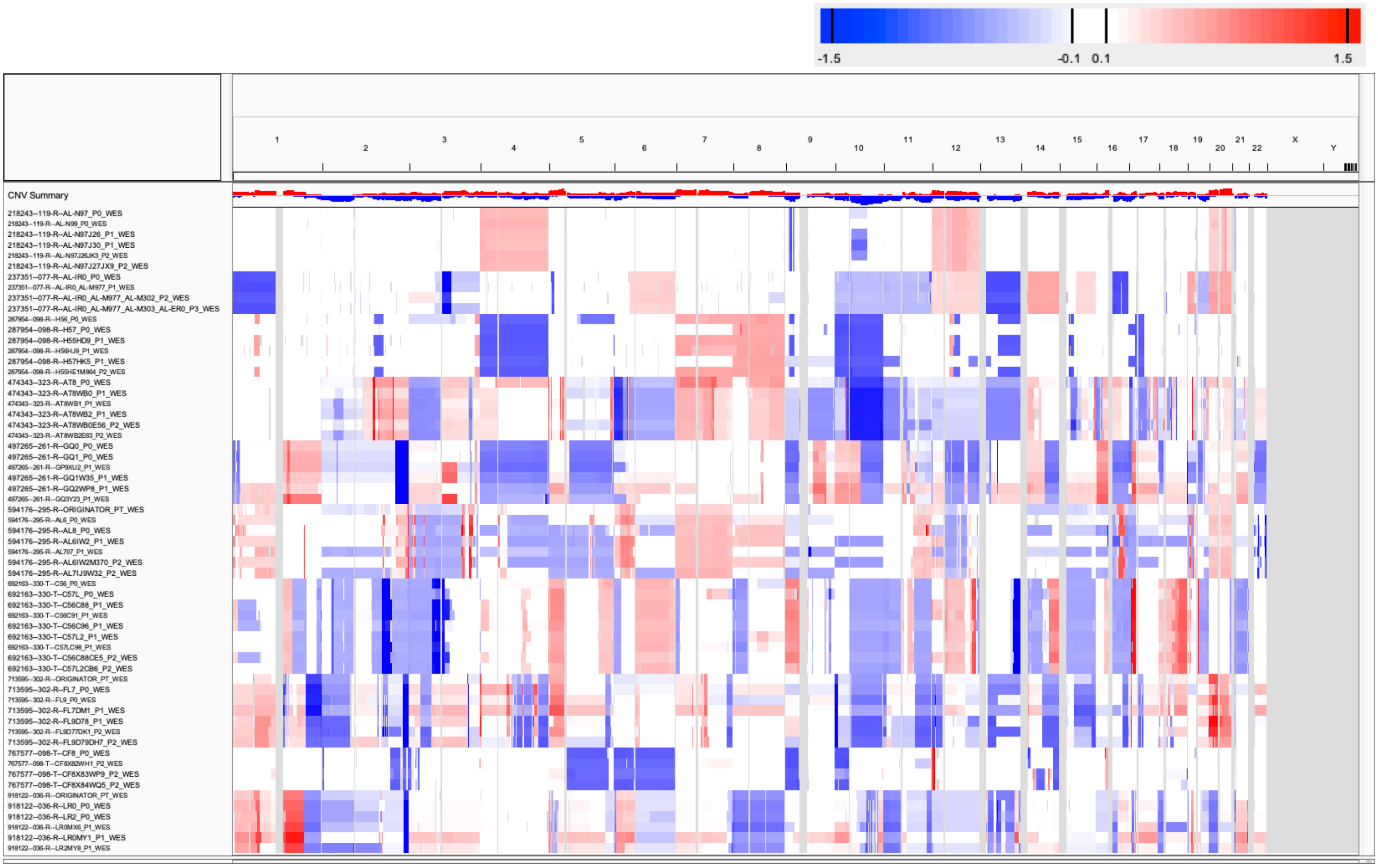

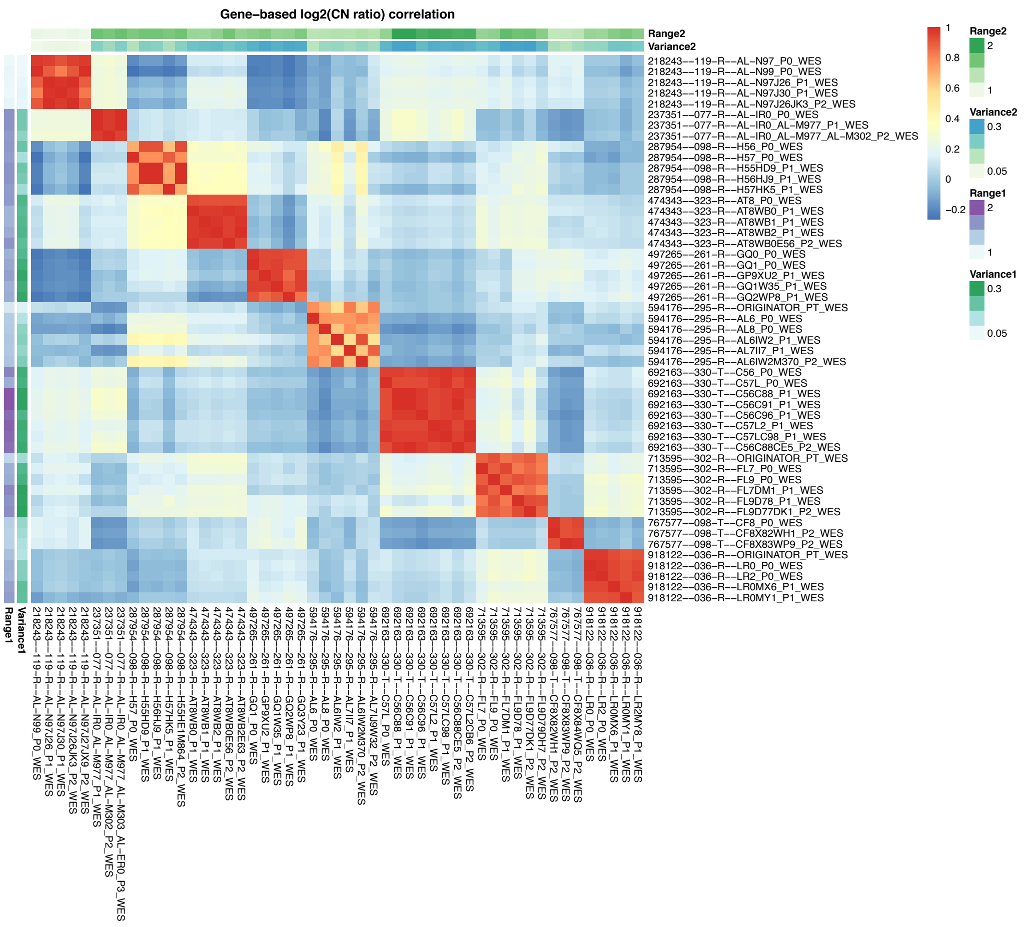

Supplementary Figure 56: CNA profiles (IGV heatmap) and correlation heatmap of gene-based copy number (log2(CN ratio), median centered) of samples from PDMR WES sarcoma dataset.

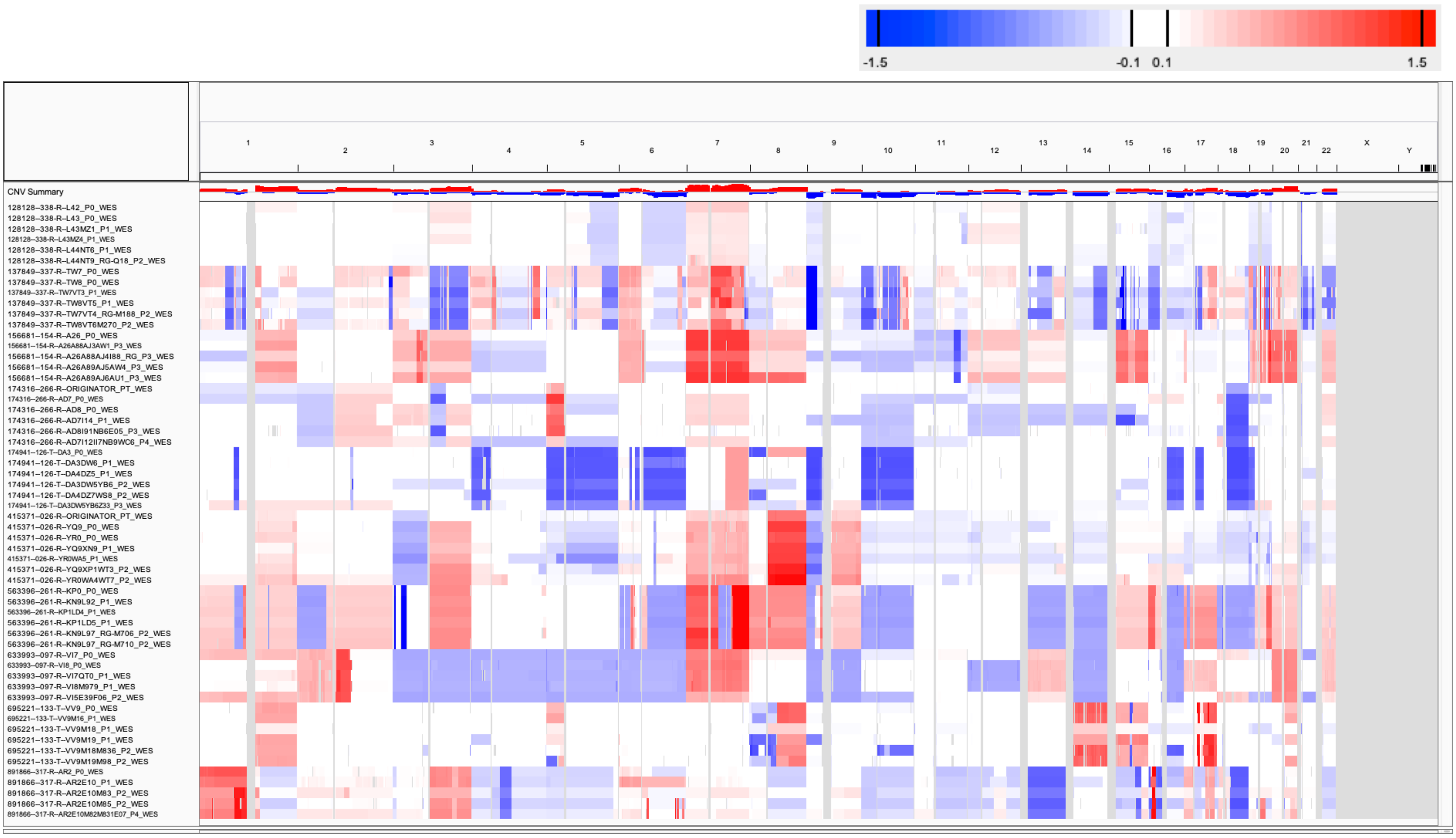

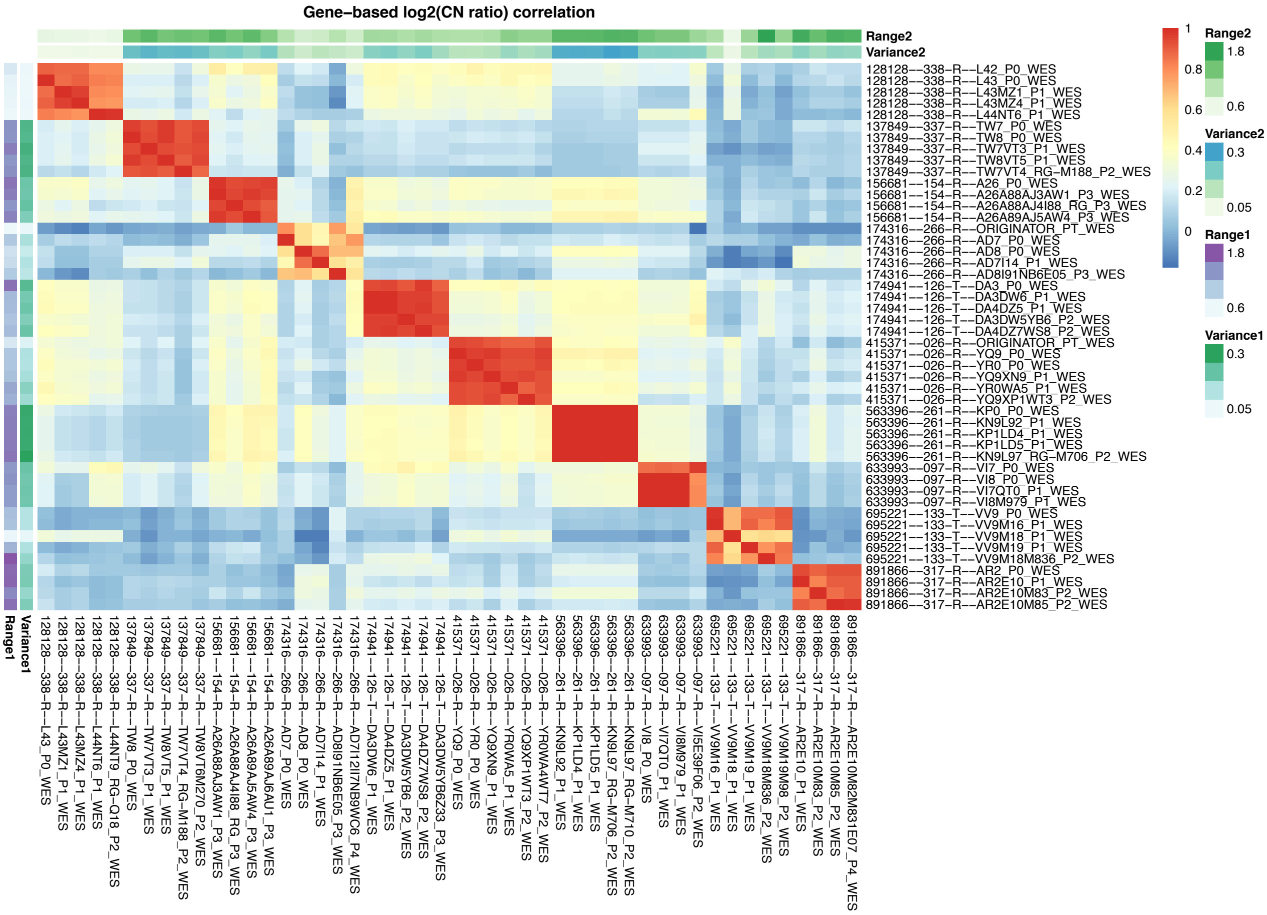

Supplementary Figure 57: CNA profiles (IGV heatmap) and correlation heatmap of gene-based copy number (log2(CN ratio), median centered) of samples from PDMR WES skin cancer dataset.

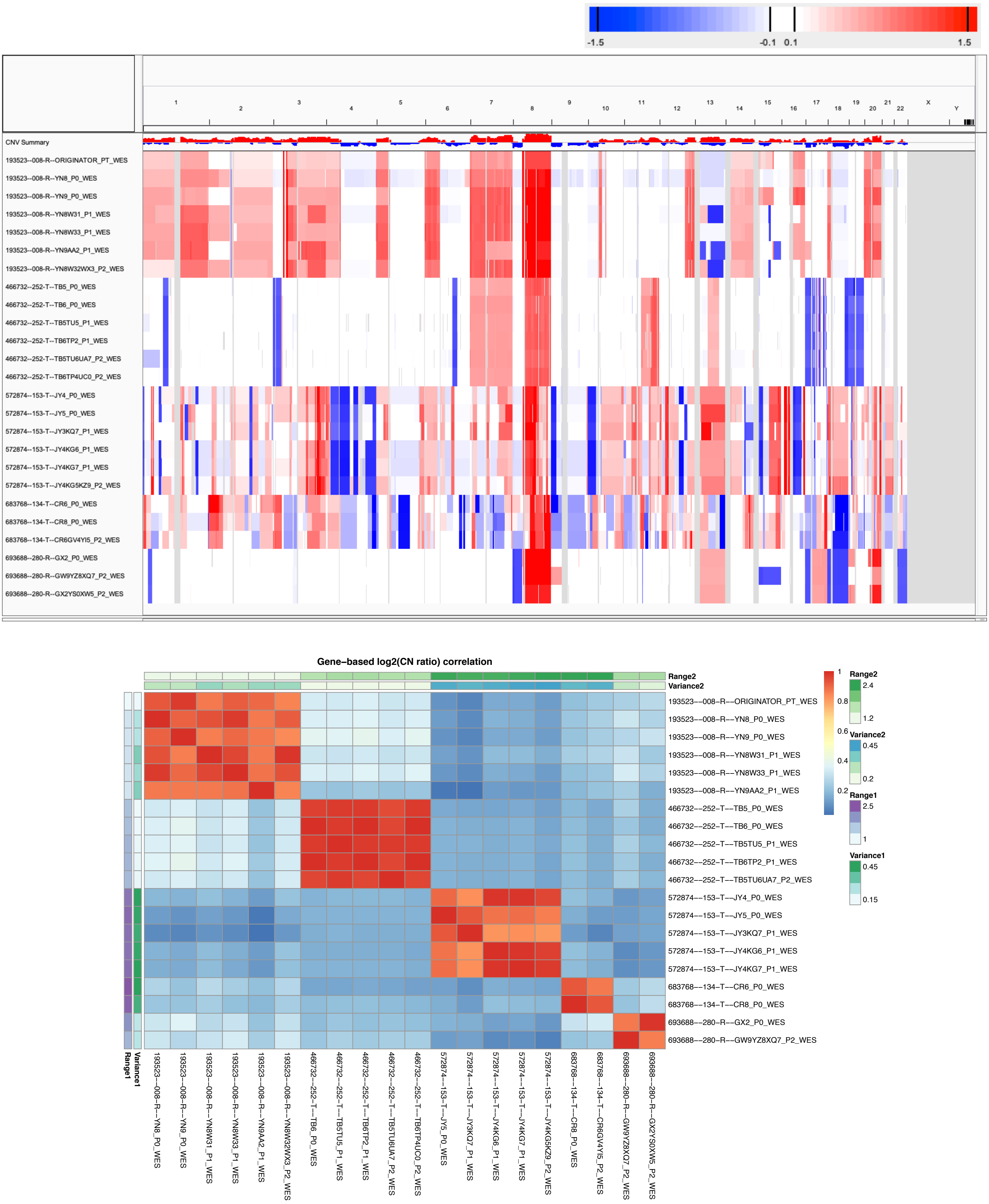

Supplementary Figure 58: CNA profiles (IGV heatmap) and correlation heatmap of gene-based copy number (log2(CN ratio), median centered) of samples from PDMR WES other cancers dataset.

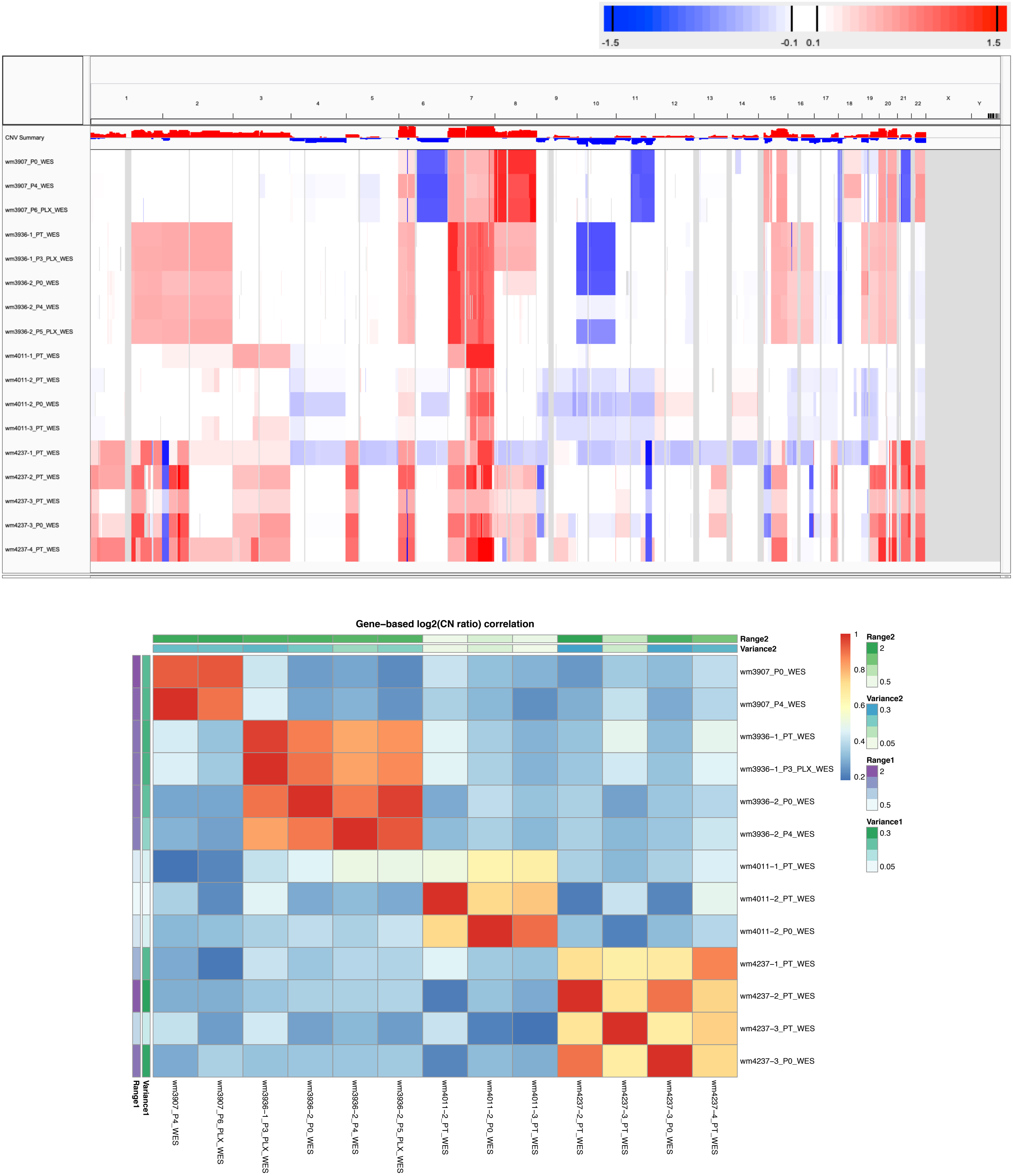

Supplementary Figure 59: CNA profiles (IGV heatmap) and correlation heatmap of gene-based copy number (log2(CN ratio), median centered) of samples from WISTAR WES skin melanoma dataset.

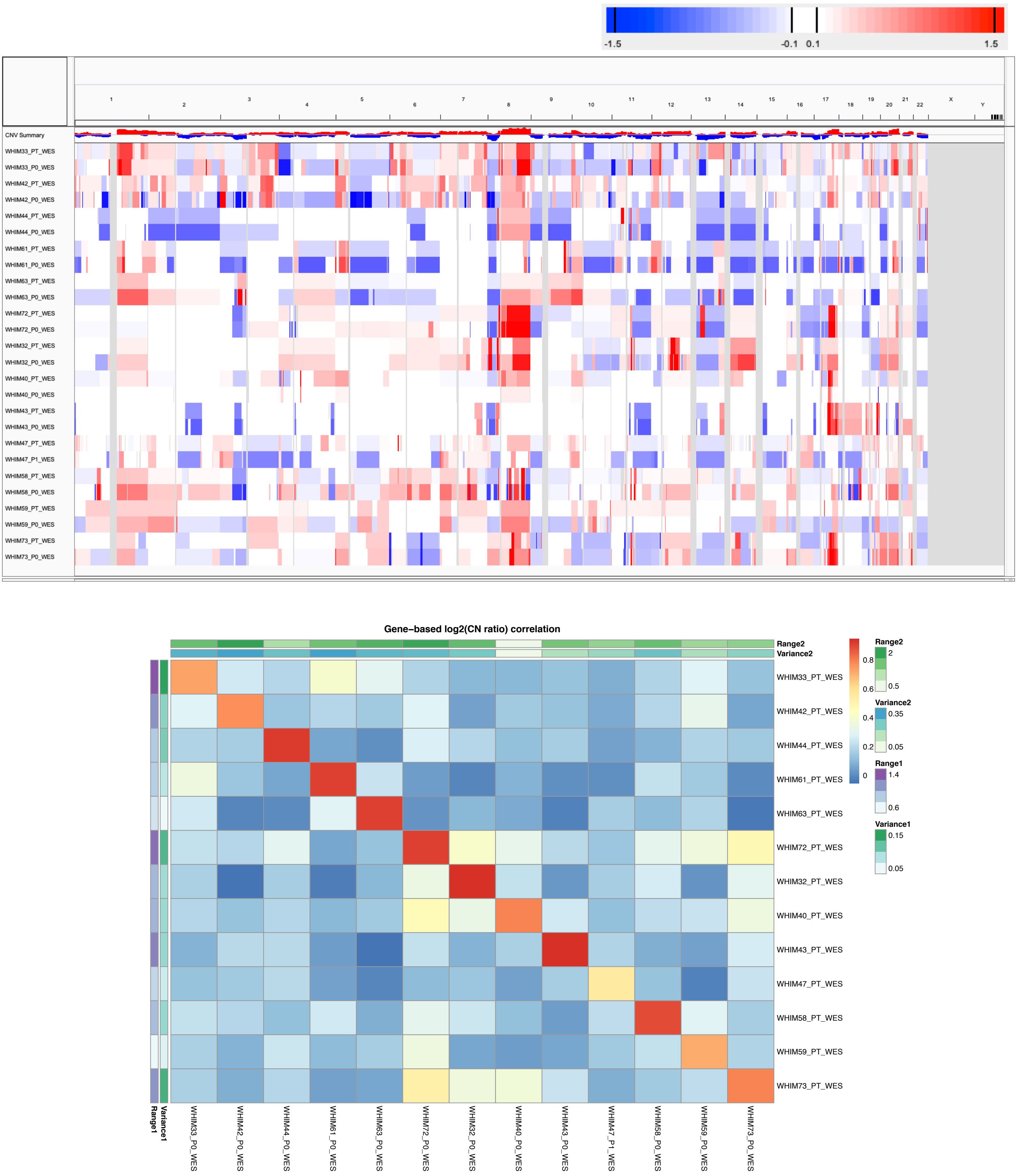

Supplementary Figure 60: CNA profiles (IGV heatmap) and correlation heatmap of gene-based copy number (log2(CN ratio), median centered) of samples from WUSTL WES breast cancer dataset.

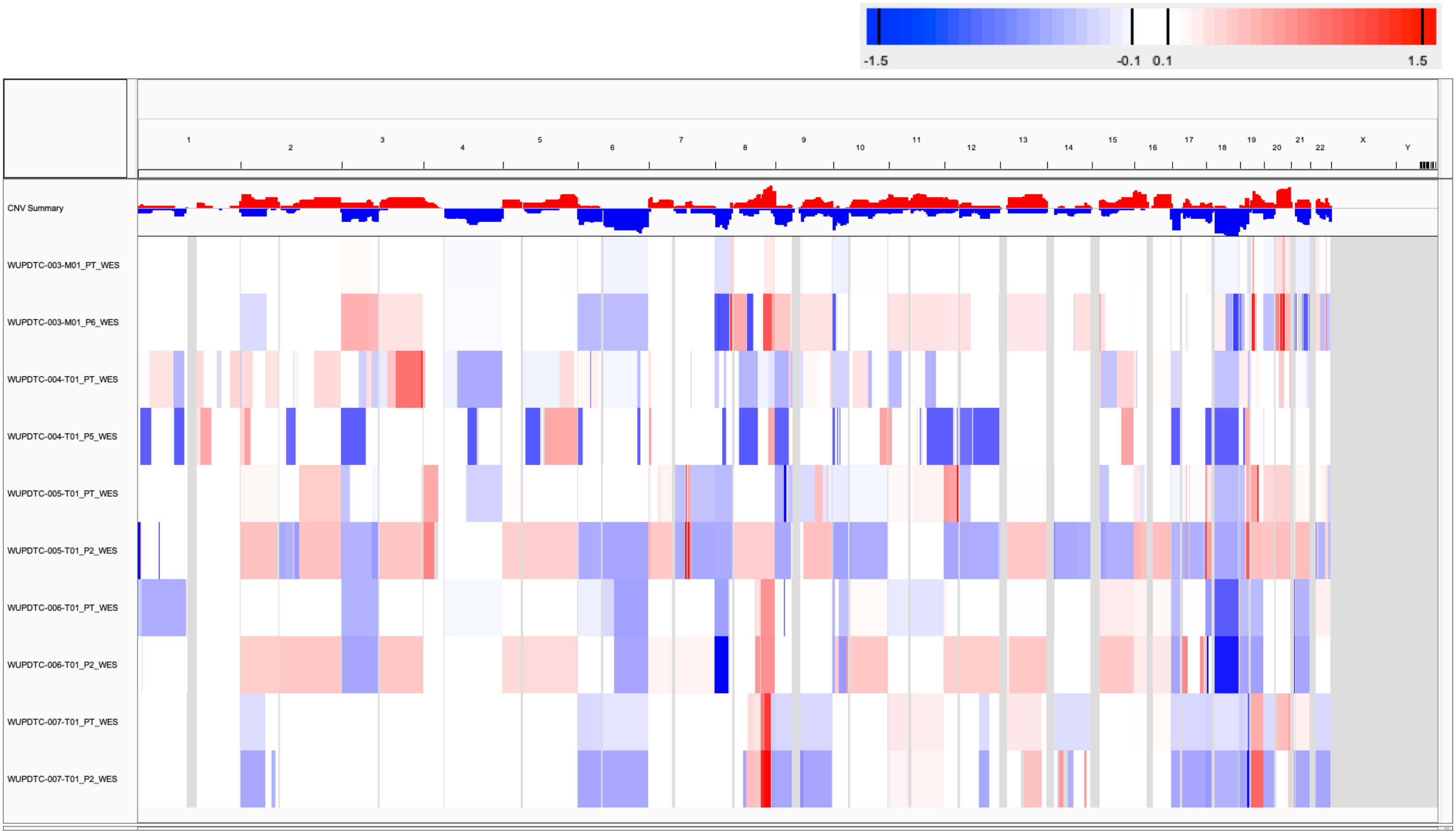

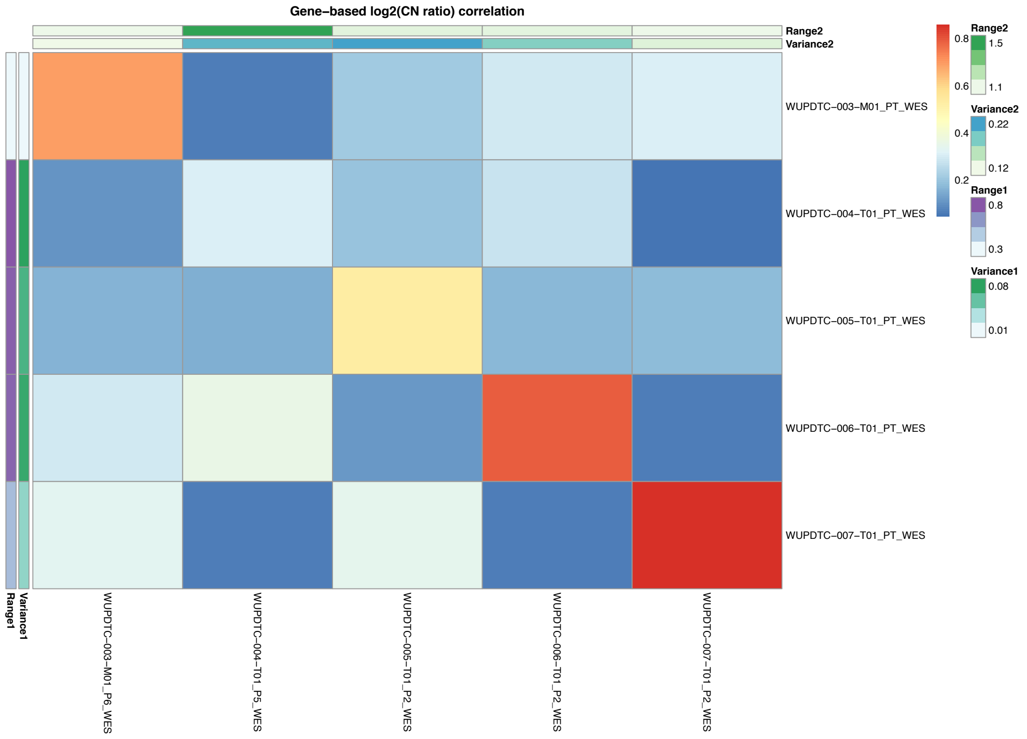

Supplementary Figure 61: CNA profiles (IGV heatmap) and correlation heatmap of gene-based copy number (log2(CN ratio), median centered) of samples from WUSTL WES pancreatic cancer dataset.

Supplementary Figure 62: (Continue next page)

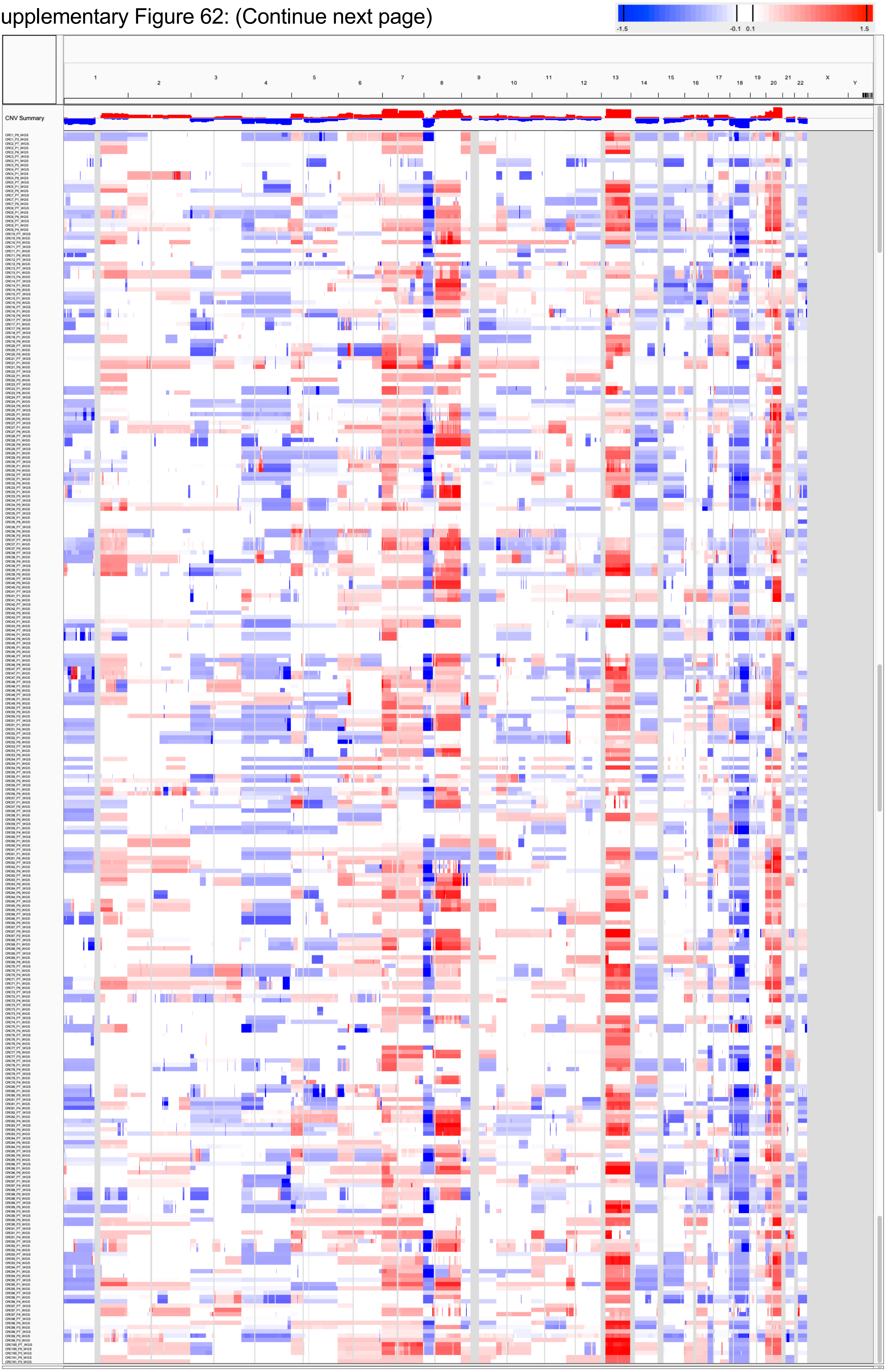

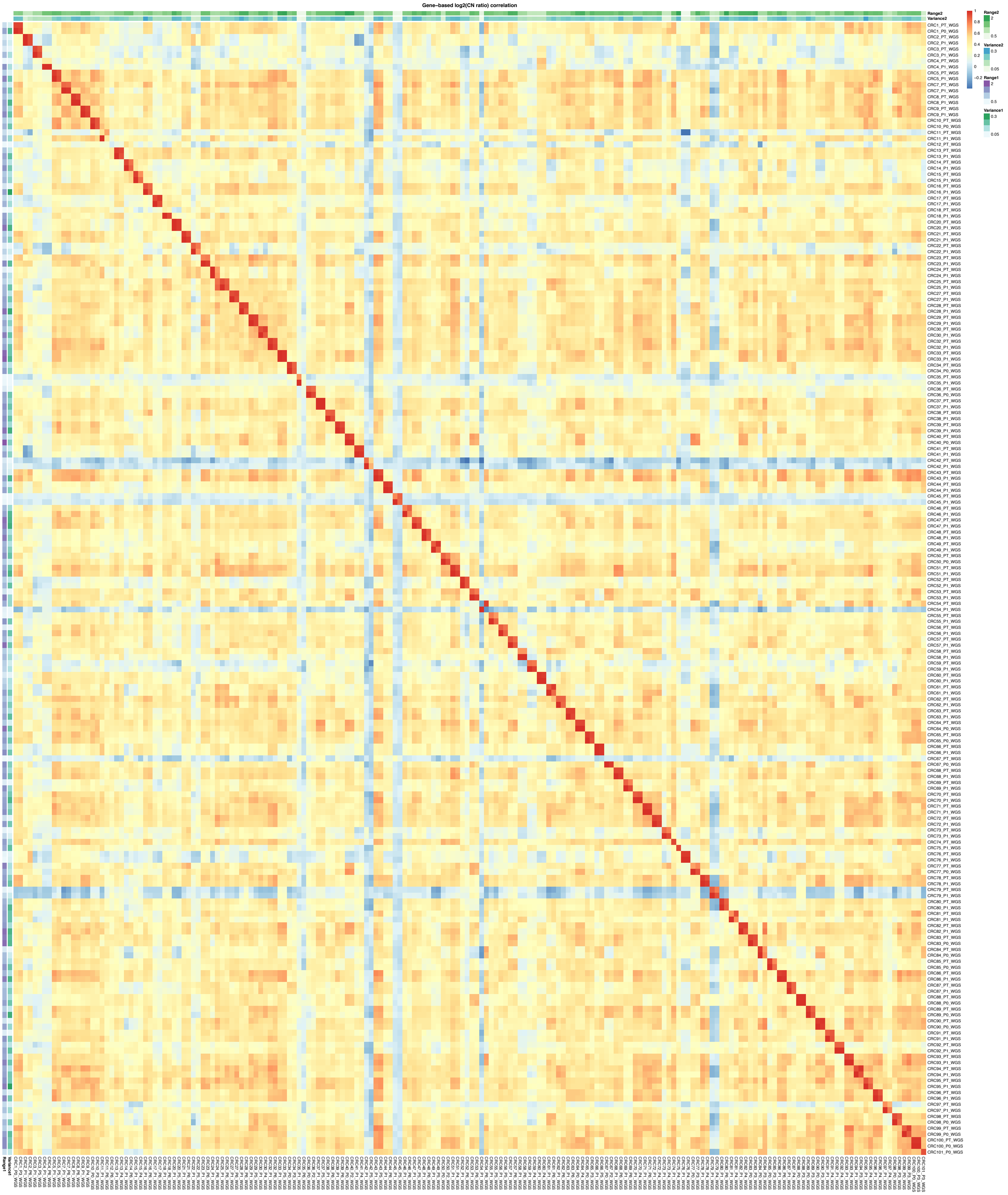

Supplementary Figure 62: CNA profiles (IGV heatmap) and correlation heatmap of gene-based copy number (log2(CN ratio), median centered) of samples from EuroPDX WGS colorectal cancer (liver metastases) dataset.

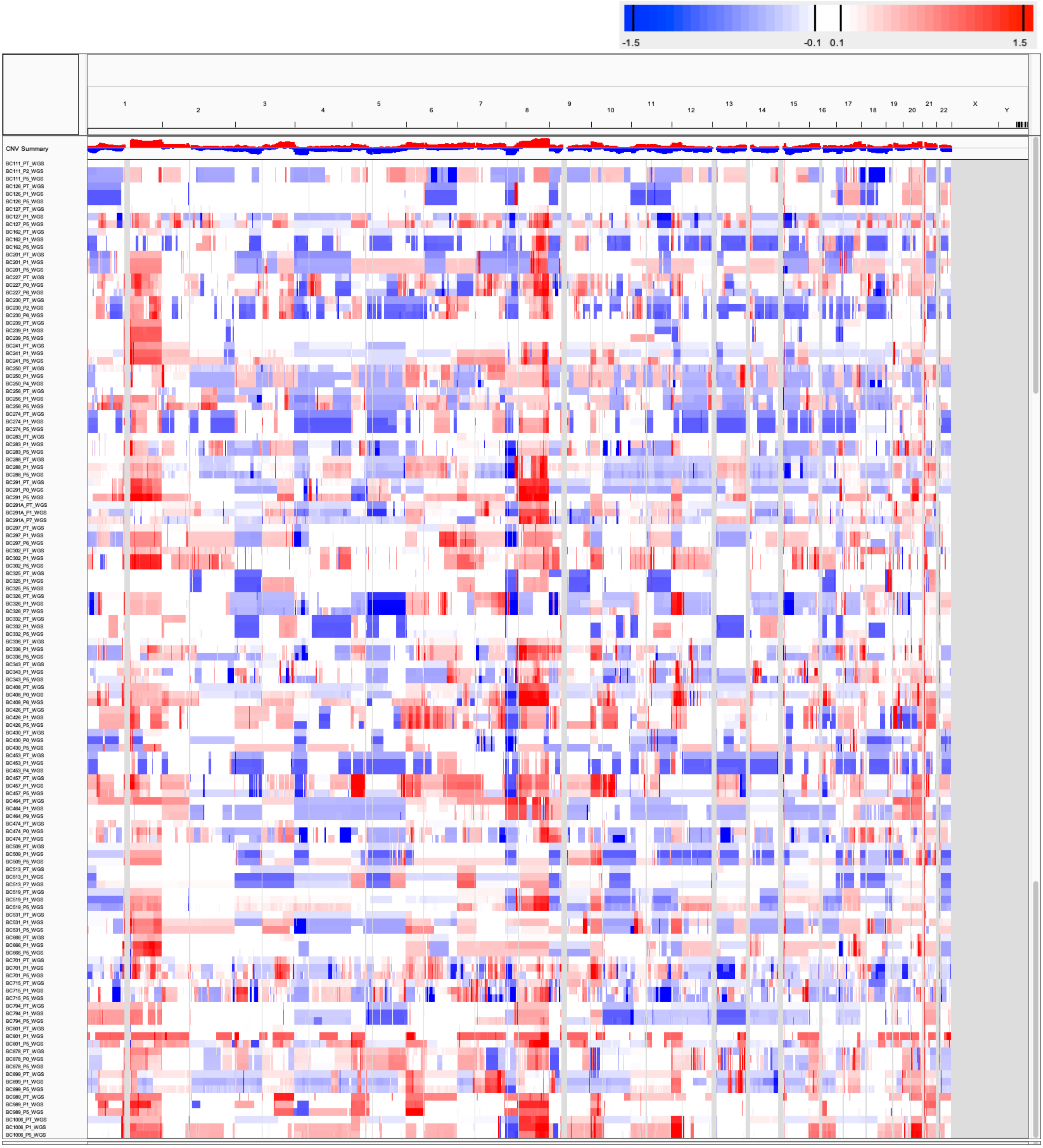

Supplementary Figure 63: (Continue next page)

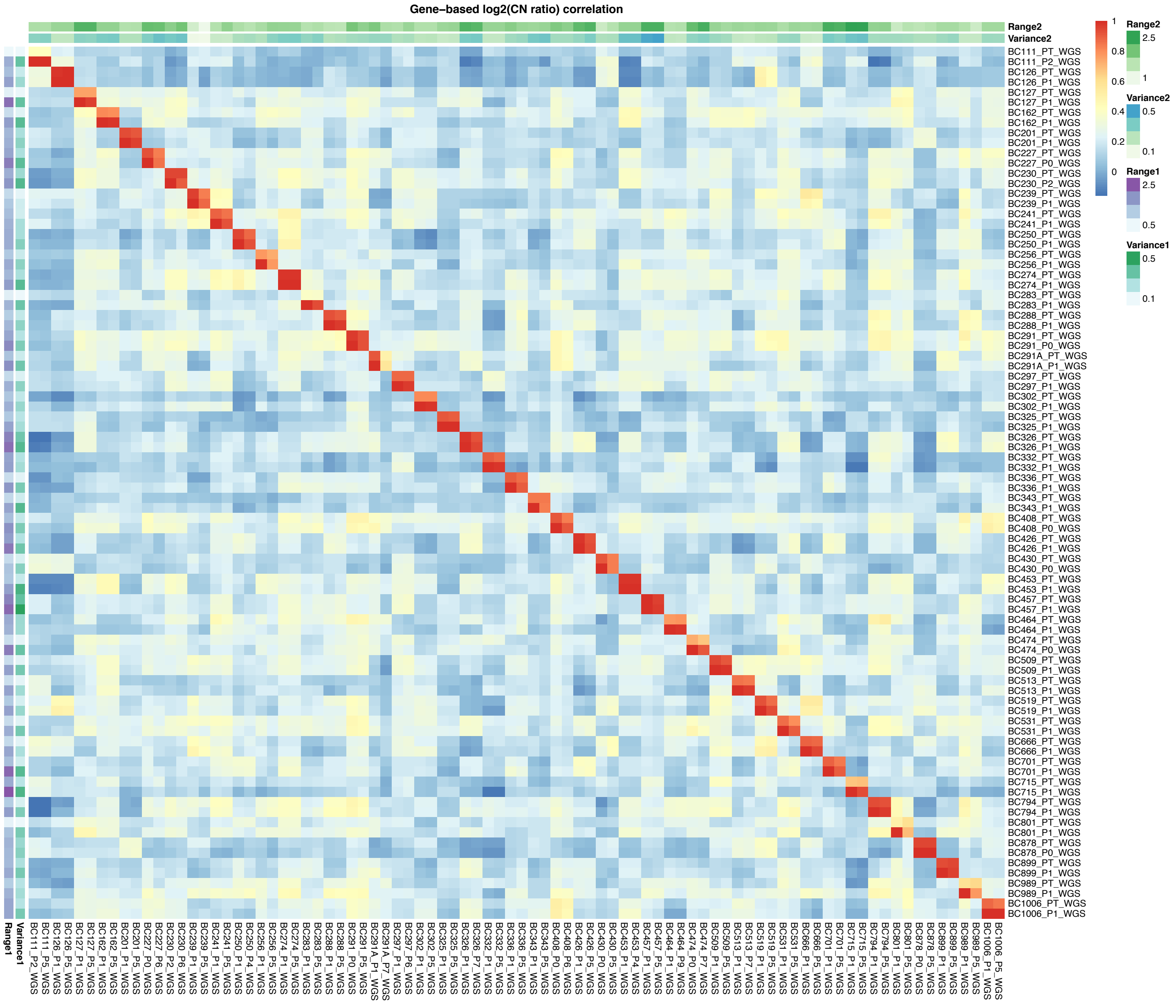

Supplementary Figure 63: CNA profiles (IGV heatmap) and correlation heatmap of gene-based copy number (log2(CN ratio), median centered) of samples from EuroPDX WGS breast cancer dataset.

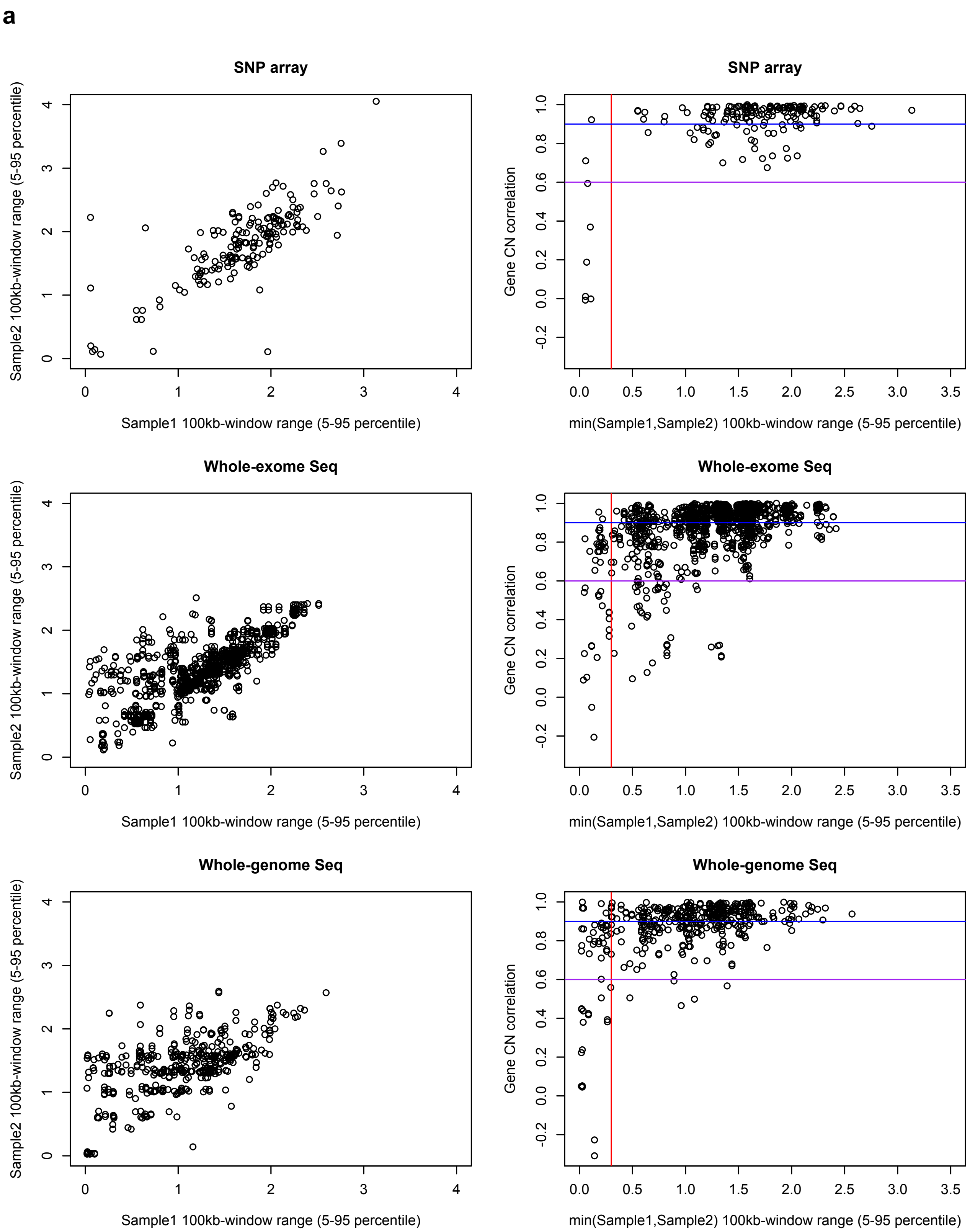

Supplementary Figure 64: (Continue next page)

**b**

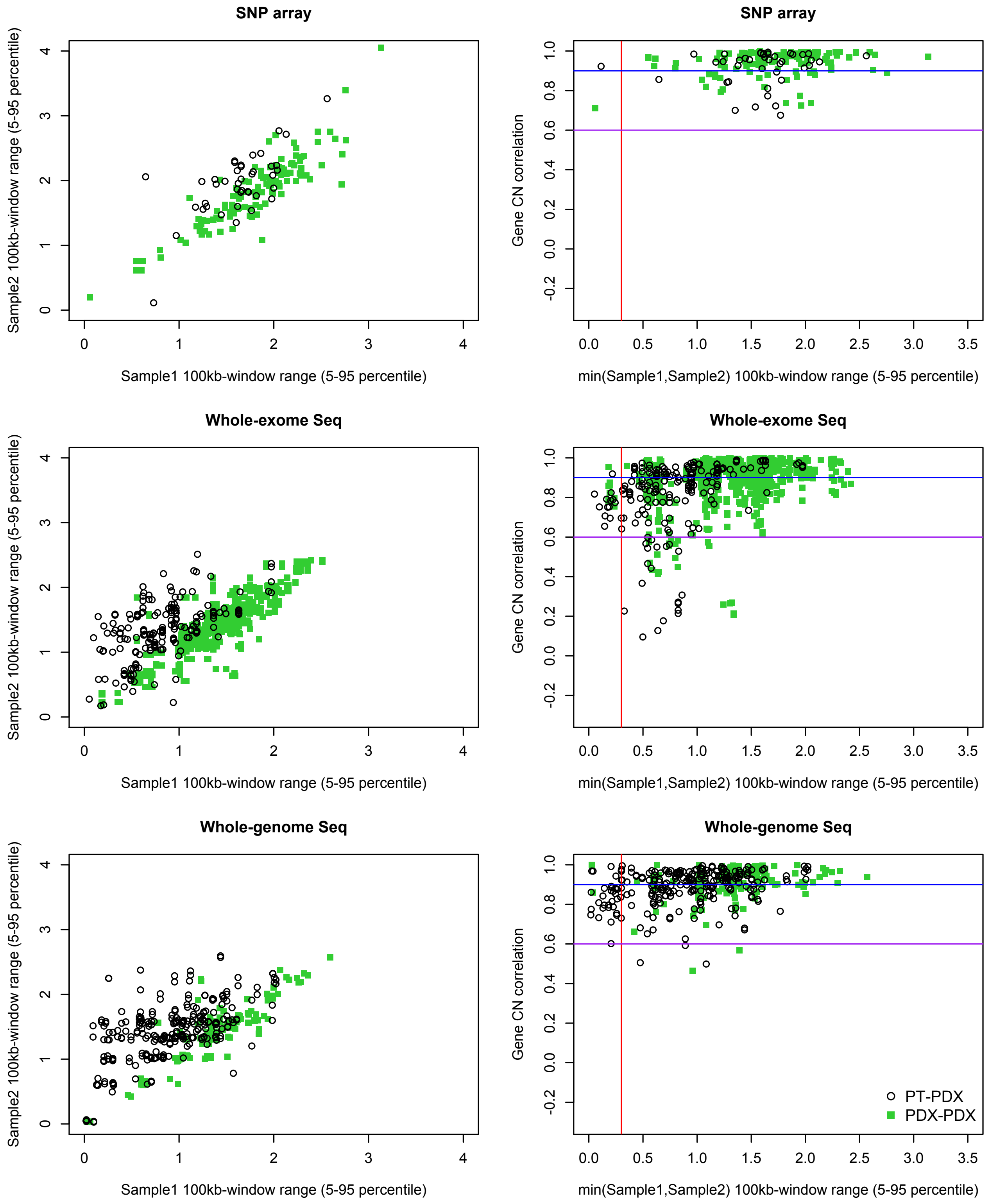

Supplementary Figure 64: PT samples have a lower range of CNA values than PDX samples. Comparison of range of CNA in pairs of samples (PT or PDX) from the same model (left panel). Pearson correlation of the samples versus the minimum range of the two samples (right panel). Samples with lower range tend to have lower correlations with other samples. For a given sample, range is defined as $log_2$(CN ratio) of the 95$^{th}$ percentile minus $log_2$(CN ratio) of the 5$^{th}$ percentile value of median-centered copy number values across 100-kb windows binned from copy number segments of each sample. (**a**) All data; (**b**) After removing comparisons of low correlation (< 0.6) due to non-aberrant samples (range < 0.3). (Sample 1: PT or lower passage PDX, Sample 2: later passage PDX or same passage PDX of different lineage)

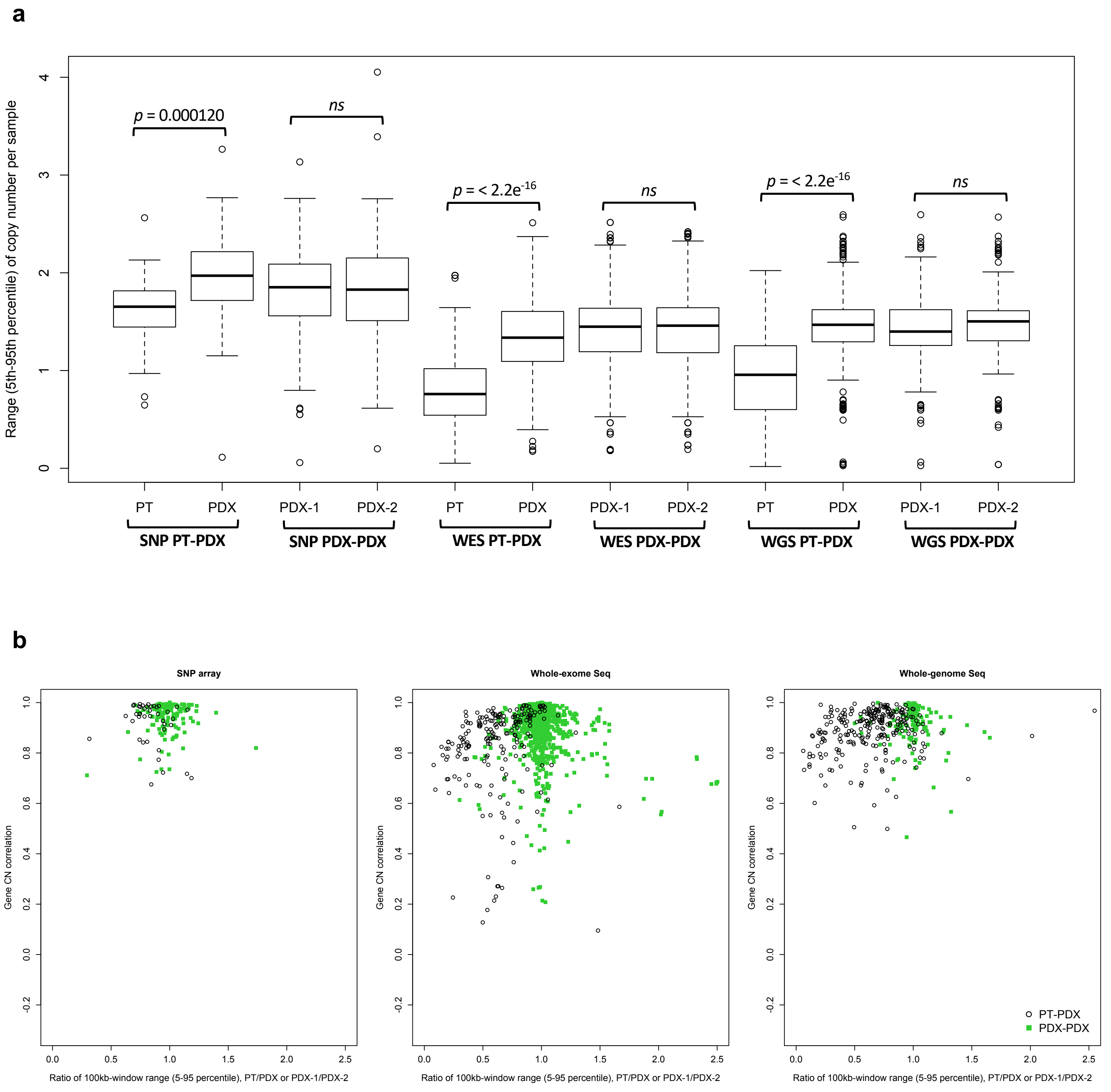

Supplementary Figure 65: (**a**) Range of CNA profiles between PT-PDX or PDX-PDX sample pairs from the same model. (**b**) Pearson correlation of the samples versus the ratio of range between two samples (PT/PDX or PDX-1/PDX-2). Samples pairs with ratio of range much greater or less than 1 i.e. one sample is much less aberrant than the other, tend to have lower correlations. For a given sample, range is defined as $\log_2$(CN ratio) of the 95$^{th}$ percentile minus $\log_2$(CN ratio) of the 5$^{th}$ percentile value of median-centered copy number values across 100-kb windows binned from copy number segments of each sample. P-values were computed by Wilcoxon rank sum test (ns: non-significant, $p > 0.05$). (PDX-1: lower passage PDX, PDX-2: later passage PDX or same passage PDX of different lineage)

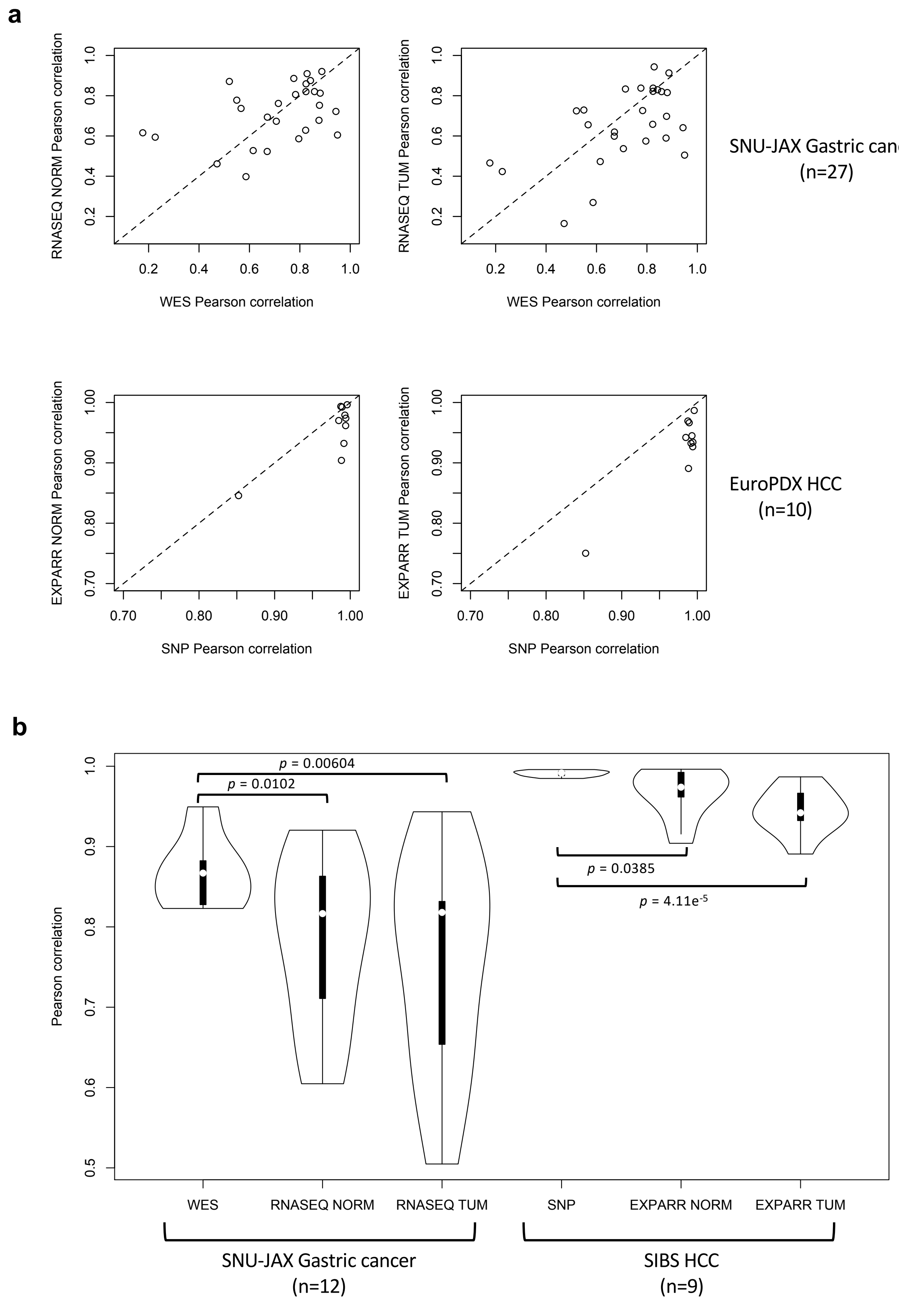

Supplementary Figure 66: (**a**) Comparison of Pearson correlation coefficients of gene-based copy number using DNA-based (WES or SNP array) versus RNA-based (RNA-Seq or gene expression array) methods. (**b**) Distribution of Pearson correlation coefficients of gene-based copy number, using DNA-based (WES or SNP array) and RNA-based (RNA-Seq or gene expression array) methods, for pairs of samples with high correlation by the DNA-based method (>0.8 for SNU-JAX Gastric cancer, >0.9 for SIBS HCC). P-values were computed by Wilcoxon rank sum test.

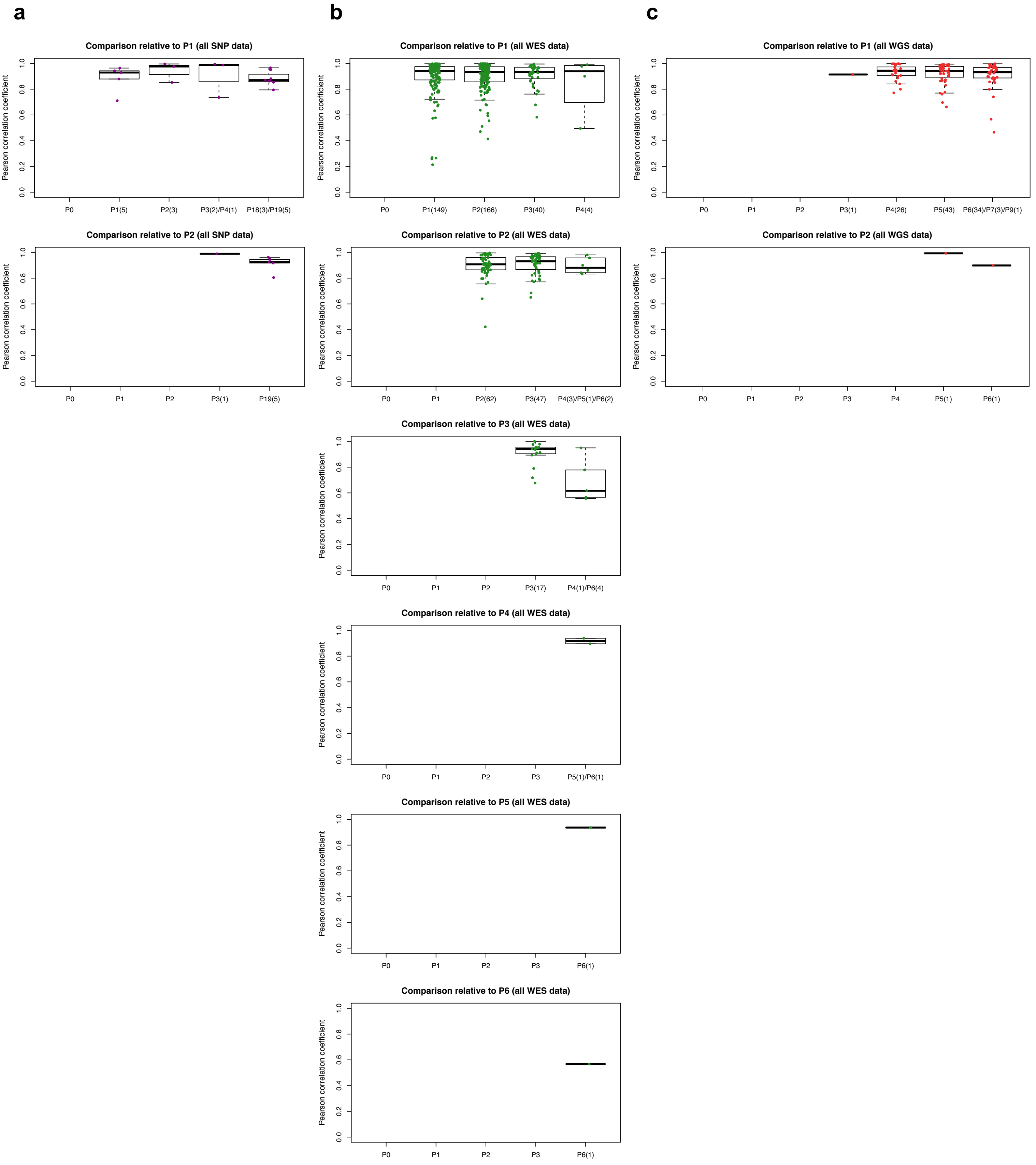

Supplementary Figure 67: Distribution of Pearson correlation coefficients of gene-based copy number, estimated by (**a**) SNP array, (**b**) WES, (**c**) WGS, between different combinations of patient tumor and PDX passages of the same model. Comparisons relative to passages P1 or later (refer to Fig. 3d – f for comparisons with PT and P0).

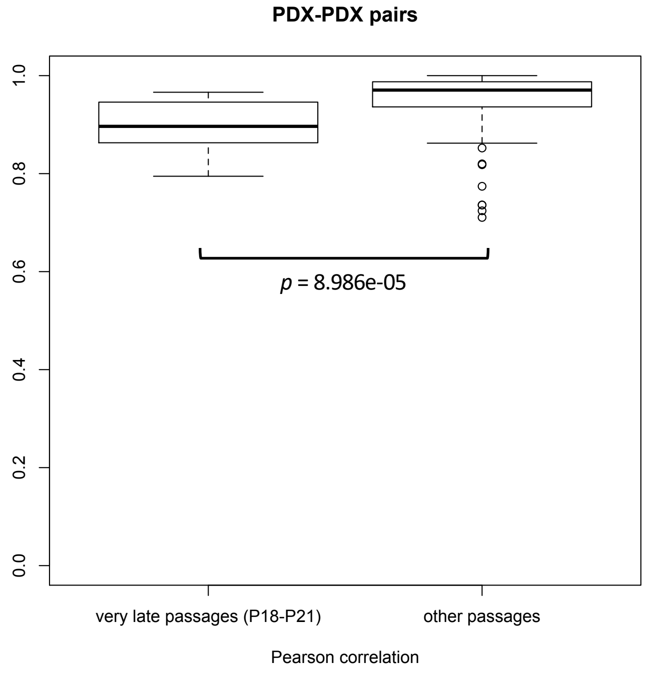

Supplementary Figure 68: Distribution of Pearson correlation coefficients of gene-based copy number between early and very-late passages of the same model for the BCM SNP array breast cancer dataset. Correlation for “other passages” are based on models from all other SNP array datasets.

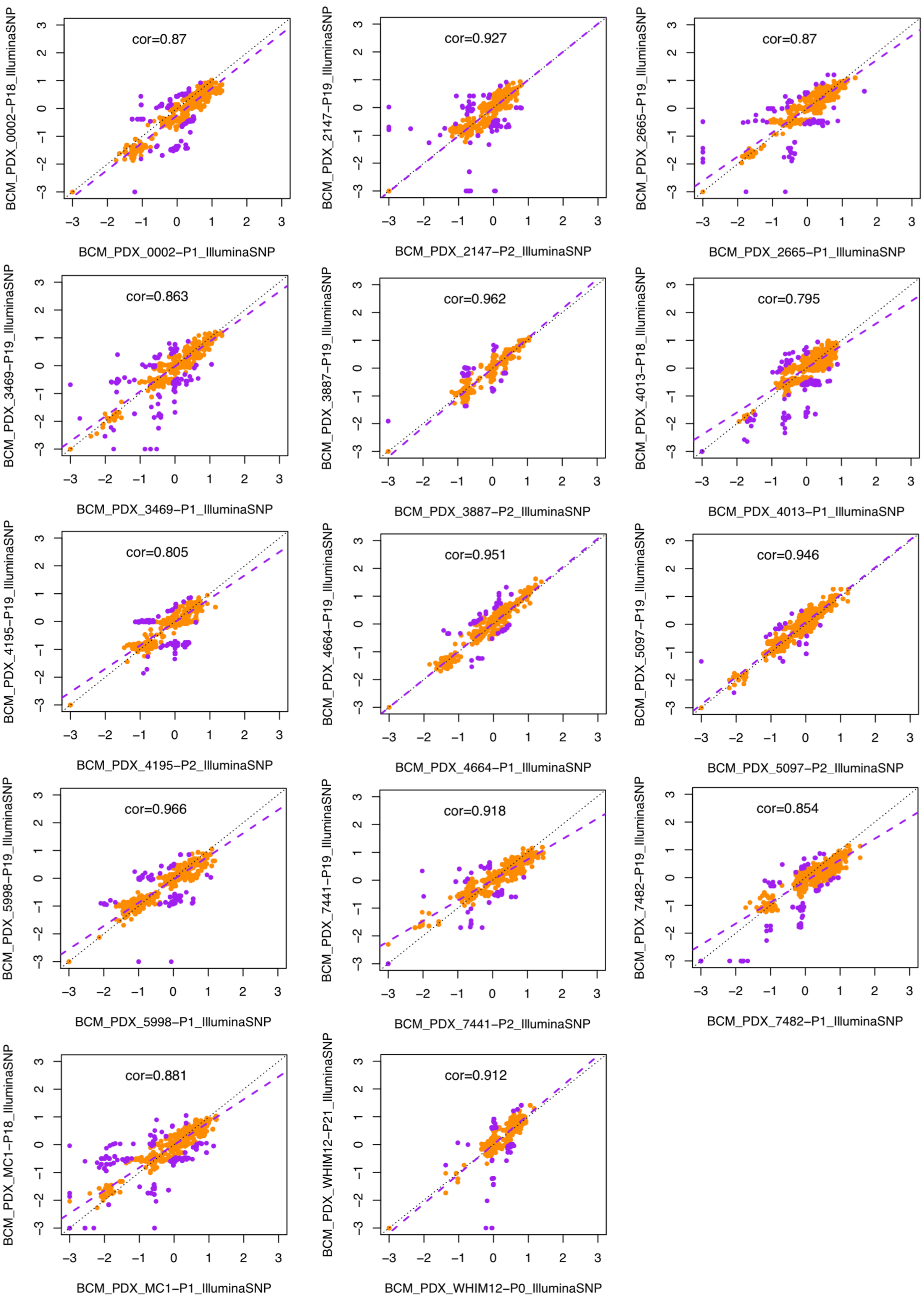

Supplementary Figure 69: Correlation and regression of gene-based copy number between early and very-late passages of the same model for the BCM SNP array breast cancer dataset. Genes with copy number changes between the passages are identified by |residual| > 0.5 (purple dots). Some genes show signs of complete deletion ($log_2$(CN ratio) < -2) but then reappear in later passages, suggesting bottlenecks from minor populations.

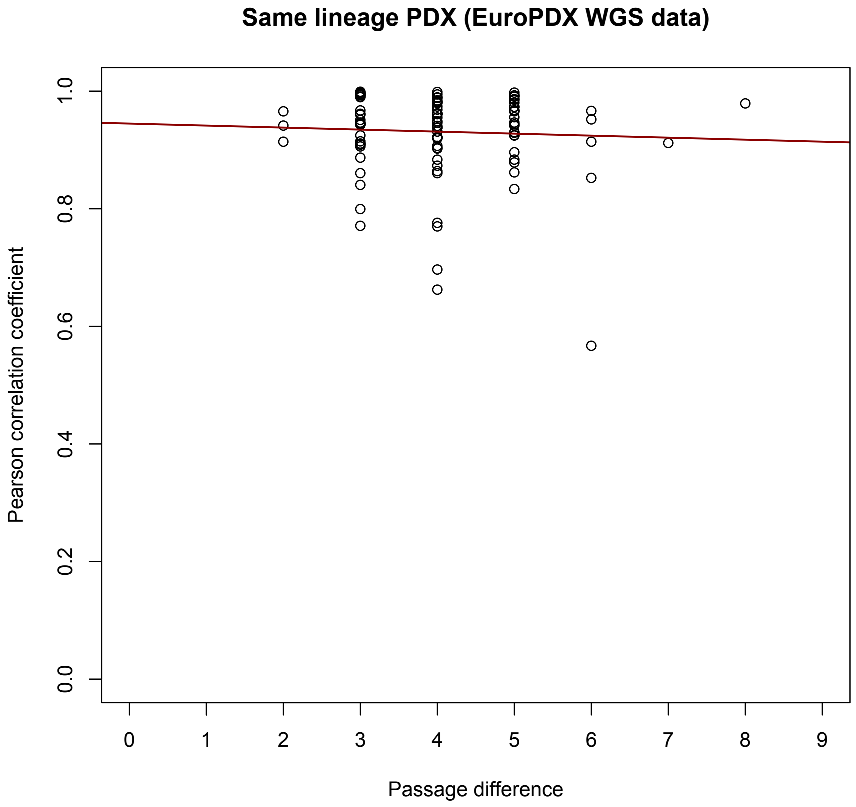

Supplementary Figure 70: Scatter plot of Pearson correlation between samples of PDX-early and PDX-late versus the corresponding passage difference for same lineage samples.

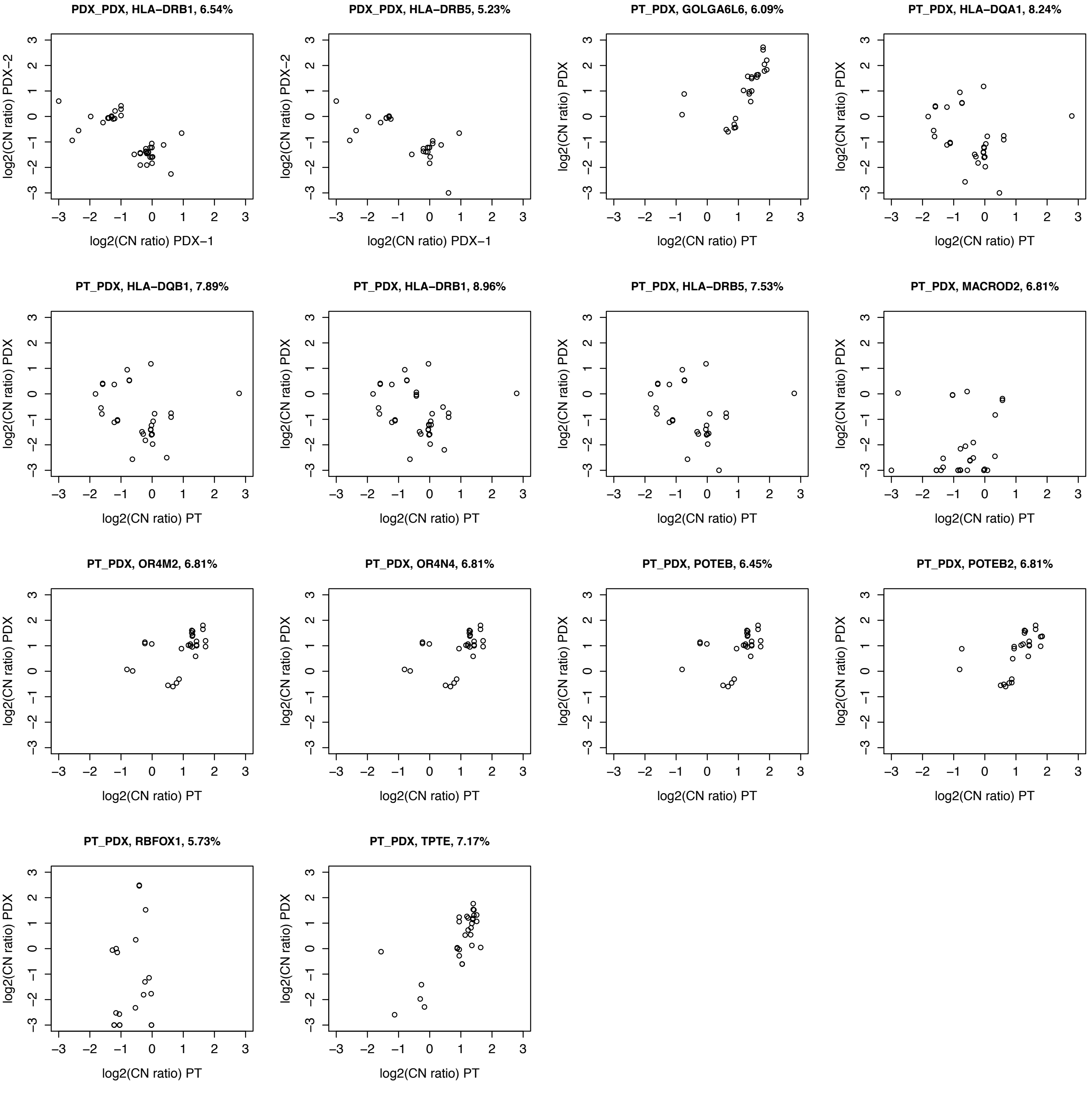

Supplementary Figure 71: $\log_2$(CN ratio) values between each pair of samples of recurrent genes (see Supplementary Table 4). PDX-1: earlier passage, PDX-2: same passage but different lineage or later passage.

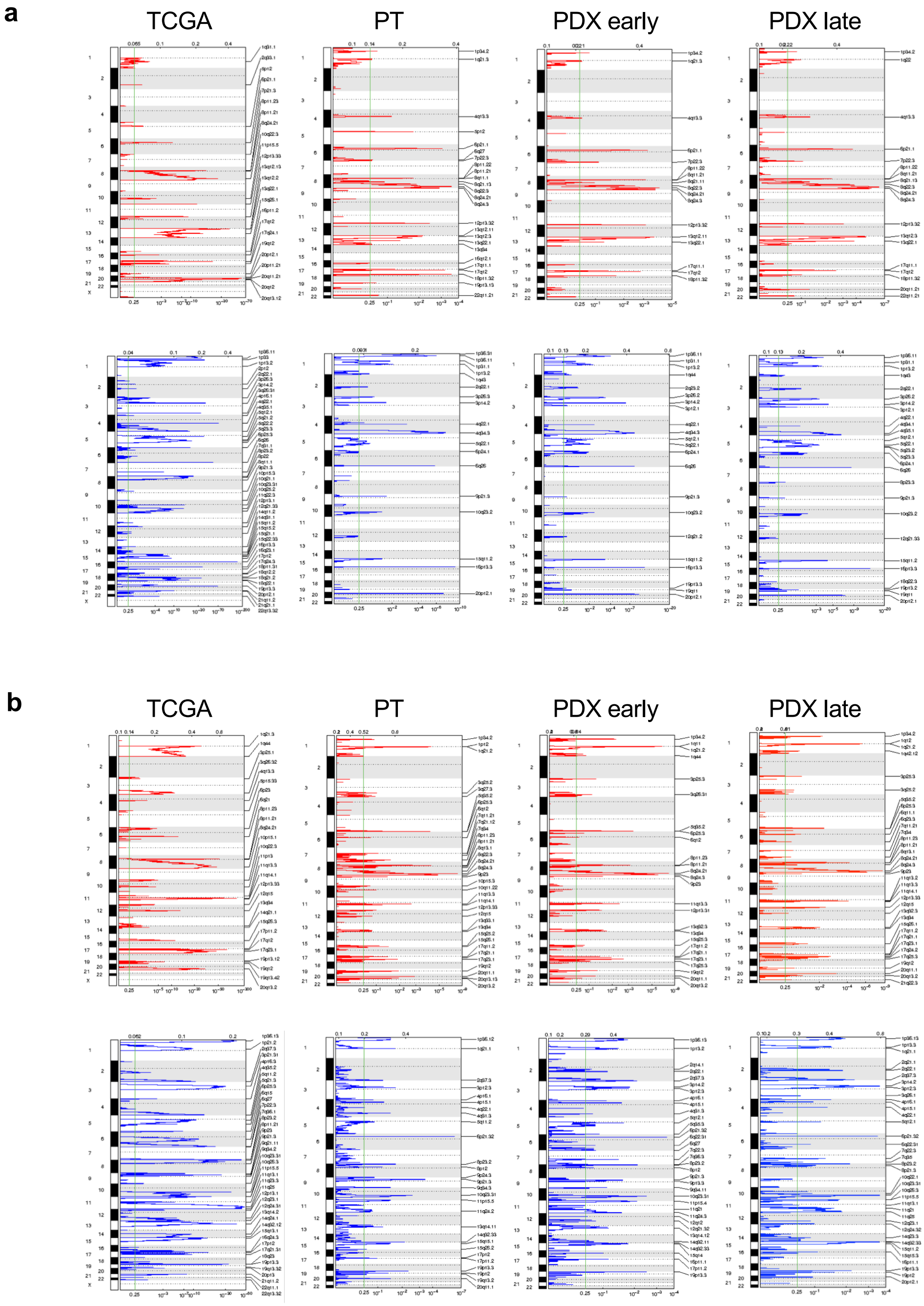

Supplementary Figure 72. GISTIC analysis of recurrent CNAs in TCGA primary tumors and EurOPDX collections of PTs and derived PDXs, at early and late passages, of (**a**) colorectal cancer and (**b**) breast cancer. For each GISTIC plot the top axis reports the G-score and the bottom axis the q-value. Red line plots: amplifications, blue line plots: deletions.

**a**

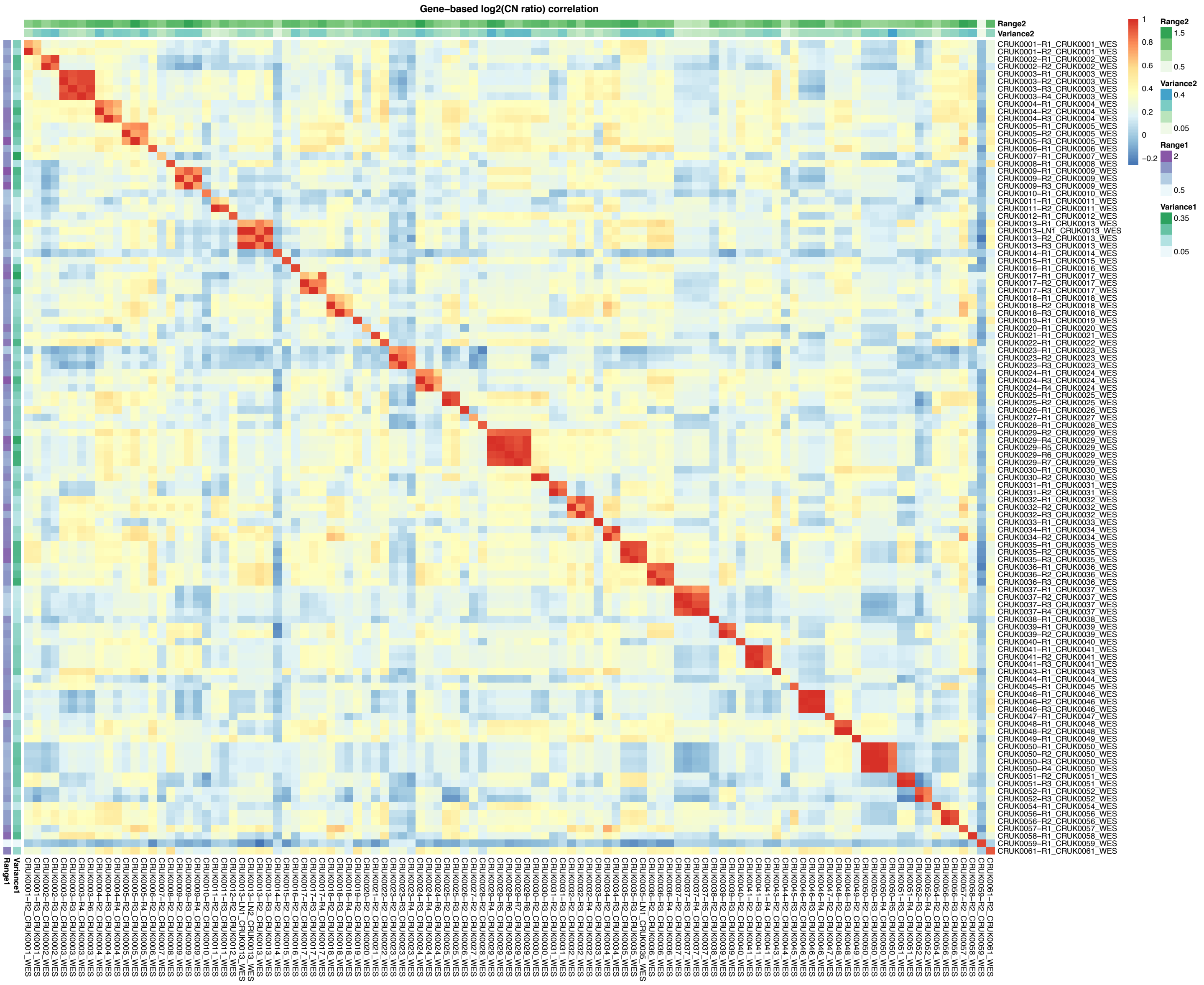

Supplementary Figure 73: (Continue next page)

**b**

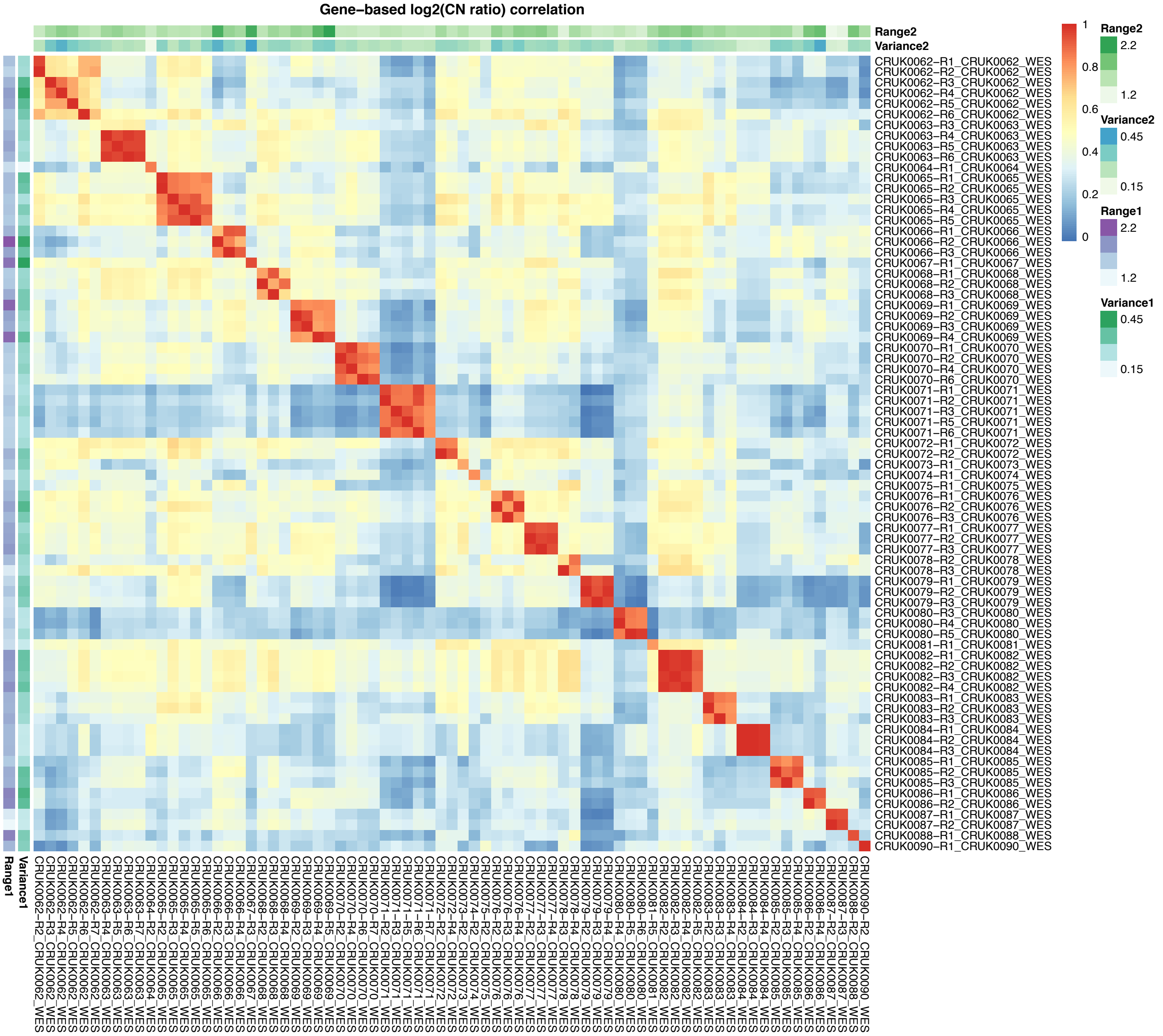

Supplementary Figure 73: (Continue next page)

**c**

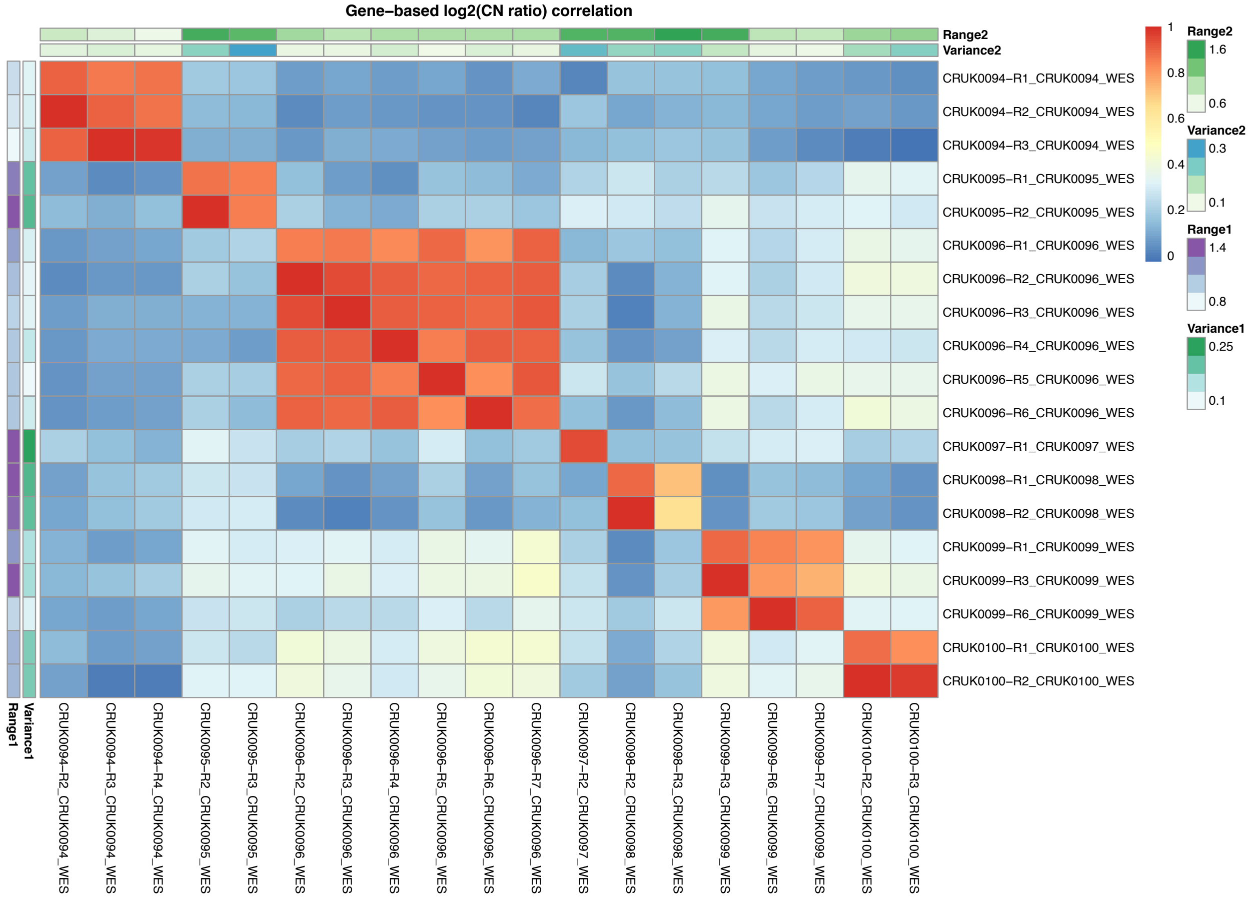

Supplementary Figure 73: Correlation heatmap of gene-based copy number (log2(CN ratio), median centered) of multi-region samples of the same tumor from TRACERx (**a**) lung adenocarcinoma (LUAD), (**b**) lung squamous cell carcinoma (LUSC) and (**c**) other lung cancer subtypes.

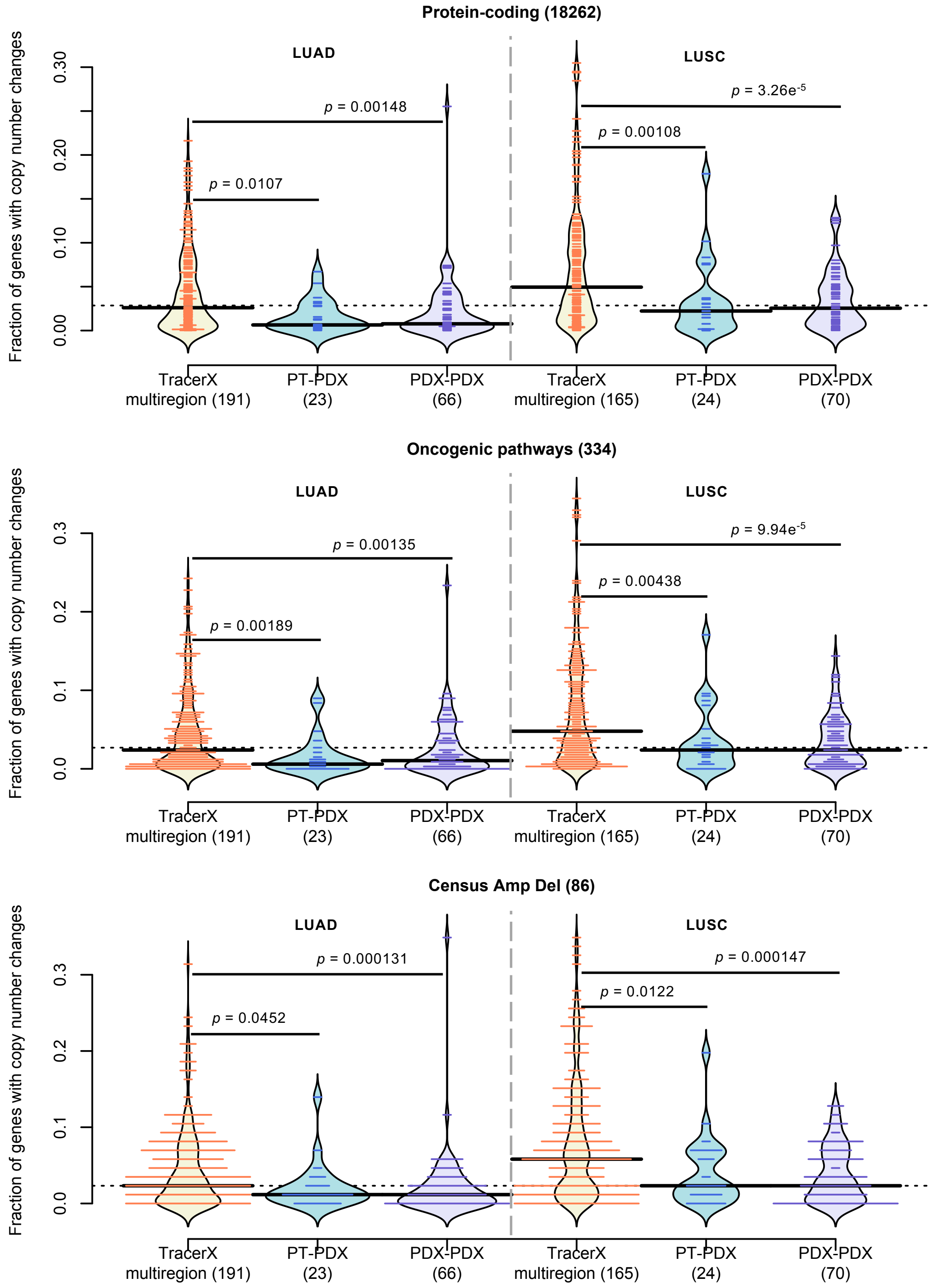

Supplementary Figure 74: Comparison of distribution of proportion of altered genes between multi-region tumor pairs from TRACERx, and PT-PDX and PDX-PDX pairs for various gene sets for LUAD and LUSC. Copy number altered genes were identified by |residual| > 0.5 from linear regression model for each pairwise comparison. P-values were computed by Wilcoxon rank sum test. Protein-coding: protein-coding genes annotated by Ensembl; Oncogenic pathways: genes in oncogenic signaling pathways identified by TCGA; Census Amp Del: genes with frequent amplifications or deletions annotated in the Cancer Gene Census

| | Model | BCM_PDX_0002 | BCM_PDX_2147 | BCM_PDX_2665 | BCM_PDX_3469 | BCM_PDX_3887 | BCM_PDX_4013 | BCM_PDX_4195 | BCM_PDX_4664 | BCM_PDX_5097 | BCM_PDX_5998 | BCM_PDX_7441 | BCM_PDX_7482 | BCM_PDX_MC1 | BCM_PDX_WHIM12 |
|---|---|---|---|---|---|---|---|---|---|---|---|---|---|---|---|
| | Early passage | 1 | 2 | 1 | 1 | 2 | 1 | 2 | 1 | 2 | 1 | 2 | 1 | 1 | 0 |
| | Late passage | 18 | 19 | 19 | 19 | 19 | 18 | 19 | 19 | 19 | 19 | 19 | 19 | 18 | 21 |
| | Correlation | 0.87 | 0.93 | 0.87 | 0.86 | 0.96 | 0.79 | 0.8 | 0.95 | 0.95 | 0.97 | 0.92 | 0.85 | 0.88 | 0.91 |
| Fraction of genes with CN changes | Protein coding genes | 0.05 | 0.02 | 0.04 | 0.05 | 0.01 | 0.04 | 0.11 | 0.01 | 0.01 | 0.02 | 0.04 | 0.02 | 0.06 | 0.01 |
| | Oncogenic signalling pathways | 0.04 | 0.03 | 0.04 | 0.03 | 0.01 | 0.06 | 0.12 | 0.02 | 0.01 | 0.01 | 0.03 | 0.02 | 0.07 | 0.02 |
| | JAX CKB Amp/Over-expr | 0.05 | 0.04 | 0.02 | 0.06 | 0.02 | 0.05 | 0.13 | 0.02 | 0.01 | 0.01 | 0.03 | 0.02 | 0.04 | 0.02 |
| | JAX CKB Del/Under-expr | 0.02 | 0.01 | 0.06 | 0.04 | 0.01 | 0.06 | 0.12 | 0.01 | 0.02 | 0 | 0.04 | 0.02 | 0.03 | 0.01 |
| | Cancer Gene Census (Amp/Del) | 0.1 | 0.02 | 0.02 | 0.06 | 0.01 | 0.01 | 0.09 | 0 | 0.03 | 0 | 0.05 | 0.05 | 0.02 | 0.02 |
| | TCGA Breast Gistic Amp | 0 | 0.07 | 0.01 | 0.3 | 0.11 | 0.01 | 0.01 | 0.02 | 0 | 0 | 0.03 | 0.01 | 0 | 0 |
| | TCGA Breast Gistic Del | 0.03 | 0.03 | 0.03 | 0.03 | 0 | 0.1 | 0.07 | 0.01 | 0.02 | 0.01 | 0.03 | 0.02 | 0.06 | 0.01 |

Supplementary Figure 75: Fraction of genes of different gene sets (see Fig. 4) with copy number changes (|residual| > 0.5) between early and late passages of each breast cancer model from the BCM breast cancer dataset.

**a**

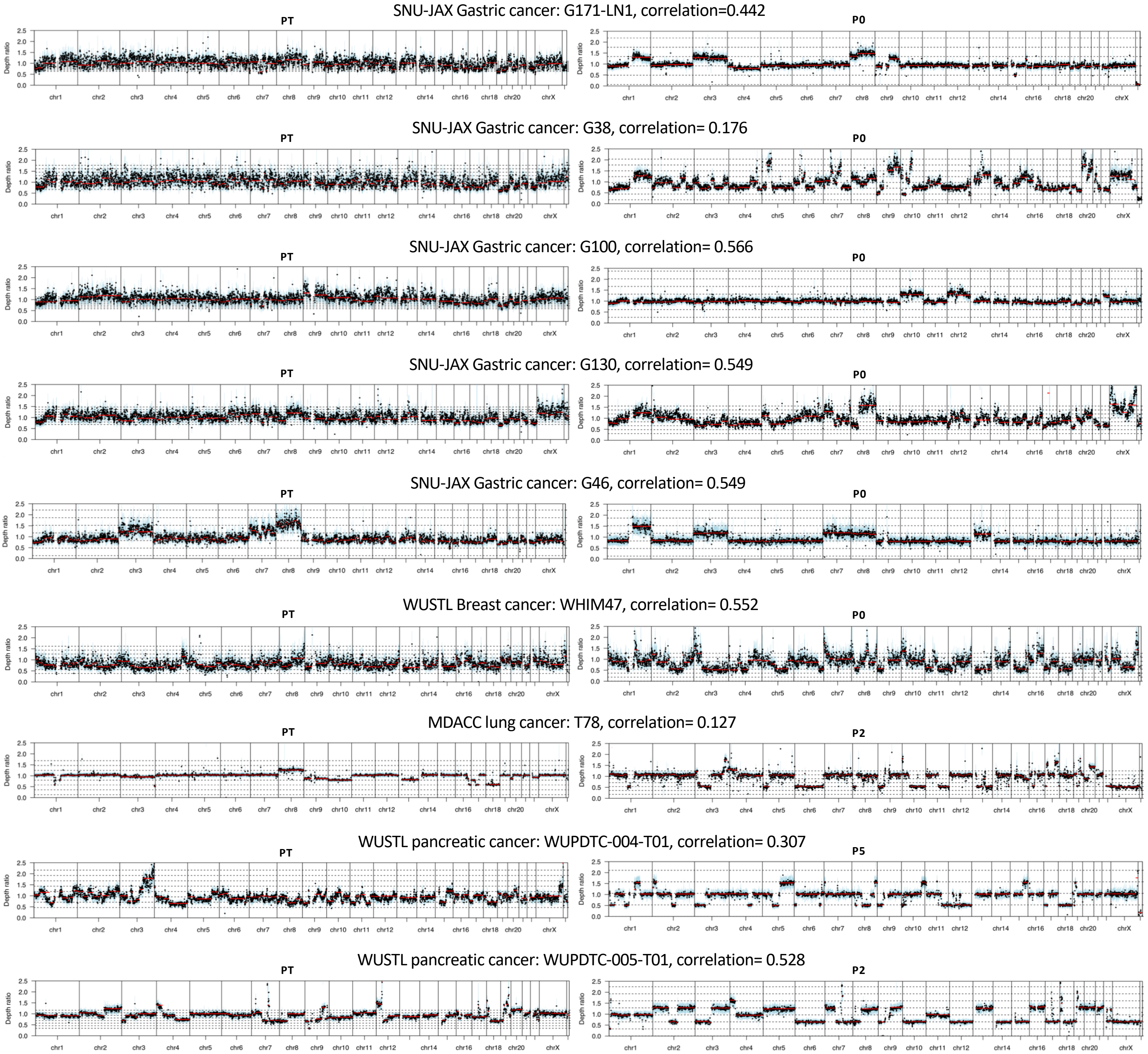

Supplementary Figure 76: (Continue next page)

**b**

EuroPDX BRCA: BC291A
Correlation=
PT vs P1: 0.907
PT vs P7: 0.498
P1 vs P7: 0.566

EuroPDX BRCA: BC989
Correlation=
PT vs P1: 0.592
PT vs P5: 0.625
P1 vs P5: 0.769

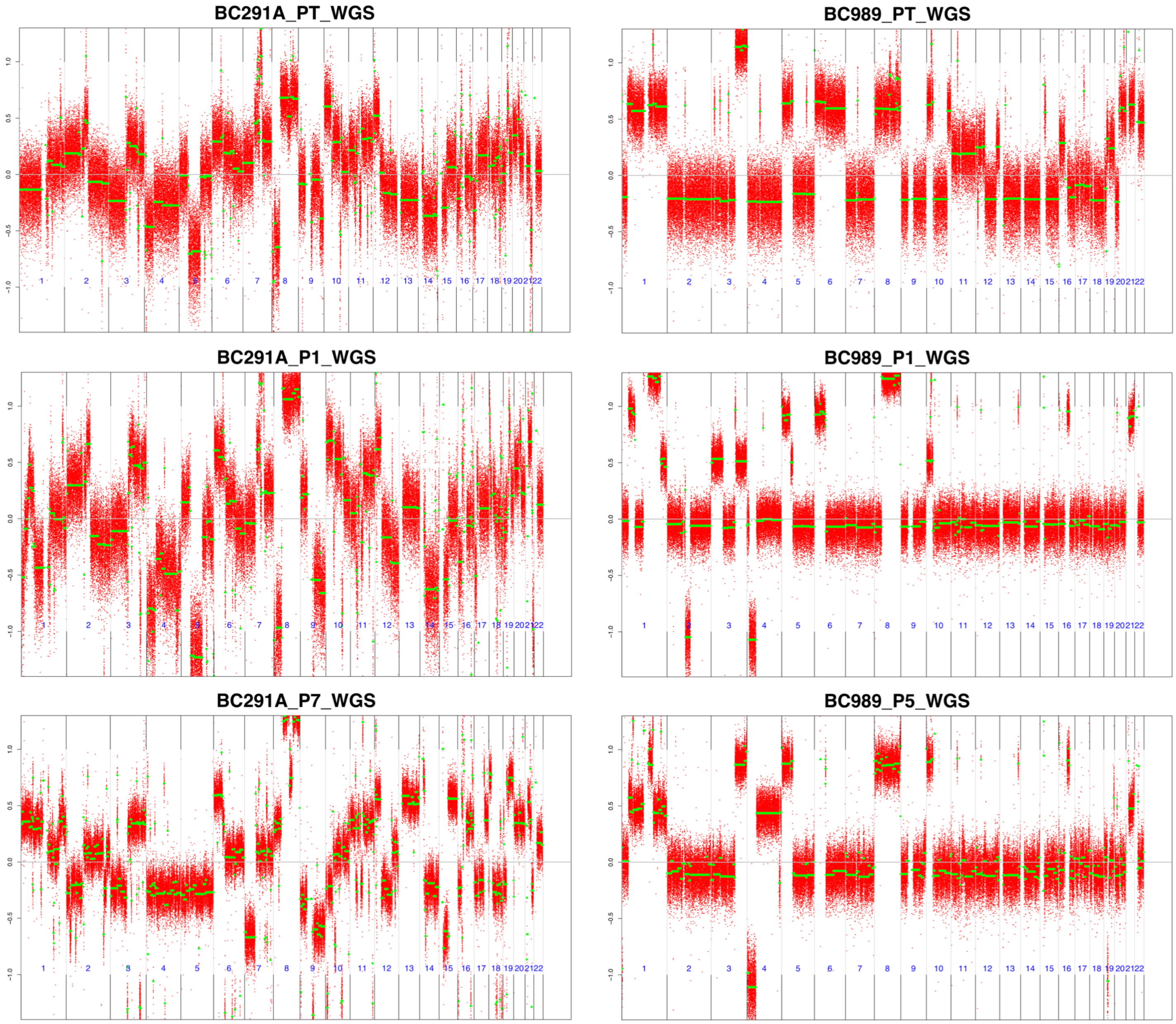

Supplementary Figure 76: Window-based and segmented copy number estimated as (**a**) depth ratio by Sequenza for WES, and (**b**) $\log_2$ (ratio) by ASCAT for WGS, for same lineage samples identified to be aberrant (range > 0.5 for both samples) and low correlation (Pearson correlation coefficient < 0.6) between the sample pairs

**a**

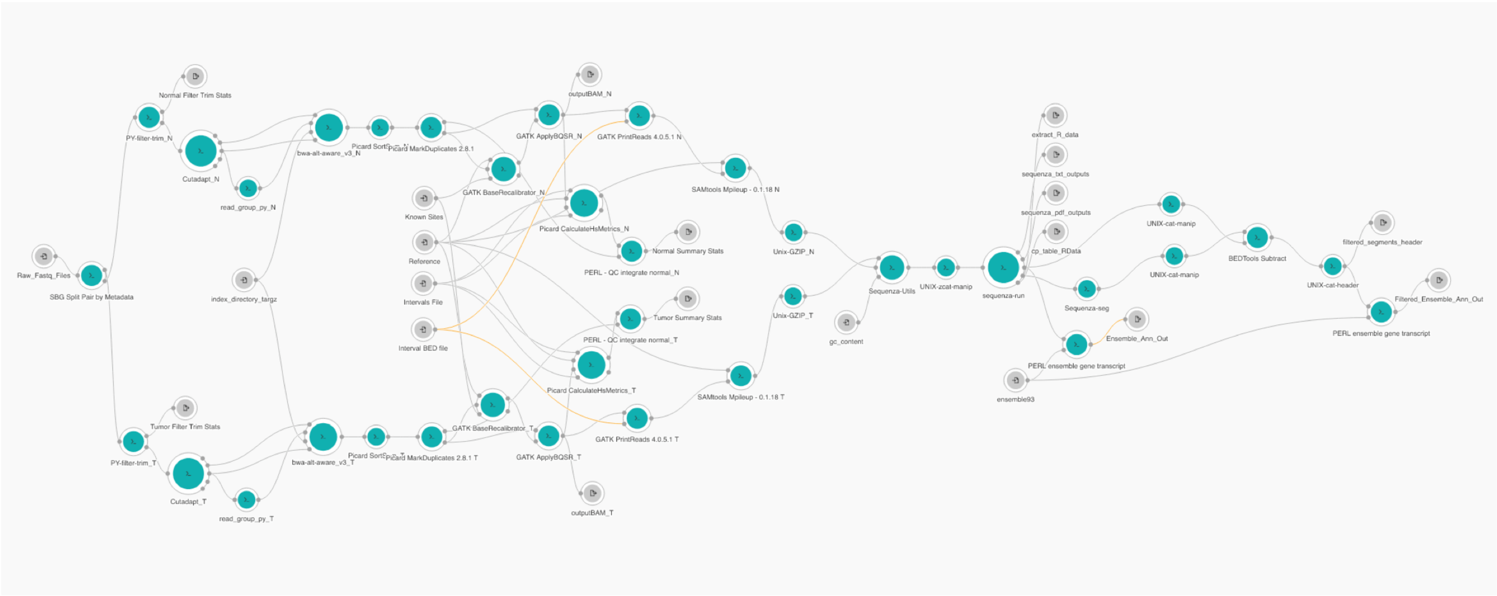

**b**

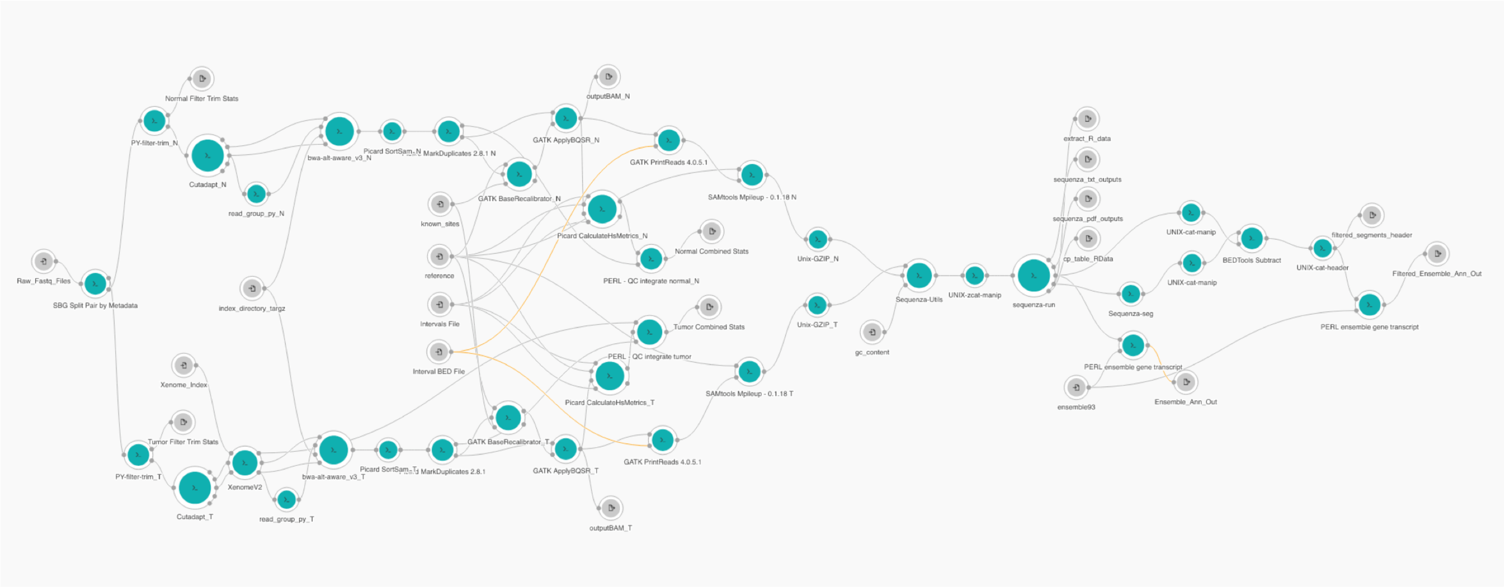

Supplementary Figure 77: Workflow for copy number estimation by Sequenza from whole-exome sequencing data with paired-normal for (a) patient tumor (without Xenome) and (**b**) PDX tumor (with Xenome for mouse reads removal).

| Data source | Number of unique models | SNP array (single tumor) | | | Whole-exome sequencing (tumor-normal) | | | Whole-genome sequencing (single tumor) | | | RNA-Seq or Microarray (tumor-normal) | | | Number of unique samples | |
|---|---|---|---|---|---|---|---|---|---|---|---|---|---|---|---|
| | | Number of models | Number of patient tumor samples | Number of PDX samples | Number of models | Number of patient tumor samples | Number of PDX samples | Number of models | Number of patient tumor samples | Number of PDX samples | Number of models | Number of patient tumor samples | Number of PDX samples | PT | PDX |
| JAX PDX resource (various tumor types) | 105 | 105 | 27 | 203 | 3 | 3 | 0 | | | | | | | 27 | 203 |
| SNU-JAX gastric cancer | 48 | | | | 33 | 33 | 33 | | | | 42 | 42 | 42 | 48 | 48 |
| HCI breast cancer | 18 | 13 | 12 | 16 | 10 | 9 | 9 | | | | | | | 16 | 25 |
| BCM breast cancer | 14 | 14 | 0 | 28 | | | | | | | | | | 0 | 28 |
| MDACC lung cancer | 45 | | | | 45 | 45 | 45 | | | | | | | 45 | 45 |
| Wistar melanoma | 10 | | | | 10 | 8 | 9 | | | | | | | 8 | 9 |
| NCI PDMR (various tumor types) | 83 | | | | 83 | 21 | 439 | | | | | | | 21 | 439 |
| WUSTL pancreatic cancer | 5 | | | | 5 | 5 | 5 | | | | | | | 5 | 5 |
| WUSTL breast cancer | 13 | | | | 13 | 13 | 13 | | | | | | | 13 | 13 |
| SIBS hepatocellular carcinoma | 28 | 28 | 2 | 33 | | | | | | | 28 | 3 | 34 | 3 | 34 |
| EuroPDX colorectal cancer (liver metastasis) | 97 | | | | | | | 97 | 95 | 192 | | | | 95 | 192 |
| EuroPDX breast cancer | 43 | | | | | | | 43 | 43 | 86 | | | | 43 | 86 |
| Total | **509** | **160** | **41** | **280** | **202** | **137** | **553** | **140** | **138** | **278** | **70** | **45** | **76** | **324** | **1127** |

Supplementary Table 1: Summary of datasets collected from various centers in the PDXNET consortium, EuroPDX consortium and published datasets.

| Tumor type | Total models | Models with patient tumor samples | Models with multiple PDX samples | Number of PDX samples |
|---|---|---|---|---|
| Colorectal cancer | 130 | 100 | 123 | 299 |
| Breast cancer | 96 | 74 | 69 | 167 |
| Gastric cancer | 48 | 48 | 0 | 48 |
| Lung squamous cell carcinoma | 34 | 19 | 17 | 69 |
| Lung adenocarcinoma | 33 | 23 | 11 | 59 |
| Hepatocellular carcinoma | 28 | 3 | 5 | 34 |
| Skin melanoma and other skin cancers | 28 | 13 | 17 | 79 |
| Head and neck cancer | 22 | 5 | 22 | 106 |
| Sarcoma | 19 | 7 | 15 | 73 |
| Uninary Bladder cancer | 16 | 6 | 15 | 61 |
| Other lung cancers | 11 | 7 | 5 | 16 |
| Brain glioblastoma multiforme | 10 | 9 | 2 | 13 |
| Ovarian cancer and other female reproducitve organ cancers | 10 | 1 | 10 | 31 |
| Pancreatic cancer | 10 | 6 | 5 | 27 |
| Renal cell carcinoma and other kidney cancers | 8 | 2 | 7 | 29 |
| Other cancers | 6 | 1 | 5 | 16 |
| **Total** | **509** | **324** | **328** | **1127** |

Supplementary Table 2: Summary of datasets by tumor type.

| Comparison dataset | | Data source | Number of PT samples | Number of PDX samples |
|---|---|---|---|---|
| SNP vs WES | Comparison of CNA profiles estiimated from SNP array and whole-exome sequencing | JAX PDX resource and HCI breast cancer | 8 | 0 |
| WES vs RNASEQ (NORM/TUM) | Comparison of CNA profiles estimated from whole-exome sequencing and RNA-sequencing, either normalized by median expression of normal samples of the same tumor type or median expression of the same set of tumor samples | SNU-JAX gastric cancer | 27 | 27 |
| SNP vs EXPARR (NORM/TUM) | Comparison of CNA profiles estimated from SNP array and gene expression array, either normalized by median expression of normal samples of the same tumor type or median expression of the same set of tumor samples | SIBS hepatocellular carcinoma | 2 | 33 |
| RNASEQ NORM vs TUM | Comparison of CNA profiles estimated from RNA-sequencing with different normalizations, by median expression of normal samples of the same tumor type or median expression of the same set of tumor samples | SNU-JAX gastric cancer | 42 | 42 |
| EXPARR NORM vs TUM | Comparison of CNA profiles estimated from gene expression array with different normalizations, by median expression of normal samples of the same tumor type or median expression of the same set of tumor samples | SIBS hepatocellular carcinoma | 3 | 34 |

Supplementary Table 3: Benchmarking dataset which comprises copy number alteration profiles estimated for matched samples assayed across multiple platforms

| Genes with >5% recurrent frequency (\|residual\| > 1) | Gene location (GRCh38) | Recurrent frequency (%) | |
|---|---|---|---|
| | | PT vs PDX | PDX vs PDX |
| GOLGA6L6 | 15q11.2 | 6.09 | |
| HLA-DQA1 | 6p21.32 | 8.24 | |
| HLA-DQB1 | 6p21.32 | 7.89 | |
| HLA-DRB1 | 6p21.32 | 8.96 | 6.54 |
| HLA-DRB5 | 6p21.32 | 7.53 | 5.23 |
| MACROD2 | 20p12.1 | 6.81 | |
| OR4M2 | 15q11.2 | 6.81 | |
| OR4N4 | 15q11.2 | 6.81 | |
| POTEB | 15q11.2 | 6.45 | |
| POTEB2 | 15q11.2 | 6.81 | |
| RBFOX1 | 16p13.3 | 5.73 | |
| TPTE | 21p11.2 | 7.17 | |

Supplementary Table 4: Recurrent frequency (based on models) of genes with >5% recurrence with large copy number deviation (|residual| > 1) from linear regression model for PT-PDX (279 models) and PDX-PDX (306 models) comparisons.
